# Chemigenetic far-red labels and Ca^2+^ indicators optimized for photoacoustic imaging

**DOI:** 10.1101/2024.05.23.595278

**Authors:** Alexander Cook, Nikita Kaydanov, Begoña Ugarte-Uribe, Juan Carlos Boffi, Gretel B. Kamm, Robert Prevedel, Claire Deo

## Abstract

Photoacoustic imaging is an emerging modality with significant promise for biomedical applications such as neuroimaging, owing to its capability to capture large fields of view, deep inside complex scattering tissue. However, the widespread adoption of this technique has been hindered by a lack of suitable molecular reporters for this modality. In this work, we introduce chemigenetic labels and calcium sensors specifically tailored for photoacoustic imaging, using a combination of synthetic dyes and HaloTag-based self-labelling proteins. We rationally design and engineer far-red “acoustogenic” dyes, showing high photoacoustic turn-ons upon binding to HaloTag, and develop a suite of tunable calcium indicators based on these scaffolds. These first-generation photoacoustic reporters show excellent performance in tissue-mimicking phantoms, with the best variants outperforming existing sensors in terms of signal intensity, sensitivity and photostability. We demonstrate the application of these ligands for labelling HaloTag-expressing neurons in mouse brain tissue, producing strong, specifically targeted photoacoustic signal, and provide a first example of *in vivo* labelling with these chemigenetic photoacoustic probes. Together, this work establishes a new approach for the design of photoacoustic reporters, paving the way towards deep tissue functional imaging.

## Introduction

The understanding of neural encoding and information processing, spanning spatial scales from individual neurons to entire brain regions, is a fundamental challenge in neuroscience. A variety of direct and indirect imaging methods can be used to monitor neural activity across these scales, from macroscopic techniques such as MRI hemodynamic contrast,^1^ to single cell fluorescence microscopy.^2^ Among these, a prevalent approach is the monitoring of fluctuations in neuronal calcium concentration, using indicators that exhibit fluorescence changes upon binding to calcium ions.^3^ Calcium imaging began with the development of small-molecule BAPTA-based probes by Roger Tsien,^4^ with this field undergoing a revolutionary shift with the introduction of genetically encoded calcium indicators (GECIs), now routinely used for complex experiments in living animals.^5-7^ Nevertheless, despite continuous improvements, fluorescence imaging is inherently limited to superficial tissue layers due to light scattering, which restricts imaging depth to a couple of millimeters at best.^8^ In contrast, photoacoustic imaging (PAI), which relies on light absorption and subsequent ultrasound emission due to the non-radiative deexcitation of chromophores, can achieve centimeter-deep imaging of large fields of view (up to ∼15×15x10 mm) at ∼50-100 µm resolution, hence standing as an ideal modality for mesoscopic-scale neuroimaging of large volumes such as the entire mouse brain.^9, 10^ However, despite the availability of highly performant fluorescent calcium indicators, there are to date no effective photoacoustic Ca^2+^ biosensors, which has hindered the widespread adoption of PAI in the neuroscience field. This can be explained by the stringent requirements for such reporters, including a high extinction coefficient (ε) in the far-red/near-infrared (NIR) to minimize background from endogenous chromophores such as hemoglobin, a low fluorescence quantum yield (Φ) to maximize radiationless relaxation and thus photoacoustic signal, high photostability, and the possibility to genetically target them to specific cells or subcellular features.^11^ Moreover, essential requirements including high sensitivity, selectivity, physiologically-relevant binding affinity, and fast kinetics need to be fulfilled by effective biosensors. The design principles of photoacoustic Ca^2+^ indicators have mirrored those of their fluorescent analogues, built from either synthetic or genetically encoded chromophores. However, given the fundamentals of the photoacoustic effect,^12^ sensors relying on absorption changes inherently possess the potential for markedly higher sensitivity than quantum yield-modulated sensors.^13^ This renders mechanisms such as photoinduced electron transfer and Förster resonance energy transfer less promising for the development of photoacoustic reporters. To date, only a limited number of calcium sensors for PAI have been reported, and suffer from important limitations. Current small-molecule calcium indicators lack genetic targetability, are generally difficult to deliver in complex biological systems, and exhibit modest sensitivity.^14, 15^ Owing to their large increase in absorption upon Ca^2+^ binding, fluorescent GECIs such as GCaMPs and photoswitchable analogues have been shown to provide a photoacoustic response to calcium, but their high fluorescence quantum yields and blue excitation wavelengths limit their applicability to superficial brain regions.^16-18^ Sensitive far-red GECIs have been notoriously difficult to engineer, and the few reported ones have not yet been used in PAI,^19-21^ likely due to low sensitivity and/or insufficient photostability, which highlights the need for alternative approaches towards PA calcium sensors.

In this work, we report the first generation of chemigenetic reporters for photoacoustic imaging. This approach, which amalgamates small-molecule dyes with self-labelling proteins such as HaloTag,^22, 23^ was recently established as a platform for the design of multicolor fluorescent Ca^2+^ biosensors.^24, 25^ Recognizing the potential of this approach for PAI, we rationally engineered a series of far-red/ NIR ‘acoustogenic’ ligands, showing high turn-ons in photoacoustic signal upon binding to HaloTag (Figure 1). Adapting these ligands to calcium-sensitive HaloTag-based proteins, we developed a suite of calcium sensors with excellent performance in tissue-mimicking phantoms. Importantly, our novel acoustogenic ligands can label HaloTag-expressing neurons in mouse brain tissue, providing high and specific PA contrast.

**Figure 1.**
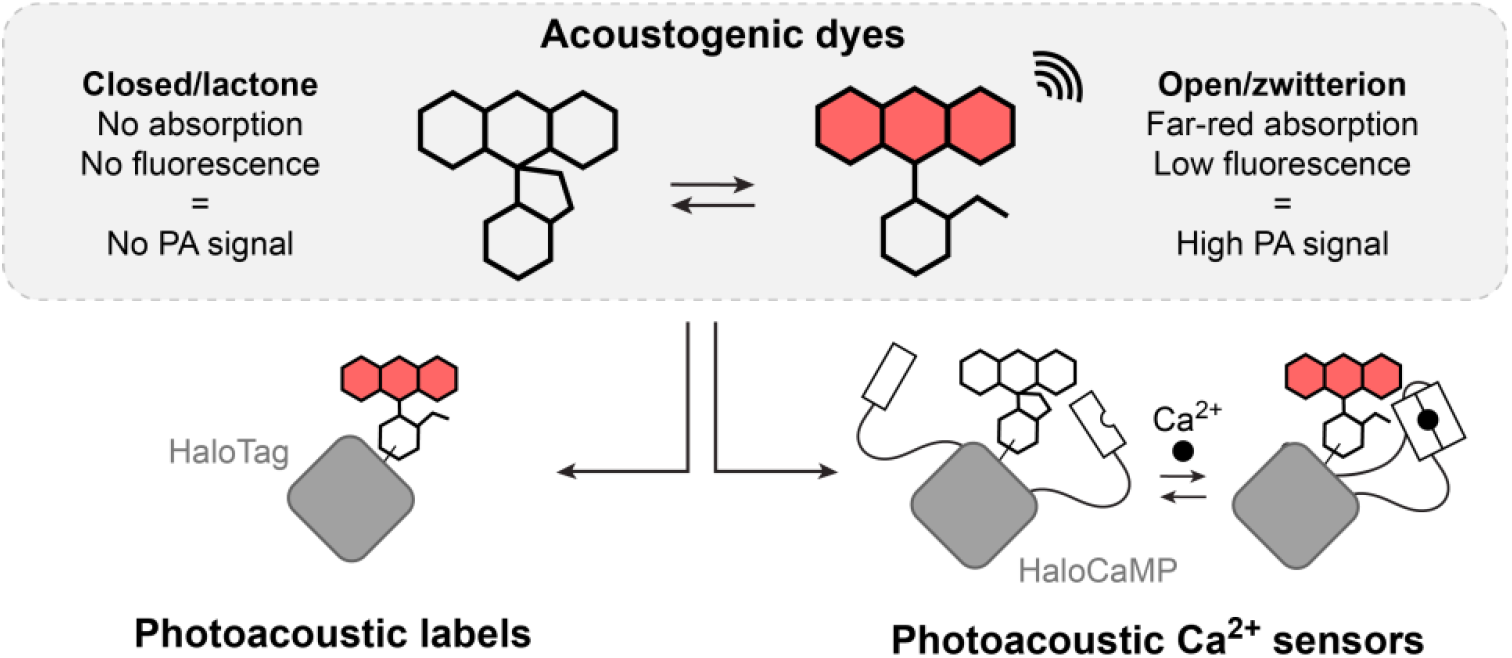
General approach for the design of chemigenetic photoacoustic labels and calcium sensors based on the open-closed equilibrium of acoustogenic dyes.

## Results

To develop efficient labels and calcium sensors for PAI, we sought to adapt the recently established “chemigenetic” approach, initially designed for fluorescent reporters.^26, 27^ The most common strategy uses the self-labelling protein HaloTag, with rhodamine-based ligands, which exist in equilibrium between a non-absorbing lactone form and a highly absorbing and fluorescent zwitterionic form. When preferentially adopting the closed form in solution, dyes can exhibit fluorogenic behavior, i.e. a substantial fluorescence turn-on upon binding to the HaloTag protein, due to a shift in the equilibrium towards the open form. Together, this platform has provided bright multicolor fluorescent labels suitable for *in vivo* imaging.^28-30^ Importantly, the HaloTag protein can be modified and appended with genetically encoded sensing motifs, yielding analyte-responsive self-labelling tags. In these tags, the conformational change of the protein upon binding to its target leads to a change in absorption and emission of the fluorogenic dye ligand.^24, 25, 31^ We reasoned that meticulous engineering of far-red/NIR absorbing and non-fluorescent dyes, featuring a similar open-closed equilibrium, would lead to “acoustogenic” HaloTag ligands. These dyes, while maintaining low fluorescence, would show large turn-ons in photoacoustic signal upon binding to the HaloTag protein, and could in turn function as acoustic reporters in HaloTag-based calcium sensors.

### Design, synthesis and properties of acoustogenic scaffolds

To design highly acoustogenic ligands, we investigated different families of dyes, focusing on scaffolds combining a far-red/NIR absorption maximum with a minimal Φ, and bearing the paradigmatic *o*-carboxylic acid group on the pendant phenyl ring responsible for the open-closed equilibrium in rhodamine derivatives (Figure 2, Figure S1).

**Figure 2.**
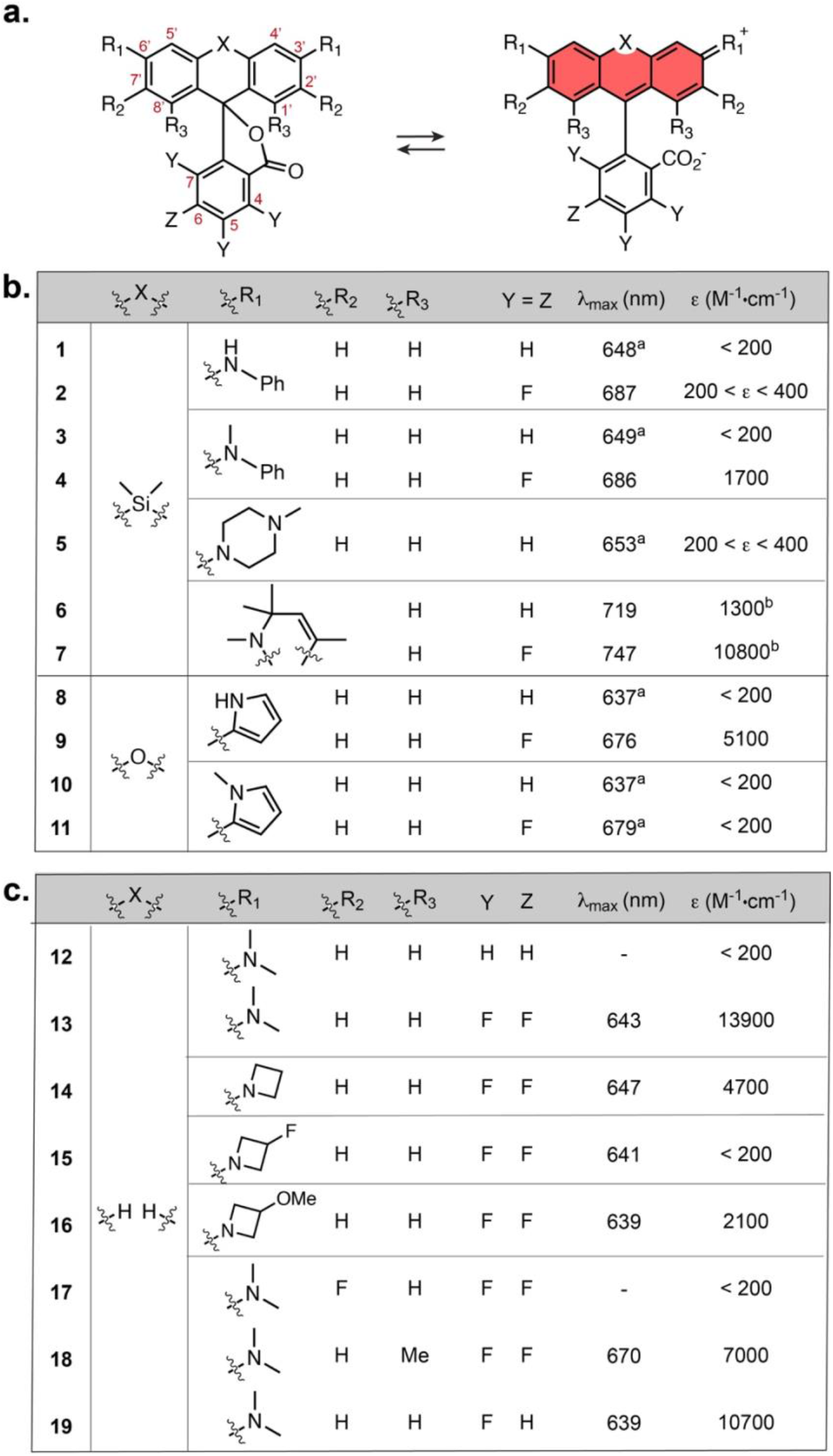
(**a**) General structure and open-closed equilibrium of the dyes studied in this work; Absorption properties of (**b**) Si-rhodamine and pyrrole-xanthene derivatives and (**c**) Malachite Green lactone derivatives; All measurements were made in 10 mM HEPES, pH = 7.4 except for ^a^in 0.1% 2,2,2-trifluoroethanol in trifluoroacetic acid. ^b^apparent value due to aggregation. Values are the mean of 3 replicates.

As analogous far-red scaffolds are often predominantly closed, we systematically examined both H- and F-substitution on the pendant phenyl ring, as fluorination at these positions leads to a shift of the equilibrium towards the open form.^30^ The absorption properties of the free dyes were measured in aqueous buffer (Figure 2b,c, Figure S2), and in MeCN/H_2_O mixtures (Figure S3), to characterize their open-closed equilibrium and therefore assess their potential acoustogenicity.

First, we examined well-established far-red Si-rhodamines, bearing *N*-aryl or *N*-methylpiperazine auxochromes to minimize fluorescence (compounds **1-5**).^32-34^ We also examined recently reported dihydroquinoline-fused Si-rhodamines, which display NIR absorption and low Φ (compounds **6, 7**).^34^ Compounds **1-7** were synthesized using reported methods (Synthetic Schemes in SI).^34, 35^ Among them, compounds **1** and **3** showed no detectable absorption (*ε* < 200 M^-1^·cm^-1^), supporting that these are too closed to show exploitable chromogenicity. In contrast, compounds **2, 4, 5, 6** displayed a weak but detectable absorption band (200 < *ε* < 2000 M^-1^·cm^-1^), which is indicative of an absorption turn-on of the corresponding HaloTag ligands upon binding to the protein.^30^ Compound **7** showed a higher extinction coefficient, and measurements in MeCN/H_2_O mixtures revealed that compounds **6** and **7** have a high propensity for aggregation in water, with **7** presenting *ε* = 98000 M^-1^·cm^-1^ in 40% MeCN/H_2_O, suggesting that the dye is fully open but strongly aggregates in aqueous buffer (Figure S3).

Next, we investigated the behavior of pyrrole-functionalised O-xanthenes, a recently reported class of NIR dyes,^36^ and synthesized compounds **8-11** by Suzuki coupling from the fluorescein ditriflates. These dyes presented absorption maxima between 637 and 679 nm (a 90-110 nm red-shift compared to their rhodamine relatives), due to an extended electronic conjugation. **8, 10** and **11** were strongly shifted towards the closed form, but tetrafluorinated **9** showed a higher *ε* = 5100 M^-1^·cm^-1^. This value is far lower than the expected maximum for these scaffolds (∼10^5^ M^-1^·cm^-1^), characteristic of possible chromogenic behavior.

Finally, we examined triarylmethane lactones (compounds **12**-**19**), closely resembling Malachite Green, which absorbs in the far-red and is virtually non-fluorescent in solution due to rotational flexibility.^37, 38^ The parent nonfluorinated (**12**) and the tetrafluorinated (**13**) Malachite Green lactones were synthesised in one step by Friedel-Crafts acylation of *N,N*-dimethylaniline with the corresponding phthalic anhydride. While **12** was too closed, compound **13** with *ε* = 13900 M^-1^·cm^-1^, was structurally reminiscent of the landmark fluorogenic dye SiR,^39^ and had substantial room for further functionalization. We hypothesized that strategies to shift the dye towards the closed form which have been developed for rhodamines could be applied to this scaffold, in order to decrease the absorption of the free dye, and in turn lead to higher chromogenicity and acoustogenicity. Following a well-established approach, we set out to replace the *N,N*-dimethyl substituents with various substituted azetidines (**14, 15, 16**).^28^ The reactivity of the azetidines prevented synthesis of these target compounds by Friedel-Crafts acylation or Pd-catalyzed cross coupling, and they were synthesized using a recently reported synthetic method involving lithiation of 2,3,4,5-tetrafluorobenzoic acid before addition to the corresponding benzophenones.^34^ The trend observed was very similar as for SiR, with electron withdrawing substituents shifting the equilibrium towards the closed form. Among these azetidine-substituted derivatives, compounds **14** and **16** displayed extinction coefficients in the desired range (4700 M^-1^·cm^-1^ and 2100 M^-1^·cm^-1^, respectively). As a second approach to tune the equilibrium, we examined the effect of substituents on the aromatic rings, introducing fluorines at the 2’,7’ positions (**17**), previously shown to shift the equilibrium towards the closed form.^40^ Unfortunately the fluorination had too strong an effect, completely closing the dye. We also introduced methyl groups on the 1’,8’ positions (**18**), a previously unexplored modification to alter the equilibrium of dyes. Interestingly, this resulted in a small shift towards the closed form, with a two-fold lower extinction coefficient for **18** compared to **13**, along with a favorable ∼30-nm red-shift in absorption. Finally, we also investigated the impact of partial defluorination of the bottom ring with compound **19**, serendipitously isolated as a by-product during the synthesis of the HaloTag ligand. As expected, this previously unexplored modification provided a desired small shift of the equilibrium towards the closed form (*ε* = 10700 M^-1^·cm^-1^).

### Acoustogenic HaloTag ligands as photoacoustic labels

Based on the absorption properties of the free dyes, we synthesized and characterized the HaloTag ligands of selected compounds in each family (presenting 200 < *ε* < 15000 M^-1^·cm^-1^, Figure S4). These were synthesized by standard amide coupling from the 6-CO_2_H precursor for non-fluorinated ligands, and directly from the free dyes for the fluorinated compounds, using an adapted version of the MAC chemistry.^41^ We measured their photophysical properties in the absence or presence of HaloTag protein (Figure 3). As expected, all ligands showed an absorption increase upon binding to HaloTag7, albeit to different extents (Figure 3b,c, Figure S5, Table S1). Si-rhodamine **4-HTL** showed a high turn-on upon binding to HaloTag, with ΔA/A_0_ = 62. This compound however showed very slow binding (> 24 hours in solution), which is prohibitive for biological applications (Figure S6). We hypothesized that this was primarily due to unfavourable interactions of the phenyl rings on the protein surface, so attempting to minimize this deleterious effect, we synthesized the unsymmetrical compound **20-HTL**, by replacing one aniline with an azetidine (Figure S4). This compound maintained high absorption turn-on upon binding (ΔA/A_0_ = 42) and low Φ < 0.01, and showed substantially faster binding to the protein than the symmetrical **4-HTL**, although still slower than conventional HaloTag ligands (Figure S6). Piperazine-SiR **5-HTL** was too closed, with low absorption when bound to HaloTag, and additionally presented a relatively high protein-bound Φ = 0.11, so was therefore deemed unsuitable for PA. Compound **6-HTL** showed fast binding kinetics and a high ΔA/A_0_ of 94, but similarly a high Φ = 0.13 with HaloTag. In the pyrrole-xanthene family, fluorinated **9-HTL** showed a good ΔA/A_0_ = 9.2. Finally, we investigated the behavior of the Malachite Green-derived HaloTag ligands. To further expand the series of fine-tuned derivatives, we additionally synthesized the hydroxy-substituted **21-HTL**, hypothesized to present improved water solubility. We also synthesized the difluoro derivative **22-HTL**, by hydrodefluorination of the ester intermediate. We note that this approach had not been previously explored to fine-tune dyes, and the regioselectivity of the reaction was confirmed by X-Ray crystallography, showing removal of the fluorine at the 4-position in the major isolated product (see Crystallography in SI). All the Malachite Green derivatives showed rapid binding kinetics, with good ΔA/A_0_ up to 21 and generally followed the expected trend, with dyes more strongly shifted towards the closed form showing higher ΔA/A_0_, but lower protein-bound absorption. Importantly, these compounds remained virtually non-fluorescent after binding (Φ < 0.01), demonstrating that HaloTag does not elicit the conformational restriction observed with fluorogen activating proteins for Malachite Green which results in a fluorescence turn-on undesirable for PAI.^42^ In an attempt to reduce the trade-off in HaloTag-bound absorption, we also evaluated selected ligands with HaloTag9, a recently published mutant which shows higher absorption turn-on with certain rhodamine derivatives (Table S1, Figure S5).^43^ Although we observed similar absorption as with HaloTag7 for the HTLs of **14, 16** and **22**, the absorption of the bound dye to HaloTag9 was up to 70% higher for compounds **9-HTL, 13-HTL** and **18-HTL**, showing that our ligands have potential for even greater turn-ons with tailored protein engineering. Overall, six of the ligands **9-HTL, 13-HTL, 14-HTL, 16-HTL, 18-HTL, 22-HTL** showed fast labelling kinetics, large absorption turn-on, high far-red/NIR absorption and low Φ upon binding to HaloTag, suggesting excellent performance as photoacoustic labels.

**Figure 3.**
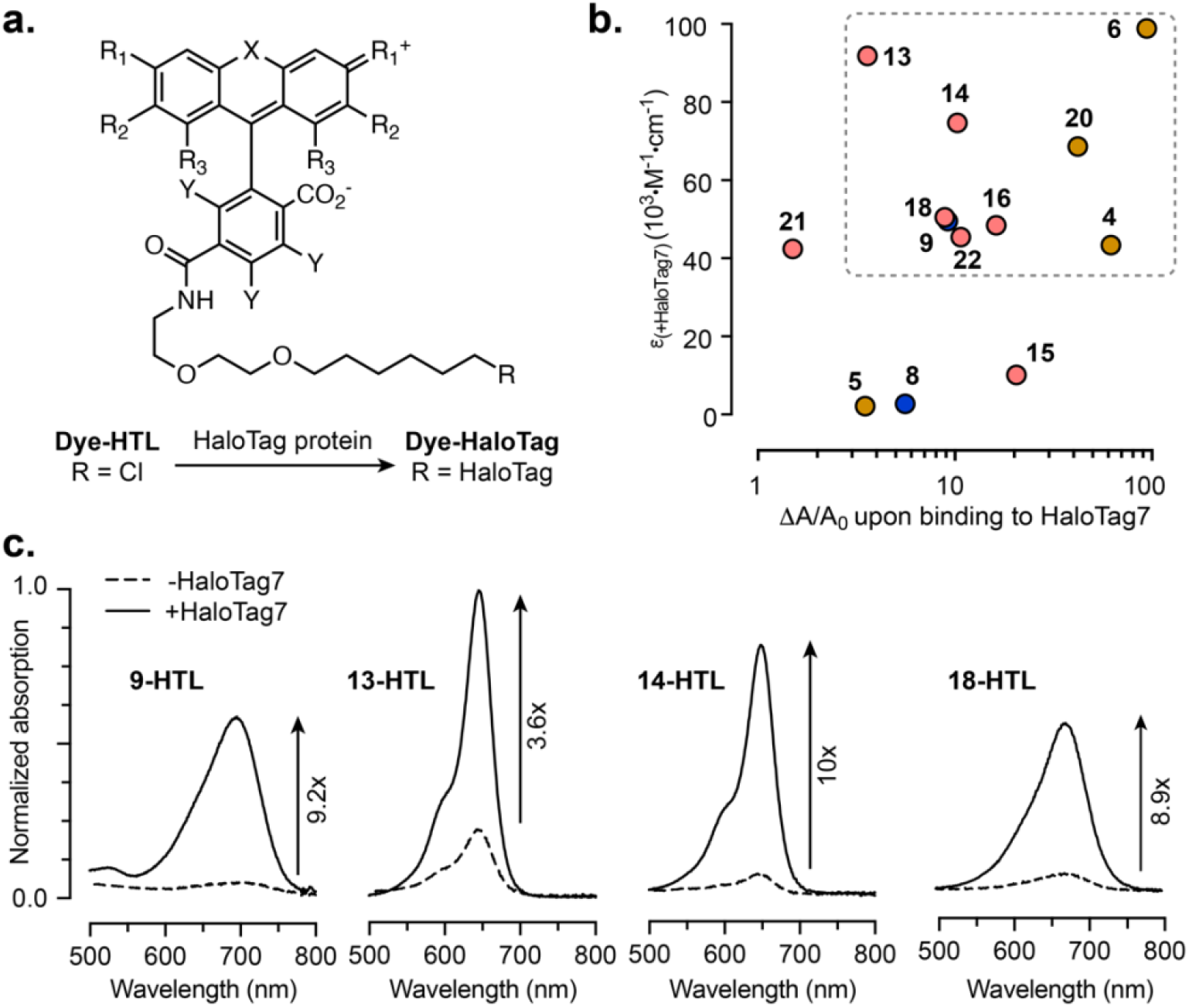
(**a**) General structure of HaloTag ligands and formation of the dye-HaloTag conjugate; (**b**) Extinction coefficient of the HaloTag-bound dye vs. absorption turn-on upon binding for the HaloTag ligands. “HTL” was omitted from the labels for clarity. Yellow: Si-rhodamines, blue: pyrrole-xanthenes, red: Malachite Green derivatives. The dashed box highlights compounds presenting both large absorption turn-ons and high extinction coefficients bound to HaloTag; Values are the mean of 3 replicates; (**c**) Normalized absorption spectra of selected HaloTag ligands in the absence or presence of HaloTag protein.

We therefore measured their photoacoustic properties, using a custom-built spectrometer (Figure S7). Specifically, the spectral features and photoacoustic turn-ons upon HaloTag binding (ΔPA/PA_0_) at λ_PA_ = λ_max_, were in excellent agreement with absorption measurements, confirming that absorption can serve as a robust proxy for the design of such photoacoustic reporters (Figure S8). To support the proof-of-principle of our approach, we compared the photoacoustic signal intensity of **13-HTL** with the spectrally-matched highly fluorescent **JF635-HTL**^28^ (Figure S9). While both HaloTag conjugates have comparable extinction coefficients, the photoacoustic signal was 17-fold larger for **13**-HaloTag7, explained by the fact that its fluorescence quantum yield (Φ ∼ 0.001) is about 750-fold smaller than for **JF635**-HaloTag7 (Φ = 0.75). This shows that strong suppression of fluorescence is a paramount requirement for high photoacoustic signal in this type of chromophore.

### Design and properties of photoacoustic calcium sensors

Given the high performance of our acoustogenic HaloTag ligands *in vitro*, we then set out to use them for the design of Ca^2+^ sensors. For this purpose, we focused on the Ca^2+^-sensitive self-labelling proteins HaloCaMP and rHCaMP.^24, 25^ In these sensors, the conformational change of the protein upon binding calcium ions alters the local environment of a HaloTag-ligand fluorophore, hereby shifting its equilibrium towards the open form with concomitant absorption increase. We reasoned that these calcium-binding self-labelling proteins would lead to sensitive photoacoustic sensors when used with our acoustogenic ligands. Indeed, all novel ligands tested showed a calcium-dependent change in absorption and photoacoustic signal when used with these platforms (Table S2, Figure S10, Figure S11). Generally, the two HaloCaMP variants, 1a and 1b, afforded positive-going sensors, while rHCaMP led to negative-going sensors with the same dyes. Importantly, absorption and photoacoustic properties were again in excellent agreement for the calcium sensors (Figure 4a, Figure S12).

**Figure 4.**
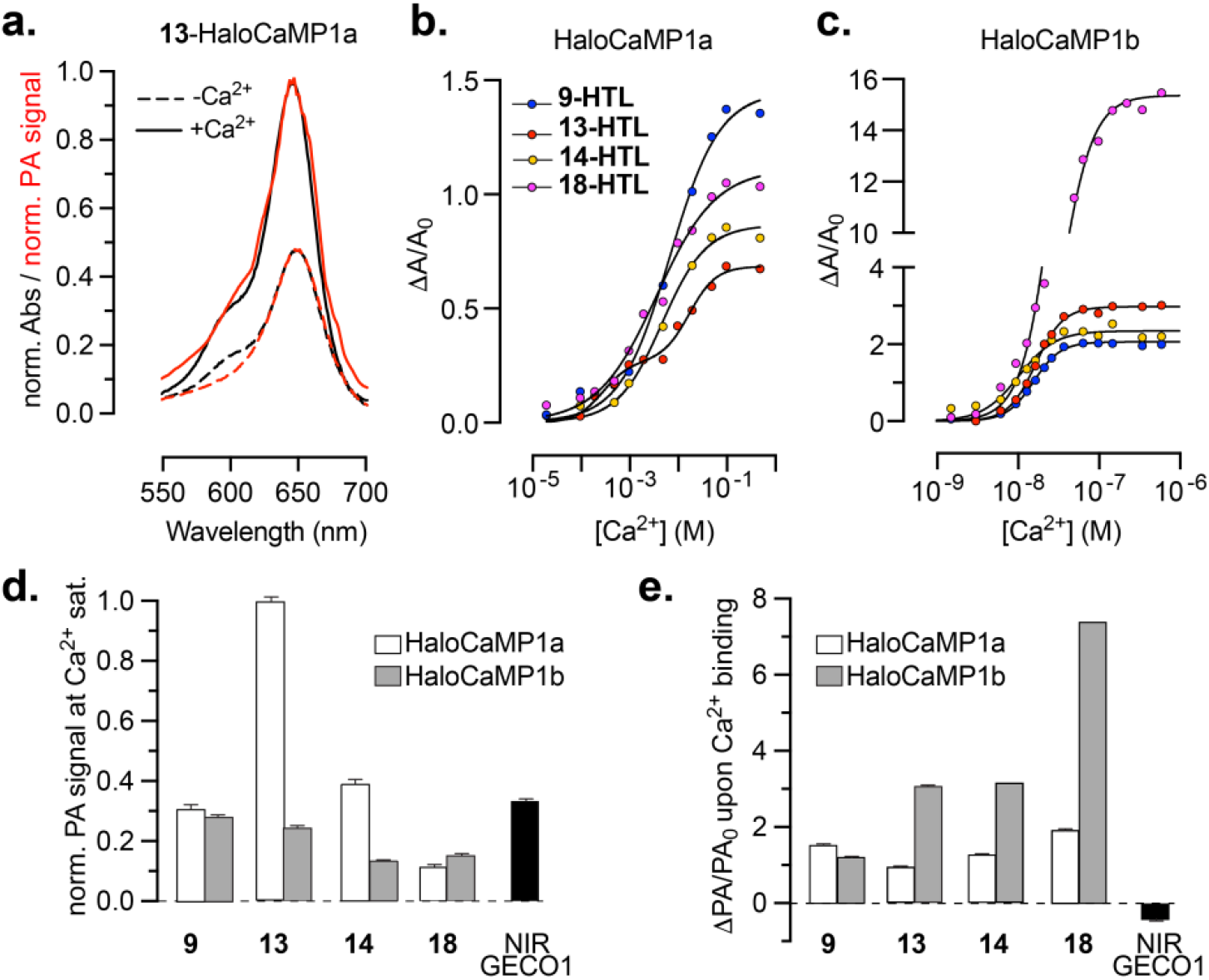
(**a**) Normalized absorption (black) and photoacoustic (red) spectra of **13-HTL** bound to HaloCaMP1a in the Ca^2+^-free (dashed lines) and Ca^2+^-bound (solid lines) states; Ca^2+^ titrations of (**b**) HaloCaMP1a and (**c**) HaloCaMP1b labelled with **9-HTL, 13-HTL, 14-HTL** or **18-HTL**; (**d**) Normalized PA signal in the Ca^2+^-bound state; (**e**) ΔPA/PA_0_ upon binding calcium, measured at λ_PA_ = λ_max_ for the same sensors compared to NIR-GECO1; (**d**,**e**) show mean and SEM for 3 replicates. “HTL” was omitted from the labels in (**d**,**e**) for clarity.

Compound **9-HTL** and Malachite Green derivatives **13-HTL, 14-HTL** and **18-HTL** showed comparable sensitivity with HaloCaMP1a (ΔA/A_0_ between 0.7 and 1.4, ΔPA/PA_0_ between 1.1 and 2.0), and larger sensitivity with HaloCaMP1b, with ΔPA/PA_0_ up to 7.5. Following the trend observed with HaloTag, compounds **16-HTL** and **22-HTL** showed low absorption and photoacoustic signal in the Ca^2+^-bound state, and were therefore excluded.

Overall, the results align with the observations made with the Janelia Fluor ligands which HaloCaMP was originally evolved against, with variant 1a generally resulting in higher absorption, and variant 1b leading to larger ΔA/A_0_ with the same dye.^24^ With ligand **13-HTL**, both HaloCaMP variants showed low pH sensitivity and high selectivity for Ca^2+^ over Mg^2+^ (Figure S13, Figure S14). We then performed calcium titrations to determine the calcium affinity of these sensors (Figure 4b,c, Table S2). The four ligands tested provide high affinity sensors with HaloCaMP1b with K_d_ ranging from 10 to 33 nM, which is in the same range as the original HaloCaMP1b sensors,^24^ and GECIs such as jGCaMP7s,^44^ thus well-suited for *in vivo* imaging of cytosolic neuronal calcium.

Surprisingly, using the same dyes with the HaloCaMP1a scaffold led to low affinity sensors, with K_d_ orders of magnitude larger than with the original JF derivatives (∼60-900 nM).^24^ **13**-HaloCaMP1a and **18**-HaloCaMP1a clearly showed two binding phases (K_d_1 = 300 µM and 980 µM, K_d_2 = 18 mM and 10 mM, respectively), while **9**-HaloCaMP1a and **14**-HaloCaMP1a presented an apparent single binding event, with a K_d_ of 6.9 mM and 4.8 mM, respectively. The mechanism by which the dye ligand determines the apparent affinity of the sensor is unclear.

Nevertheless, millimolar affinity sensors can prove useful for monitoring extracellular calcium dynamics,^45^ or visualizing calcium flux inside subcellular compartments such as the endoplasmic reticulum.^46-48^ Overall, **13-** HaloCaMP1a provided the largest photoacoustic signal in the Ca^2+^-bound state (Figure 4d), and **18-**HaloCaMP1b showed the highest sensitivity, with ΔPA/PA_0_ = 7.5 (Figure 4e). Our probes are also far more photostable than competing protein sensors like the far-red, biliverdin-dependent, fluorescent GECI NIR-GECO1^19^ (Figure S15). After illumination for 1 h at λ_max_, the photoacoustic signal of NIR-GECO1 decreased to 65%, while **13**-HaloCaMP1a and **13**-HaloCaMP1b retained 90% to 95% of their original signal. The parent chromophores show a similar trend, with **13**-HaloTag7 substantially more photostable than mIFP (Figure S15a,c), and the relative photostabilities between mIFP and NIR-GECO1 are consistent with reported values.^19, 49^ Overall, ligands **9, 13, 14** and **18** used in combination with HaloCaMP variants provide a series of eight photoacoustic Ca^2+^-indicators with a wide range of affinities, signal intensities and sensitivities. Importantly, these positive-going sensors clearly outperform existing sensors such as NIR-GECO1 and published synthetic PA calcium sensors in sensitivity (Figure 4e, Table S2).

### Photoacoustic tomography in tissue-mimicking phantoms

We then set out to characterize and evaluate our new acoustogenic probes in tissue-mimicking phantoms, using a custom-built all-optical Fabry-Pérot-based photoacoustic tomography set-up, which is ideally suited for deep-tissue imaging in mice.^50, 51^ The phantoms were prepared by immobilizing 1 mm diameter hollow polyethylene tubes in coupling medium composed of either pure H_2_O (non-scattering medium), or 60% v/v milk/H_2_O, which mimics the scattering of the living mouse brain (Figure 5a).^52-54^ The tubes were filled with solutions of purified dye-protein conjugates, and positioned at the desired depth from the Fabry-Pérot detector. We evaluated the performance of our HaloTag-based photoacoustic labels and compared it against mIFP, the chromophore component of NIR-GECO1.^19^ Ligand **13** bound to HaloTag7 gave ∼6-fold higher PA signal than mIFP (Figure S16), with detectable signal down to 1.2-cm depth in 60% milk phantom, much deeper than the mouse brain (∼ 6 mm) (Figure S17). In contrast, mIFP gave a much lower signal and was not detectable below 8 mm in these scattering conditions. At a depth of 5 mm, equivalent to close to the bottom of the mouse brain, a solution of 5 μM **13**-HaloTag7 gave a signal-to-background ratio of 2, suggesting a detection limit of 2.5 μM in those brain-scattering representative conditions (Figure 5b, Figure S18), which is well within the protein concentration range achievable *in vivo* with adeno-associated viruses.^55, 56^ Notably, the HaloCaMP-based calcium sensors led to similar turn-ons in the phantoms as in spectroscopy, outperforming NIR-GECO1 which only showed a small negative response (Figure 5c, Figure S19). In the Ca^2+^-bound state, **13**-HaloCaMP1a gave twice the photoacoustic signal of NIR-GECO1 and **13**-HaloCaMP1b, in scattering medium at ∼5 mm depth (Figure S20).

**Figure 5.**
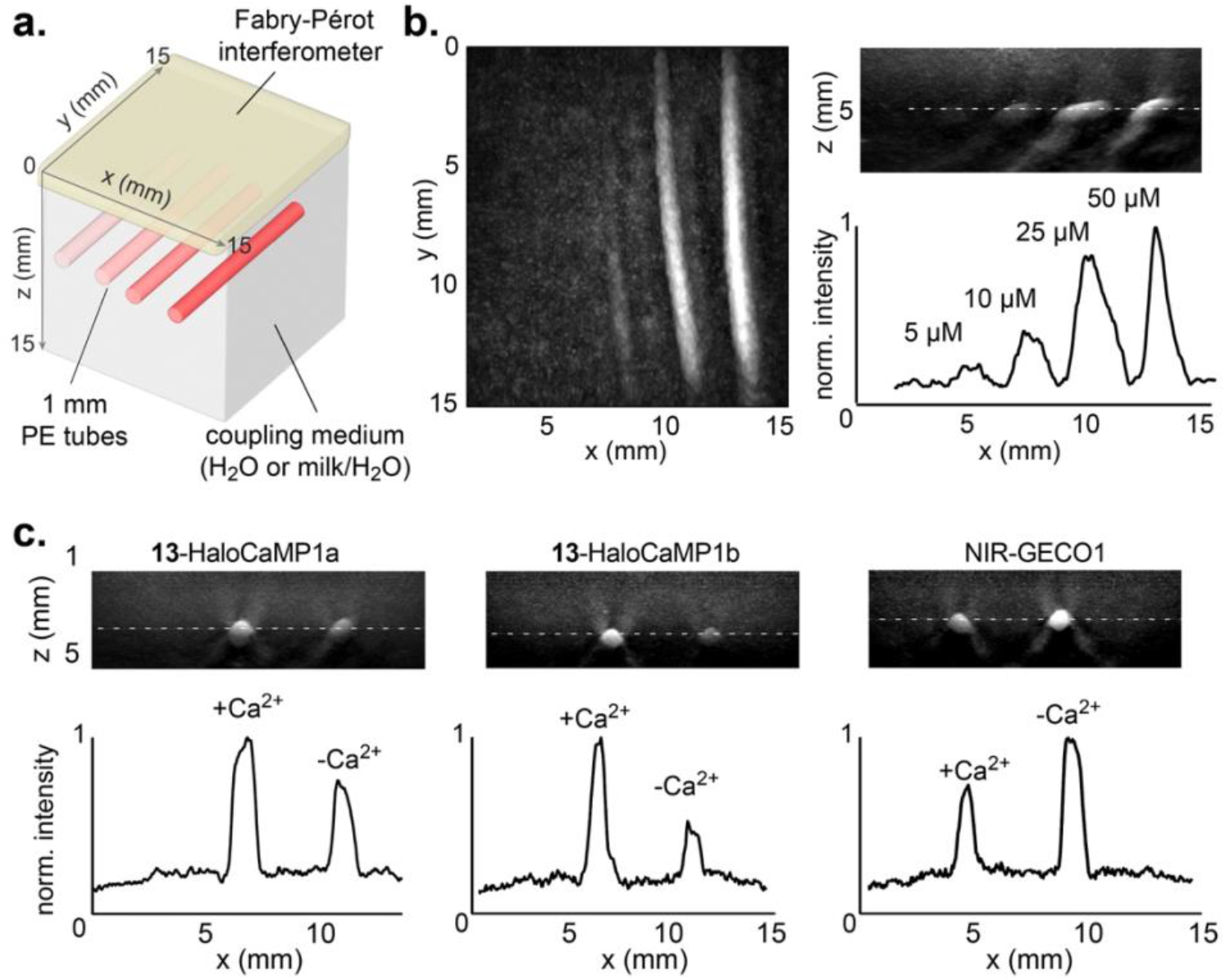
(**a**) Schematic representation of the tissue-mimicking phantom set-up showing relative position of the polyethylene (PE) tubes and the Fabry-Pérot interferometer; (**b**) PAT maximum intensity projections and line profile (250 µm thick) quantification of **13**-HaloTag at 5, 10, 25 or 50 µM, at ∼5 mm depth in 60% milk/H_2_O scattering medium; (**c**) PAT maximum intensity projections and quantification of **13**-HaloCaMP1a, **13**-HaloCaMP1b or NIR-GECO1 in the calcium-bound and the calcium-free states at 50 µM, at ∼3.5 mm depth in 60% milk/H_2_O scattering medium; PAT was performed at λ_PA_ = λ_max_ for each probe.

### Labelling and photoacoustic tomography of mouse brain tissue

Consequently, we set out to demonstrate the use of the newly developed reporters in biological systems. We first assessed the cell permeability and labelling efficiency of the ligands in U2OS cells expressing HaloTag7-EGFP (Figure S21). Very faint fluorescence signal from the dyes could be observed in the far-red channel. The bright competitor **JF549-HTL** was subsequently added to these pre-labelled cells,^28^ and the absence of fluorescence signal in this channel confirmed that the ligands tested **9-HTL, 13-HTL** and **14-HTL** efficiently bind to HaloTag in living cells. Moving to more complex tissues, we then tried to label mouse brain slices for photoacoustic imaging. For this purpose, mice were injected with an AAV1 HaloTag7-EGFP under the synapsin promoter into the hippocampus in the righthand hemisphere of the brain. Following expression, brains were excised, sliced coronally and labelled with **13-HTL** by bath loading. Successful specific labelling of HaloTag-expressing neurons in the hippocampus was observed via widefield fluorescence microscopy, as shown by the correlation of the EGFP fluorescence and the weak far-red fluorescence from **13-HTL**, which was absent prior to labelling (Figure 6, Figure S22). Photoacoustic tomography showed strong specific photoacoustic signal from the labelled hippocampus. In contrast, brain slices lacking HaloTag7 expression showed no detectable fluorescence or photoacoustic signal from **13-HTL** after incubation with the dye, demonstrating specificity of the labelling (Figure 6b).

**Figure 6.**
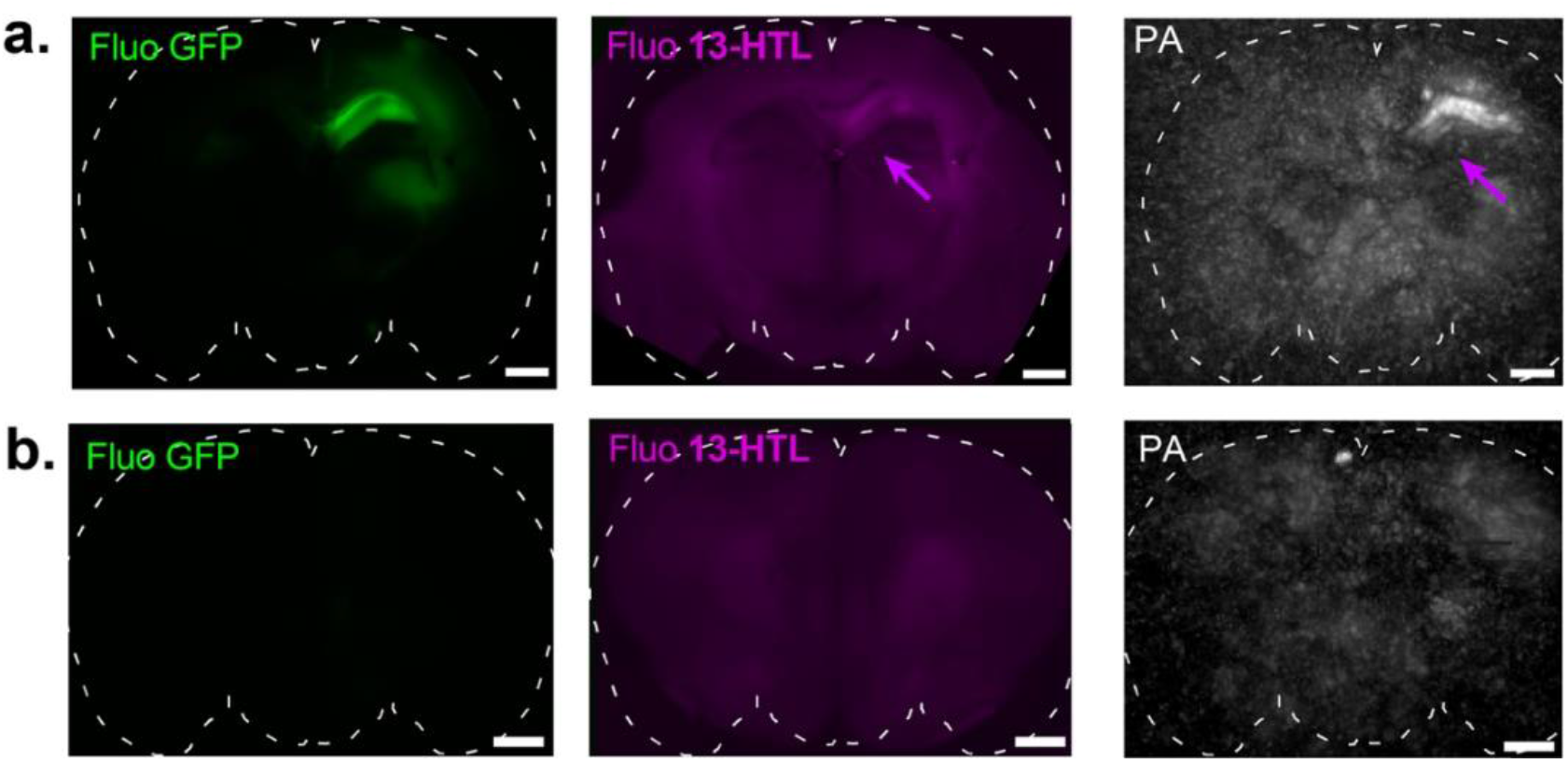
Fluorescence microscopy and photoacoustic tomography of mouse coronal brain slices labelled *ex vivo* with **13-HTL**; (**a**) Slice with high expression of HaloTag-EGFP in the hippocampus and (**b**) slice with no detectable expression of HaloTag-EGFP. Fluorescence images of EGFP (left panels) and**13-HTL** (middle panels), photoacoustic tomography maximum intensity projections (∼2 mm thickness) at λ_PA_ = 646 nm (right panels). Magenta arrows highlight signal from the bound dye ligand. Scale bars: 1 mm.

Finally, we set out to investigate the labelling of mouse brains *in vivo* for photoacoustic imaging. Following stereotactic AAV1 HaloTag7-EGFP injection into the righthand hippocampus and thalamus, dye labelling was performed *in vivo* via intracerebroventricular injection of **13-HTL** in the left lateral ventricle. Photoacoustic tomography of the entire excised brains showed no signal in the control mice (no dye injected, Figure 7a). In contrast, strong photoacoustic signal at λ_PA_ = 646 nm was observed in the dye-injected mice, in close proximity to the viral injection site in the hippocampus (Figure 7b). To confirm that the signal specifically arises from the HaloTag-bound dye, the brain was fixed, sliced and fluorescence imaging was performed. Based on the mouse brain atlas,^57^ the slices were mapped to their corresponding coronal section in the PAT volume. The PA signal unambiguously correlated with the far-red fluorescence signal from the dye in the hippocampus, where HaloTag is strongly expressed, as evidenced by EGFP fluorescence (Figure 7c,d,e, Figure S23). *In vivo* labelling in the hippocampus was also achieved with compound **14-HTL**, showing specific labelling of individual neurons (Figure 7f,g, Figure S24). While EGFP signal was also visible in the thalamus, no fluorescence signal from the dye was visible in that region, suggesting that **13-HTL** might show limited bioavailability. In addition, we note that attempts to repeat this labelling resulted in substantially less fluorescence signal in the hippocampus (Figure S25).

**Figure 7.**
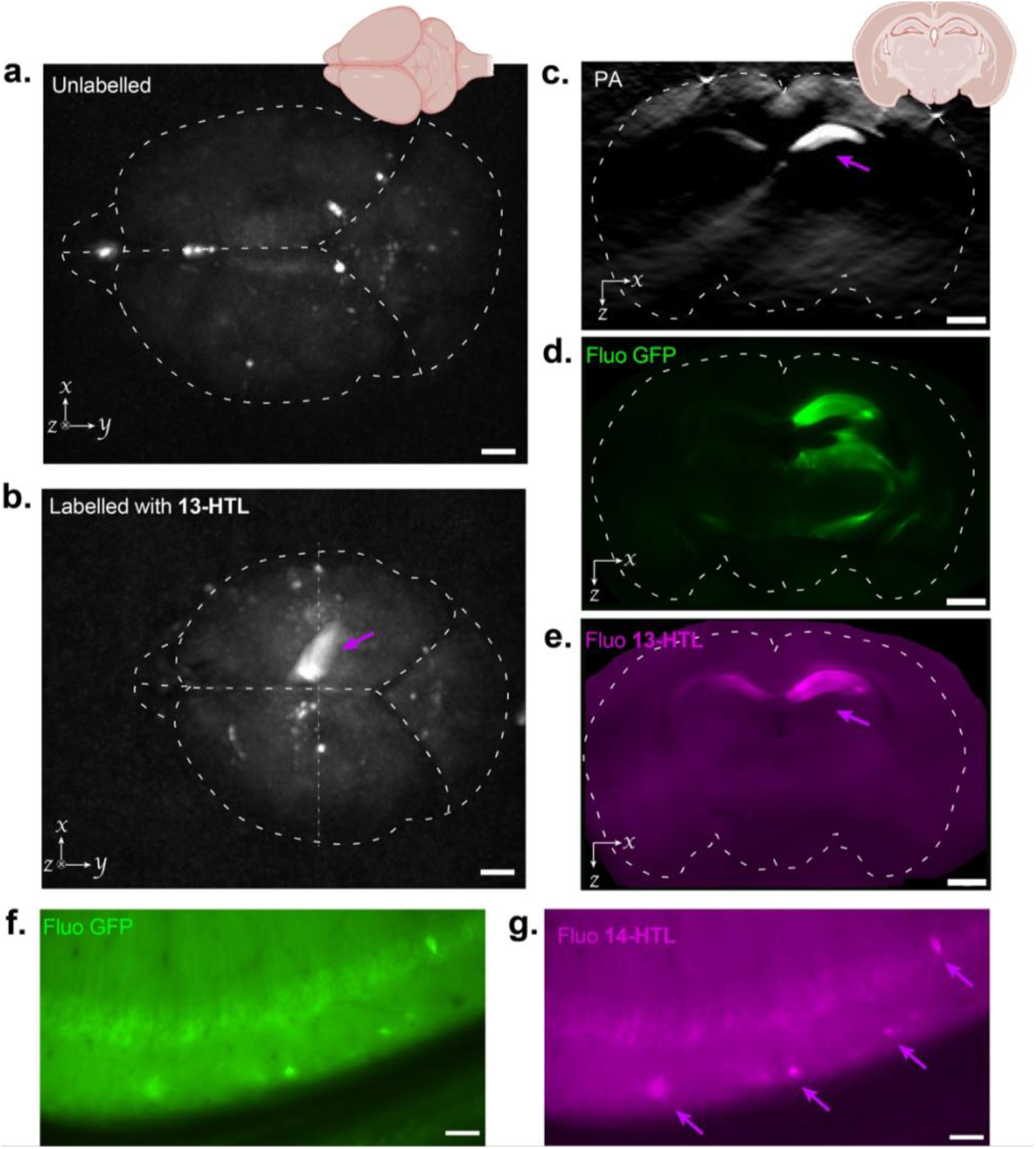
Photoacoustic tomography of a whole *ex vivo* mouse brain expressing HaloTag7-EGFP in neurons, labelled with **13-HTL** delivered by intracerebroventricular injection *in vivo*, and fluorescence images of coronal slices. (**a, b**) PAT maximum intensity axial projection (λ_PA_ = 646 nm) of the entire brain between 2.5 and 5.75 mm from the surface of Fabry-Pérot interferometer to exclude strong endogenous signal from the olfactory bulbs: (**a**) control, unlabelled mouse and (**b**) mouse labelled with **13-HTL** *in vivo* via intracerebroventricular injection. The magenta arrow indicates signal from **13**-HaloTag7; (**c**) selected coronal slice image from the PAT volume and corresponding fluorescence images of (**d**) EGFP and (**e**) **13-HTL**; (**f, g**) Fluorescence images of HaloTag-EGFP expressing neurons labelled with **14-HTL** delivered by intracerebroventricular injection *in vivo*. Images of (**f**) EGFP and (**g**) **14-HTL**. Magenta arrows highlight signal from the bound dye ligand. Scale bar: 1 mm in **a-e**, 50 µM in **f, g**.

Together, these results demonstrate that our acoustogenic ligands can specifically label neurons in mouse brain tissues, and highlight that further improvements will be needed for reliable *in vivo* delivery, ideally systemically.

## Discussion

While a variety of photoacoustic probes have been reported, such as contrast agents for imaging vasculature, tumors and anatomical structures, and reporters showing irreversible activation by biomolecules or analytes of interest,^58, 59, 60^ the field is still lacking specific, dynamic reporters, that can be easily targeted to cells and subcellular features of interest. To resolve these issues, we have repurposed the chemigenetic strategy for the design of genetically targeted photoacoustic probes and calcium sensors, based on the HaloTag self-labelling protein and acoustogenic dye ligands. We synthesized and systematically investigated the photophysical properties of a library of 20 dyes. Through synthetic modifications, we rationally optimized the scaffolds for strong photoacoustic signal in the far-red/NIR region, and high calcium sensitivity when used in combination with the calcium-sensitive HaloCaMP protein. In the process, we establish the century-old dyestuff Malachite Green as a robust photoacoustic probe, and additionally introduce novel synthetic methods to modify its properties, specifically 1’,8’-methylation and mono-hydrodefluorination. These modifications, previously unexplored, are likely to be broadly applicable to rhodamine derivatives, hereby expanding the toolbox of general methods to tune dyes. Overall, this work resulted in a series of acoustogenic dye-ligands with up to 12-fold photoacoustic turn-on, and ultimately 8 calcium indicators with ΔPA/PA_0_ up to 7.5 and affinities ranging from nM to mM. These first-generation chemigenetic probes for PAI constitute, to the best of our knowledge, the first example of positive-going, far-red calcium sensors optimized for this modality. Our probes exhibit superior *in vitro* performance, demonstrated through both spectroscopic and tomographic photoacoustic measurements, outperforming existing PA sensors. Notably, we achieved specific labelling of HaloTag-expressing neurons in mouse brain tissue, leading to strong photoacoustic signal that could be readily visualized via PAT, showing high potential for *in vivo* neuroimaging applications. The variable *in vivo* labelling observed suggests that the bioavailability of the dye ligand is an important limiting factor, affected by solubility, slow diffusion and tissue permeation. Current efforts in our group focus on improving the bioavailability of these acoustogenic dye ligands, a critical property to achieve robust whole brain labelling.

A key feature of the chemigenetic approach is the high tunability of the system afforded by the synthetic component, precluding the need for re-engineering the protein scaffold. Here, by developing photoacoustic ligands that exhibit high performance with established HaloTag and HaloCaMP proteins, we extend the “plug-and-play” versatility of these systems to imaging across various modalities, simply by varying the small-molecule dye ligand. This feature stands as a powerful asset for imaging across spatial scales using different contrast mechanisms. Nevertheless, further refinements of these first-generation sensors are conceivable through protein engineering, which will require innovative high-throughput screening methods and instrumentation, tailored for this modality. We envision that continued improvements in photoacoustic hardware, along with further molecular engineering to refine sensor properties and improve dye bioavailability, will pave the way towards non-invasive calcium imaging across the entire mouse brain, thus positioning photoacoustic imaging to reach its full potential for functional neuroimaging.

## SUPPLEMENTARY MATERIAL

Supplementary figures, general methods, synthetic procedures and characterization for all new compounds (pdf).

CCDC 2336812 contains the supplementary crystallographic data for this paper. These data can be obtained free of charge from The Cambridge Crystallographic Data Centre via www.ccdc.cam.ac.uk/data_request/cif.

## ACKNOWLEDGMENT

This work was supported by the European Molecular Biology Laboratory (EMBL) and the Chan Zuckerberg Initiative (Deep Tissue Imaging grants no. 2020-225346 and no. 2024-337799).

The authors acknowledge: the Lavis group (Janelia Research Campus HHMI) for sharing dyes and for useful discussions, and in particular Luke Lavis for critical reading of this manuscript; the Schreiter group (Janelia Research Campus, HHMI) for plasmids and HaloTag-EGFP virus; the Johnsson group (MPI for Medical Research, Heidelberg) for plasmids and U2OS cell line; Dieter Schollmeyer (Johannes Gutenberg Universität Mainz) for crystallography; the Core Facilities at EMBL: Protein Expression and Purification, Metabolomics, Advanced Light Microscopy Facility, Mechanical Workshop, Gene Editing & Virus Facility for their services and support.

## ABBREVIATIONS

GECI: genetically encoded calcium indicator
HTL: HaloTag ligand
NIR: near-infrared
PA: photoacoustic
PAI: photoacoustic imaging
PAT: photoacoustic tomography
SiR: silicon-rhodamine

