## Supplementary material for "Chemigenetic far-red labels and Ca^2+^ indicators optimized for photoacoustic imaging"

### Supplementary Figures and Tables

**Figure S1**. Chemical structures of the free dyes synthesized and studied in this work.

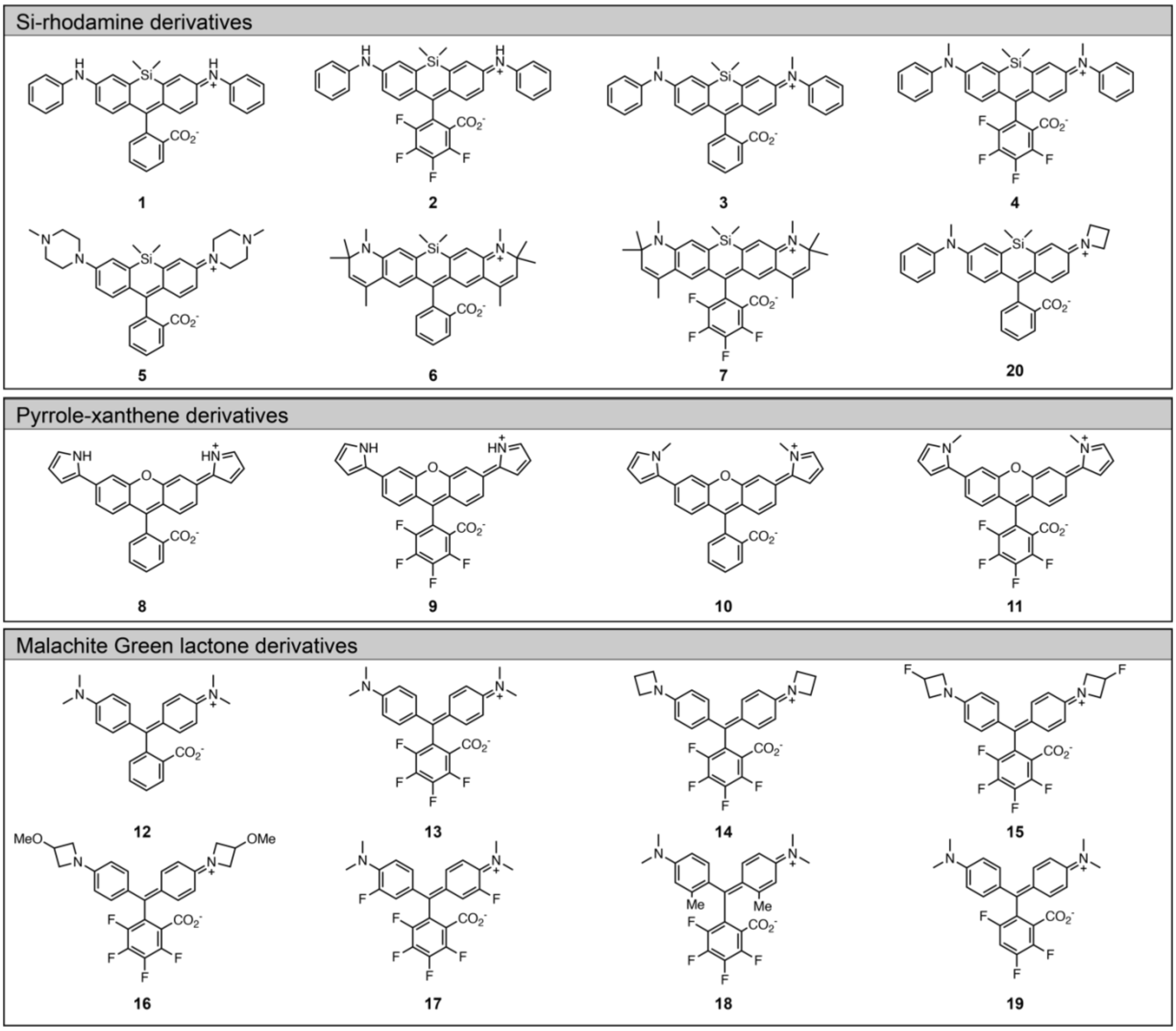

**Figure S2.** Absorption spectra of free dyes **1-20**. All measurements were performed in 10 mM HEPES, pH 7.4 at 1.25 μM dye concentration.

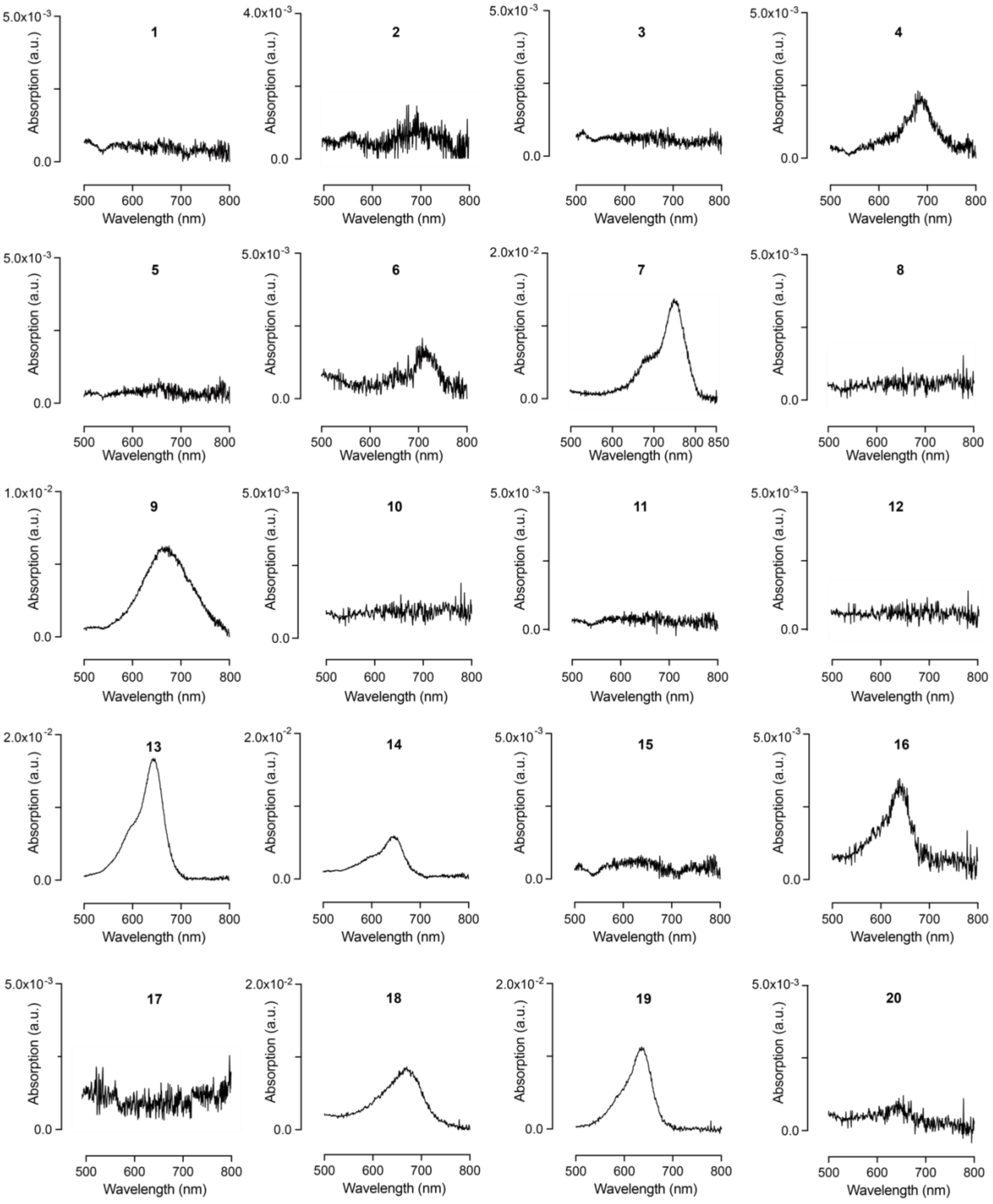

**Figure S3.** Absorption at $\lambda_{max}$ of free dyes **1-20** in MeCN/$H_2O$ mixtures.[1] (**a**) Si-rhodamines and pyrrole-xanthene derivatives; **(b**) Malachite Green lactone derivatives. All measurements were performed at 5 μM dye concentration, values are mean of 3 replicates.

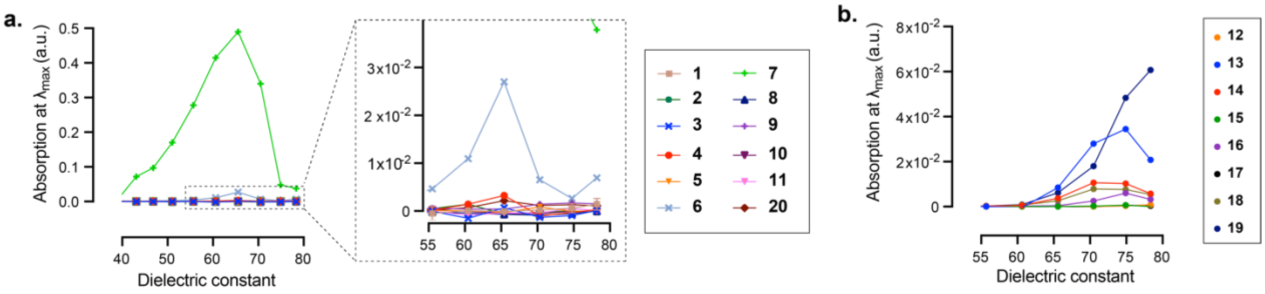

**Figure S4**. Chemical structures of the HaloTag ligands synthesized and studied in this work.

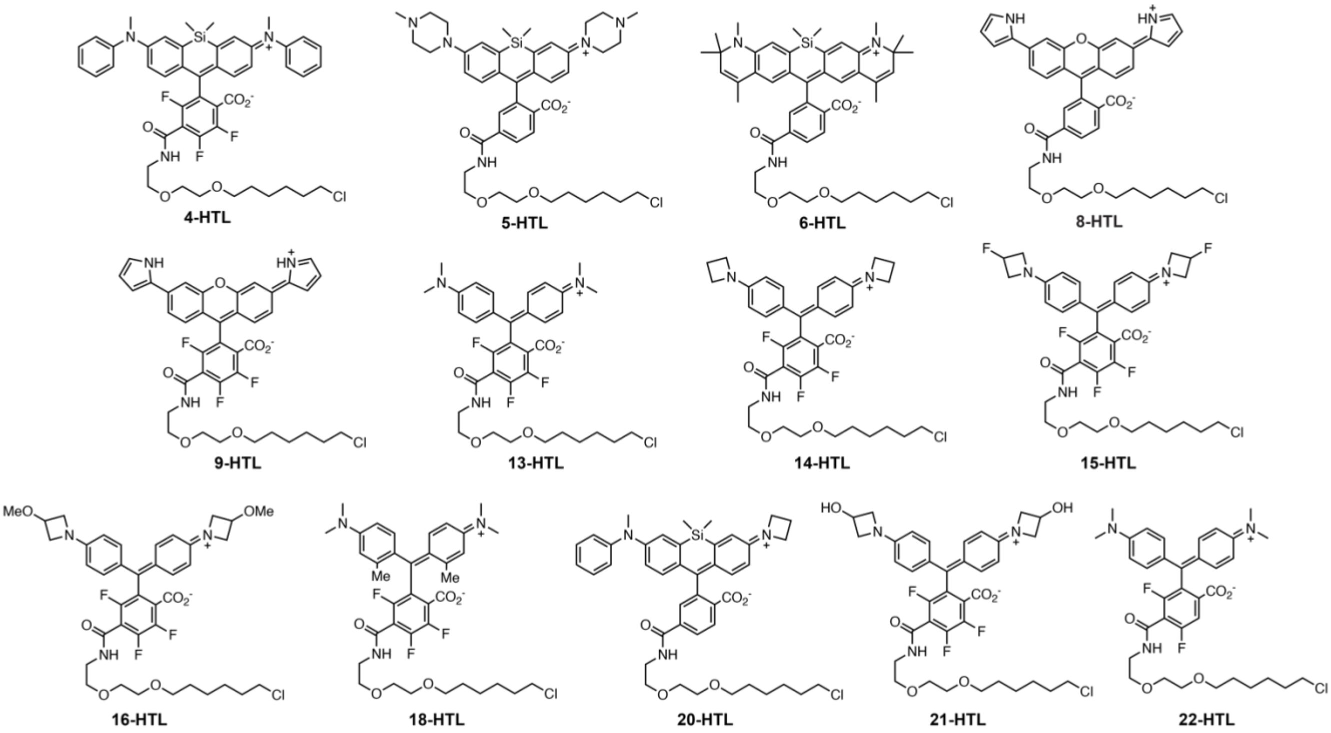

**Table S1.** Properties of the HaloTag ligands bound to HaloTag protein. All measurements were performed in 10 mM HEPES, pH 7.4, containing 0.1 mg·mL$^{-1}$ of CHAPS. Values are the mean of 3 replicates.

| Compound | Protein | $\lambda_{max}$ (nm) | $\lambda_{em}$ (nm) | $\varepsilon$ ($M^{-1}$·$cm^{-1}$) | $\Phi$ | $\Delta A/A_0$ | $\Delta PA/PA_0$ |
|---|---|---|---|---|---|---|---|
| **4-HTL** | +HaloTag7 | 683 | 705 | 44300 | < 0.01 | 62 | - |
| **5-HTL** | +HaloTag7 | 648 | 658 | 3000 | 0.11 | 3.5 | - |
| **6-HTL** | +HaloTag7 | 726 | 749 | 99000 | 0.13 | 94 | - |
| **8-HTL** | +HaloTag7 | 655 | 710 | 3600 | < 0.01 | 5.6 | - |
| **9-HTL** | +HaloTag7 | 695 | 742 | 50300 | < 0.01 | 9.2 | 6.9 |
| | +HaloTag9 | 700 | - | 86300 | - | 18 | 8.6 |
| **13-HTL** | +HaloTag7 | 646 | 666 | 92400 | 0.001 | 3.6 | 2.8 |
| | +HaloTag9 | 648 | - | 114100 | - | 4.6 | 2.9 |
| **14-HTL** | +HaloTag7 | 648 | 662 | 75100 | < 0.01 | 10 | 8.1 |
| | +HaloTag9 | 650 | - | 81600 | - | 11 | - |
| **15-HTL** | +HaloTag7 | 638 | 650 | 11400 | < 0.01 | 21 | - |
| **16-HTL** | +HaloTag7 | 643 | 661 | 48800 | < 0.01 | 16 | 12 |
| | +HaloTag9 | 645 | - | 53200 | - | 17 | - |
| **18-HTL** | +HaloTag7 | 672 | 713 | 50700 | < 0.01 | 8.9 | 5.7 |
| | +HaloTag9 | 670 | - | 76800 | - | 13 | - |
| **20-HTL** | +HaloTag7 | 654 | 679 | 68800 | 0.01 | 42 | - |
| **21-HTL** | +HaloTag7 | 644 | 658 | 43100 | < 0.01 | 1.5 | 1.0 |
| **22-HTL** | +HaloTag7 | 640 | 656 | 45900 | < 0.01 | 11 | 5.7 |
| | +HaloTag9 | 641 | - | 50500 | - | 13 | - |

**Figure S5**. Absorption spectra of HaloTag ligands in the absence of protein (dashed black lines), bound to HaloTag7 (red lines) or bound to HaloTag9 (blue lines). All measurements were performed in 10 mM HEPES, pH 7.4, containing 0.1 mg·mL$^{-1}$ of CHAPS at a dye ligand concentration of 1.25 µM and protein concentration of 1.9 µM, in triplicate.

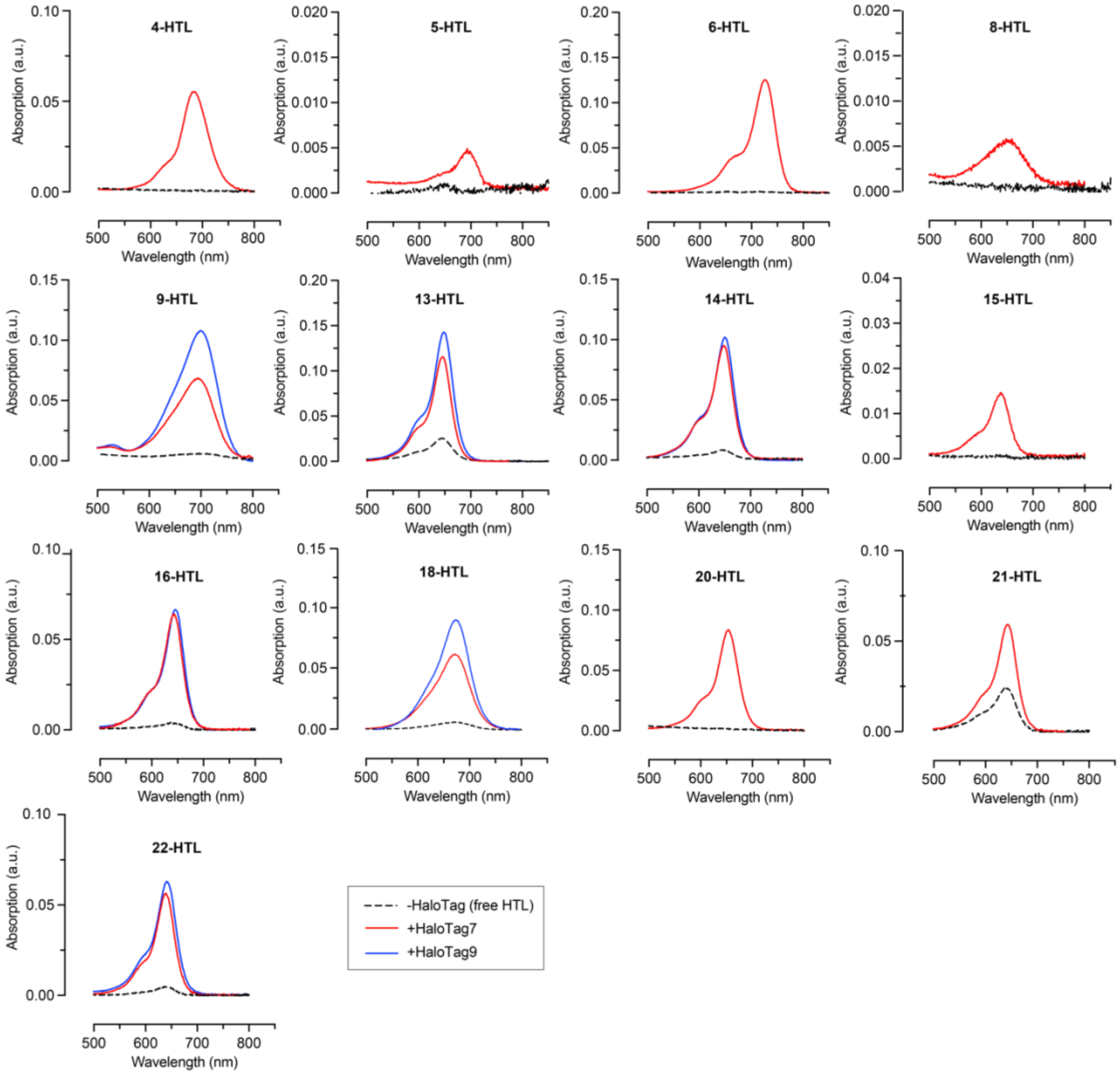

**Figure S6.** Binding kinetics of selected HaloTag ligands to HaloTag7. All measurements were performed in 10 mM HEPES, pH 7.4, containing 0.1 mg·mL$^{-1}$ of CHAPS at 5 µM of HaloTag ligand and 7.5 µM of HaloTag7 protein.

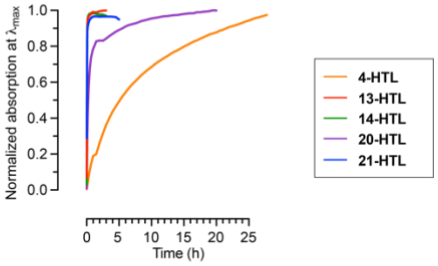

**Figure S7.** Photoacoustic spectra of selected HaloTag ligands in the absence of protein (dashed black lines) or bound to HaloTag7 (red lines). All measurements were performed in 10 mM HEPES, pH 7.4, containing 0.1 mg·mL$^{-1}$ CHAPS at a dye ligand concentration of 1.25 µM and protein concentration of 1.9 µM, in triplicate.

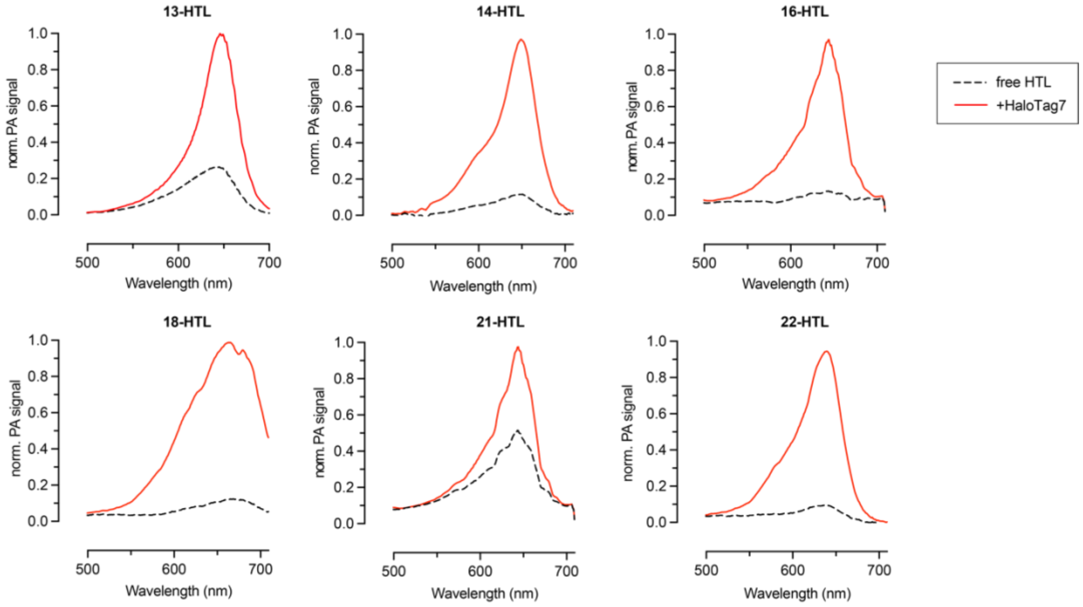

**Figure S8.** ΔPA/PA$_0$ vs ΔA/A$_0$ for selected HaloTag ligands upon binding to HaloTag protein. Measurements were performed in triplicate at 1.25 µM HaloTag ligand and 1.9 µM protein, in 10 mM HEPES, pH 7.4 containing 0.1 mg·mL$^{-1}$ CHAPS. "HTL" was omitted from the labels in the figure for clarity.

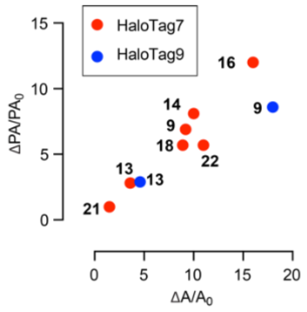

**Figure S9**. Normalized absorption at $\lambda_{max}$ (black) and normalized photoacoustic signal at $\lambda_{PA}=\lambda_{max}$ (red) for **JF635-HTL** and **13-HTL** bound to HaloTag7 (HT7). Measurements were performed in triplicate at 1.25 µM HaloTag ligand and 1.9 µM HaloTag7, in 10 mM HEPES, pH 7.4 containing 0.1 mg·mL$^{-1}$ CHAPS. Values were normalized to **JF635-HTL**.

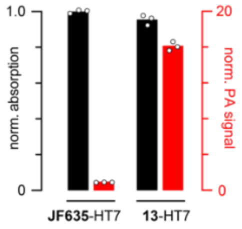

**Table S2**. Properties of photoacoustic calcium sensors. All measurements were performed in 30 mM MOPS, 100 mM KCl, pH 7.2, containing 0.1 mg·mL$^{-1}$ of CHAPS, to which either 10 mM EGTA (for the calcium free state) or excess $CaCl_2$ was added (for the calcium saturated state, i.e. 500 mM $CaCl_2$ for HaloCaMP1a or 10 mM $CaCl_2$ for HaloCaMP1b and rHCaMP) was added. Values are the mean of 3 replicates. [a]Data from ref [2]. [b]Data from ref [3]. [c]Data from ref [4].

| Compound | protein | $\lambda_{max}$ (nm) | $\varepsilon_{sat}$ ($M^{-1}\cdot cm^{-1}$) | $\Delta A/A_0$ | $\Delta PA/PA_0$ | $K_d$ |
|---|---|---|---|---|---|---|
| **9-HTL** | HaloCaMP1a | 700 | 23600 | 1.4 | 1.8 | 6.9 mM |
| | HaloCaMP1b | 704 | 36000 | 2.1 | 1.3 | 16 nM |
| | rHCaMP | 693 | 5400 | -0.5 | - | - |
| **13-HTL** | HaloCaMP1a | 646 | 69100 | 0.7 | 1.1 | 300 µM; 18 mM |
| | HaloCaMP1b | 645 | 29300 | 3.3 | 3.1 | 17 nM |
| | rHCaMP | 645 | 17000 | -0.4 | - | - |
| **14-HTL** | HaloCaMP1a | 649 | 30500 | 0.8 | 1.5 | 4.8 mM |
| | HaloCaMP1b | 650 | 15200 | 2.4 | 3.2 | 10 nM |
| | rHCaMP | 648 | 7500 | -0.3 | - | - |
| **16-HTL** | HaloCaMP1a | 643 | 13200 | 4.1 | - | - |
| | HaloCaMP1b | 642 | 4100 | 1.5 | - | - |
| | rHCaMP | 645 | 5300 | -0.4 | - | - |
| **18-HTL** | HaloCaMP1a | 674 | 20900 | 1.0 | 2.0 | 980 µM; 10 mM |
| | HaloCaMP1b | 675 | 15300 | 16 | 7.5 | 33 nM |
| | rHCaMP | 670 | 5900 | 0.01 | - | - |
| **22-HTL** | HaloCaMP1a | 640 | 14700 | 0.9 | - | - |
| | HaloCaMP1b | 638 | 7300 | 0.2 | - | - |
| | rHCaMP | 640 | 6100 | -0.4 | - | - |
| NIR-GECO1[a] | | 678[a] | 20000[a] | -0.68[a] | -0.5 | 215 nM[a] |
| CASPA_550[b] | | 550[b] | 77745[b] | ~-0.5[b] | ~-0.5[b] | ~4 µM[b] |
| "L"[c] | | 765[c] | 194000[c] | ~-0.5[c] | ~-0.5[c] | 11.3 µM[c] |

**Figure S10.** Absorption spectra of the chemigenetic calcium sensors in the calcium-free (dashed black lines) and calcium-bound states (solid red lines). All measurements were performed in 30 mM MOPS, 100 mM KCl, pH 7.2, containing 0.1 mg·mL$^{-1}$ CHAPS to which either 10 mM EGTA (for the calcium free state) or excess $CaCl_2$ (for the calcium saturated state, i.e. 500 mM $CaCl_2$ for HaloCaMP1a or 10 mM $CaCl_2$ for HaloCaMP1b and rHCaMP) was added. For HaloCaMP1a and HaloCaMP1b, solutions were prepared with 1.25 µM HaloTag ligand and 1.9 µM protein. For rHCaMP, solutions were prepared with 2.5 µM HaloTag ligand and 3.8 µM protein**.** Measurements were performed in triplicate.

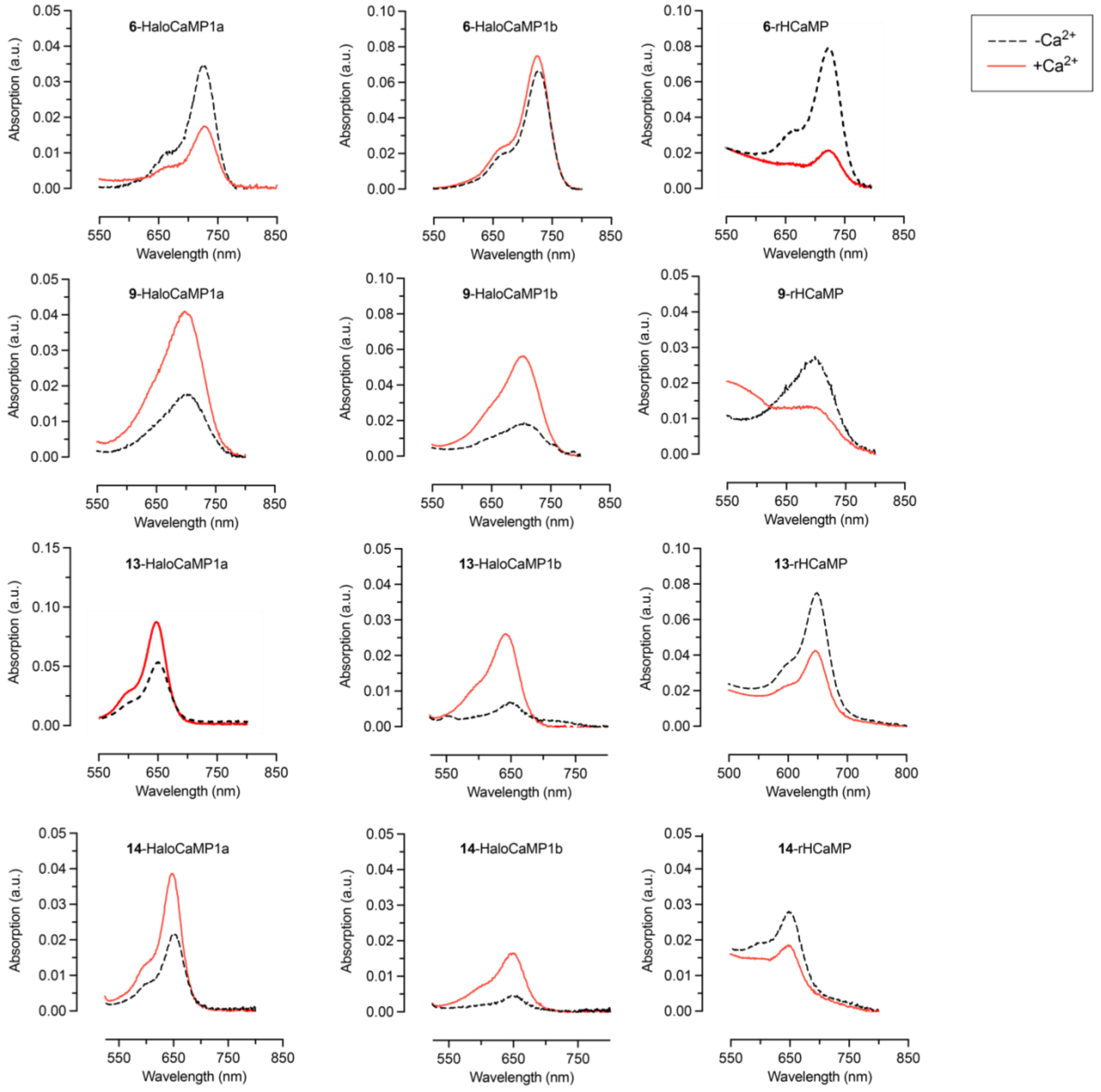

**Figure S10 –** continued.

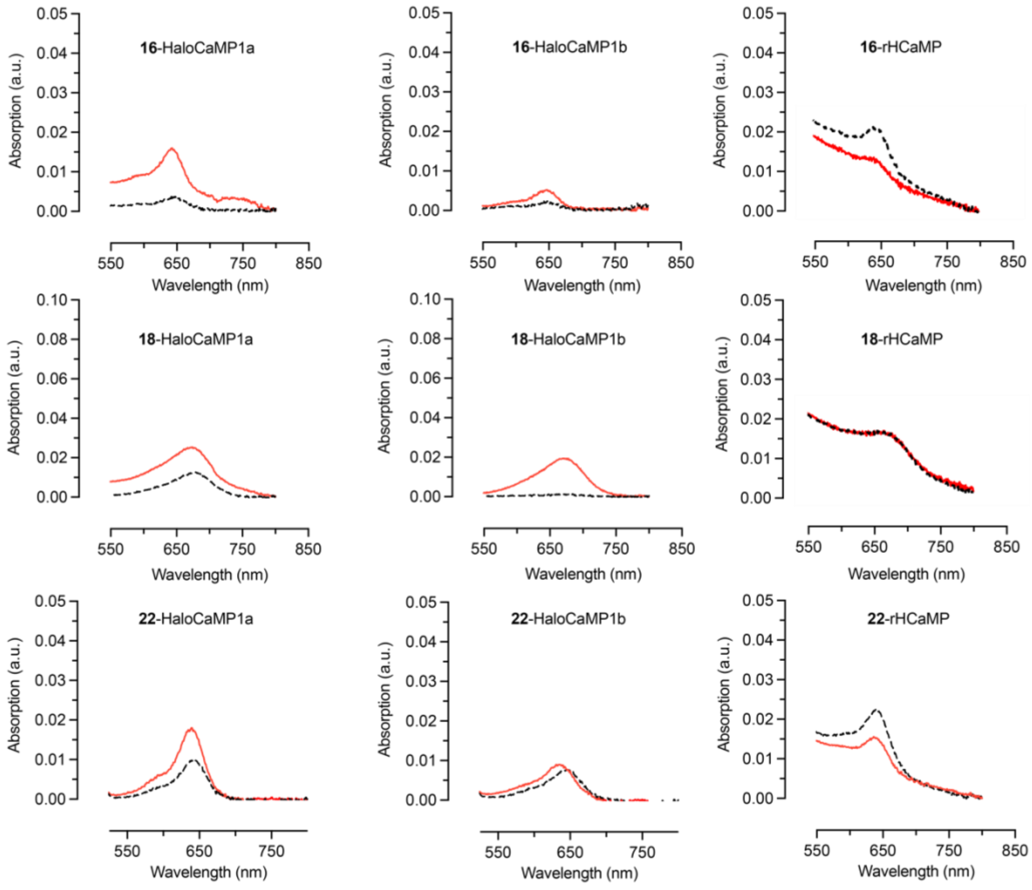

**Figure S11**. Photoacoustic spectra of selected chemigenetic calcium sensors in the calcium-free (dashed black lines) and calcium-bound states (solid red lines). All measurements were performed in 30 mM MOPS, 100 mM KCl, pH 7.2, containing 0.1 mg·mL$^{-1}$ CHAPS to which 10 mM EGTA was added (for the calcium free state), or excess $CaCl_2$ was added (for the calcium saturated state, 500 mM $CaCl_2$ for HaloCaMP1a or 10 mM $CaCl_2$ for HaloCaMP1b and rHCaMP). For HaloCaMP1a and HaloCaMP1b, solutions were prepared with 1.25 µM HaloTag ligand and 1.9 µM protein. Measurements were performed in triplicate.

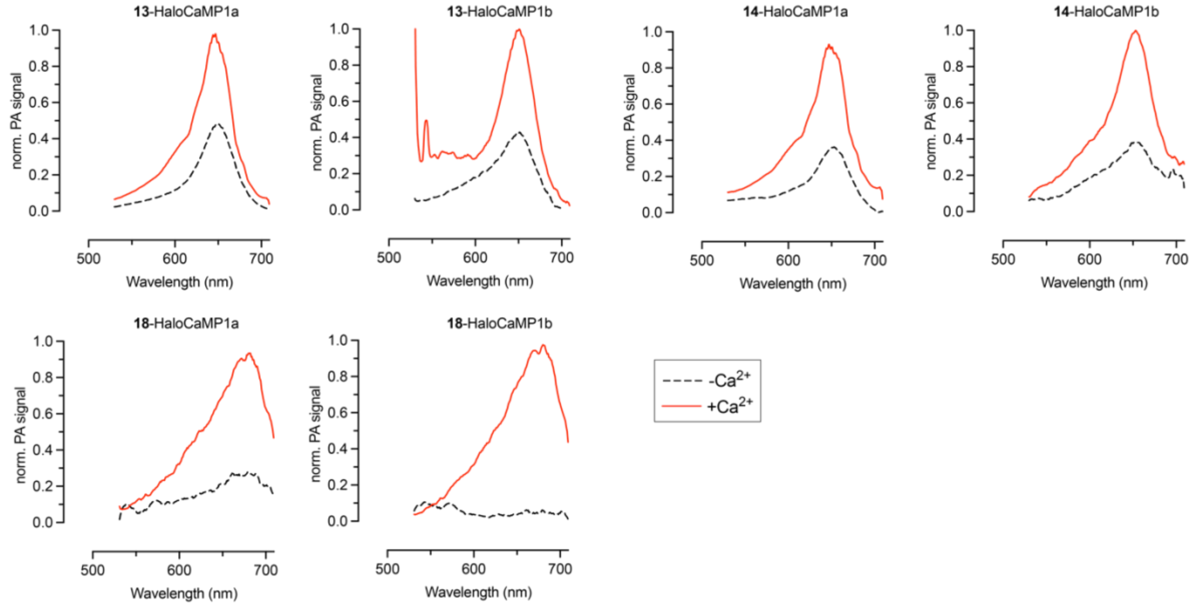

**Figure S12.** $\Delta PA/PA_0$ vs $\Delta A/A_0$ for selected chemigenetic calcium sensors upon binding $Ca^{2+}$. Measurements were performed in triplicate at 1.25 µM HaloTag ligand and 1.9 µM protein, in 10 mM HEPES, pH 7.4 containing 0.1 mg·mL$^{-1}$ CHAPS. “HTL” was omitted from the labels for clarity.

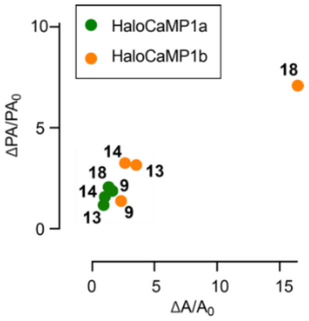

**Figure S13.** pH sensitivity of calcium sensors **13**-HaloCaMP1a (**a**) and **13**-HaloCaMP1b (**b**). Absorbance at $\lambda_{max}$ measured in citrate, phosphate and TRIS buffers in the absence (black) or presence (red) of calcium. Measurements were performed in duplicate at 1.25 µM dye ligand and 1.9 µM protein.

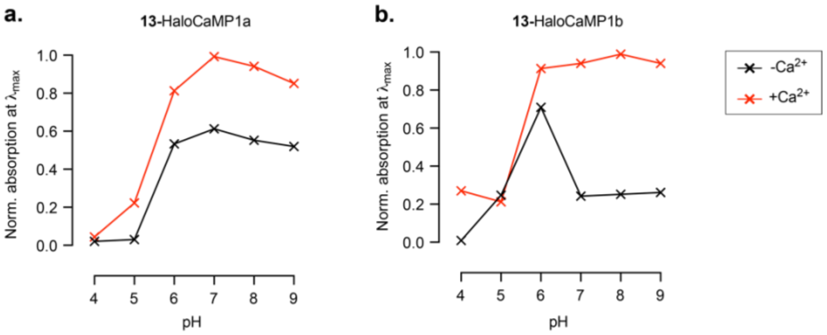

**Figure S14**. Selectivity of sensors **13**-HaloCaMP1a (**a**) and **13**-HaloCaMP1b (**b**) for $Ca^{2+}$ over $Mg^{2+}$. All measurements were performed at 1.25 µM dye and 1.9 µM protein. Measurements were performed in 30 mM MOPS, 100 mM KCl, pH 7.2 containing EGTA (10 µM for HaloCaMP1a, 10 mM for HaloCaMP1b), in duplicate.

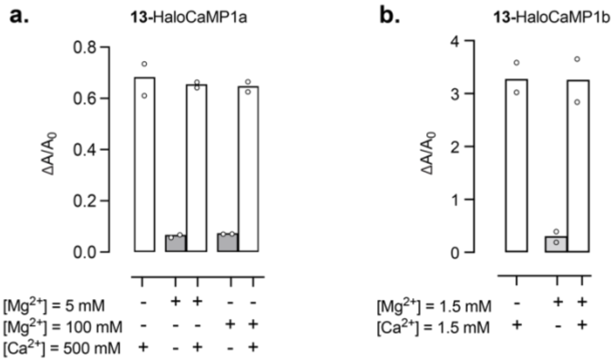

**Figure S15**. Photoacoustic signal of absorbance-matched samples during irradiation at $\lambda_{PA} = \lambda_{max}$ for 1 h at 100 Hz pulse repetition rate. Normalized photoacoustic signal plotted against cumulative illumination energy. (**a**) Photoacoustic labels: mIFP ($\lambda_{PA}$ = 683 nm), Cy5 ($\lambda_{PA}$ = 646 nm), **13**-HaloTag7 ($\lambda_{PA}$ = 646 nm); (**b**) Calcium sensors in the $Ca^{2+}$ saturated state: NIR-GECO1 ($\lambda_{PA}$ = 678 nm), **13**-HaloCaMP1a ($\lambda_{PA}$ = 646 nm), **13**-HaloCaMP1b ($\lambda_{PA}$ = 645 nm); Normalised photoacoustic signal plotted against time of the photoacoustic labels (**c**), and calcium sensors (**d**).

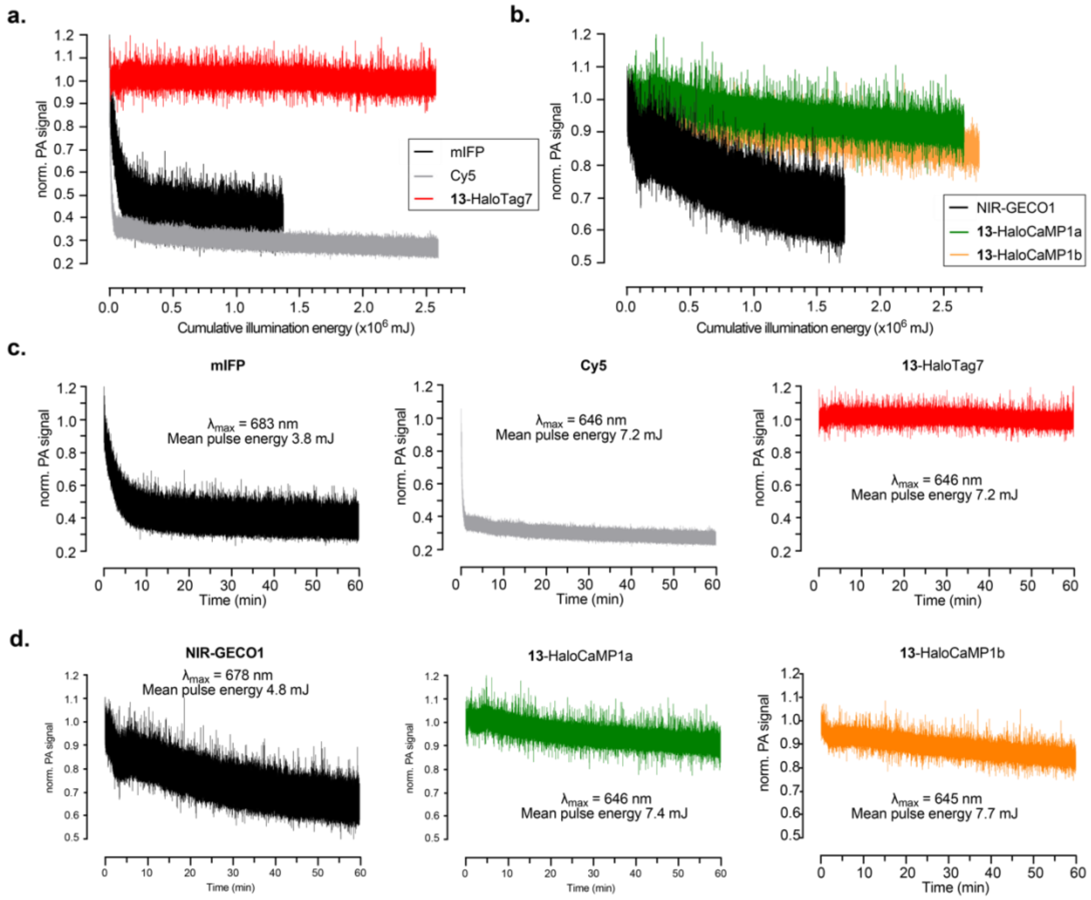

**Figure S16**. Representative photoacoustic tomography images of tissue-mimicking phantoms of mIFP (right tube) and **13**-HaloTag7 (left tube). The tubes were filled with a 50 µM protein solution, and immobilized in coupling medium which was either $H_2O$ or 60% v/v milk/$H_2O$. (xy) images are maximum intensity projections and (xz) images are maximum intensity projections of the reconstructed side view. All images are displayed to their respective minimum and maximum intensities. The Fabry-Pérot interferometer is located on the z = 0 plane, $\lambda_{PA}$ = 646 nm. Intensity measurement was performed using a line profile (250 µm thick) on the (xz) maximum intensity projections.

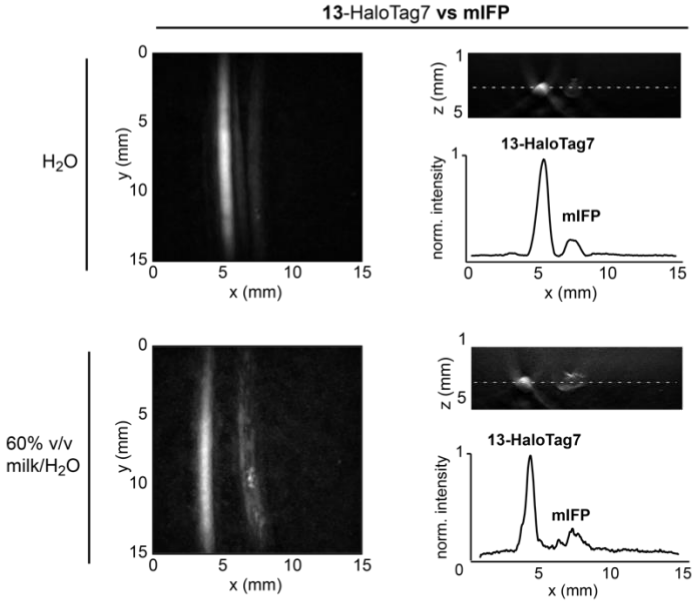

**Figure S17.** Representative photoacoustic tomography images of tissue-mimicking phantoms of mIFP (**a**) and **13**-HaloTag7 (**b**)**.** The tubes were filled with a 50 µM protein solution and immobilized at different depths in coupling medium which was either $H_2O$, or 60% v/v milk/$H_2O$. (xy) images are maximum intensity projections and (xz) images are maximum intensity projections of the reconstructed side view. All images are displayed to their respective minimum and maximum intensities. The Fabry-Pérot interferometer is located on the z = 0 plane, $\lambda_{PA}$ = 683 nm for mIFP, $\lambda_{PA}$ = 646 nm for **13**-HaloTag7.

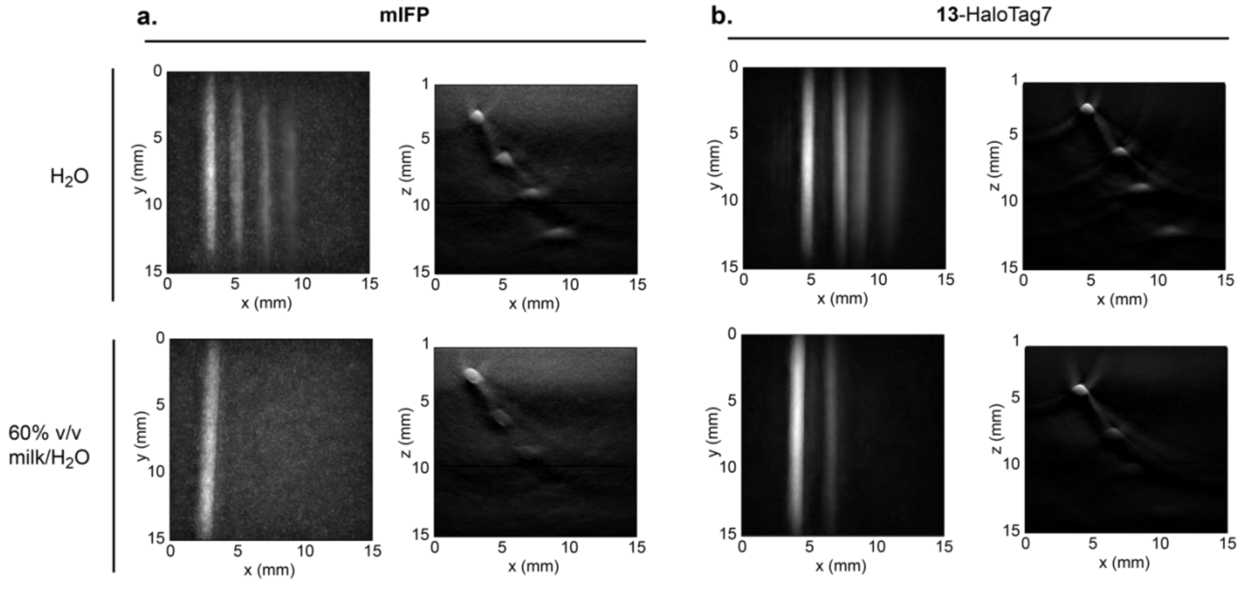

**Figure S18.** Representative photoacoustic tomography images of tissue-mimicking phantoms of **13**-HaloTag7 at different concentrations at 5 mm depth. The tubes were filled with 5, 10, 25 or 50 µM protein solutions and immobilized in coupling medium which was either $H_2O$, or 60% v/v milk/$H_2O$. (xy) images are maximum intensity projections and (xz) images are maximum intensity projections of the reconstructed side view. All images are displayed to their respective minimum and maximum intensities. The Fabry-Pérot interferometer is located on the z = 0 plane, $\lambda_{PA}$ = 646 nm. Intensity measurement was performed using a line profile (250 µm thick) on the (xz) maximum intensity projections.

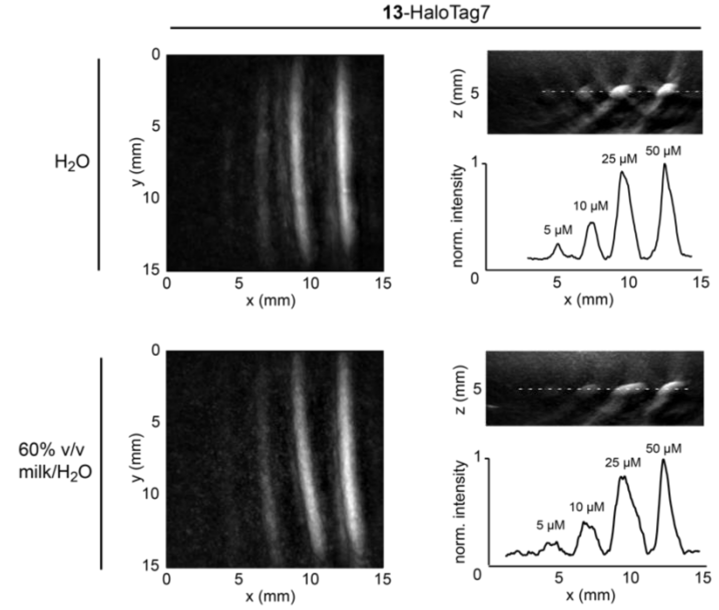

**Figure S19.** Representative photoacoustic tomography images of tissue-mimicking phantoms of **13**-HaloCaMP1a, **13**-HaloCaMP1b and NIR-GECO1 in the calcium-bound (left tubes) or calcium-free (right tubes) states. The tubes were filled with 50 µM protein solution to which either excess $CaCl_2$ or excess EGTA was added. The tubes were immobilized in coupling medium which was either $H_2O$, or 60% v/v milk/$H_2O$. The (xz) images are maximum intensity projections of the reconstructed side view. All images are displayed to their respective minimum and maximum intensities. The Fabry-Pérot interferometer is located on the z = 0 plane, $\lambda_{PA}$ = 645 nm for **13-**HaloCaMP1a/b and $\lambda_{PA}$ = 678 nm for NIR-GECO1. Intensity measurement was performed using a line profile (250 µm thick) on the (xz) maximum intensity projections.

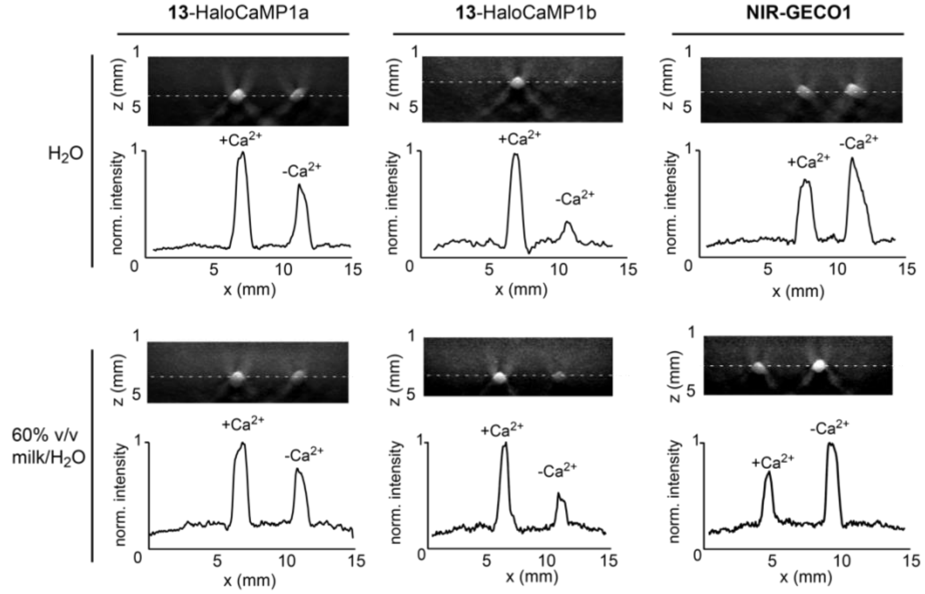

**Figure S20**. Representative photoacoustic tomographic images of tissue-mimicking phantoms of side-by-side comparison of the calcium-bound states of **13**-HaloCaMP1a **(13-1a**, left tubes), **13**-HaloCaMP1b **(13-1b**, middle tubes) and NIR-GECO1 (right tubes). The tubes were filled with a 50 µM protein solution containing excess $CaCl_2$ and embedded in coupling medium which was either $H_2O$ or 60% v/v milk/$H_2O$. (xy) images are maximum intensity projections and (xz) images are maximum intensity projections of the reconstructed side view. All images are displayed to their respective minimum and maximum intensities. The Fabry-Pérot interferometer is located on the z = 0 plane, $\lambda_{PA}$ = 645 nm. Intensity measurement was performed using a line profile (250 µm thick) on the (xz) maximum intensity projections.

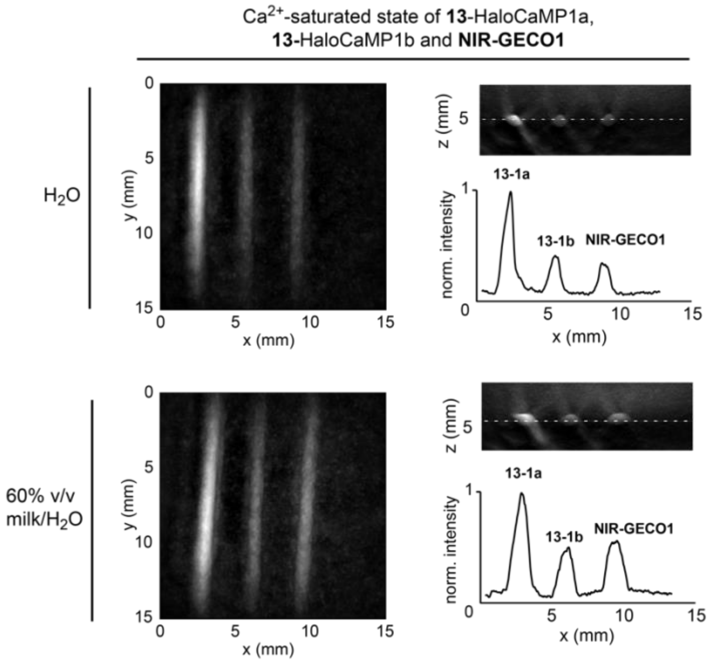

**Figure S21.** Representative widefield fluorescence images of live U2OS cells expressing HaloTag7-EGFP labelled with acoustogenic dye ligands, followed by **JF549-HTL** to assess labelling efficiency. Cells were incubated with 1 µM of either **JF549-HTL**, **9-HTL**, **13-HTL**, or **14-HTL** for 2 h, washed, then incubated with 0.5 µM **JF549-HTL** for 30 mins and washed again. Images in three channels correspond to the fluorescence from EGFP, **JF549-HTL** and from the acoustogenic dye. Representative images from 2 wells for each labelling condition, 4 images per well. Scale bars: 100 µm.

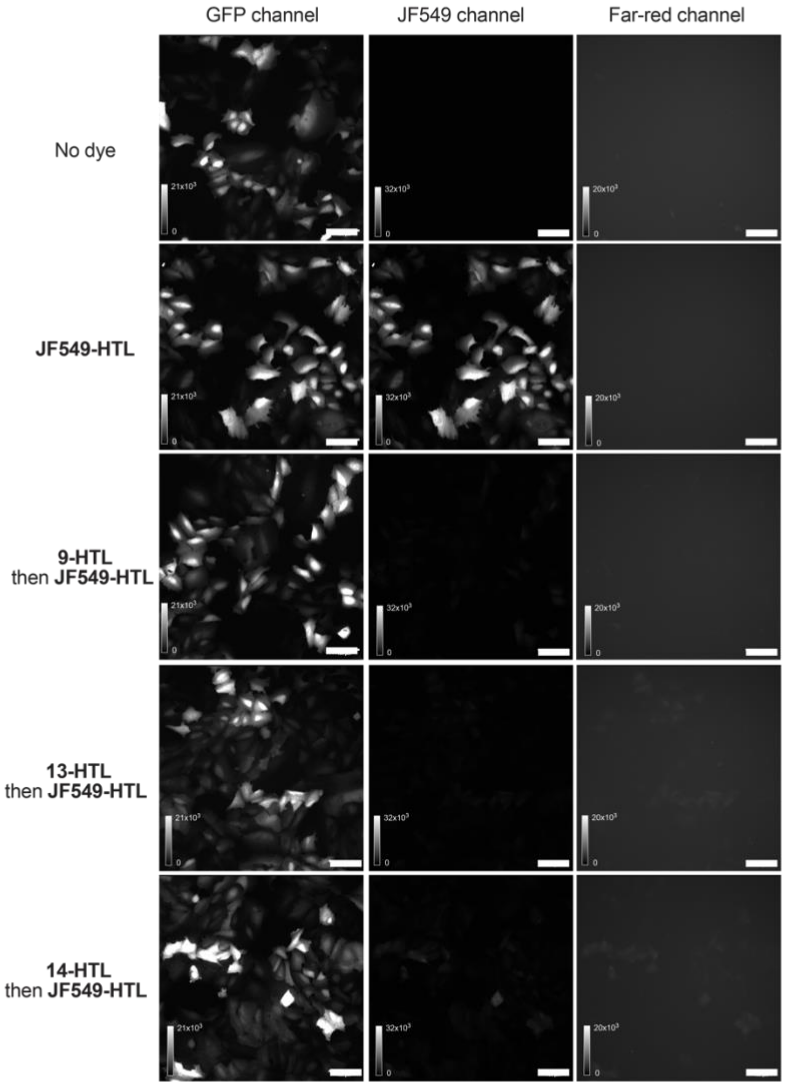

**Figure S22.** Fluorescence microscopy images of the coronal mouse brain slices in Figure 6, before bath labelling with **13-HTL**. Fluorescence images of EGFP channel (left panel) and **13-HTL** channel (right panel) in coronal brain slices expressing HaloTag7-EGFP in the hippocampus before bath labelling with **13-HTL**. Scale bars: 1 mm.

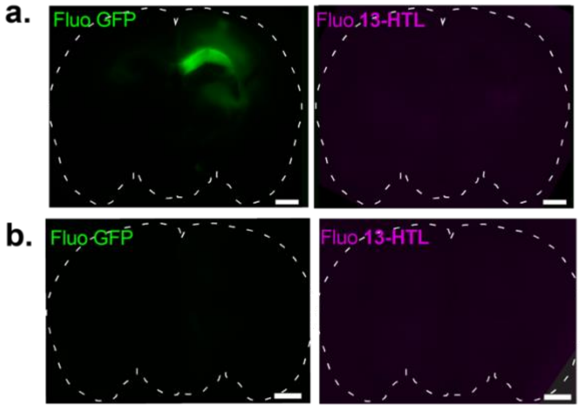

**Figure S23.** Photoacoustic tomography of a whole *ex vivo* mouse brain expressing HaloTag7-EGFP in neurons, labelled with **13-HTL** delivered by intracerebroventricular injection *in vivo,* and fluorescence images of coronal slices. (**a**) PAT maximum intensity axial projection of the entire brain between 2.5 and 5.75 mm from the surface of Fabry-Pérot interferometer to exclude strong endogenous signal from the olfactory bulbs. The magenta dashed line indicates the position of the coronal slice ($\lambda_{PA}$ = 646 nm); (**b**) Widefield fluorescence image in the EGFP channel; (**c**) Widefield fluorescence image in the far-red channel; (**d**) Overlay of the EGFP and **13-HTL** channels; (**e**) Corresponding coronal slice from the PAT volume. Scale bars: 1 mm.

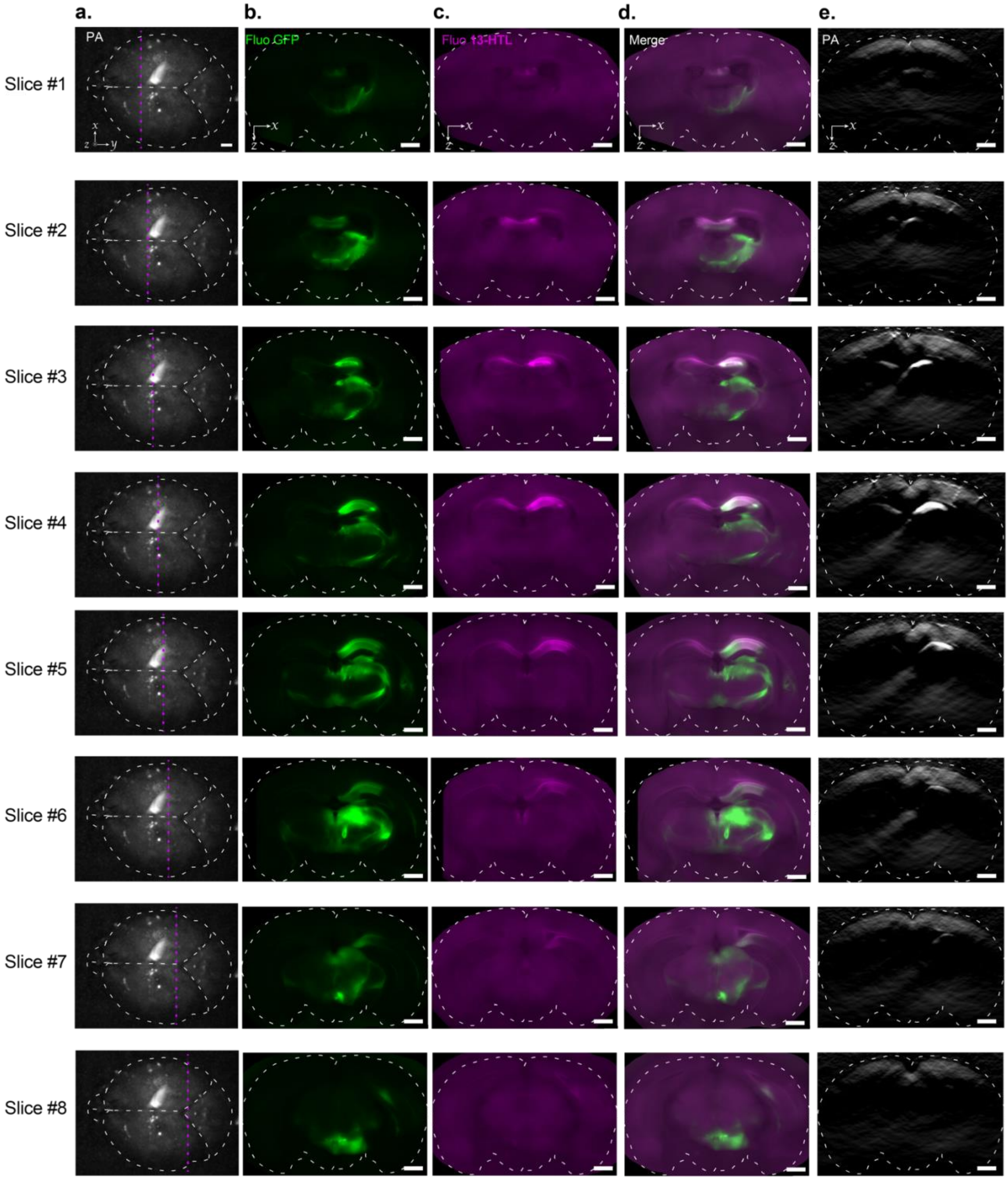

**Figure S24**. Additional representative widefield fluorescence images of coronal slices of the mouse brain. HaloTag7-EGFP expressing neurons were labelled with **14-HTL** delivered through intracerebroventricular injection *in vivo*. (**a**) Green channel corresponding to the EGFP signal; (**b**) Far-red channel corresponding to the fluorescence of **14-HTL**. Scale bars: 50 µM.

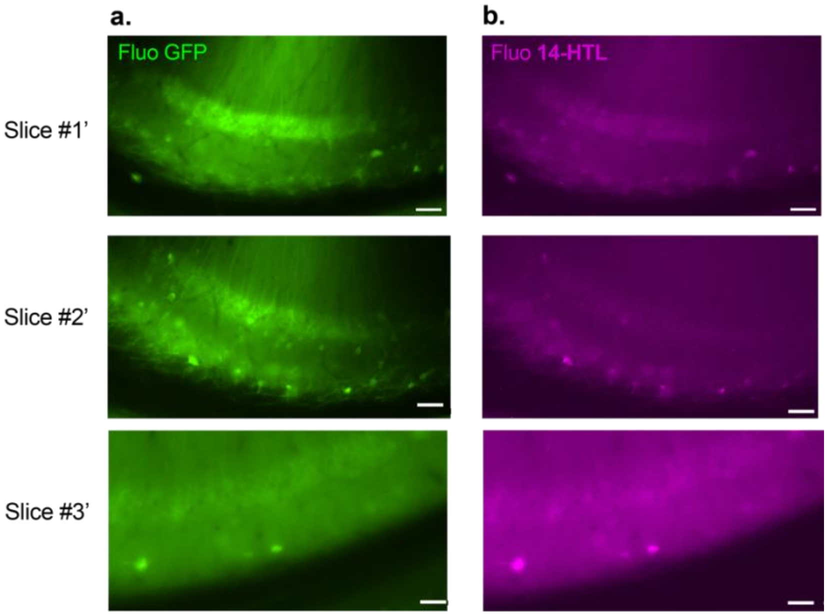

**Figure S25**. Additional widefield fluorescence images of coronal slices of a mouse brain showing partial labelling with **13-HTL,** delivered by intracerebroventricular injection *in vivo***.** Images in green channel corresponding to the EGFP signal (left panels) and images in the far-red channel corresponding to the fluorescence of **13-HTL** (right panels). Scale bars: 1 mm.

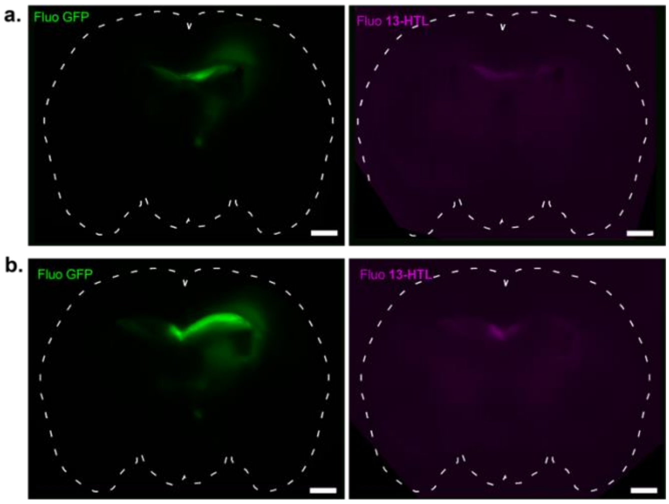

### Synthetic Schemes

Synthesis of **1**, **2**, **3**, **4**:

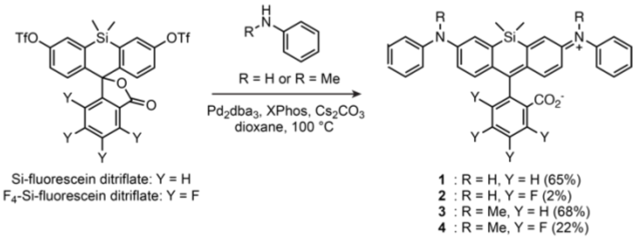

Synthesis of **5**:

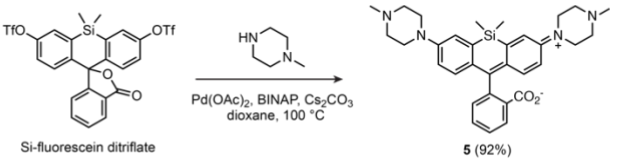

Synthesis of **8**, **9**, **10**, **11**:

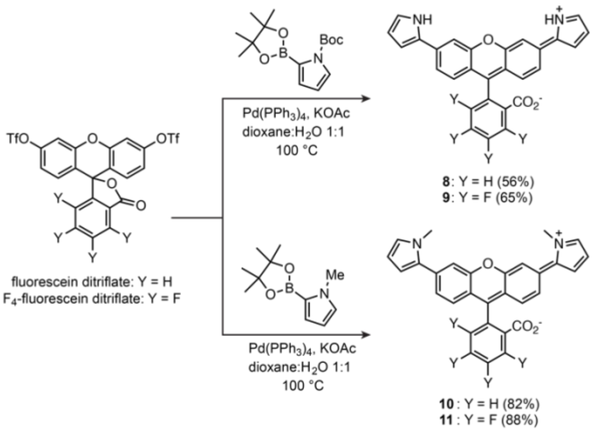

Synthesis of **12**, **13**:

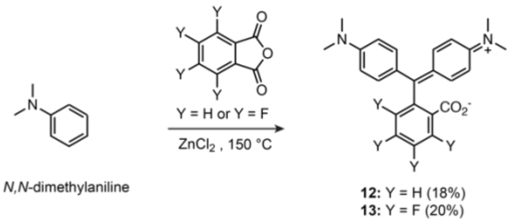

Synthesis of **S1**, **S2**, **S3**, **S4, S5**:

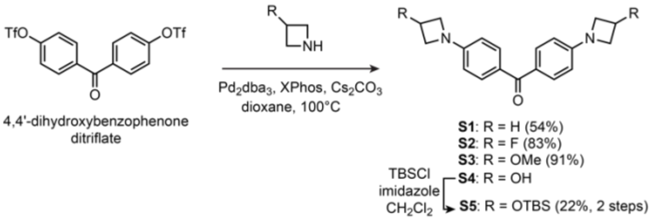

Synthesis of **14**, **15**, **16**, **S6**:

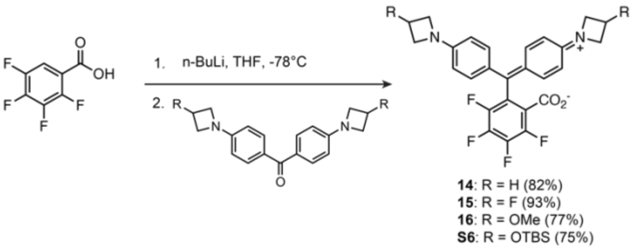

Synthesis of **17, 18**:

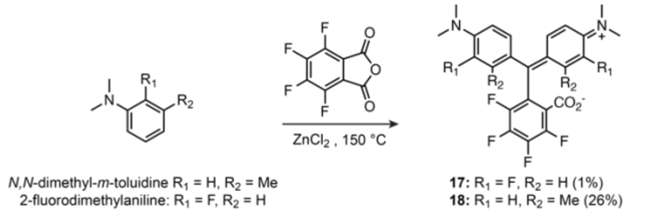

Synthesis of **19**:

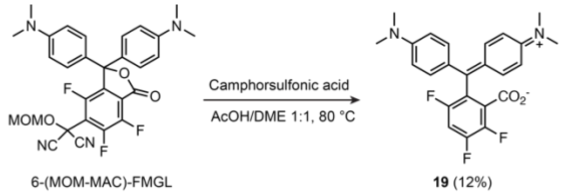

Synthesis of **20**:

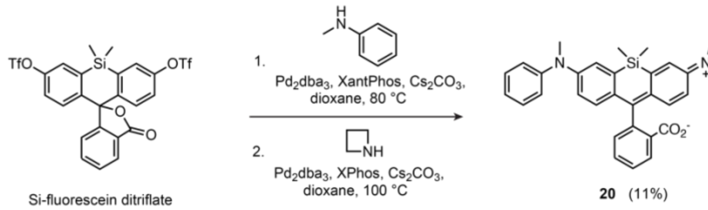

Synthesis of **20-HTL**:

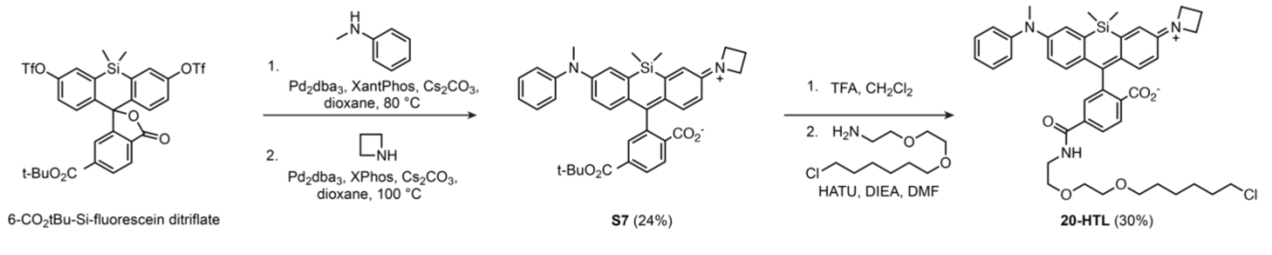

Synthesis of **5-HTL**:

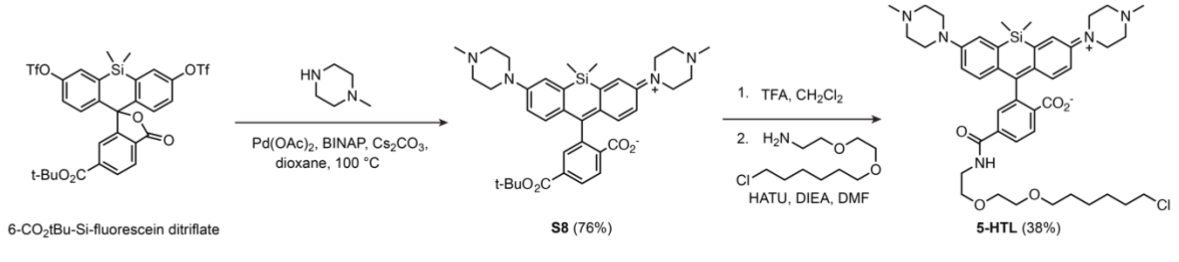

Synthesis of **8-HTL**:

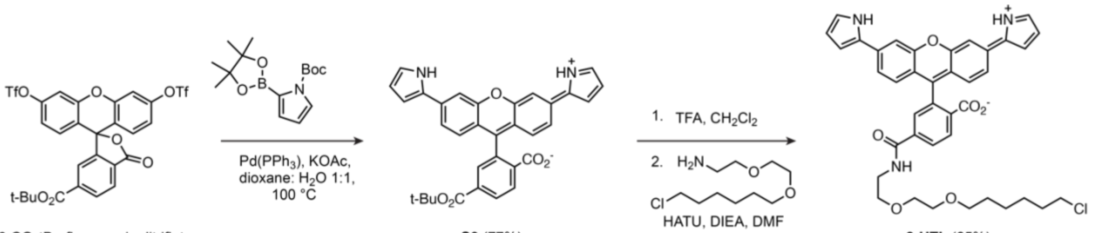

6-CO2tBu-fluorescein ditriflate **S9** (77%) **8-HTL** (65%)

Synthesis of **S10**:

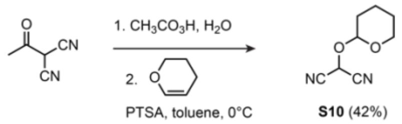

Synthesis of **9-HTL**:

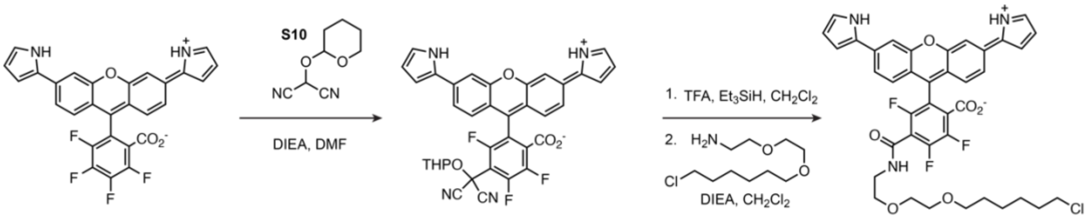

**9** **S11** **9-HTL** (9%, 3 steps)

Synthesis of **14-HTL, 15-HTL, 16-HTL, 21-HTL**:

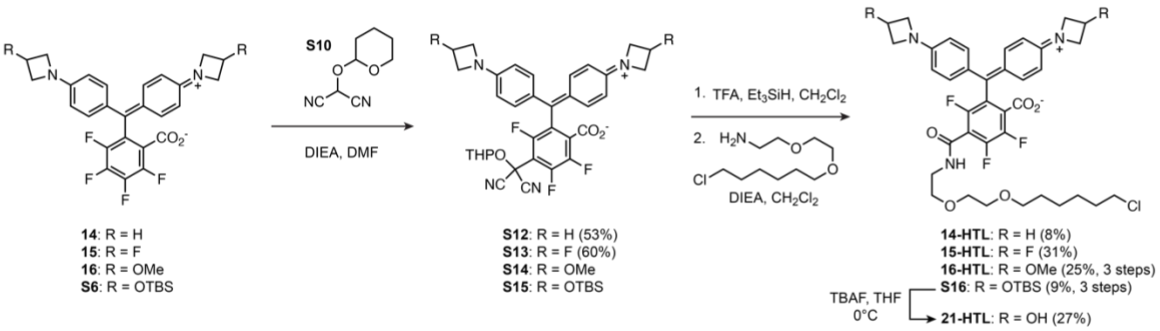

Synthesis of **18-HTL**:

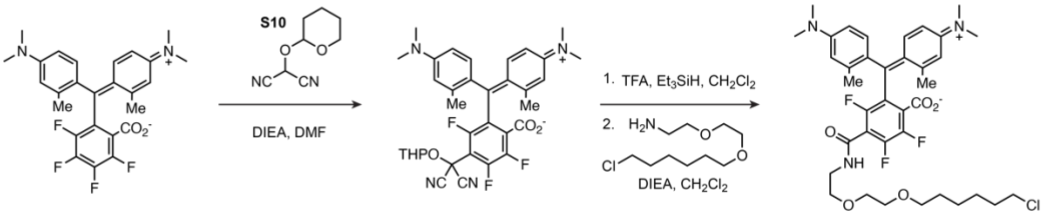

**18** **S17** **18-HTL** (22%, 3 steps)

Synthesis of **22-HTL:**

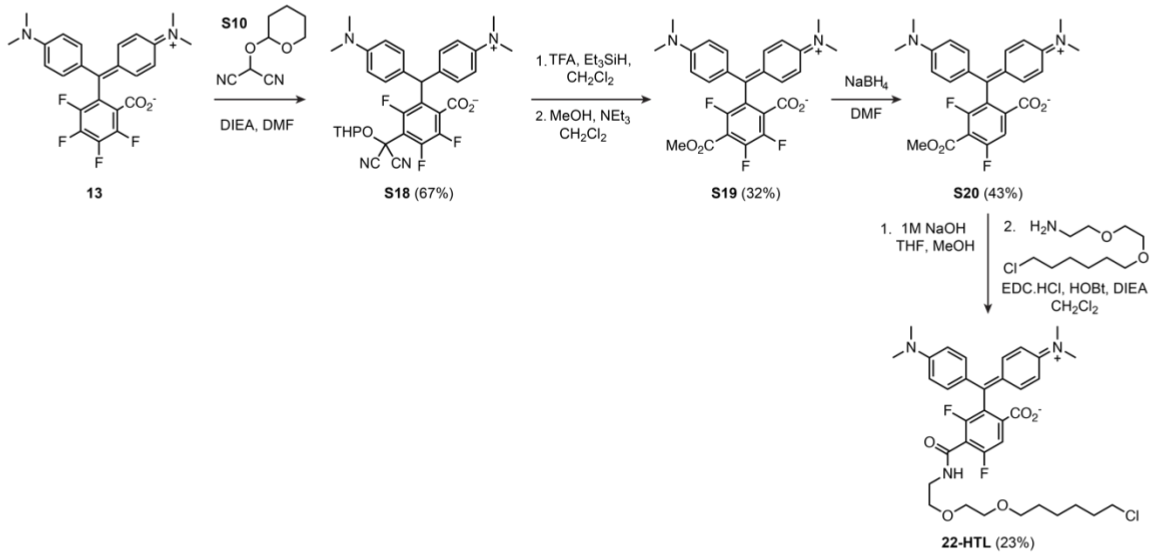

### General Experimental Information

#### Synthesis

Compounds **6**, **7**, **4-HTL**, **6-HTL, 7-HTL**, **13-HTL, 6-(MOM-MAC)-FMGL** were synthesized as previously reported[5] or gifted by the Lavis group (Janelia Research Campus, HHMI).

Commercial reagents were obtained from reputable suppliers (e.g. Merck, TCI, BLDpharm) and used as received. All solvents used for chemical reactions were of anhydrous grade, purchased in septum-sealed bottles stored under an inert atmosphere. Reactions under an inert atmosphere were sealed with septa and purged under vacuum/argon on a Schlenk line. Reactions were performed in round-bottomed flasks or septum-sealed vials.

Reactions were monitored by thin layer chromatography (TLC) on precoated aluminium plates (silica gel 60 F254, 200 µm thickness) or by LCMS (Agilent 1260 Infinity II; ZORBAX SB-C18 18 µm 80 Å, 2.1x50 mm column, 5 to 20 µL injection, 5–95% $CH_3CN/H_2O$ gradient with constant 0.1% v/v $HCO_2H$ additive, 8 to 10 min run, 0.5 mL/min flow, ESI positive ion mode, detection at 254 nm). TLC plates were visualized by UV illumination (254 nm) or developed with stains (e.g. cerium ammonium molybdate or Seebach stain). Compounds were purified by flash chromatography on an automated purification system (Biotage Isolera One) using pre-packed silica cartridges (Biotage Sfär Duo, 60 Å pores, 60 µm particles size) or by preparative HPLC (Agilent 1260 Infinity II, Phenomenex Gemini NX 21.2x150 mm, 5 µm C18 column). High-resolution mass spectrometry was performed by the Metabolomics Core Facility at EMBL Heidelberg.

NMR spectra were recorded on a 400 MHz spectrometer (Bruker 400 UltraShield) in deuterated solvents. All spectra were recorded at 298 K. $^1$H and $^{13}$C chemical shifts (δ) were referenced to residual solvent peaks, and $^{19}$F chemical shifts (δ) were referenced to $CFCl_3$. Data for $^1$H NMR spectra are reported as chemical shift (δ in ppm), multiplicity (s = singlet, d = doublet, t = triplet, q = quartet, p = pentuplet, dd = doublet of doublets, ddd= doublet of doublets of doublets, dt = doublet of triplets, tt = triplet of triplets, dtt = doublet of triplets of triplets, m = multiplet, br s = broad signal), coupling constant (*J* in Hz), integration. Data for $^{13}$C NMR spectra are reported by chemical shift (δ in ppm) with hydrogen substitution (C, CH, $CH_2$, $CH_3$) information obtained from DEPT spectra. Data for $^{19}$F NMR are reported by chemical shift (δ in ppm) and coupling constant (*J* in Hz). Data was processed using Mnova from Mestrelab. The $^{13}$C NMR spectra are not reported for compounds containing fluorinated aryl rings due to complex couplings. Determination of sample concentration for UV-Vis and fluorescence spectroscopy was performed by $^1$H NMR of samples in DMSO-$d_6$ containing 5 mM DMF for relative integration.

#### Cloning, Protein Expression and Purification

pET51b-HaloTag7 (Addgene, 167266), pBR322-HaloTag9 (Addgene, 169324) and pET51b-rHCaMP (Addgene, 187106) plasmids were received as gifts from Kai Johnsson (MPI for Medical Research, Heidelberg). pRSET-HaloCaMP1a-EGFP (Addgene, 138327) and pRSET-HaloCaMP1b-EGFP (Addgene, 138328) were received as gifts from Eric Schreiter (Janelia Research Campus, HHMI). The HaloCaMP1a and HaloCaMP1b inserts were cloned via Gibson assembly into the pET51b backbone for controlled protein overexpression. Sanger sequencing was used to verify DNA sequences in the obtained plasmid constructs. The pDuEx2-NIRGECO1 plasmid (Addgene, 113680) was used for NIR-GECO1 insert amplification followed by Gibson assembly into the pET51b backbone. For the preparation of viruses, pAAV-HaloTag7-EGFP plasmid was a gift from Eric Schreiter (Janelia Research Campus, HHMI) and adeno-associated viruses (AAV1) were prepared by Janelia Virus Services or the EMBL Rome Genetic & Viral Engineering Facility.

HaloTag7, HaloCaMP1a and HaloCaMP1b proteins were expressed and purified as previously described.[6] In summary, pET51b plasmids containing either HaloTag7, HaloCaMP1a-EGFP or HaloCaMP1b-EGFP, were transformed into T7 Express Competent *E. coli* cells (NEB). Cell colonies were grown in LB medium containing ampicillin (100 µg·mL$^{-1}$) overnight at 37°C at 200 r.p.m. The starting culture was diluted 100-fold in LB containing ampicillin (100 µg·mL$^{-1}$) and incubated at 37°C, 200 r.p.m until the O.D. reached 0.4-0.6. Protein expression was then induced by IPTG (1 mM) and the culture incubated at 18°C, 200 r.p.m for 14-20 h. After centrifugation, cell pellets were lysed [50 mM Tris-HCl, pH = 8.0, 300 mM NaCl, 5 mM imidazole, 5 mM β-mercaptoethanol, protease inhibitor (cOmplete, Roche, 25x stock), 0.01 mg·mL$^{-1}$ DNase, 5 mM $MgCl_2$, 5 mg·mL$^{-1}$ n-octyl-β-D-glucopyranoside] in a microfluidizer and the lysate was clarified by centrifugation. Purification from lysate was achieved by nickel-affinity chromatography using HisTrap™ HP column [(GE Healthcare), equilibration buffer 50 mM Tris-HCl pH 8.0, 300 mM NaCl, 10-20 mM imidazole, 5 mM β-mercaptoethanol; elution buffer 50 mM Tris-HCl pH 8.0, 100mM NaCl, 50-250 mM imidazole (stepwise gradient), 5 mM β-mercaptoethanol]. Selected protein fractions were then pooled, followed by size exclusion chromatography using a Superdex 200 16/600 column (GE Healthcare) at a flow rate of 0.5 mL·min$^{-1}$ in 50 mM Tris-HCl, 100 mM NaCl, pH = 7.4. Protein concentration was estimated using the extinction coefficient of EGFP ($\varepsilon$(488 nm) = 55900 M$^{-1}$·cm$^{-1}$) or using the extinction coefficient at 280 nm calculated from the protein sequence (Expasy).

HaloTag9 and rHCaMP were expressed as described above. Purification from lysate was achieved by gravity nickel-affinity chromatography using PureCube Ni-NTA Agarose (Cube Biotech), and the same buffers as above except the elution buffer (50 mM Tris-HCl pH 8.0, 100mM NaCl, 500 mM imidazole), followed by PD-10 desalting column (Cytiva) for buffer exchange to 50 mM Tris-HCl, 100 mM NaCl, pH 7.4. Protein concentration was estimated using the extinction coefficient at 280 nm calculated from the protein sequence (Expasy).

NIR-GECO1[2] and mIFP[7] (Addgene 54620) were expressed and purified as for HaloTag9 and rHCaMP, except with an additional incubation step with the biliverdin cofactor. For this, 100 mL of cell lysates were incubated on ice with 300 µL of biliverdin (20 mg·mL$^{-1}$ stock solution in DMSO), for 1h on ice before loading onto the Ni-NTA Agarose column.

### UV-Vis, Fluorescence and Photoacoustic Spectroscopy

#### UV–Vis and Fluorescence Spectroscopy

All measurements were performed at room temperature (23±2°C). Compounds were prepared as stock solutions at 1 mM in DMSO which were diluted in solvents and buffers to a final DMSO concentration not exceeding 1% v/v. Spectroscopy in buffer solutions was performed using 1-cm path length polystyrene cuvettes (Ratiolab GmbH). With organic solvents 1-cm path length quartz cuvettes were used (Hellma Suprasil Quartz). Absorption spectra were recorded on a Cary Model 60 spectrophotometer (Agilent). Fluorescence spectra were recorded on a JASCO spectrofluorometer (FP-8500). Data was analysed and graphs were plotted using Prism (GraphPad). Absorption spectra, maximum absorption wavelength ($\lambda_{max}$), extinction coefficient (ε) and maximum emission wavelength ($\lambda_{em}$) were measured in triplicate under the following conditions, reported values for ε are the mean of 3 replicates. Spectra were recorded at 1.25 µM dye concentration, unless otherwise stated. For the free dyes, measurements were made in 10 mM HEPES pH 7.4. For the corresponding HaloTag ligands, 0.1 mg·ml$^{-1}$ of 3-((3-cholamidopropyl)dimethylammonio)-1-propanesulfonate (CHAPS) was added to the buffer. For measurements in the presence of proteins, the dye ligand was incubated with 1.5 equivalent (eq) of purified protein for at least 3 h at room temperature.

To further quantify the open-closed equilibrium, we measured absorption spectra of dyes at 5 µM in MeCN/$H_2O$ mixtures[8] and 0.1% v/v trifluoroacetic acid in 2,2,2-trifluoroethanol.[9] The Malachite Green scaffold did not fully open in either acidic or basic conditions.

Fluorescence quantum yields (Φ) were measured using an absolute quantum yield measurement system (Quantaurus, C11347, Hamamatsu). Measurements were carried out using dilute samples (A<0.1), self-absorption corrections were performed using the Quantaurus software. The fluorescence quantum yield of compound **13-HTL** bound to HaloTag7 was calculated precisely using the relative method with Oxazine 1 (Φ = 0.11 in EtOH) as a standard.[10]

The following notations are used:
$\lambda_{max}$ (nm): wavelength at maximal absorption.
$\lambda_{em}$ (nm): wavelength at maximal fluorescence emission.
$\lambda_{PA}$ (nm): wavelength used for photoacoustic excitation.
ε ($M^{-1} \cdot cm^{-1}$): molar extinction coefficient.
Φ: fluorescence quantum yield.
$K_d$ (nM, µM or mM): apparent dissociation constant of the calcium sensors.
$\Delta A/A_0$: change in absorption divided by the basal absorption.
$\Delta PA/PA_0$: change in photoacoustic signal divided by the basal photoacoustic signal.

**Photoacoustic Spectroscopy**

A custom-built multimodal spectroscopy setup was used to measure photoacoustic spectra. The setup consists of a water tank holding a standard 1-cm glass cuvette containing the sample. The system utilizes a 100 Hz pulsed tunable laser source (EVO I OPO, Innolas Laser GmbH) to excite photoacoustic signals, which are subsequently detected using a focused water-immersion transducer operating at a frequency of 0.5 MHz (V301, Olympus IMS). The acquired ultrasound signals were recorded via a data acquisition (DAQ) card with a sampling rate of 125 MSa/s (ATS9440, Alazar Technologies Inc.) and processed with a custom MATLAB code. The peak-to-peak value of the ultrasound waveform was used as a metric of photoacoustic signal. The photoacoustic spectra were measured by sweeping through the wavelength range of the excitation laser (420-709 nm) and acquiring photoacoustic signals at each wavelength. Each spectrum was corrected for changes in laser pulse energy with wavelength, and the background photoacoustic signal from a cuvette with corresponding buffer solution was subtracted to remove any signal from the glass cuvette. Single wavelength measurements were recorded at the maximum absorption wavelength and averaged for 1000 pulses.

For photostability measurements, the photoacoustic spectroscopy set-up was adapted into a well plate reader configuration. The laser light was adjusted to illuminate the entire well and photobleach the sample homogenously. Single wavelength photobleaching were recorded at the maximum absorption wavelength for 1 h at 100 Hz pulse repetition rate with average power ranging from 100 to 500 mW depending on the wavelength. The photoacoustic amplitudes were corrected for power fluctuations between pulses and normalised to initial values. The photobleaching data was then plotted against the cumulative deposited energy to compare bleaching rates of samples at different wavelengths where the laser power is different. 1 mL samples were prepared and absorption matched to ~0.065 $cm^{-1}$ using the Cary spectrometer. Absorption was measured again after irradiation for validation.

**Calcium Titrations and $K_d$ Determination.**

For the high affinity sensors, calcium titrations were performed using a commercial EGTA/Ca·EGTA buffer system (Invitrogen) to which 0.1 $mg \cdot ml^{-1}$ CHAPS was added, following the associated protocol. For the low affinity sensors, buffer solutions (30 mM MOPS, 100 mM KCl, pH 7.2) containing various calcium concentrations (0 µM, 20 µM, 100 µM, 200 µM, 500 µM, 1 mM, 2 mM, 5 mM, 10 mM, 20 mM, 50 mM, 100 mM, 500 mM) were prepared.

For calcium titrations of the HaloTag ligands in the presence of HaloCaMP protein, the dye was incubated with 1.5 eq of purified HaloCaMP protein for 3 h at room temperature. All the calcium titrations were performed in duplicate. $K_d$ values were obtained by fitting the curve of the absorption at $\lambda_{max}$ as a function of free [$Ca^{2+}$] with the Hill equation.

**Calcium Selectivity**

**13-HTL** (60 µM) was incubated with purified HaloCaMP1a and HaloCaMP1b (90 µM, 1.5 eq) at room temperature for 3 h. For HaloCaMP1a, the labelled protein was diluted to 1.25 µM in 30 mM MOPS pH 7.2, 100 mM KCl, 10 µM EGTA. $CaCl_2$ or $MgCl_2$ were added to final concentrations of either 500 mM $CaCl_2$, 5 mM $MgCl_2$ or 100 mM $MgCl_2$. $CaCl_2$ was then added to the $MgCl_2$ containing solutions to reach 500 mM $Ca^{2+}$ concentration.

Absorption was measured at each step. For HaloCaMP1b, the labelled protein was diluted to 1.25 µM in 30 mM MOPS pH 7.2, 100 mM KCl, 10 mM EGTA. $CaCl_2$ or $MgCl_2$ were added to a concentration of 1.5 mM (sufficient $Ca^{2+}$ concentration to reach the calcium-saturated state of the sensor). To the $MgCl_2$-containing solution, 1.5 mM $CaCl_2$ was subsequently added. All measurements were performed in duplicate.

#### pH Stability

Purified HaloCaMP1a and HaloCaMP1b (90 µM, 1.5 eq) and **13-HTL** (60 µM) were incubated at room temperature for 3 h. The labelled protein was diluted to 1.25 µM, into the following buffer systems: citrate (pH = 4.0, 5.0), phosphate (pH = 6.0, 7.0), Tris (pH = 8.0, 9.0), containing 150 mM NaCl and 0.1 mg·mL$^{-1}$ CHAPS. 10 mM EGTA was added for the calcium-free state, or excess $CaCl_2$ was added for the calcium-bound state (i.e. 500 mM for HaloCaMP1a or 10 mM for HaloCaMP1b). Absorbance spectra were recorded, all measurements were performed in duplicate.

### Microscopy and Tomography

#### Fluorescence Microscopy in Cultured Cells

Widefield imaging was performed at the Advanced Light Microscopy Facility (EMBL Heidelberg), on a Nikon Ti-E microscope equipped with a Spectra X light engine (Lumencore) with a 20x objective (CFI P-Apo 20x Lambda/ 0.75/ 1,00) and imaged onto a scientific complementary metal–oxide–semiconductor camera (pco.edge 4.2 CL). A quad bandpass filter cube was used to image GFP (excitation 485/20, emission 525/50), **JF549-HTL** (excitation 531/40, emission 593/40) and far-red ligands (excitation 650/13, emission 692/40).

#### General Cell Culture Methods

U2OS cells stably expressing HaloTag–EGFP fusion protein (gift from the Johnsson group, MPI for Medical Research, Heidelberg) were cultured in Dulbecco's modified Eagle medium [(DMEM, high glucose (4.5 g·L$^{-1}$) with phenol red, supplemented with 10% v/v fetal bovine serum, penicillin (100 units·mL$^{-1}$), streptomycin (100 µg·mL$^{-1}$), hygromycin B (100 µg·mL$^{-1}$), 2 mM L-Glutamine (Gibco) and 1 mM sodium pyruvate], and maintained at 37°C in a humidified 5% v/v $CO_2$ environment. All labelled cells were imaged 18–24 h post-plating.

U2OS HaloTag7-EGFP cells were seeded in a chambered coverslip with 8 individual wells (Ibidi, 80806). The next day cells were incubated with 1 µM HaloTag ligand **(9-HTL**, **13-HTL**, **14-HTL** or positive control **JF549-HTL**) for 2 h at 37°C, washed three times with imaging buffer [DMEM low glucose (1 g·L$^{-1}$) without phenol red, supplemented with 10% v/v fetal bovine serum, 2 mM L-Glutamine (Gibco) and 1 mM sodium pyruvate]. Following this, 0.5 µM of **JF549-HTL** was added to the medium, and the cells were incubated for 30 min more at 37°C. The cells were subsequently washed twice with PBS, incubated for 15 minutes at 37°C and then washed three times with imaging buffer before widefield fluorescence imaging as described above.

#### Photoacoustic Tomography

A custom-built photoacoustic tomography setup based on a transparent Fabry-Pérot ultrasound sensor was used for PA imaging. Briefly, the setup is based on a 100 Hz pulsed tunable laser source (EVO I OPO, Innolas Laser GmbH) that excites the photoacoustic signal in a 420-709 nm range. The ultrasound waves are then detected by a home-made Fabry-Pérot sensor[11] interrogated with a 1520-1620 nm CW laser (Venturi TLB-8800, Newport). The Fabry-Pérot sensor was raster-scanned with a 15x15 mm FOV and 50 µm step resulting in a 301x301 ultrasound sampling points. The photoacoustic image was then reconstructed with either a back-projection or time-reversal algorithm from a k-Wave toolbox for MATLAB.[12]

For the tissue-mimicking phantoms, protein solutions at 50 µM were loaded in PE tubing (ID 0.86 mm, OD 1.27 mm, Hugo Sachs Elektronik), secured in a 3D-printed holder and fixed in a petri dish. They were submerged in coupling media of water or milk-water mixtures (10% or 60% whole cow’s milk). The photoacoustic data was acquired at respective excitation wavelengths and the resulting image was formed with a k-Wave reconstruction algorithm and then exported to ImageJ, where the maximum intensity projections of a 3D image were extracted for xy, xz and yz views. The maximum intensity projections were then analysed with a line profile to quantify the values from each sample.

#### Mouse *in vivo* Experiments

This work followed the European Communities Council Directive (2010/63/EU) to minimize animal pain and discomfort. All procedures described in this paper were approved by EMBL's committee for animal welfare and institutional animal care and use, under license 22-004_HD_RP. Experiments were performed on 12-week-old male C57BL/6 mice from the EMBL Heidelberg core colonies or Charles River Laboratories. During the course of the study, mice were housed in groups of 1–5 in makrolon type 2L or 3H cages, in ventilated racks at room temperature and 50% humidity while kept in a 12/12 h light/dark cycle. Food and water were available *ad libitum*.

For neuronal-specific expression of HaloTag7-EGFP, AAV1-synapsin1-NES-HaloTag7-EGFP ($6.0\times10^{13}$ viral genomes per mL). For stereotaxic injection of AAVs, mice were anesthetized with isoflurane (Baxter) vapor mixed with $O_2$ (5% for induction and 1–1.5% for maintenance). Eye ointment was applied (Bepanthen, Bayer) and Xyolcain 1% (AstraZeneca) was subcutaneously injected in the scalp for preincisional local anesthesia. During anesthesia, body temperature was monitored and maintained with a RWD ThermoStar system. A small incision was produced in the scalp to expose the dorsal cranium and, using the bregma as a landmark, a small burr hole centered at ML 1.3 mm, AP -2 mm was made with a dental drill (Microtorque, Harvard Apparatus). AAV injections were performed with a RWD 68803 stereotaxic fame equipped with a 68055 mouse adapter in the hippocampus and thalamus on the left hemisphere at depths -1.2 mm and 3 mm respectively, using a syringe at a rate of ~4 $\mu L \cdot h^{-1}$. Approximatively 300 nL were injected per spot. Mice were sutured, meloxican administered subcutaneously (Metacan, Boehringer Ingelheim) dosed at 1 $\mu g \cdot g^{-1}$ for pain relief and single housed after the viral injection surgery and had a recovery period of at least 4 weeks before further experiments for HaloTag7-EGFP expression.

A 15 mM solution of **13-HTL or 14-HTL** in DMSO was prepared. 2 µL was added to 2 µL of F127 pluronic acid in DMSO (20% w/w), and 16 µL of sterile saline was added to give a final concentration of 1.5 mM of dye ligand. 2 µL (3 nmol) of this dye ligand solution was administered via intracerebroventricular injection (N = 3 mice for **13-HTL** and N = 2 mice for **14-HTL**). 24 hours after dye injection, animals were sacrificed with $CO_2$ and transcardially perfused with ~35 mL of PBS without $Ca^{2+}$ and $Mg^{2+}$ to remove blood from tissues. The perfused brain was then acutely dissected fresh in ice cold PBS without $Ca^{2+}$ and $Mg^{2+}$. Immediately after dissection the brain was imaged in our photoacoustic tomography setup by gluing to the bottom of a petri dish and submersion in PBS without $Ca^{2+}$ and $Mg^{2+}$ for acoustic coupling.

After PAI, the brain was retrieved from the photoacoustic tomography setup and directly sliced 300 µm thick with a vibratome (Leica VT1200) in PBS without $Ca^{2+}$ and $Mg^{2+}$. Based on the mouse brain atlas[13], and using morphological landmarks, including the splitting of coronal hemispheres, each slice was mapped to their corresponding coronal section in the PAT volume.

#### Fluorescence Microscopy and Photoacoustic Imaging of Brain Slices

Alternatively, after PAI, brains were fixated by immersion in PBS with 4% paraformaldehyde for 72 h at 4ºC. The fixed brain was washed three times with PBS and then sliced (200-300 µm thick) with a vibratome (Leica VT1200) in PBS without $Ca^{2+}$ and $Mg^{2+}$. The slices were imaged via widefield fluorescence imaging (as described above) with the 4x objective in transmission, GFP channel and Cy5 channel.

For dye labelling of brain slices, the slices were soaked in a bath of PBS containing 10 µM of **13-HTL** for 2 h at 37°C, rotating gently on a tilt shaker to ensure homogenous labelling (N = 3 mice). The slices were washed with PBS without $Ca^{2+}$ and $Mg^{2+}$ (3 x 20 min) at 37°C before photoacoustic imaging.

For photoacoustic imaging the brain slices were fixed in low-gelling temperature agarose (A9045-10G, Sigma) in a petri dish. The sample was submerged in PBS without $Ca^{2+}$ and $Mg^{2+}$ for acoustic coupling and placed in the photoacoustic tomography setup for imaging. The wide-field fluorescence image of the agarose-fixed slice was taken in the GFP channel for co-registering the PAI image with the original fluorescence images.

### X-Ray Crystallography

To confirm the position of the two fluorine atoms on compound **22-HTL**, we solved a crystal structure of the isolated intermediate **S20**. Suitable crystals were grown by vapour diffusion of cyclohexane in a solution of MeOH/$CH_2Cl_2$, and X-Ray diffraction data was collected by Dr. Dieter Schollmeyer at the Johannes Gutenberg Universität Mainz.

X-Ray diffraction revealed the structure of **S20** (CCDC Deposition Number 2336812), the major product present at 90%, bearing the hydrogen at the 4-position (C25 in the atom numbering scheme, Figures C1 and C2). In addition, the crystal revealed the presence of another regioisomer **S20'** present at 10% proportion, and bearing the hydrogen at the 5-position. The two regioisomers could not be fully separated at that stage, and the 90:10 mixture was used for the synthesis of the HaloTag ligand **22-HTL**. This compound could be purified as a single isomer, as shown by the $^{1}$H NMR spectrum and LCMS trace, and the NMR characteristics (chemical shifts and coupling constants for the fluorines and the hydrogen on the bottom ring) are in good accordance with that of compound **S20**, confirming the chemical structure proposed for **22-HTL**.

**Figure C1**. Chemical structures and proportions of the isomers observed in the single crystal of compound **S20**.

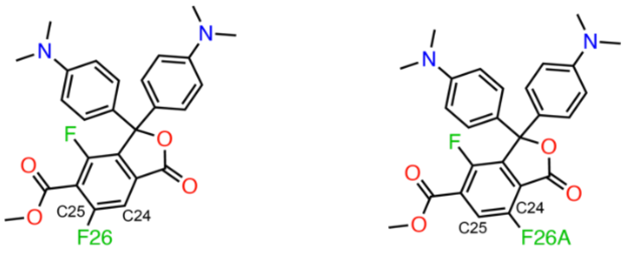

**S20** (major - 90%)

**S20'** (minor - 10%)

**Figure C2**. Molecular structure with atom numbering scheme. The crystal was composed of 90% of compound bearing fluorine at C25 (= F26), and 10% of compound bearing fluorine at C24 (= F26A).

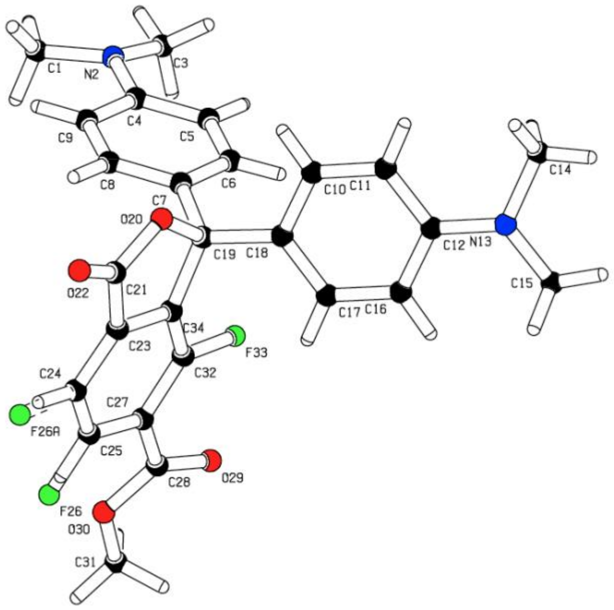

**Table C1.** Crystal data and structure refinement

| | | |
|---|---|---|
| Identification code | f2mgome | |
| Empirical formula | $C_{26}H_{24}F_2N_2O_4$ | |
| moiety formula | $C_{26}H_{24}F_2N_2O_4$ | |
| Formula weight | 466.47 | |
| Temperature | 120(2) K | |
| Wavelength, radiation type | 1.54178Å, CuKα | |
| Diffractometer | STOE STADIVARI | |
| Crystal system | Monoclinic | |
| Space group name, number | P $2_1$/c, (14) | |
| Unit cell dimensions | a = 19.0356(5) Å | α = 90° |
| | b = 6.76850(10) Å | β = 118.528(2)° |
| | c = 19.9025(5) Å | γ = 90° |
| Volume | 2252.94(10) $Å^3$ | |
| Number of reflections | 15982 | |
| and range used for lattice parameters | 4.45° <=θ<= 68.46° | |
| Z | 4 | |
| Density (calculated) | 1.375 $Mg/m^3$ | |
| Absorption coefficient | 0.872 $mm^{-1}$ | |
| Absorption correction | Integration | |
| Max. and min. transmission | 0.9773 and 0.9080 | |
| F(000) | 976 | |
| Crystal size, colour and form | 0.030 x 0.060 x 0.120 $mm^3$, colorless needle | |
| Theta range for data collection | 4.449 to 68.725°. | |
| Index ranges | -22<=h<=22, -7<=k<=8, -24<=l<=23 | |
| Number of reflections: | | |
| collected | 24162 | |
| independent | 4119 [R(int) = 0.0426] | |
| observed [I>2sigma(I)] | 2870 | |
| Completeness to theta = 67.7° | 99.5 % | |
| Refinement method | Full-matrix least-squares on $F^2$ | |
| Data / restraints / parameters | 4119 / 1 / 321 | |
| Goodness-of-fit on $F^2$ | 0.977 | |
| Final R indices [I>2sigma(I)] | R1 = 0.0421, wR2 = 0.1052 | |
| R indices (all data) | R1 = 0.0638, wR2 = 0.1126 | |
| Largest diff. peak and hole | 0.365 and -0.275 $eÅ^{-3}$ | |
| Remark | position of F26 split at C24 (10%) and C25 (90%) | |

### Synthetic Procedures and Characterizations

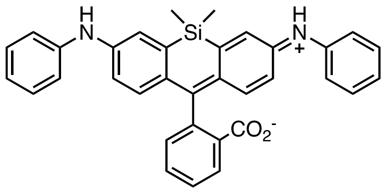

**(1):** Si-fluorescein ditriflate (40.1 mg, 0.063 mmol, 1 eq), $Pd_2dba_3$ (5.7 mg, 0.006 mmol, 0.1 eq), XPhos (9.0 mg, 0.019 mmol, 0.3 eq) and $Cs_2CO_3$ (57.1 mg, 0.175 mmol, 2.8 eq) were loaded into a vial. The vial was sealed and evacuated/backfilled with argon. 1,4-Dioxane (1 mL) and subsequently aniline (17 µL, 0.150 mmol, 2.4 eq) were added then the reaction mixture was stirred at 100°C for 3 h. After cooling to room temperature, the reaction mixture was filtered through Celite, washed with $CH_2Cl_2$ and the solvent was evaporated under reduced pressure. Purification was performed by silica gel column chromatography (5% MeOH/$CH_2Cl_2$) followed by reverse phase HPLC (10–95% $CH_3CN/H_2O$, linear gradient with constant 0.1% v/v TFA additive). The pooled HPLC product fractions were partially concentrated to remove $CH_3CN$ then lyophilized overnight to afford the title compound as a blue-green solid (26.0 mg, 65%, TFA salt). $^1H$ NMR ($CDCl_3$, 400 MHz) δ 8.20 (br s, 2H), 8.09 (d, *J* = 7.6 Hz, 1H), 7.71, (t, *J* = 7.6 Hz, 1H), 7.60 (t, *J* = 7.6 Hz, 1H), 7.38 – 7.28 (m, 7H), 7.15 (d, *J* = 7.7 Hz, 4H), 7.07 (t, *J* = 7.7 Hz, 2H), 6.91 – 6.83 (m, 4H), 0.58 (s, 3H), 0.47 (s, 3H). $^{13}C$ NMR ($CDCl_3$, 101 MHz) δ 170.7 (C), 153.6 (C), 143.0 (C), 142.3 (C), 137.8 (C), 136.0 (C), 134.0 (CH), 129.6 (CH), 129.2 (CH), 128.6 (CH), 126.9 (C), 126.2 (CH), 124.9 (CH), 122.1 (CH), 121.9 (CH), 118.6 (CH), 117.8 (CH), 77.4 (C), 0.3 (Si-$CH_3$), -1.6 (Si-$CH_3$). Analytical HPLC: $t_R$ = 5.4 min, 98% purity (5–95% $CH_3CN$ /$H_2O$, gradient with constant 0.1% v/v formic acid additive, 10 min run, 0.5 mL/min flow, detection at 254 nm. HRMS (ESI) calcd for $C_{34}H_{29}N_2O_2Si$ $[M+H]^+$ 525.1993, found 525.2001.

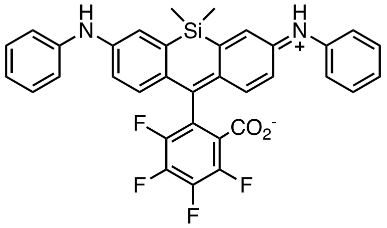

**(2):** $F_4$-Si-fluorescein ditriflate (42.6 mg, 0.060 mmol, 1 eq), $Pd_2dba_3$ (5.5 mg, 0.006 mmol, 0.1 eq), XPhos (8.6 mg, 0.018 mmol, 0.3 eq) and $Cs_2CO_3$ (54.8 mg, 0.168 mmol, 2.8 eq) were loaded into a vial. The vial was sealed and evacuated/backfilled with argon. 1,4-Dioxane (1 mL) and subsequently aniline (13 µL, 0.144 mmol, 2.4 eq) were added then the reaction mixture was stirred at 100°C for 19 h. After cooling to room temperature, the reaction mixture was filtered through Celite, washed with EtOAc and the solvent was evaporated under reduced pressure. Purification was performed by silica gel column chromatography (Biotage Sfär Duo 5 g, 0–15% EtOAc/cyclohexane, linear gradient) followed by reverse phase HPLC (30–95% $CH_3CN/H_2O$, linear gradient with constant 0.1% v/v TFA additive). The pooled HPLC product fractions were partially concentrated to remove $CH_3CN$ then lyophilized overnight to afford the title compound as a blue solid (0.89 mg, 2%, TFA salt). $^1H$ NMR (400 MHz, $(CD_3)_2CO$) δ 7.71 (s, 2H), 7.51 (d, *J* = 2.6 Hz, 2H), 7.32 – 7.25 (m, 4H), 7.22 – 7.16 (m, 4H), 7.07 (dd, *J* = 8.7, 2.6 Hz, 2H), 6.99 (dd, *J* = 8.6, 1.6 Hz, 2H), 6.93 (tt, *J* = 7.3, 1.2 Hz, 2H), 0.59 (s, 3H), 0.54 (s, 3H). $^{19}F$ NMR (376 MHz, $(CD_3)_2CO$) δ -139.83 – -139.97 (m, 1F), -141.76 – -141.91 (m, 1F), -146.03 – -146.18 (m, 1F), -154.05 – -154.20 (m, 1F). Analytical HPLC: $t_R$ = 4.8 min, >99% purity (5–95% $CH_3CN/H_2O$, gradient with constant 0.1% v/v formic acid additive, 8 min run, 0.5 mL/min flow, detection at 254 nm. HRMS (ESI) calcd for $C_{34}H_{24}F_4N_2O_2Si$ $[M+H]^+$ 597.1616, found 597.1600.

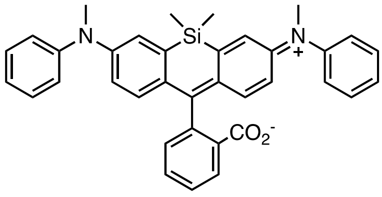

**(3):** Si-fluorescein ditriflate (38.5 mg, 0.060 mmol, 1 eq), $Pd_2dba_3$ (5.5 mg, 0.006 mmol, 0.1 eq), XPhos (8.6 mg, 0.018 mmol, 0.3 eq) and $Cs_2CO_3$ (54.9 mg, 0.169 mmol, 2.8 eq) were loaded into a vial. The vial was sealed and evacuated/backfilled with argon. 1,4-Dioxane (1 mL) and subsequently *N*-methylaniline (16 µL, 0.144 mmol, 2.4 eq) were added then the reaction mixture was stirred at 100°C for 3 h. After cooling to room temperature, the reaction mixture was filtered through Celite, washed with $CH_2Cl_2$ and the solvent was evaporated under reduced pressure. Purification by silica gel column chromatography (10% EtOAc/cyclohexane) afforded the title compound as a blue-green solid (22.5 mg, 68%). $^1H$ NMR ($CDCl_3$, 400 MHz) δ 7.98 (d, *J* = 7.6 Hz, 1H), 7.68 (t, *J* = 7.6 Hz, 1H), 7.57 (t, *J* = 7.6 Hz, 1H), 7.38 (d, *J* = 7.6 Hz, 1H), 7.30 (t, *J* = 7.7 Hz, 4H), 7.25 (d, *J* = 2.8 Hz, 2H), 7.09 (d, *J* = 7.7 Hz, 4H), 7.04 (t, *J* = 7.7 Hz, 2H), 6.81 (d, *J* = 8.8 Hz, 2H), 6.77 (dd, *J* = 8.8, 2.8 Hz, 2H), 3.33 (s, 6H), 0.57 (s, 3H), 0.54 (s, 3H). $^{13}C$ NMR $(CD_3)_2CO$, 101 MHz) δ 170.4 (C), 155.0 (C), 149.3 (C), 149.1 (C), 137.4 (C), 136.5 (C), 135.1 (CH), 130.3 (CH), 130.1 (CH), 128.5 (CH), 127.1 (C), 126.3 (CH), 125.5 (CH), 123.8 (CH), 123.5 (CH), 123.4 (CH), 119.9 (CH), 91.3 (C), 40.3 (N-$CH_3$), 0.2 (Si-$CH_3$), -1.5 (Si-$CH_3$). Analytical HPLC: $t_R$ = 4.7 min, >99% purity (5–95% $CH_3CN/H_2O$, gradient with constant 0.1% v/v formic acid additive, 10 min run, 0.5 mL/min flow, detection at 254 nm). HRMS (ESI) calcd for $C_{36}H_{33}N_2O_2Si$ $[M+H]^+$ 553.2306, found 553.2314.

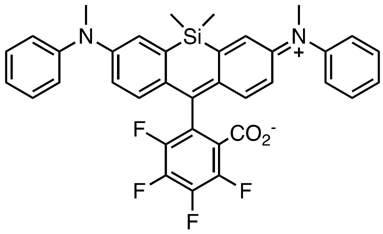

**(4):** $F_4$-Si-fluorescein ditriflate (33.5 mg, 0.047 mmol, 1 eq), $Pd_2dba_3$ (4.3 mg, 0.005 mmol, 0.1 eq), XPhos (6.8 mg, 0.014 mmol, 0.3 eq) and $Cs_2CO_3$ (43.1 mg, 0.132 mmol, 2.8 eq) were loaded into a vial. The vial was sealed and evacuated/backfilled with argon. 1,4-Dioxane (1 mL) and subsequently N-methylaniline (12 µL, 0.113 mmol, 2.4 eq) were added then the reaction mixture was stirred at 100°C for 3 h. After cooling to room temperature, the reaction mixture was filtered through Celite, washed with $CH_2Cl_2$ and the solvent was evaporated under reduced pressure. Purification was performed by silica gel column chromatography (0–10% EtOAc/cyclohexane) followed by reverse phase HPLC (30–95% $CH_3CN/H_2O$, linear gradient with constant 0.1% v/v TFA additive). The pooled HPLC product fractions were partially concentrated to remove $CH_3CN$ then lyophilized overnight to afford the title compound as a blue-green solid (7.7 mg, 22%, TFA salt). $^1H$ NMR ($CDCl_3$, 400 MHz) δ 7.39 (t, *J* = 7.9 Hz, 4H), 7.22 – 7.15 (m, 6H), 7.14 (d, *J* = 2.8 Hz, 2H), 6.86 (d, *J* = 8.9 Hz, 2H), 6.74 (dd, *J* = 8.9, 2.8 Hz, 2H), 3.42 (s, 6H), 0.47 (s, 3H), 0.45 (s, 3H). $^{19}F$ NMR ($CDCl_3$, 376 MHz) δ -137.5 – -137.7 (m, 1F), -138.6 (td, *J* = 19.6, 3.6 Hz, 1F), -145.7 – -146.0 (m, 1F), -151.6 (td, *J* = 19.6 Hz, 3.6 Hz, 1F). HRMS (ESI) calcd for $C_{36}H_{29}F_4N_2O_2Si$ $[M+H]^+$ 625.1929, found 625.1920.

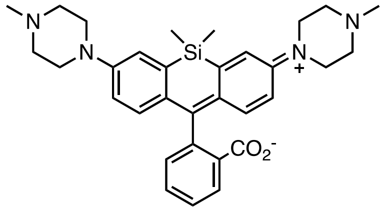

**(5):** Si-fluorescein ditriflate (41.0 mg, 0.048 mmol, 1 eq), $Pd(OAc)_2$ (2.2 mg, 0.010 mmol, 0.2 eq), BINAP (9.1 mg, 0.015 mmol, 0.3 eq) and $Cs_2CO_3$ (44.4 mg, 0.136 mmol, 2.8 eq) were loaded into a vial. The vial was sealed and evacuated/backfilled with argon. Toluene (1 mL) and subsequently 1-methylpiperazine (13 µL, 0.117 mmol, 2.4 eq) were added and the reaction mixture was stirred at 100°C for 48 h. After cooling to room temperature, the reaction mixture was filtered through Celite, washed with MeOH and the solvent was evaporated under reduced pressure. Purification by silica gel column chromatography (10% MeOH/$CH_2Cl_2$) afforded the title compound as a pale green solid (23.8 mg, 92%). $^1$H NMR (400 MHz, $CD_3CN$) δ 7.93 (d, *J* = 7.6 Hz, 1H), 7.71 (td, *J* = 7.5, 1.2 Hz, 1H), 7.61 (td, *J* = 7.5, 0.9 Hz, 1H), 7.28 – 7.32 (m, 2H), 7.22 (d, *J* = 7.7 Hz, 1H), 6.83 – 6.89 (m, 4H), 3.82 (br s, 4H), 3.49 (br s, 4H), 3.12 (m, 8H), 2.80 (s, 6H), 0.65 (s, 3H), 0.56 (s, 3H). $^{13}$C NMR ($CD_3CN$, 101 MHz) δ 171.2 (CO), 155.6 (C), 149.9 (C), 137.5 (C), 136.6 (C), 135.7 (CH), 130.3 (CH), 129.0 (CH), 126.6 (CH), 126.4 (C), 125.0 (CH), 121.7 (CH), 118.5 (CH), 91.1 (C), 53.7 ($CH_2$), 46.5 ($CH_2$), 43.5 ($CH_3$), 0.0 (Si-$CH_3$), -1.1 (Si-$CH_3$). Analytical HPLC: $t_R$ = 2.5 min, >99% purity, (5–95% $CH_3CN/H_2O$, gradient with constant 0.1% formic acid additive, 8 min run, 0.5 mL/min flow, UV detection at 254 nm). HRMS (ESI) calcd for $C_{32}H_{39}N_4O_2Si$ $[M+H]^+$ 539.2837, found 539.2825.

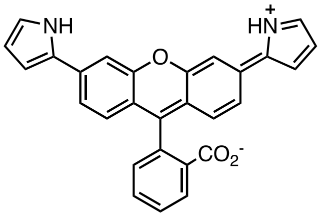

**(8):** Fluorescein ditriflate (92.4 mg, 0.155 mmol, 1 eq), *N*-Boc-pyrrole-2-boronic acid pinacol ester (136 mg, 0.465 mmol, 3.0 eq), $Pd(PPh_3)_4$ (10.7 mg, 0.009 mmol, 0.06 eq) and KOAc (91.2 mg, 0.930 mmol, 6.0 eq) were loaded into a vial. The vial was sealed and evacuated/backfilled with argon. A mixture of 1,4-dioxane/$H_2O$ (2/1, 2.85 mL) was added and the reaction mixture was stirred at 100°C for 23 h. After cooling to room temperature, the reaction mixture was extracted with $CHCl_3$. The organic layers were combined, washed with brine, dried over $Na_2SO_4$ and the solvent was evaporated under reduced pressure. Silica gel column chromatography (Biotage Sfär Duo 5 g, 0–30% $CH_2Cl_2$/cyclohexane, linear gradient) afforded the title compound as a blue solid (37.7 mg, 56%). $^1$H NMR (400 MHz, $CDCl_3$) δ 8.87 (s, 2H), 8.06 – 8.00 (m, 1H), 7.69 – 7.58 (m, 2H), 7.31 (d, *J* = 1.8 Hz, 2H), 7.16 – 7.07 (m, 3H), 6.87 – 6.91 (m, 2H), 6.73 (d, *J* = 8.3 Hz, 2H), 6.58 – 6.52 (m, 2H), 6.28 (q, *J* = 2.8 Hz, 2H). $^{13}$C NMR (101 MHz, $CDCl_3$) δ 170.1 (CO), 153.1 (C), 151.8 (C), 135.6 (C), 135.4 (CH), 130.7 (C), 130.0 (CH), 128.6 (CH), 126.4 (C), 125.4 (CH), 124.2 (CH), 120.3 (CH), 119.6 (CH), 116.0 (C), 111.5 (CH), 110.5 (CH), 107.7 (CH), 75.2 (C). Analytical HPLC: $t_R$ = 4.3 min, >99% purity (5–95% $CH_3CN/H_2O$, gradient with constant 0.1% formic acid additive, 8 min run, 0.5 mL/min flow, UV detection at 254 nm. HRMS (ESI) calcd for $C_{28}H_{19}N_2O_3$ $[M+H]^+$ 431.1390, found 431.1388.

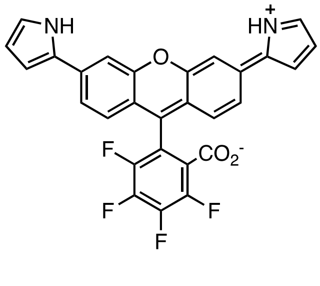

**(9):** $F_4$-Fluorescein ditriflate (30.8 mg, 0.046 mmol, 1 eq), *N*-Boc-pyrrole-2-boronic acid pinacol ester (40.6 mg, 0.139 mmol, 3.0 eq), $Pd(PPh_3)_4$ (3.2 mg, 0.003 mmol, 0.06 eq) and KOAc (27.2 mg, 0.277 mmol, 6.0 eq) were loaded into a vial. The vial was sealed and evacuated/backfilled with argon. A mixture of 1,4-dioxane/$H_2O$ (2/1, 1.05 mL) was added and the reaction mixture was stirred at 100°C for 46 h. After cooling to room temperature, the reaction mixture was extracted with $CHCl_3$. The organic layers were combined, washed with brine, dried over $Na_2SO_4$ and the solvent was evaporated under reduced pressure. Silica gel column chromatography (30% EtOAc/cyclohexane) afforded the title compound as a blue solid (15.0 mg, 65%). $^1$H NMR (400 MHz, $(CD_3)_2CO$) δ 10.72 (br s, 2H), 7.62 (d, *J* = 1.8 Hz, 2H), 7.48 (dd, *J* = 8.4, 1.8 Hz, 2H), 7.22 (d, *J* = 8.3 Hz, 2H), 6.96 (td, *J* = 2.7, 1.4 Hz, 2H), 6.70 – 6.73 (m, 2H), 6.20 – 6.24 (m, *J* = 3.5, 2.4 Hz, 2H). $^{19}$F NMR (376 MHz, $(CD_3)_2CO$) δ -

139.98 (td, *J* = 19.6, 9.3 Hz, 1F), -143.84 (td, *J* = 19.3, 4.3 Hz, 1F), -144.45 (td, *J* = 18.8, 9.0 Hz, 1F), -152.37 (td, *J* = 18.6, 4.3 Hz, 1F). Analytical HPLC: $t_R$ = 4.9 min, 98% purity, 5–95% $CH_3CN/H_2O$, gradient with constant 0.1% formic acid additive, 10 min run, 0.5 mL/min flow, UV detection at 254 nm. HRMS (ESI) calcd for $C_{28}H_{15}F_4N_2O_3$ $[M+H]^+$ 503.1013, found 503.1017.

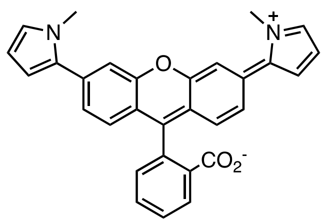

**(10):** Fluorescein ditriflate (77.7 mg, 0.130 mmol, 1 eq), 1-methyl-2-pyrroleboronic acid pinacol ester (81.0 mg, 0.391 mmol, 3.0 eq), $Pd(PPh_3)_4$ (9.0 mg, 0.007 mmol, 0.06 eq) and KOAc (76.7 mg, 0.781 mmol, 6.0 eq) were loaded into a vial. The vial was sealed and evacuated/backfilled with argon. A mixture of 1,4-dioxane/$H_2O$ (2/1, 2.40 mL) was added and the reaction mixture was stirred at 100°C for 23 h. After cooling to room temperature, the reaction mixture was extracted with $CH_2Cl_2$. The organic layers were combined, washed with brine, dried over $Na_2SO_4$ and the solvent was evaporated under reduced pressure. Silica gel column chromatography (Biotage Sfär Duo 5 g, 0–20% EtOAc/cyclohexane, linear gradient) afforded the title compound as a grey solid (49.1 mg, 82%). $^1H$ NMR (400 MHz, $CDCl_3$) δ 8.07 (d, *J* = 7.5 Hz, 1H), 7.75 – 7.63 (m, 2H), 7.35 (d, *J* = 1.7 Hz, 2H), 7.28 (d, *J* = 7.6 Hz, 1H), 7.10 (dd, *J* = 8.2, 1.7 Hz, 2H), 6.85 (d, *J* = 8.3 Hz, 2H), 6.75 (t, *J* = 2.3 Hz, 2H), 6.29 – 6.33 (m, 2H), 6.22 (t, *J* = 3.1 Hz, 2H), 3.73 (s, 6H). $^{13}C$ NMR (101 MHz, ($CDCl_3$) δ 169.5 (CO), 153.2 (C), 151.4 (C), 136.1 (C), 135.3 (CH), 133.2 (C), 130.1 (CH), 128.2 (CH), 126.7 (C), 125.4 (CH), 125.0 (CH), 124.2 (CH), 124.1 (CH), 117.0 (C), 116.4 (CH), 109.9 (CH), 108.3 (CH), 82.7 (C), 35.5 ($CH_3$). Analytical HPLC: $t_R$ = 4.7 min, 98% purity (5–95% $CH_3CN/H_2O$, gradient with constant 0.1% formic acid additive, 8 min run, 0.5 mL/min flow, UV detection at 254 nm. HRMS (ESI) calcd for $C_{30}H_{23}N_2O_3$ $[M+H]^+$ 459.1703, found 459.1701.

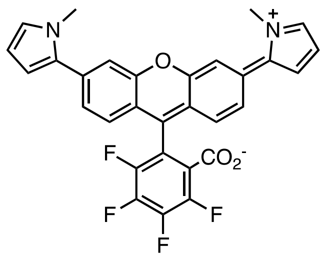

**(11):** $F_4$-Fluorescein ditriflate (32.4 mg, 0.046 mmol, 1 eq), 1-methyl-2-pyrroleboronic acid pinacol ester (40.6 mg, 0.139 mmol, 3.0 eq), $Pd(PPh_3)_4$ (3.2 mg, 0.003 mmol, 0.06 eq) and KOAc (27.2 mg, 0.277 mmol, 6.0 eq) were loaded into a vial. The vial was sealed and evacuated/backfilled with argon. A mixture of 1,4-dioxane/$H_2O$ (2/1, 1.05 mL) was added and the reaction mixture was stirred at 100°C for 22 h. After cooling to room temperature, the reaction mixture was extracted with $CH_2Cl_2$. The organic layers were combined, washed with brine, dried over $Na_2SO_4$ and the solvent was evaporated under reduced pressure. Silica gel column chromatography (15% EtOAc/cyclohexane) afforded the title compound as an off-white solid (21.4 mg, 88%). $^1H$ NMR (400 MHz, $(CD_3)_2SO$, 5 mM DMF) δ 7.51 (t, *J* = 1.1 Hz, 2H), 7.31 (d, *J* = 1.1 Hz, 4H), 6.93 (dd, *J* = 2.7, 1.8 Hz, 2H), 6.38 (dd, *J* = 3.7, 1.8 Hz, 2H), 6.11 (dd, *J* = 3.7, 2.6 Hz, 2H), 3.75 (s, 6H). $^{19}F$ NMR ($CDCl_3$, 376 MHz) δ -137.72 – -138.11 (m, 1F), -141.36 – -141.74 (m, 2F), -149.48 – -149.81 (m, 1F). Analytical HPLC: $t_R$ = 5.4 min, 98% purity, 5–95% $CH_3CN/H_2O$, gradient with constant 0.1% formic acid additive, 10 min run, 0.5 mL/min flow, UV detection at 254 nm. HRMS (ESI) calcd for $C_{30}H_{19}F_4N_2O_3$ $[M+H]^+$ 531.1326, found 531.1331.

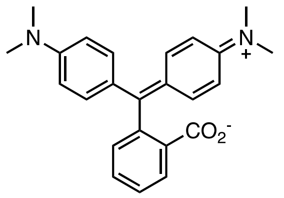

**(12):** *N,N*-dimethylaniline (1.03 mL, 8.10 mmol, 2 eq), phthalic anhydride (600 mg, 4.05 mmol, 1 eq) and $ZnCl_2$ (1.14 mg, 8.10 mmol, 2 eq) were loaded into a sealed vial and stirred vigorously at 150°C for 5 h. The reaction mixture was cooled to room temperature then dissolved progressively in organic solvents ($CH_3CN$ and EtOAc). Aqueous HCl (1 M, 10 mL) was added and the organic solvents evaporated under reduced pressure. The aqueous phase was neutralised to pH = 5-6 with NaOH (1M), which formed a green precipitate. The precipitate was dissolved with EtOAc. The layers were separated and the aqueous layer was extracted with EtOAc (3x). The organic layers were combined, dried over $Na_2SO_4$, filtered and evaporated to dryness. Trituration with cyclohexane to remove excess *N,N*-dimethylaniline followed by purification by silica gel column chromatography (Biotage Sfär Duo 25 g, 0–30% EtOAc/cyclohexane, linear gradient) afforded the title compound as a yellow solid (271 mg, 18%). For characterization by NMR and UV/Vis spectroscopy, purification of an analytical fraction was performed by reverse phase HPLC (10–50% $CH_3CN/H_2O$, linear gradient with constant 0.1% v/v TFA additive). The pooled HPLC product fractions were combined, and lyophilized to afford a fraction of the pure compound as a TFA salt. $^1H$ NMR (400 MHz, $CDCl_3$) δ 7.94 (d, *J* = 7.6 Hz, 1H), 7.75 (t, *J* = 7.6, 1H), 7.61 (t, *J* = 7.6 Hz, 1H), 7.55 (d, *J* = 7.6 Hz, 1H), 7.41 – 7.35 (m, 8H), 3.13 (s, 12H). $^{13}C$ NMR (101 MHz, $CDCl_3$) δ 169.3 (C), 150.8 (C), 145.1 (C), 139.1 (C), 134.9 (CH), 130.2 (CH), 129.1 (CH), 126.5 (CH), 125.3 (C), 124.1 (CH), 119.0 (CH), 90.5 (C), 45.0 ($CH_3$). Analytical HPLC: $t_R$ = 4.2 min, >99% purity, 5–95% $CH_3CN/H_2O$, gradient with constant 0.1% formic acid additive, 8 min run, 0.5 mL/min flow, UV detection at 254 nm. HRMS (ESI) calcd for $C_{24}H_{21}N_2O_2$ $[M+H]^+$ 373.1911, found 373.1907.

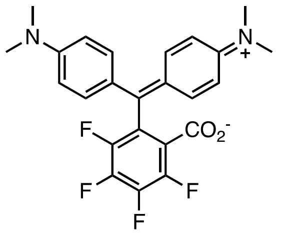

**(13):** *N,N*-dimethylaniline (2.00 g, 16.53 mmol, 2 eq), tetrafluorophthalic anhydride (1.82 g, 8.27 mmol, 1 eq) and $ZnCl_2$ (2.25 g, 8.27 mmol, 2 eq) were loaded into a sealed vial and stirred vigorously at 150°C for 4 h. The reaction mixture was cooled to room temperature then dissolved progressively in organic solvents ($CH_3CN$ and EtOAc). Aqueous HCl (1 M, 50 mL) was added and the organic solvents evaporated under reduced pressure. The aqueous layer was neutralised to pH 5-6 with NaOH (1M), which formed a green precipitate. The precipitate was dissolved with EtOAc. The layers were separated and the aqueous layer was extracted with EtOAc (3x). The organic layers were combined, dried over $Na_2SO_4$, filtered and evaporated to dryness. Purification by silica gel column chromatography (Biotage Sfär Duo 50 g, 0-30% EtOAc/cyclohexane) afforded the title compound as a blue-green solid (746 mg, 20%). $^1H$ NMR (400 MHz, $CDCl_3$) δ 7.15 (d, *J* = 8.6 Hz, 4H), 6.65 (d, *J* = 8.6 Hz, 4H), 2.96 (s, 12H). $^{19}F$ NMR (376 MHz, $CDCl_3$) δ -137.87 (t, *J* = 20.2 Hz, 1F), -138.82 – -139.06 (m, 1F), -143.33 – -143.59 (m, 1F), -151.88 – -152.18 (m, 1F). Analytical HPLC: $t_R$ = 4.7 min, 98% purity, 5–95% $CH_3CN/H_2O$, gradient with constant 0.1% formic acid additive, 8 min run, 0.5 mL/min flow, UV detection at 254 nm. HRMS (ESI) calcd for $C_{24}H_{21}F_4N_2O_2$ $[M+H]^+$ 445.1534, found 445.1530.

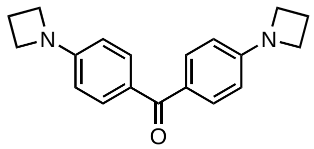

**(S1):** 4,4'-Dihydroxybenzophenone ditriflate (1.00 g, 2.10 mmol, 1 eq), $Pd_2(dba)_3$ (193 mg, 0.210 mmol, 0.1 eq), XPhos (301 mg, 0.631 mmol, 0.3 eq) and $Cs_2CO_3$ (1.92 g, 5.89 mmol, 2.8 eq) were loaded into a vial. The vial was sealed and evacuated/backfilled with argon. 1,4-Dioxane (8 mL) and subsequently azetidine (340 µL, 5.05 mmol, 2.4 eq) were added then the reaction mixture was stirred at 100°C for 5 h. After cooling to room temperature, the mixture was filtered through Celite, washed with EtOAc, and the solvent evaporated under reduced pressure. Purification by silica gel column chromatography (Biotage Sfär Duo 10 g, 0–30% EtOAc/cyclohexane, linear gradient) followed by recrystallisation from MeOH afforded the title compound as a yellow solid (333 mg, 54%). $^{1}$H NMR (400 MHz, $(CD_3)_2CO$) δ 7.62 (d, *J* = 8.6 Hz, 4H), 6.43 (d, *J* = 8.6 Hz, 4H), 3.98 (t, *J* = 7.3 Hz, 8H), 2.42 (p, *J* = 7.3 Hz, 4H). $^{13}$C NMR (101 MHz, $(CD_3)_2CO$) δ 193.4 (C), 155.0 (C), 132.3 (CH), 127.8 (C), 110.3 (CH), 52.4 ($CH_2$), 17.2 ($CH_2$). Analytical HPLC: $t_R$ = 4.1 min, 97% purity, 5–95% $CH_3CN/H_2O$, gradient with constant 0.1% formic acid additive, 8 min run, 0.5 mL/min flow, UV detection at 254 nm. HRMS (ESI) calcd for $C_{19}H_{21}N_2O$ $[M+H]^+$ 293.1648, found 293.1643.

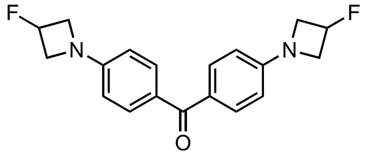

**(S2):** 4,4'-Dihydroxybenzophenone ditriflate (1.50 g, 3.13 mmol, 1 eq), $Pd_2(dba)_3$ (287 mg, 0.313 mmol, 0.1 eq), XPhos (448 mg, 0.939 mmol, 0.3 eq), $Cs_2CO_3$ (5.30 g, 16.28 mmol, 5.2 eq) and 3-fluoroazetidine hydrochloride (838 mg, 7.52 mmol, 2.4 eq) were loaded into a vial. The vial was sealed and evacuated/backfilled with argon. 1,4-Dioxane (10 mL) was added and the reaction mixture was stirred at 100°C for 4 h. After cooling to room temperature, the reaction mixture was filtered through Celite, washed with EtOAc and the solvent evaporated under reduced pressure. Purification by silica gel column chromatography (Biotage Sfär Duo 25 g, 0–30% EtOAc/cyclohexane, linear gradient) followed by recrystallisation from MeOH afforded the title compound as a pale-yellow solid (833 mg, 81%). $^{1}$H NMR (400 MHz, $(CD_3)_2CO$) δ 7.65 (d, *J* = 8.7 Hz, 4H), 6.54 (d, *J* = 8.7 Hz, 4H), 5.56 (dtt, $^{2}J_{HF}$ = 57.3 Hz, *J* = 8.8, 3.3 Hz, 2H), 4.39 – 4.29 (m, 4H), 4.11 – 4.00 (m, 4H). $^{13}$C NMR (101 MHz, $(CD_3)_2CO$) δ 193.5 (C), 154.3 (d, $^{4}J_{CF}$ = 1.5 Hz, C), 132.4 (CH), 128.6 (C), 119.8 (C), 116.5 (C), 111.2 (CH), 84.1 (d, $^{1}J_{CF}$ = 203.0 Hz, CFH), 59.9 (d, $^{2}J_{CF}$ = 24.2 Hz, $CH_2$). $^{19}$F NMR (376 MHz, $(CD_3)_2CO$) δ -179.83 – -179.86 (m, 2F). Analytical HPLC: $t_R$ = 4.3 min, 95% purity, 5–95% $CH_3CN/H_2O$, gradient with constant 0.1% formic acid additive, 8 min run, 0.5 mL/min flow, UV detection at 254 nm. HRMS (ESI) calcd for $C_{19}H_{19}N_2O_2$ $[M+H]^+$ 329.1460, found 329.1454.

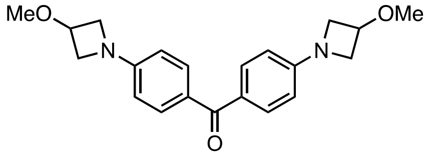

**(S3):** 4,4'-Dihydroxybenzophenone ditriflate (1.01 g, 2.12 mmol, 1 eq), $Pd_2(dba)_3$ (0.212 g, 0.212 mmol, 0.1 eq), XPhos (0.303 g, 0.636 mmol, 0.3 eq), $Cs_2CO_3$ (3.59 g, 11.02 mmol, 5.2 eq) and 3-methoxyazetidine hydrochloride (0.628 g, 5.085 mmol, 2.4 eq) were loaded into a vial. The vial was sealed and evacuated/backfilled with argon. 1,4-Dioxane (8 mL) was added then the reaction mixture was stirred at 100°C for 4 h. After cooling to room temperature, the reaction mixture was filtered through Celite, washed with EtOAc, and the solvent evaporated under reduced pressure. Purification by silica gel column chromatography (Biotage Sfär Duo 25 g, 0–50% EtOAc/cyclohexane, linear gradient) afforded the title compound as a yellow solid (679 mg, 91%).$^{1}$H NMR (400 MHz, $CDCl_3$) δ 7.71 (d, *J* = 8.5 Hz, 4H), 6.43 (d, *J* = 8.5 Hz, 4H), 4.40 – 4.34 (m, 2H), 4.09 – 4.03 (m, 4H), 3.85 – 3.80 (m, 4H), 3.35 (s, 6H). $^{13}$C NMR (101 MHz, $CDCl_3$) δ 194.3 (CO), 153.5 (C), 132.2 (CH), 127.7 (C), 110.2 (CH), 70.0 (CH), 58.6 ($CH_2$), 56.3 ($CH_3$). Analytical HPLC: $t_R$ = 3.8 min, 97% purity, 5–95% $CH_3CN/H_2O$, gradient with constant 0.1% formic acid additive, 8 min run, 0.5 mL/min flow, UV detection at 254 nm. HRMS (ESI) calcd for $C_{21}H_{25}N_2O_3$ $[M+H]^+$ 353.1860, found 353.1853.

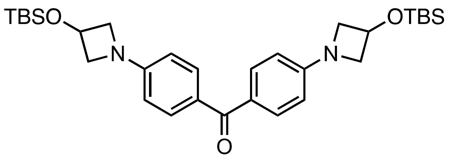

**(S5):** 4,4'-Dihydroxybenzophenone ditriflate (1.09 g, 2.29 mmol, 1 eq), $Pd_2(dba)_3$ (209 mg, 0.229 mmol, 0.1 eq), XPhos (327 mg, 0.686 mmol, 0.3 eq), $Cs_2CO_3$ (4.32 g, 13.3 mmol, 6 eq) and 3-hydroxyazetidine hydrochloride (642 mg, 6.86 mmol, 3 eq) were loaded into a vial. The vial was sealed and evacuated/backfilled with argon. 1,4-Dioxane (10 mL) was added then the reaction mixture was stirred at 100°C for 4 h. After cooling to room temperature, the reaction mixture was filtered through Celite, washed with EtOAc and MeOH, and the solvents evaporated under reduced pressure. Partial purification by silica gel column chromatography (Biotage Sfär Duo 25 g, 0–50% EtOAc/cyclohexane, linear gradient) afforded the hydroxy intermediate **S4** as an orange-red solid. The intermediate was dissolved in $CH_2Cl_2$ (80 mL), cooled to 0°C, and imidazole (1.47 g, 21.6 mmol, 6 eq) followed by *tert*-butyldimethylsilyl chloride (1.63 g, 10.8 mmol, 3 eq) were added. The reaction mixture was stirred at room temperature under argon for 14 h, after which it was washed with $H_2O$ and brine. The organic layer was dried over anhydrous $Na_2SO_4$, filtered, and evaporated to dryness. Purification by silica gel chromatography (Biotage Sfär Duo 25 g, 0–15% EtOAc/cyclohexane, linear gradient) followed by recrystallisation from MeOH afforded the title compound as a pale-yellow solid (268 mg, 21% over two steps). $^1H$ NMR (400 MHz, $CDCl_3$) δ 7.70 (d, *J* = 8.5 Hz, 4H), 6.43 (d, *J* = 8.5 Hz, 4H), 4.82 – 4.74 (m, 2H), 4.27 – 4.18 (m, 4H), 3.79 – 3.74 (m, 4H), 0.90 (s, 18H), 0.09 (s, 12H). $^{13}C$ NMR (101 MHz, $CDCl_3$) δ 194.3 (CO), 153.6 (C), 132.2 (CH), 127.7 (C), 110.5 (CH), 62.4 ($CH_2$), 61.9 ($CH_2$) 25.9 ($CH_3$), 18.1 (C), -4.8 ($CH_3$). HRMS (ESI) calcd for $C_{31}H_{49}N_2O_3Si_2$ $[M+H]^+$ 553.3276, found 553.3267.

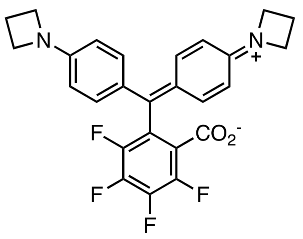

**(14):** A solution of 2,3,4,5-tetrafluorobenzoic acid (2.01 g, 10.3 mmol, 10 eq) in anhydrous THF (20 mL) was cooled to -78°C under argon. *n*-Butyllithium (2.5 M in hexanes, 8.27 mL, 20.6 mmol, 20 eq) was added, and the reaction was stirred at -78°C for 3 h. **S1** (302 mg, 1.03 mmol, 1 eq) in anhydrous THF (20 mL) was added, the reaction was allowed to warm to room temperature and stirred for 16 h. The reaction mixture was subsequently diluted with saturated $NH_4Cl$ and $H_2O$ and extracted with EtOAc (×3). The combined organic layers were washed with saturated $NaHCO_3$ and brine, dried over anhydrous $Na_2SO_4$, filtered, and evaporated to dryness. Purification by silica gel chromatography (Biotage Sfär Duo 25 g, 0–30% TBME/cyclohexane, linear gradient) afforded the title compound as a blue-green solid (360 mg, 74%). $^1H$ NMR (400 MHz, $(CD_3)_2SO$, 5 mM DMF) δ 7.03 (d, *J* = 8.4 Hz, 4H), 6.42 – 6.37 (d, *J* = 8.4 Hz, 4H), 3.81 (t, *J* = 7.2 Hz, 8H), 2.30 (p, *J* = 7.2 Hz, 4H). $^{19}F$ NMR (376 MHz, $(CD_3)_2CO$) δ -138.94 (t, *J* = 20.0 Hz, 1F), -140.60 (td, *J* = 19.3, 8.8 Hz, 1F), -144.83 (ddd, *J* = 20.9, 18.0, 8.8 Hz, 1F), -153.52 (ddd, *J* = 20.4, 17.9, 4.5 Hz, 1F). Analytical HPLC: $t_R$ = 4.7 min, >99% purity, 5–95% $CH_3CN/H_2O$, gradient with constant 0.1% formic acid additive, 8 min run, 0.5 mL/min flow, UV detection at 254 nm. HRMS (ESI) calcd for $C_{26}H_{21}F_4N_2O_2$ $[M+H]^+$ 469.1534, found 469.1525.

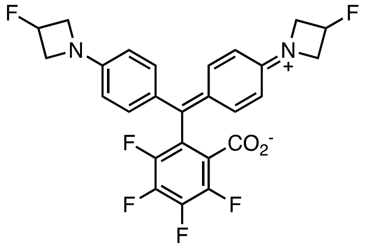

**(15):** A solution of 2,3,4,5-tetrafluorobenzoic acid (1.21 g, 6.24 mmol, 10 eq) in anhydrous THF (15 mL) was cooled to -78°C under argon. *N*-Butyllithium (2.5 M in hexanes, 5.00 mL, 12.5 mmol, 20 eq) was added, and the reaction was stirred at -78°C for 3 h. A solution of **S2** (205 mg, 0.624 mmol, 1 eq) in anhydrous THF (15 mL) was added, and the reaction was allowed to warm to room temperature and stirred for 18 h. The reaction mixture was subsequently diluted with saturated $NH_4Cl$ and water and extracted with EtOAc (×3). The combined organic layers were washed with saturated $NaHCO_3$ and brine, dried over anhydrous $Na_2SO_4$, filtered, and evaporated to dryness. Purification by silica gel chromatography (Biotage Sfär Duo 25 g, 0–30% TBME/cyclohexane, linear gradient) afforded the title compound as a blue-green solid (232 mg, 74%). $^1$H NMR (400 MHz, $(CD_3)_2SO$, 5 mM DMF) δ 7.07 (d, $J$ = 8.5 Hz, 4H), 6.48 (d, $J$ = 8.5 Hz, 4H), 5.48 (dtt, $^2J_{HF}$ = 57.5 Hz, $J$ = 8.8, 3.0 Hz, 2H), 4.23 – 4.09 (m, 4H), 3.95 – 3.85 (m, 4H). $^{19}$F NMR (376 MHz, $(CD_3)_2CO$) δ -138.83 – -139.07 (m, 1F), -139.52 – -139.76 (m, 1F), -142.56 – -151.87 (m, 1F), -151.51 – -151.81 (m, 1F), -178.49 – -178.93 (m, 2F). Analytical HPLC: $t_R$ = 4.4 min, 97% purity, 5–95% $CH_3CN/H_2O$, gradient with constant 0.1% formic acid additive, 8 min run, 0.5 mL/min flow, UV detection at 254 nm. HRMS (ESI) calcd for $C_{26}H_{19}F_4N_2O_2$ $[M+H]^+$ 505.1345, found 505.1338.

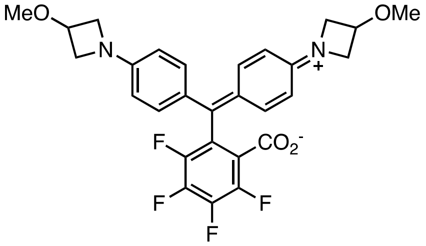

**(16):** A solution of 2,3,4,5-tetrafluorobenzoic acid (3.45 g, 17.8 mmol, 10 eq) in anhydrous THF (42 mL) was cooled to -78°C under argon. *n*-Butyllithium (2.5 M in hexanes, 14.2 mL, 35.5 mmol, 20 eq) was added, and the reaction was stirred at -78°C for 3 h. A solution of **S3** (636 mg, 1.78 mmol, 1 eq) in anhydrous THF (35 mL) was added, the reaction was allowed to warm to room temperature and stirred for 18 h. The reaction mixture was subsequently diluted with saturated $NH_4Cl$ and water and extracted with EtOAc (3x). The combined organic layers were washed with saturated $NaHCO_3$ and brine, dried over anhydrous $Na_2SO_4$, filtered, and evaporated to dryness. Purification by silica gel chromatography (Biotage Sfär Duo 25 g, 0–30% EtOAc/cyclohexane, linear gradient) afforded the title compound as a blue-green solid (656 mg, 77%). For full characterization by NMR and UV/Vis spectroscopy an analytical fraction of the product was further purified by reverse phase HPLC (10–100% $CH_3CN/H_2O$, linear gradient with constant 0.1% v/v TFA additive). The pooled HPLC product fractions were neutralised with saturated $NaHCO_3$ and extracted with EtOAc (3x). The combined organic layers were dried over $Na_2SO_4$, filtered and evaporated to dryness to afford a pure sample. $^1$H NMR (400 MHz, $CD_3CN$) δ 7.12 – 7.04 (m, 4H), 6.46 – 6.38 (m, 4H), 4.30 (tt, $J$ = 6.1, 4.2 Hz, 2H), 4.07 – 4.02 (m), 3.68 – 3.63 (m, 4H), 3.27 (s, 6H). $^{19}$F NMR (376 MHz, $(CD_3)_2SO$) δ -138.89 – -139.20 (m, 1F), -139.83 – -140.08 (m, 1F), -142.91 – -143.44 (m, 1F), -151.96 – -152.41 (m, 1F). Analytical HPLC: $t_R$ = 4.5 min, >99 % purity, 5–95% $CH_3CN/H_2O$, gradient with constant 0.1% formic acid additive, 8 min run, 0.5 mL/min flow, UV detection at 254 nm. HRMS (ESI) calcd for $C_{28}H_{25}F_4N_2O_4$ $[M+H]^+$ 529.1745, found 529.1735.

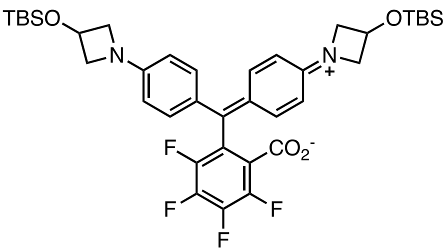

**(S6):** A solution of 2,3,4,5-tetrafluorobenzoic acid (941 mg, 4.85 mmol, 10 eq) in anhydrous THF (10 mL) was cooled to -78°C under argon. *n*-Butyllithium (2.5 M in hexanes, 3.88 mL, 9.70 mmol, 20 eq) was added, and the reaction was stirred at -78°C for 3 h. A solution of **S4** (268 mg, 0.485 mmol, 1 eq) in anhydrous THF (20 mL) was added, the reaction was allowed to warm to room temperature and stirred for 17 h. The reaction mixture was subsequently diluted with saturated $NH_4Cl$ and water and extracted with EtOAc (3x). The combined organic layers were washed with saturated $NaHCO_3$ and brine, dried over anhydrous $Na_2SO_4$, filtered, and evaporated to dryness. Purification by silica gel chromatography (Biotage Sfär Duo 25 g, 0–15% EtOAc/cyclohexane, linear gradient) afforded the title compound as a blue-green solid (155 mg, 44%). For full characterization by NMR and UV/Vis spectroscopy an analytical fraction of the product was further purified by column chromatography (Biotage Sfär Duo 5 g, 0–15% EtOAc/cyclohexane, linear gradient) to afford a pure sample. $^1$H NMR (400 MHz, $(CD_3)_2SO$, 5 mM DMF) δ 7.04 (d, *J* = 8.4 Hz, 4H), 6.43 (d, *J* = 8.7 Hz, 4H), 4.77 (p, *J* = 5.5 Hz, 2H), 4.13 (t, *J* = 7.1 Hz, 4H), 3.52 (dd, *J* = 8.0, 4.7 Hz, 4H), 0.86 (s, 18H), 0.06 (s, 12H). $^{19}$F NMR (376 MHz, $(CD_3)_2SO$) δ -138.85 – -139.05 (m, 1F), -139.69 – -139.87 (m, 1F), -142.79 – -143.00 (m, 1F), -151.81 – -151.97 (m, 1F). Analytical HPLC: $t_R$ = 7.7 min, 97% purity, 5–95% $CH_3CN/H_2O$, gradient with constant 0.1% formic acid additive, 8 min run, 0.5 mL/ min flow, UV detection at 254 nm. HRMS (ESI) calcd for $C_{38}H_{49}F_4N_2O_4Si_2$ $[M+H]^+$ 729.3162, found 729.3139.

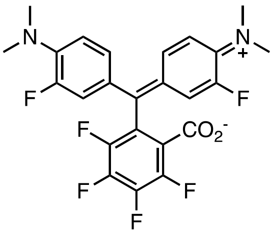

**(17):** 2-fluorodimethylaniline (510 mg, 3.67 mmol, 2 eq), tetrafluorophthalic anhydride (400 mg, 1.84 mmol, 1 eq) and $ZnCl_2$ (500 mg, 3.67 mmol, 2 eq) were loaded into a sealed vial then stirred vigorously at 150°C for 4 h. The reaction mixture was cooled to room temperature then dissolved progressively in organic solvents ($CH_3CN$ and EtOAc). Aqueous HCl (1 M, 12 mL) was added and the organic solvents were evaporated under reduced pressure. The aqueous phase was neutralised to pH 5-6 with NaOH (1M), which formed a green precipitate. The precipitate was dissolved with EtOAc. The layers were separated and the aqueous layer was extracted with EtOAc (3x). The organic layers were combined, dried over $Na_2SO_4$, filtered and evaporated to dryness. Trituration with cyclohexane removed some excess 2-fluorodimethylaniline, and purification was performed by silica gel column chromatography (Biotage Sfär Duo 25 g, 0–15% EtOAc/cyclohexane, linear gradient), followed by reverse phase HPLC (10–50% $CH_3CN/H_2O$, linear gradient with constant 0.1% v/v TFA additive). The pooled HPLC product fractions were combined, and lyophilized to afford the title compound as a white solid (5.4 mg, 1%, TFA salt). $^1$H NMR (400 MHz, $CDCl_3$) δ 7.01 – 6.89 (m, 4H), 6.82 (t, *J* = 8.9 Hz, 2H), 2.89 (d, *J* = 1.1 Hz, 12H). $^{19}$F NMR (376 MHz, $CDCl_3$) δ -118.96 (m, 2F), -136.03 – -136.23 (m, 1F), -136.55 – -136.71 (m, 1F), -140.30 – -140.49 (m, 1F), -147.32 (td, *J* = 39.0, 4.9 Hz, 1F). Analytical HPLC: $t_R$ = 4.8 min, 97% purity, 5–95% $CH_3CN/H_2O$, gradient with constant 0.1% formic acid additive, 8 min run, 0.5 mL/min flow, UV detection at 254 nm. HRMS (ESI) calcd for $C_{24}H_{22}F_3N_2O_2$ $[M+H]^+$ 481.1345, found 481.1340.

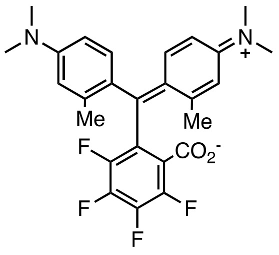

**(18):** *N,N*-dimethyl-*m*-toluidine (1.33 mL, 9.13 mmol, 2 eq), tetrafluorophthalic anhydride (1.00 g, 4.57 mmol, 1 eq) and $ZnCl_2$ (1.25 g, 9.13 mmol, 2 eq) were loaded into a sealed vial then stirred vigorously at 150°C for 3.5 h. The reaction mixture was cooled to room temperature then dissolved progressively in organic solvents ($CH_3CN$ and EtOAc). Aqueous HCl (1 M, 20 mL) was added and the organic solvents evaporated under reduced pressure. The aqueous phase was neutralised to pH 5-6 with NaOH (1M), which formed a green precipitate. The precipitate was dissolved with EtOAc, the layers were separated and the aqueous layer was extracted with EtOAc (3x). The organic layers were combined, dried over $Na_2SO_4$, filtered and evaporated to dryness. Trituration with cyclohexane removed residual *N,N*-dimethylaniline and purification by silica gel column chromatography (Biotage Sfär Duo 25 g, 0-30% EtOAc/cyclohexane, linear gradient) afforded the title compound as a blue-green solid (476 mg, 22%). For full characterization by NMR and UV/Vis spectroscopy, purification of an analytical fraction was performed by reverse phase HPLC (10–95% $CH_3CN/H_2O$, linear gradient with constant 0.1% v/v TFA additive). The pooled HPLC product fractions were combined, and lyophilized to afford the title compound as a pale green solid (73.3 mg, 3%, TFA salt). $^1H$ NMR (400 MHz, $CDCl_3$) δ 6.80 (dd, *J* = 8.8, 2.8 Hz, 2H), 6.53 (s, 2H), 6.42 (dd, *J* = 8.8, 2.8 Hz, 2H), 2.95 (d, *J* = 1.2 Hz, 12H), 2.11 (s, 6H). $^{19}F$ NMR (376 MHz, $CDCl_3$) δ -137.22 – -137.95 (td, *J* = 20.0, 8.4 Hz, 1F), -139.28 – -139.68 (td, *J* = 20.0, 8.4 Hz, 1F), -143.49 – -143.96 (m, 1F), -151.44 (td, *J* = 19.5, 4.3 Hz, 1F). Analytical HPLC: $t_R$ = 4.8 min, 98% purity, 5–95% $CH_3CN/H_2O$, gradient with constant 0.1% formic acid additive, 8 min run, 0.5 mL/min flow, UV detection at 254 nm. HRMS (ESI) calcd for $C_{26}H_{25}F_4N_2O_2$ $[M+H]^+$ 473.1847, found 473.1841.

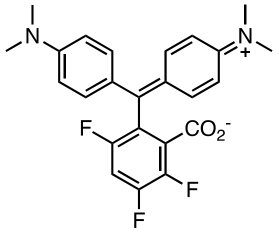

**(19):** Compound **19** was isolated as a by-product when attempting to synthesize the HaloTag ligand. **6-(MOM-MAC)-FMGL**[5] (223 mg, 0.431 mmol, 1 eq), and camphorsulfonic acid (110 mg, 0.475 mmol, 1.1 eq) were loaded into a vial which was sealed and evacuated/backfilled with argon. AcOH/DME (1/1, 6.4 mL) was added and the reaction stirred at 80°C for 68 h. The solvent was evaporated under reduced pressure, and the aqueous phase was neutralised with saturated $NaHCO_3$ before the reaction mixture was extracted with EtOAc (3x). The organic layers were combined, dried over $Na_2SO_4$, filtered and evaporated to dryness. To the crude intermediate, a solution of HaloTag(O2) amine (as the TFA salt, 229 mg, 0.678 mmol, 3 eq), DIEA (394 µL, 2.26 mmol, 10 eq) and HATU (258 mg, 0.678 mmol, 3 eq) in DMF (3.5 mL) were added. The reaction mixture was stirred for 18 h at room temperature and then evaporated to dryness. Purification was performed by silica gel column chromatography (Biotage Sfär Duo 25 g, 0–20% EtOAc/cyclohexane, linear gradient) followed by reverse phase HPLC (20–80% $CH_3CN/H_2O$, linear gradient with constant 0.1% v/v TFA additive). The pooled HPLC product fractions were neutralised with saturated $NaHCO_3$, extracted with EtOAc (3x), dried over $Na_2SO_4$, filtered and evaporated to dryness to afford the title compound as a blue-green solid (20.6 mg, 11%). $^1H$ NMR (400 MHz, $CDCl_3$) δ 7.36 – 7.31 (m, 4H), 7.29 (d, *J* = 5.2 Hz, 1H), 7.20 – 7.15 (m, 4H), 3.11 (s, 12H). $^{19}F$ NMR (376 MHz, $CDCl_3$) δ -113.82 (d, *J* = 19.5 Hz, 1F), -129.4 – -129.3 (m, 1F), -142.13 (t, *J* = 20.9 Hz, 1F). Analytical HPLC: $t_R$ = 4.5 min, 96% purity, 5–95% $CH_3CN/H_2O$, gradient with constant 0.1% formic acid additive, 8 min run, 0.5 mL/min flow, UV detection at 254 nm. HRMS (ESI) calcd for $C_{24}H_{22}F_3N_2O_2$ $[M+H]^+$ 427.1628, found 427.1624.

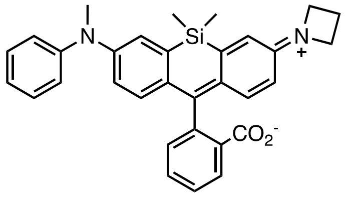

**(20):** Si-fluorescein ditriflate (100 mg, 0.157 mmol, 1 eq), $Pd_2(dba)_3$ (14.4 mg, 0.016 mmol, 0.1 eq), XantPhos (27.3 mg, 0.048 mmol, 0.3 eq) and $Cs_2CO_3$ (102.4 mg, 0.314 mmol, 2.0 eq) were loaded into a vial. The vial was sealed and evacuated/backfilled with argon. 1,4-Dioxane (2 mL) and subsequently *N*-methylaniline (17.0 µL, 0.157 mmol, 1 eq) were added then the reaction mixture was stirred at 80°C for 2 h. After cooling to room temperature, the reaction mixture was filtered through Celite, washed with EtOAc and the solvent was evaporated under reduced pressure. Partial purification was performed by silica gel column chromatography (cyclohexane:$CH_2Cl_2$:toluene (2:2:1)) and afforded the intermediate as a yellow-green solid. The intermediate showed moderate stability and was directly engaged in the next step.
The intermediate (~40 mg, 0.067 mmol), $Pd_2dba_3$ (6.1 mg, 0.007 mmol, 0.1 eq), XPhos (9.5 mg, 0.020 mmol, 0.3 eq) and $Cs_2CO_3$ (60.7 mg, 0.186 mmol, 2.8 eq) were loaded into a vial. The vial was sealed and evacuated/backfilled with argon. 1,4-Dioxane (1 mL) and subsequently azetidine (11 µL, 0.186 mmol, 2.4 eq) were added then the reaction mixture was stirred at 100°C for 3 h. After cooling to room temperature, the reaction mixture was filtered through Celite, washed with EtOAc and the solvent was evaporated under reduced pressure. The crude material was purified by silica gel column chromatography (10–15% TBME/cyclohexane, linear gradient) followed by reverse phase HPLC (10–50% $CH_3CN/H_2O$, linear gradient with constant 0.1% v/v TFA additive). The pooled HPLC product fractions were combined and lyophilized to afford the title compound as a blue-green solid (10.7 mg, 11% yield over 2 steps, TFA salt). $^1H$ NMR ($CDCl_3$, 400 MHz) δ 7.99 (d, *J* = 7.6 Hz, 1H), 7.69 (td, *J* = 7.5, 1.1 Hz, 1H), 7.58 (td, *J* = 7.5, 1.1 Hz, 1H), 7.36 – 7.28 (m, 3H), 7.19 (dd, *J* = 14.7, 2.7 Hz, 2H), 7.14 – 7.03 (m, 3H), 6.97 (d, *J* = 8.8 Hz, 1H), 6.82 (d, *J* = 8.8 Hz, 1H), 6.72 – 6.77 (m, 2H), 4.24 (t, *J* = 7.7 Hz, 4H), 3.33 (s, 3H), 2.54 (p, *J* = 7.7 Hz, 2H), 0.592 (s, 3H), 0.587 (s, 3H). $^{13}C$ NMR (101 MHz, $CDCl_3$) δ 170.2 (C), 152.8 (C), 148.6 (C), 148.1 (C), 146.3 (C), 139.3 (C), 139.2 (C), 137.1 (C), 134.4 (C), 134.0 (CH), 129.6 (CH), 129.4 (CH), 129.0 (CH), 128.7 (CH), 127.0 (C), 126.5 (CH), 125.0 (CH), 123.6 (CH), 123.2 (CH), 122.3 (CH), 120.7 (CH), 118.8 (CH), 116.6 (CH), 77.4 (C), 54.8 ($CH_2$), 40.2 ($CH_3$), 16.8 ($CH_2$), 0.3 (Si-$CH_3$), -1.7 (Si-$CH_3$). Analytical HPLC: $t_R$ = 5.2 min, 95% purity, 5–95% $CH_3CN/H_2O$, gradient with constant 0.1% formic acid additive, 8 min run, 0.5 mL/min flow, UV detection at 254 nm. HRMS (ESI) calcd for $C_{32}H_{31}N_2O_2Si$ $[M+H]^+$ 503.2149, found 503.2145.

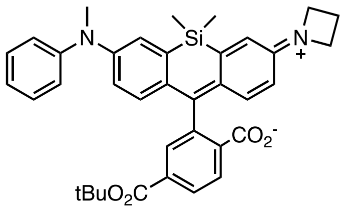

**(S7):** 6-$CO_2$tBu-fluorescein ditriflate (202 mg, 0.273 mmol, 1 eq), $Pd_2(dba)_3$ (25.0 mg, 0.027 mmol, 0.1 eq), XantPhos (47.5 mg, 0.082 mmol, 0.3 eq) and $Cs_2CO_3$ (178 mg, 0.547 mmol, 2.0 eq) were loaded into a vial. The vial was sealed and evacuated/backfilled with argon. 1,4-Dioxane (4 mL) and subsequently *N*-methylaniline (29.6 µL, 0.273 mmol, 1 eq) were added then the reaction mixture was stirred at 80°C for 2 h. After cooling to room temperature, the reaction mixture was filtered through Celite, washed with EtOAc and the solvent was evaporated under reduced pressure. Purification by silica gel column chromatography (Biotage Sfar Duo 10 g, cyclohexane:$CH_2Cl_2$: toluene (10:0:0–5:3.5:1.5), linear gradient) afforded the intermediate as a pale green solid. The intermediate showed moderate stability and was directly engaged in the next step.
This intermediate (~63 mg, 0.091 mmol), $Pd_2(dba)_3$ (8.3 mg, 0.009 mmol, 0.1 eq), XPhos (12.9 mg, 0.027 mmol, 0.3 eq) and $Cs_2CO_3$ (82.6 mg, 0.254 mmol, 2.8 eq) were loaded into a vial. The vial was sealed and evacuated/backfilled with argon. 1,4-Dioxane (2 mL) and subsequently azetidine (14.7 µL, 0.217 mmol, 2.4 eq) were added then the reaction mixture was stirred at 100°C for 3.5 h. After cooling to room temperature, the

reaction mixture was filtered through Celite, washed with EtOAc and the solvent was evaporated under reduced pressure. Purification by silica gel column chromatography (Biotage Sfär Duo 5 g, 0–20% EtOAc/cyclohexane, linear gradient) afforded the title compound as a blue solid (35 mg, 21% over 2 steps). $^1$H NMR (400 MHz, $CDCl_3$) δ 8.13 (dd, *J* = 8.1, 1.3 Hz, 1H), 7.97 (d, *J* = 8.0 Hz, 1H), 7.87 (s, 1H), 7.33 – 7.27 (m, 2H), 7.23 (d, *J* = 2.6 Hz, 1H), 7.12 – 7.07 (m, 2H), 7.04 (t, *J* = 7.3 Hz, 1H), 6.88 – 6.82 (m, 2H), 6.79 (dd, *J* = 8.9, 2.7 Hz, 1H), 6.71 (br s, 1H), 6.35 (br s, 1H), 3.93 (t, *J* = 7.4 Hz, 4H), 3.33 (s, 3H), 2.39 (q, *J* = 7.2 Hz, 2H), 1.56 (s, 9H), 0.62 (s, 3H), 0.56 (s, 3H). Analytical HPLC: $t_R$ = 5.6 min, 95% purity (5–95% $CH_3CN/H_2O$, gradient with constant 0.1% formic acid additive, 8 min run, 0.5 mL/min flow, UV detection at 254 nm. HRMS (ESI) calcd for $C_{37}H_{39}N_2O_4Si$ $[M+H]^+$ 603.2674, found 603.2664.

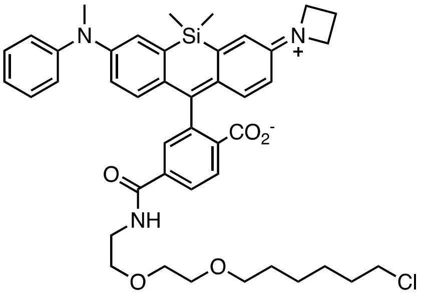

**(20-HTL): S7** (56.3 mg, 0.093 mmol) was dissolved in $CH_2Cl_2$ (1.95 mL). TFA (0.39 mL) was added and the reaction was stirred at room temperature for 6 h. Toluene (2 mL) was added, the reaction mixture evaporated to dryness and then azeotroped with MeOH (3x) to give the crude carboxylic acid as a blue solid. This intermediate, HaloTag(O2) amine (TFA salt, 47.3 mg, 0.140 mmol, 1.5 eq) and HATU (53.3 mg, 0.140 mmol, 1.5 eq) were loaded into a vial. The vial was sealed and evacuated/backfilled with argon. DMF (2 mL) and subsequently DIEA (81 µL, 0.467 mmol, 5 eq) were added. The reaction mixture was stirred at room temperature for 24 h. The solvent was evaporated under reduced pressure before purification by silica gel column chromatography (Biotage Sfär Duo 5 g, 0–100% TBME/cyclohexane, linear gradient) followed by reverse phase HPLC (10–95% $CH_3CN/H_2O$, linear gradient with constant 0.1% v/v TFA additive). The pooled HPLC product fractions were partially concentrated to remove $CH_3CN$ then lyophilized overnight to afford the title compound as a blue solid (24.1 mg, 30%, TFA salt). $^1$H NMR (400 MHz, $CDCl_3$) δ 7.99 (d, *J* = 7.9 Hz, 1H), 7.90 (dd, *J* = 8.0, 1.4 Hz, 1H), 7.74 – 7.72 (m, 1H), 7.30 (t, *J* = 7.9 Hz, 2H), 7.23 (d, *J* = 2.4 Hz, 1H), 7.10 (d, *J* = 7.6 Hz, 2H), 7.04 (t, *J* = 7.3 Hz, 1H), 6.83 – 6.74 (m, 4H), 6.72 (d, *J* = 2.7 Hz, 1H), 6.33 (dd, *J* = 8.7, 2.6 Hz, 1H), 3.93 (t, *J* = 7.3 Hz, 4H), 3.68–3.60 (m, 6H), 3.55 (dd, *J* = 5.8, 3.1 Hz, 2H), 3.50 (t, *J* = 6.6 Hz, 2H), 3.40 (t, *J* = 6.7 Hz, 2H), 3.33 (s, 3H), 2.39 (p, *J* = 7.3 Hz, 2H), 1.76 – 1.67 (m, 2H), 1.52 (p, *J* = 6.9 Hz, 2H), 1.43 – 1.35 (m, 2H), 1.34 – 1.26 (m, 2H), 0.60 (s, 3H), 0.55 (s, 3H). $^{13}$C NMR (101 MHz, $(CD_3)_2CO$) δ 170.0 (CO), 166.1 (CO), 155.8 (C), 152.1 (C), 149.2 (C), 149.1 (C), 141.3 (C), 137.4 (C), 136.8 (C), 136.1 (C), 132.9 (C), 130.3 (CH), 129.0 (C), 128.9 (CH), 128.6 (CH), 128.5 (CH), 126.3 (CH), 124.1 (CH), 124.0 (CH), 123.8 (CH), 123.0 (CH), 119.8 (CH), 116.4 (CH), 113.4 (CH), 91.8 (C), 71.5 ($CH_2$), 70.9 ($CH_2$), 70.8 ($CH_2$), 70.0 ($CH_2$), 53.8 ($CH_2$), 45.8 ($CH_2$), 40.6 ($CH_2$), 40.3 (N-$CH_3$), 33.3 ($CH_2$), 30.3 ($CH_2$), 27.3 ($CH_2$), 26.1 ($CH_2$), 17.4 ($CH_2$), 0.2 (Si-$CH_3$), -1.2 (Si-$CH_3$). Analytical HPLC: $t_R$ = 3.7 min, 97% purity, 5–95% $CH_3CN/H_2O$, gradient with constant 0.1% formic acid additive, 8 min run, 0.5 mL/ min flow, UV detection at 254 nm. HRMS (ESI) calcd for $C_{43}H_{51}ClN_3O_5Si$ $[M+H]^+$ 752.3281, found 752.3268.

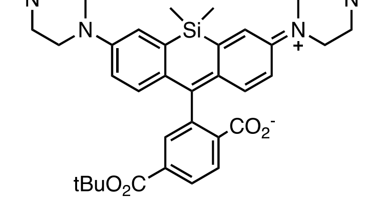

**(S8):** *6*-$CO_2$tBu-Si-fluorescein ditriflate (88.1 mg, 0.119 mmol, 1 eq), $Pd(OAc)_2$ (5.4 mg, 0.024 mmol, 0.2 eq), BINAP (22.3 mg, 0.036 mmol, 0.3 eq) and $Cs_2CO_3$ (109 mg, 0.334 mmol, 2.8 eq) were added to a vial. The vial was sealed and evacuated/backfilled with argon. Toluene (1 mL) and subsequently *N*-methylpiperazine (32 µL, 0.286 mmol, 2.4 eq) were added then the reaction mixture was stirred at 100°C for 48 h. After cooling to room temperature, the reaction mixture was filtered through Celite, washed with MeOH and the solvent was evaporated under reduced pressure. Purification by silica gel column chromatography (10% MeOH/$CH_2Cl_2$) afforded the title compound as a pale blue solid (57.7 mg, 76%). For NMR characterisation, an analytical fraction of the product was further purified by reverse phase HPLC (10–95% $CH_3CN/H_2O$, linear gradient with constant 0.1% v/v TFA additive). The pooled HPLC product fractions were neutralised with saturated $NaHCO_3$ and extracted with EtOAc (3x). The combined organic layers were dried over $Na_2SO_4$, filtered and evaporated to dryness to afford a pure sample. $^1$H NMR (400 MHz, $CD_3OD$) δ 8.13 (dd, *J* = 8.1, 1.3 Hz, 1H), 7.99 (d, *J* = 8.0 Hz, 1H), 7.69 (t, *J* = 1.0 Hz, 1H), 7.29 (t, *J* = 1.6 Hz, 2H), 6.90 – 6.83 (m, 4H), 3.26 (t, *J* = 5.1 Hz, 8H), 2.68 (t, *J* = 5.1 Hz, 8H), 2.40 (s, 6H), 0.68 (s, 3H), 0.58 (s, 3H). $^{13}$C NMR ($(CD_3)_2CO$, 101 MHz) δ 170.3 (CO), 164.6 (CO), 157.0 (C), 150.9 (C), 138.3 (C), 136.3 (C), 135.1 (C), 130.7 (CH), 128.9 (C), 128.3 (CH), 126.5 (CH), 125.0 (CH), 120.9 (CH), 118.0 (CH), 90.9 (C), 82.9 (C), 55.1 ($CH_2$), 48.1 ($CH_2$) 45.4 ($CH_3$), 28.1 ($CH_3$), -0.2 (d, Si-$CH_3$). Analytical HPLC: $t_R$ = 2.9 min, >99% purity, (5–95% $CH_3CN/H_2O$, gradient with constant 0.1% formic acid additive, 8 min run, 0.5 mL/ min flow, UV detection at 254 nm). HRMS (ESI) calcd for $C_{37}H_{47}N_4O_4Si$ [M+H]$^+$ 639.3361, found 639.3347.

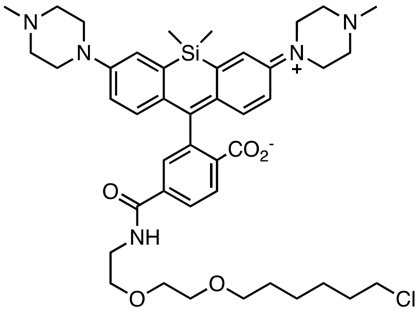

**(5-HTL): S8** (57.7 mg, 0.090 mmol) was dissolved in $CH_2Cl_2$ (2 mL) and TFA (0.4 mL) was added. The reaction was stirred at room temperature for 4 h. Toluene (2 mL) was added, the reaction mixture evaporated to dryness and then azeotroped with MeOH (3x) to give the resulting acid as a blue solid. This acid intermediate (53.9 mg, 0.077 mmol), HaloTag(O2) amine (TFA salt, 39.2 mg, 0.116 mmol, 1.5 eq) and HATU (44.1 mg, 0.116 mmol, 1.5 eq) were loaded into a vial. The vial was sealed and evacuated/backfilled with argon. DMF (1 mL) and subsequently DIEA (67 µL, 0.387 mmol, 5 eq) were added then the reaction mixture was stirred at room temperature for 4.5 h. The reaction mixture was evaporated to dryness before purification by silica gel column chromatography (10% MeOH/$CH_2Cl_2$) followed by reverse phase HPLC (10–50% $CH_3CN/H_2O$, linear gradient with constant 0.1% v/v TFA additive). The pooled HPLC product fractions were partially concentrated to remove $CH_3CN$ then lyophilized overnight to afford the title compound as a blue solid (26.8 mg, 38%, TFA salt). $^1$H NMR ($(CD_3)_2CO$, 400 MHz) δ 8.12 (dd, *J* = 7.95, 1.40 Hz, 1H), 8.01 (dd, *J* = 8.0, 2.0 Hz, 1H), 7.78 (s, 1H), 7.48 (d, *J* = 2.8 Hz, 2H), 7.00 – 6.94 (m, 2H), 6.89 (dd, *J* = 8.8, 1.8 Hz, 2H), 3.70 – 3.16 (m, 16H), 3.13 – 3.03 (m, 4H), 2.92 (s, 6H), 1.80 – 1.66 (m, 2H), 1.49 – 1.25 (m, 8H), 0.72 (s, 3H), 0.58 (s, 3H). $^{13}$C NMR ($CD_3CN$, 101 MHz) δ 170.5 (CO), 166.8 (CO), 155.7 (C), 150.0 (C), 141.8 (C), 137.6 (C), 136.1 (C), 129.2 (CH), 129.1 (CH), 128.5 (C), 126.9 (CH), 123.8 (CH), 121.8 (CH), 118.5 (CH), 91.4 (C), 71.6 ($CH_2$), 70.9 ($CH_2$), 70.8 ($CH_2$), 69.9 ($CH_2$), 53.7 ($CH_2$), 46.5 ($CH_2$), 46.2 ($CH_2$), 43.5 ($CH_3$), 40.7 ($CH_2$), 33.3 ($CH_2$), 30.2 ($CH_2$), 27.3 ($CH_2$), 26.1 ($CH_2$), 0.1 (Si-$CH_3$), -1.00 (Si-$CH_3$). Analytical HPLC: $t_R$ = 2.4 min, 95% purity, 5–95% $CH_3CN/H_2O$, gradient with constant 0.1% formic acid additive, 10 min run, 0.5 mL/min flow, UV detection at 254 nm). HRMS (ESI) calcd for $C_{43}H_{59}ClN_5O_5Si$ [M+H]$^+$ 788.3969, found 788.3953.

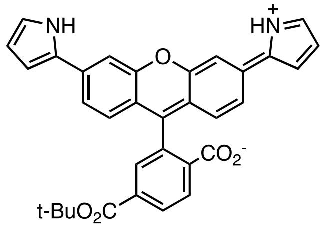

**(S9):** 6-$CO_2$tBu-fluorescein ditriflate (79.5 mg, 0.114 mmol, 1 eq), 1-*N*-Boc-pyrrole-2-boronic acid pinacol ester (100 mg, 0.342 mmol, 3.0 eq), $Pd(PPh_3)_4$ (7.9 mg, 0.069 mmol, 0.06 eq) and KOAc (67.2 mg, 0.685 mmol, 6.0 eq) were loaded into a vial. The vial was sealed and evacuated/backfilled with argon. 1,4-Dioxane/$H_2O$ (2/1, 1.2 mL) was added and the reaction mixture was stirred at 100°C for 68 h. After cooling to room temperature, the reaction mixture was extracted with $CHCl_3$. The organic layers were combined, washed with brine, dried over $Na_2SO_4$ and the solvent was evaporated under reduced pressure. Silica gel column chromatography (30% EtOAc/cyclohexane) afforded the title compound as a blue solid (46.7 mg, 77%). $^1H$ NMR (400 MHz, $CDCl_3$) δ 8.78 (br s, 2H), 8.26 (d, *J* = 8.0 Hz, 1H), 8.09 (d, *J* = 8.0 Hz, 1H), 7.75 (s, 1H), 7.35 – 7.32 (m, 2H), 7.13 (dd, *J* = 8.3, 1.8 Hz, 2H), 6.92 – 6.88 (m, 2H), 6.73 (d, *J* = 8.3 Hz, 2H), 6.61 – 6.54 (m, 2H), 6.32 – 6.28 (m, 2H), 1.53 (s, 9H). $^{13}C$ NMR (101 MHz, $(CD_3)_2CO$) δ 172.3 (CO), 164.7 (CO), 154.1 (C), 152.7 (C), 146.1 (C), 139.3 (C), 137.1 (C), 131.9 (CH), 131.0 (C), 129.5 (CH), 126.0 (CH), 125.5 (CH), 121.2 (CH), 120.4 (CH), 116.3 (C), 111.8 (CH), 110.6 (CH), 108.4 (CH), 83.3 (C), 82.9 (C), 28.1 ($CH_3$). Analytical HPLC: $t_R$ = 5.2 min, 98% purity, 5–95% $CH_3CN/H_2O$, gradient with constant 0.1% formic acid additive, 10 min run, 0.5 mL/ min flow, UV detection at 254 nm. HRMS (ESI) calcd for $C_{33}H_{27}N_2O_5$ $[M+H]^+$ 531.1914, found 531.1919.

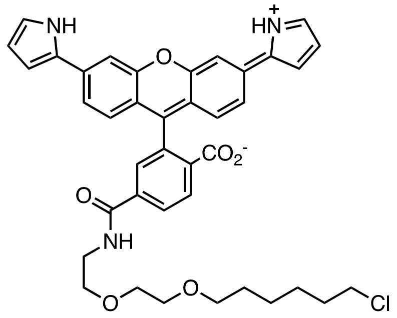

**(8-HTL): S9** (44.2 mg, 0.083 mmol) was dissolved in $CH_2Cl_2$ (2 mL), and TFA (0.4 mL) was added. The reaction was stirred at room temperature for 6 h. Toluene (2 mL) was added, the reaction mixture evaporated to dryness and then azeotroped with MeOH (3x) to give the crude carboxylic acid as a purple solid. This acid intermediate (34.9 mg, 0.059 mmol), HaloTag(O2) amine (TFA salt, 28.8 mg, 0.089 mmol, 1.5 eq) and HATU (33.8 mg, 0.089 mmol, 1.5 eq) were loaded into a vial. The vial was sealed and evacuated/backfilled with argon. DMF (1 mL) and subsequently DIEA (52 µL, 0.297 mmol, 5 eq) were added then the mixture was stirred at room temperature for 3 h. The reaction mixture was evaporated to dryness and purification by silica gel column chromatography (20% EtOAc/cyclohexane) afforded the title compound as a blue solid (38.3 mg, 68%). $^1H$ NMR ($(CD_3)_2CO$, 400 MHz) δ 10.75 ( br s, 2H), 8.25 (dd, *J* = 8.0, 1.4 Hz, 1H), 8.10 (d, *J* = 8.0 Hz, 2H), 7.80 (s, 1H), 7.61 (d, *J* = 1.8 Hz, 2H), 7.41 (dd, *J* = 8.4, 1.8 Hz, 2H), 6.96 – 6.92 (m, 2H), 6.86 (d, *J* = 8.4 Hz, 2H), 6.70 – 6.66 (m, 2H), 6.22 – 6.18 (m, 2H), 3.56 – 3.51 (m, 4H), 3.51 – 3.45 (m, 4H), 3.43 – 3.38 (m, 2H), 3.30 (t, *J* = 6.5 Hz, 2H), 1.72 – 1.63 (m, 2H), 1.46 – 1.22 (m, 6H). $^{13}C$ NMR ($(CD_3)_2CO$, 101 MHz) δ 168.9 (CO), 166.0 (CO), 154.2 (C), 152.6 (C), 142.3 (C), 137.0 (C), 130.9 (C), 130.4 (CH), 129.5 (CH), 129.3 (C), 125.8 (CH), 123.3 (CH), 121.2 (CH), 120.3 (CH), 116.3 (C), 111.7 (CH), 110.6 (CH), 108.3 (CH), 83.2 (C), 71.4 ($CH_2$), 70.8 ($CH_2$), 70.6 ($CH_2$), 70.0 ($CH_2$), 46.7 ($CH_2$), 40.5 ($CH_2$), 33.3 ($CH_2$), 30.2 ($CH_2$), 27.3 ($CH_2$), 26.1 ($CH_2$). Analytical HPLC: $t_R$ = 4.9 min, 99% purity, 5–95% $CH_3CN/H_2O$, gradient with constant 0.1% formic acid additive, 10 min run, 0.5 mL/min flow, UV detection at 254 nm. HRMS (ESI) calcd for $C_{39}H_{39}ClN_3O_6$ $[M+H]^+$ 680.2522, found 680.2515.

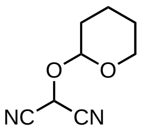

**(S10):** Acetylmalononitrile (1.39 g, 12.9 mmol, 1 eq) was dissolved in $H_2O$ (30 mL), and a peracetic acid solution (9.6% in AcOH, 30 mL) was added. The reaction was stirred at room temperature for 3 h. The solvent was evaporated under reduced pressure. The crude off-white intermediate was resuspended in toluene (33 mL) and cooled to 0°C. *p*-Toluenesulfonic acid (246 mg, 1.29 mmol, 0.1 eq) and 3,4-dihydro-2H-pyran (1.77 mL, 19.4 mmol, 1.5 eq) were added, and the mixture was stirred for 2 h at 0°C. The reaction was diluted with EtOAc and the organic layer was washed with $H_2O$, saturated $NaHCO_3$ and brine. The combined organic layers were dried over $Na_2SO_4$, filtered and evaporated to dryness. Purification by silica gel column chromatography (Biotage Sfär Duo 25 g, 0–50% $CH_2Cl_2$/cyclohexane, linear gradient) afforded the title compound as a white solid (844 mg, 40%). $^1H$ NMR (400 MHz, $CDCl_3$) δ 5.38 (s, 1H), 5.01 (t, *J* = 2.6 Hz, 1H), 3.79 (td, *J* = 10.8, 2.8 Hz, 1H), 3.74 – 3.65 (m, 1H), 1.87 – 1.51 (m, 6H). $^{13}C$ NMR (101 MHz, $CDCl_3$) δ 111.01 (CN), 110.97 (CN), 98.6 (CH), 62.7 ($CH_2$), 51.8 (CH), 29.1 ($CH_2$), 24.7 ($CH_2$), 17.8 ($CH_2$). HRMS (ESI) calcd for $C_8H_9N_2O_2$ $[M-H]^-$ 165.0670, found 165.0672.

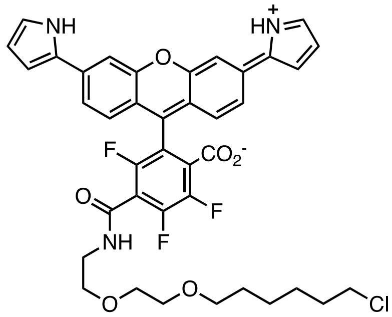

**(9-HTL):** **9** (93.3 mg, 0.186 mmol, 1 eq) and **S10** (28.6 mg, 0.186 mmol, 1 eq) were dissolved in DMF (4.1 mL) under argon, and DIEA (66.3 µL, 0.372 mmol, 2 eq) was added. The reaction was stirred at room temperature for 3 h, then evaporated to dryness. Purification by silica gel chromatography (Biotage Sfär Duo 10 g, 0–50% TBME/cyclohexane, linear gradient) afforded compound **S11** as a dark blue solid. This intermediate showed limited stability and was directly engaged in the next step.

This intermediate **S11** (~106 mg, 0.164 mmol) was dissolved in $CH_2Cl_2$ (10.9 mL). Triethylsilane (2.18 mL) and then trifluoroacetic acid (1.09 mL) were added and the reaction was stirred at room temperature for 3 h. Toluene (5 mL) was added, and the reaction mixture was evaporated to dryness. The residue was added to a solution of HaloTag(O2) amine (TFA salt, 110 mg, 0.333 mmol, 2 eq) and DIEA (292 µL, 1.64 mmol, 10 eq) in $CH_2Cl_2$ (2 mL), and the reaction was stirred at room temperature for 16 h. The solvent was evaporated under reduced pressure and purification was performed by silica gel column chromatography (Biotage Sfär Duo 5 g, 0–60% EtOAc/cyclohexane, linear gradient) followed by reverse phase HPLC (20–80% $CH_3CN/H_2O$, linear gradient with constant 0.1% v/v TFA additive). The pooled HPLC product fractions were combined, neutralised with saturated $NaHCO_3$, and extracted with EtOAc (3x). The organic layers were combined, dried over $Na_2SO_4$ and evaporated to dryness to afford the title compound as a blue-green solid (12.9 mg, 9% over 3 steps). $^1H$ NMR (400 MHz, $CD_3CN$) δ 9.94 (s, 2H), 7.52 (d, *J* = 1.8 Hz, 2H), 7.43 (s, 1H), 7.39 (dd, *J* = 8.3, 1.8 Hz, 2H), 7.12 (d, *J* = 8.3 Hz, 2H), 6.92 (q, *J* = 2.3 Hz, 2H), 6.67 (q, *J* = 2.6 Hz, 2H), 6.23 (q, *J* = 2.7 Hz, 2H), 3.55 – 3.36 (m, 8H), 3.27 (t, *J* = 6.5 Hz, 2H), 1.68 (p, *J* = 6.8 Hz, 2H), 1.44 – 1.18 (m, 8H). $^{19}F$ NMR (376 MHz, $CD_3CN$) δ -123.09 (d, *J* = 21.4 Hz, 1F), -133.62 (d, *J* = 21.5 Hz, 1F), -142.86 (t, *J* = 21.6 Hz, 1F). Analytical HPLC: $t_R$ = 4.5 min, >99% purity, 5–95% $CH_3CN/H_2O$, linear gradient with constant 0.1% formic acid additive, 8 min run, 0.5 mL/min flow, UV detection at 254 nm. HRMS (ESI) calcd for $C_{39}H_{36}ClF_3N_3O_6$ $[M+H]^+$ 734.2239, found 734.2239.

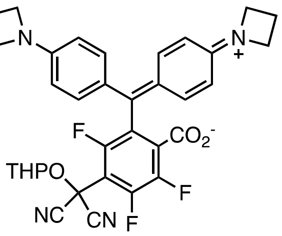

**(S12): 14** (208 mg, 0.444 mmol, 1 eq) and **S10** (68.5 mg, 0.444 mmol, 1 eq) were combined in DMF (9.9 mL) under argon, and DIEA (159 µL, 0.818 mmol, 2 eq) was added. The reaction was stirred at room temperature for 3 h, then evaporated to dryness. Purification by silica gel column chromatography (Biotage Sfär Duo 10 g, 0–30% EtOAc/cyclohexane) afforded the title compound as a blue-green solid (145 mg, 53%). For characterization by NMR spectroscopy, an analytical fraction of the product was further purified by silica gel column chromatography (Biotage Sfär Duo 5 g, 0–25% EtOAc/cyclohexane). $^{1}H$ NMR (400 MHz, $(CD_3)_2CO$) δ 7.09 – 7.14 (m, 4H), 6.38 – 6.43 (m, 4H), 4.70 (t, *J* = 3.0 Hz, 1H), 3.88 (t, *J* = 7.2 Hz, 8H), 3.67 – 3.75 (m, 1H), 3.50 – 3.57 (m, 1H), 2.36 (p, *J* = 7.3 Hz, 4H), 1.90 – 1.81 (m, 2H), 1.75 – 1.50 (m, 4H). $^{19}F$ NMR (376 MHz, $(CD_3)_2CO$) δ -112.62 (d, *J* = 21.8 Hz, 1F), -129.26 (d, *J* = 19.8 Hz, 1F), -141.66 (t, *J* = 20.8 Hz, 1F). HRMS (ESI) calcd for $C_{34}H_{30}F_3N_4O_4$ $[M+H]^+$ 615.2214, found 615.2196.

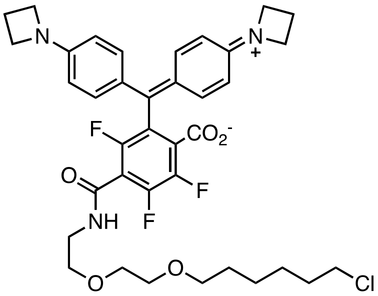

**(14-HTL): S12** (114 mg, 0.186 mmol) was dissolved in $CH_2Cl_2$ (12.4 mL) then triethylsilane (1.24 mL) and trifluoroacetic acid (2.5 mL) were added. The reaction was stirred at room temperature for 2 h. Toluene (5 mL) was added, and the reaction mixture was evaporated to dryness. The residue was added to a solution of HaloTag(O2) amine (TFA salt, 126 mg, 0.373 mmol, 2 eq) and DIEA (333 µL, 1.86 mmol, 10 eq) in $CH_2Cl_2$ (3 mL), and the reaction was stirred at room temperature for 15 h. The solvent was evaporated under reduced pressure and purification was performed by silica gel column chromatography (Biotage Sfär Duo 5 g, 0–60% EtOAc/cyclohexane, linear gradient) followed by reverse phase HPLC (20–80% $CH_3CN/H_2O$, linear gradient with constant 0.1% v/v TFA additive). The pooled HPLC product fractions were combined, neutralised with saturated $NaHCO_3$, and extracted with EtOAc (3x). The organic layers were combined, dried over $Na_2SO_4$ and evaporated to dryness to afford the title compound as a blue-green solid (8.9 mg, 7%). $^{1}H$ NMR (400 MHz, $(CD_3)_2SO$), 5 mM DMF) δ 9.14 (t, *J* = 5.6 Hz, 1H), 7.02 (d, *J* = 8.3 Hz, 4H), 6.39 (d, *J* = 8.3 Hz, 4H), 3.81 (t, *J* = 7.2 Hz, 8H), 3.60 (t, *J* = 6.6 Hz, 2H), 3.51 (m, 4H), 3.46 (m, 2H), 3.40 (q, *J* = 5.6 Hz, 2H), 2.29 (q, *J* = 7.3 Hz, 4H), 1.68 (p, *J* = 6.8 Hz, 2H), 1.46 (p, *J* = 6.7 Hz, 2H), 1.31 (m, 6H). $^{19}F$ NMR (376 MHz, $CD_3CN$) δ -118.58 (d, *J* = 21.8 Hz, 1F), -135.14 (d, *J* = 21.3 Hz, 1F), -143.24 (t, *J* = 21.6 Hz, 1F). Analytical HPLC: $t_R$ = 4.8 min, >99% purity, 5–95% $CH_3CN/H_2O$, gradient with constant 0.1% formic acid additive, 8 min run, 0.5 mL/min flow, UV detection at 254 nm. HRMS (ESI) calcd for $C_{37}H_{42}ClF_3N_3O_5$ $[M+H]^+$ 700.2760, found 700.2743.

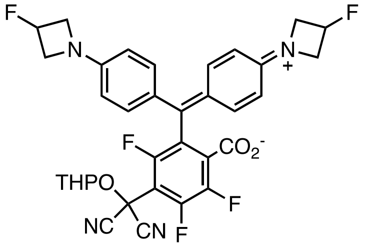

**(S13): 15** (102 mg, 0.202 mmol, 1 eq) and **S10** (31.2 mg, 0.202 mmol, 1 eq) were combined in DMF (4.5 mL) under argon, and DIEA (72 µL, 0.404 mmol, 2 eq) was added. The reaction was stirred at room temperature for 3.5 h, then evaporated to dryness. Purification by silica gel column chromatography (Biotage Sfär Duo 10 g, 0–30% EtOAc/cyclohexane, linear gradient) afforded the title compound as a blue-green solid (79.2 mg, 60%). For characterization by NMR spectroscopy, an analytical fraction of the product was further purified by silica gel column chromatography (Biotage Sfär Duo 5 g, 0–60% $CH_2Cl_2$/cyclohexane followed by Biotage Sfär Duo 5 g, 0–25% EtOAc/cyclohexane, linear gradient). $^1H$ NMR (400 MHz, $(CD_3)_2CO$) δ 7.16 (d, $J$ = 8.7, 4H), 6.56 – 6.47 (m, 4H), 5.50 (dtt, $^2J_{HF}$ = 57.3 Hz, $J$ = 8.8, 3.0 Hz, 2H), 5.47 (t, $J$ = 3.1 Hz, 1H), 4.30 – 4.19 (m, 4H), 4.01 – 3.88 (m, 4H), 3.75 – 3.67 (m, 1H), 3.57 – 3.50 (m, 1H), 1.90 – 1.78 (m, 2H), 1.75 – 1.51 (m, 4H). $^{19}F$ NMR (376 MHz, $CD_3)_2CO$) δ -112.64 (d, $J$ = 21.8 Hz), -129.01 (d, $J$ = 20.4 Hz), -141.46 (t, $J$ = 20.8 Hz), -179.60 – -179.80 (m, 2F). HRMS (ESI) calcd for $C_{34}H_{28}F_5N_4O_4$ $[M+H]^+$ 651.2025, found 651.2009.

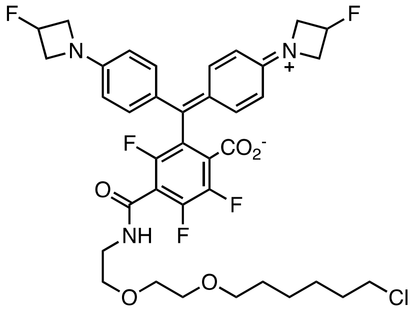

**(15-HTL): S13** (16.6 mg, 0.026 mmol) was dissolved in $CH_2Cl_2$ (3.4 mL), triethylsilane (0.34 mL) and trifluoroacetic acid (0.68 mL) were added. The reaction was stirred at room temperature for 1 h. Toluene (2 mL) was added, and the reaction mixture was evaporated to dryness. The residue was added to a solution of HaloTag(O2) amine (TFA salt, 17.2 mg, 0.051 mmol, 2 eq) and DIEA (46 µL, 0.255 mmol, 10 eq) in $CH_2Cl_2$ (1.5 mL), and the reaction was stirred at room temperature for 16 h. The solvent was evaporated under reduced pressure and purification was performed by silica gel column chromatography (Biotage Sfär Duo 5 g, 0–60% EtOAc/cyclohexane, linear gradient) followed by reverse phase HPLC (10–95% $CH_3CN/H_2O$, linear gradient with constant 0.1% v/v TFA additive). The pooled HPLC product fractions were combined, neutralised with saturated $NaHCO_3$, and extracted with EtOAc (3x). The organic layers were combined, dried over $Na_2SO_4$ and evaporated to dryness to afford the title compound as a blue-green solid (5.8 mg, 30%). $^1H$ NMR (400 MHz, $(CD_3)_2SO$, 5 mM DMF) δ 9.18 (t, $J$ = 5.8 Hz, 1H), 7.06 (d, $J$ = 8.3 Hz, 4H), 6.47 (d, $J$ = 8.3 Hz, 4H), 5.47 (m, 2H), 4.16 (m, 4H), 3.89 (ddd, $J$ = 24.2, 9.5, 3.1 Hz, 4H), 3.55 – 3.27 (m, 12H), 1.67 (p, $J$ = 6.9 Hz, 2H), 1.46 (p, $J$ = 6.8 Hz, 2H), 1.25 (m, $J$ = 11.9 Hz, 2H), 0.85 (t, $J$ = 6.6 Hz, 2H). $^{19}F$ NMR (376 MHz, $CD_3CN$) δ -118.51 (dd, $J$ = 22.2 Hz, 2.1 Hz, 1F), -134.89 (dd, $J$ = 21.1 Hz, 2.5 Hz, 1F), -143.01 (t, $J$ = 21.8 Hz, 1F), -179.95 – -179.98 (m, 2F). Analytical HPLC: $t_R$ = 4.5 min, 98% purity, 5–95% $CH_3CN/H_2O$, gradient with constant 0.1% formic acid additive, 8 min run, 0.5 mL/min flow, UV detection at 254 nm. HRMS (ESI) calcd for $C_{37}H_{40}ClF_5N_3O_5$ $[M+H]^+$ 736.2571, found 736.2554.

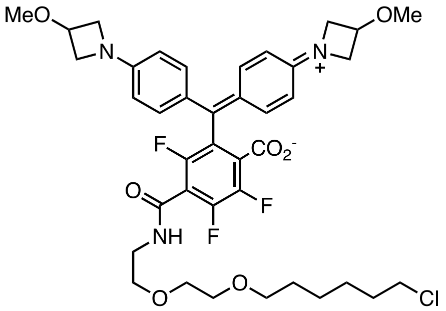

**(16-HTL)**: **16** (656 mg, 1.24 mmol, 1 eq) and **S10** (191 mg, 1.24 mmol, 1 eq) were combined in DMF (27.5 mL) under argon, and DIEA (443 µL, 2.48 mmol, 2 eq) was added. The reaction was stirred at room temperature for 4 h, then evaporated to dryness. Purification by silica gel column chromatography (Biotage Sfär Duo 25 g, 0–30% EtOAc/cyclohexane, linear gradient) afforded the compound **S14** as a blue-green solid. This intermediate showed limited stability and was directly engaged in the next step.

**S14** (~455 mg, 0.675 mmol) was taken up in $CH_2Cl_2$ (45 mL), triethylsilane (4.50 mL) and trifluoroacetic acid (9 mL) were added. The reaction was stirred at room temperature for 2 h. Toluene (5 mL) was added, and the reaction mixture was evaporated to dryness. The residue was added to a solution of HaloTag(O2) amine (TFA salt, 455 mg, 1.35 mmol, 2 eq) and DIEA (1.21 mL, 6.75 mmol, 10 eq) in $CH_2Cl_2$ (30 mL), and the reaction was stirred at room temperature for 15 h. The solvent was evaporated under reduced pressure and purification was performed by silica gel column chromatography (Biotage Sfär Duo 10 g, 0–50% EtOAc/cyclohexane, linear gradient) followed by reverse phase HPLC (10–100% $CH_3CN/H_2O$, linear gradient with constant 0.1% v/v TFA additive). The pooled HPLC product fractions were combined, neutralised with saturated $NaHCO_3$, and extracted with EtOAc (3x). The organic layers were combined, dried over $Na_2SO_4$ and evaporated to dryness to afford the title compound as a blue-green solid (243 mg, 26% over 3 steps). $^1H$ NMR (400 MHz, $(CD_3)_2CO$) δ 8.36 (br s, 1H), 7.11 (d, $J$ = 8.7 Hz, 4H), 6.44 (d, $J$ = 8.7 Hz, 4H), 4.37 – 4.31 (m, 2H), 4.12 – 4.08 (m, 4H), 3.70 – 3.65 (m, 4H), 3.65 – 3.60 (m, 2H), 3.60 - 3.54 (m, 6H), 3.51 – 3.48 (m, 2H), 3.39 (t, $J$ = 6.5 Hz, 2H), 3.28 (s, 6H), 1.74 (p, $J$ = 5.6 Hz, 2H), 1.50 (p, $J$ = 6.8 Hz, 2H), 1.45 – 1.28 (m, 4H). $^{19}F$ NMR (376 MHz, $(CD_3)_2SO$) δ -118.77 (d, $J$ = 21.8 Hz, 1F), -134.76 – -134.98 (d, $J$ = 23.2 Hz, 1F), -143.10 (t, $J$ = 22.0 Hz, 1F). Analytical HPLC: $t_R$ = 4.5 min, 98.5 % purity, 5–95% $CH_3CN/H_2O$, gradient with constant 0.1% formic acid additive, 8 min run, 0.5 mL/min flow, UV detection at 254 nm. HRMS (ESI) calcd for $C_{39}H_{46}ClF_3N_3O_7$ $[M+H]^+$ 760.2971, found 760.2950.

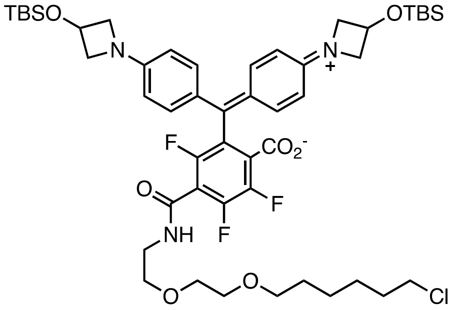

**(S16): S6** (159 mg, 0.288 mmol, 1 eq) and **S10** (44.4 mg, 0.288 mmol, 1 eq) were combined in DMF (6 mL) under argon, and DIEA (103 µL, 0.576 mmol, 2 eq) was added. The reaction was stirred at room temperature for 3 h, then evaporated to dryness. Purification by silica gel column chromatography (Biotage Sfär Duo 10 g, 0–15% EtOAc/cyclohexane, linear gradient) afforded compound **S15** as a green solid. This intermediate showed limited stability and was directly engaged in the next step.

**S15** (~85 mg, 0.098 mmol) was taken up in $CH_2Cl_2$ (6.5 mL), triethylsilane (650 µL) and trifluoroacetic acid (1.3 mL) were added. The reaction was stirred at room temperature for 4 h. Toluene (2 mL) was added, and the reaction mixture was evaporated to dryness. The residue was added to a solution of HaloTag(O2) amine (TFA salt, 65.9 mg, 0.195 mmol, 2 eq) and DIEA (175 µL, 0.977 mmol, 10 eq) in $CH_2Cl_2$ (2 mL), and the reaction was stirred at room temperature for 15 h. The solvent was evaporated under reduced pressure and purification was performed by silica gel column chromatography (Biotage Sfär Duo 5 g, 0–30% EtOAc/cyclohexane, linear gradient) to afford the title compound as a green solid (23.6 mg, 9% over 3 steps). $^1$H NMR (400 MHz, $CD_3CN$) δ 7.35 (t, *J* = 5.5 Hz, 1H), 7.07 (d, *J* = 8.3 Hz, 4H), 6.41 (d, *J* = 8.3 Hz, 4H), 4.76 (tt, *J* = 6.2, 4.7 Hz, 2H), 4.16 – 4.10 (m, 4H), 3.60 – 3.45 (m, 12H), 3.41 (t, *J* = 6.5 Hz, 2H), 3.34 (t, *J* = 6.5 Hz, 2H), 1.79 – 1.66 (m, 2H), 1.54 (p, *J* = 6.9 Hz, 2H), 1.48 – 1.24 (m, 4H), 0.89 (s, 18H), 0.08 (s, 12H). $^{19}$F NMR (376 MHz, $CD_3CN$) δ -118.45 (d, *J* = 21.8 Hz, 1F), -135.11 (d, *J* = 21.1 Hz, 1F), -143.25 (t, *J* = 21.7 Hz, 1F). Analytical HPLC: $t_R$ = 7.5 min, 95% purity, 5–95% $CH_3CN/H_2O$, gradient with constant 0.1% formic acid additive, 8 min run, 0.5 mL/min flow, UV detection at 254 nm. HRMS (ESI) calcd for $C_{49}H_{70}F_3N_3O_7Si_2$ [M+H]$^+$ 960.4387, found 960.4364.

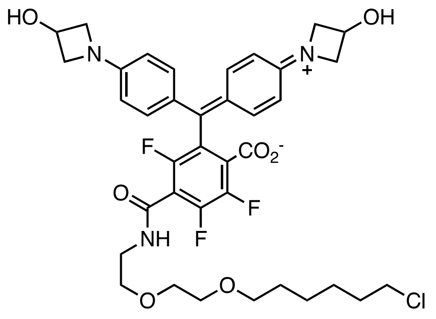

**(21-HTL): S16** (6.9 mg, 0.007 mmol, 1 eq) in THF (1 mL) was cooled to 0°C and a solution of tetrabutylammonium fluoride (36 mM in THF, 1 mL, 5 eq) was added. The reaction was stirred at 0°C for 1 h. Saturated $NH_4Cl$ was added, and the mixture was extracted with EtOAc (3x). The combined organic layers were washed with brine, dried over anhydrous $Na_2SO_4$, filtered, and evaporated to dryness. Purification was performed by silica gel chromatography (Biotage Sfär Duo 5 g, 0–5% MeOH/$CH_2Cl_2$, linear gradient) followed by reverse phase HPLC (10–100% $CH_3CN/H_2O$, linear gradient with constant 0.1% v/v TFA additive). The pooled HPLC product fractions were combined, neutralised with saturated $NaHCO_3$, and extracted with EtOAc (3x). The combined organic layers were dried over $Na_2SO_4$ and evaporated to dryness to afford the title compound as a blue solid (1.4 mg, 27%). $^1$H NMR (400 MHz, $(CD_3)_2SO$) δ 9.15 (t, *J* = 5.6 Hz, 1H), 7.02 (d, *J* = 8.5 Hz, 4H), 6.61 (br s, 2H), 6.41 (d, *J* = 8.5 Hz, 4H), 5.67 – 5.58 (m, 2H), 4.59 – 4.51 (m, 4H), 4.07 (t, *J* = 7.2 Hz, 4H), 3.62 – 3.43 (m, 12H), 1.67 (p, *J* = 7.0 Hz, 2H), 1.47 (p, *J* = 7.0 Hz, 2H), 1.39 – 1.10 (m, 4H). $^{19}$F NMR (376 MHz, $(CD_3)_2SO$) δ -118.79 (d, *J* = 20.9 Hz), -134.87 (d, *J* = 23.5 Hz), -143.05 – -143.29 (m, 1F). Analytical HPLC: $t_R$ = 3.9 min, >99% purity, 5–95% $CH_3CN/H_2O$, gradient with constant 0.1% formic acid additive, 8 min run, 0.5 mL/min flow, UV detection at 254 nm. HRMS (ESI) calcd for $C_{37}H_{42}ClF_3N_3O_7$ [M+H]$^+$ 732.2658, found 732.2637.

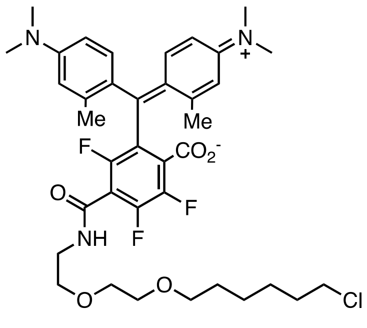

**(18-HTL): 18** (59.7 mg, 0.126 mmol, 1 eq) and **S10** (19.5 mg, 0.126 mmol, 1 eq) were combined in DMF (3 mL) under argon, and DIEA (45 µL, 0.252 mmol, 2 eq) was added. The reaction was stirred at room temperature for 3 h, then evaporated to dryness. Purification by silica gel column chromatography (Biotage Sfär Duo 5 g, 0–20% EtOAc/cyclohexane, linear gradient) afforded compound **S17** as a blue-green solid. This intermediate showed limited stability and was directly engaged in the next step.

This intermediate **S17** (~57 mg, 0.093 mmol) was dissolved in $CH_2Cl_2$ (6.2 mL). Triethylsilane (620 µL) and then trifluoroacetic acid (1.24 mL) were added and the reaction was stirred at room temperature for 3 h. Toluene (5 mL) was added, and the reaction mixture was evaporated to dryness. The residue was added to a solution of HaloTag(O2) amine (TFA salt, 62.6 mg, 0.186 mmol, 2 eq) and DIEA (165 µL, 0.932 mmol, 10 eq) in $CH_2Cl_2$ (2 mL), and the reaction was stirred at room temperature for 16 h. The solvent was evaporated under reduced pressure and purification was performed by silica gel column chromatography (Biotage Sfär Duo 5 g, 0–50% EtOAc/cyclohexane, linear gradient) followed by reverse phase HPLC (10–95% $CH_3CN/H_2O$, linear gradient with constant 0.1% v/v TFA additive). The pooled HPLC product fractions were combined, neutralised with saturated $NaHCO_3$, and extracted with EtOAc (3x). The organic layers were combined, dried over $Na_2SO_4$ and evaporated to dryness to afford the title compound as a blue-green solid (20.1 mg, 23% over 3 steps). $^1$H NMR (400 MHz, $(CD_3)_2CO$) δ 8.18 – 8.25 (m, 1H), 6.83 (d, *J* = 8.2 Hz, 2H), 6.61 (s, 2H), 6.46 (dd, *J* = 8.8, 2.8 Hz, 2H), 3.64 – 3.60 (m, 2H), 3.60 – 3.52 (m, 6H), 3.52 – 3.47 (m, 2H), 3.39 (t, *J* = 6.5 Hz, 2H), 2.94 (s, 12H), 2.07 (s, 6H), 1.78 – 1.69 (m, 2H), 1.55 – 1.47 (m, 2H), 1.47 – 1.26 (m, 4H). $^{19}$F NMR (376 MHz, $CD_3CN$) δ -118.61 (d, *J* = 21.4 Hz, 1F), -135.00 (dd, *J* = 21.1, 2.8 Hz, 1F), -143.75 (t, *J* = 21.7 Hz, 1F). Analytical HPLC: $t_R$ = 4.8 min, >99% purity, 5–95% $CH_3CN/H_2O$, gradient with constant 0.1% formic acid additive, 8 min run, 0.5 mL/min flow, UV detection at 254 nm. HRMS (ESI) calcd for $C_{37}H_{46}ClF_3N_3O_5$ [M+H]$^+$ 704.3073, found 704.3052.

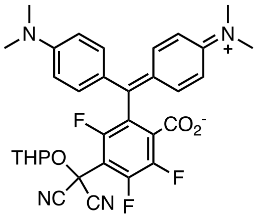

**S18**: **13** (253 mg, 0.570 mmol, 1 eq) and **S10** (87.8 mg, 0.570 mmol, 1 eq) were combined in DMF (12.6 mL) under argon, and DIEA (203 µL, 1.139 mmol, 2 eq) was added. The reaction was stirred at room temperature for 4 h, then evaporated to dryness. Purification by silica gel column chromatography (Biotage Sfär Duo 10 g, 0–30% EtOAc/cyclohexane, linear gradient) afforded the title compound as a blue-green solid (225 mg, 67%). For characterization by NMR spectroscopy, an analytical fraction of the product was further purified by silica gel column chromatography (Biotage Sfär Duo 10 g, 0–25% EtOAc/cyclohexane, linear gradient). $^1$H NMR (400 MHz, $(CD_3Cl)$ δ 7.14 (d, *J* = 8.4 Hz, 4H), 6.69 (d, *J* = 8.1 Hz, 4H), 5.52 (m, 1H), 3.90 – 3.81 (m, 1H), 3.68 (d, *J* = 13.3 Hz, 1H), 2.97 (s, 12H), 1.90 – 1.53 (m, 6H). $^{19}$F NMR (376 MHz, $CD_3Cl$) δ -112.41 (d, *J* = 22.5 Hz, 1F), -127.44 (d, *J* = 20.1 Hz, 1F), -139.61 (t, *J* = 21.5 Hz, 1F). HRMS (ESI) calcd for $C_{32}H_{30}F_3N_4O_4$ [M+H]$^+$ 591.2214, found 591.2211.

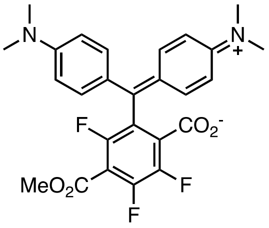

**S19: S18** (616 mg, 1.04 mmol) was taken up in $CH_2Cl_2$ (69 mL), triethylsilane (7 mL) and trifluoroacetic acid (14 mL) were added. The reaction was stirred at room temperature for 4 h. Toluene (10 mL) was added, and the reaction mixture was evaporated to dryness. To the residue was added a solution of methanol (1.06 mL, 26.1 mmol, 25 eq) and triethylamine (1.45 mL, 10.4 mmol, 10 eq) in $CH_2Cl_2$ (12 mL), and the reaction was stirred at room temperature for 15 h. The solvent was evaporated under reduced pressure and purification was performed by silica gel column chromatography (Biotage Sfär Duo 25 g, 0–20% EtOAc/cyclohexane, linear gradient) to afford the title compound as a blue-green solid (161 mg, 32%). $^1$H NMR (400 MHz, $CDCl_3$) δ 7.14 (d, *J* = 8.4 Hz, 4H), 6.65 (d, *J* = 8.4 Hz, 4H), 3.97 (s, 3H), 2.96 (s, 12H). $^{19}$F NMR (376 MHz, $CDCl_3$) δ -115.33 (d, *J* = 22.6 Hz), -131.62 (d, *J* = 17.2 Hz, 1F), -141.97 (t, *J* = 20.0 Hz, 1F). Analytical HPLC: $t_R$ = 5.2 min, 97.3% purity, 5–95% $CH_3CN/H_2O$, gradient with constant 0.1% formic acid additive, 8 min run, 0.5 mL/min flow, UV detection at 254 nm. HRMS (ESI) calcd for $C_{26}H_{24}F_3N_2O_4$ [M+H]$^+$ 485.1683, found 485.1680.

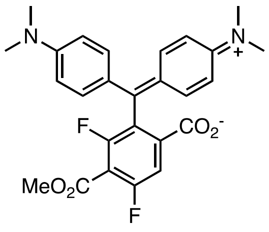

**S20: S19** (211 mg, 0.437 mmol, 1 eq) was loaded in a vial which was sealed and evacuated/backfilled with argon. DMF (3 mL) was added, followed by dropwise addition of a solution of $NaBH_4$ (82.6 mg, 2.18 mmol, 5 eq) in DMF (1.5 mL). The reaction was stirred at room temperature for 4 h and carefully quenched with $H_2O$. The mixture was extracted with $CH_2Cl_2$ (3x), the organic layers were combined, washed with brine, dried over $Na_2SO_4$, filtered and evaporated to dryness. Purification was performed by silica gel column chromatography (Biotage Sfär Duo 10 g, 0–15% EtOAc/cyclohexane, linear gradient) to afford the title compound as a blue-green solid (87.5 mg, 43%). $^1$H NMR (400 MHz, $CDCl_3$) δ 7.50 (d, *J* = 7.1 Hz, 1H), 7.15 (d, J = 8.6 Hz, 4H), 6.65 (d, *J* = 8.6 Hz, 4H), 3.96 (s, 3H), 2.96 (s, 12H). $^{19}$F NMR (376 MHz, $CDCl_3$) δ -107.66 (s, 1F), -109.96 (d, *J* = 4.7 Hz, 1F). Analytical HPLC: $t_R$ = 5.0 min, 5–95% $CH_3CN/H_2O$, gradient with constant 0.1% formic acid additive, 8 min run, 0.5 mL/min flow, UV detection at 254 nm. Purity was estimated at 90% based on crystallography. HRMS (ESI) calcd for $C_{26}H_{24}F_2N_2O_4$ $[M+H]^+$ 467.1777, found 467.1776.

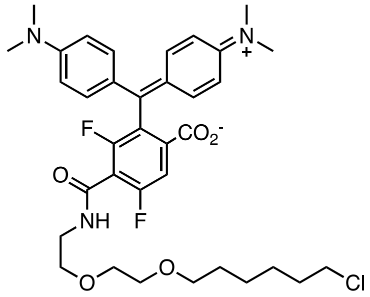

**(22-HTL): S20** (117 mg, 0.250 mmol, 1 eq) was dissolved in 1:1 THF:MeOH (22 mL) under argon. 1M NaOH (1 mL, 4 eq) was added dropwise and the reaction stirred at room temperature for 24 h. The reaction was neutralised (1M HCl), diluted in $H_2O$ and the solvents evaporated. The mixture was extracted with i-PrOH/$CHCl_3$ 15% (3x), before the organic layers were combined, dried over $Na_2SO_4$, filtered and evaporated to dryness. This crude intermediate, HaloTag(O2) amine (TFA salt, 843 mg, 2.50 mmol, 10 eq), EDC.HCl (359 mg, 1.87 mmol, 7.5 eq) and HOBt (152 mg, 1.13 mmol, 4.5 eq) were loaded into a sealed vial, which was evacuated/backfilled under argon. $CH_2Cl_2$ (5 mL) and then DIEA (0.46 mL, 2.6 mmol, 10 eq) were added and the reaction stirred at room temperature for 22 h. Saturated $NaHCO_3$ was added, and the mixture was extracted with $CH_2Cl_2$ (3x). The organic layers were combined, dried over $Na_2SO_4$, filtered and evaporated to dryness. Purification was performed by silica gel column chromatography (Biotage Sfär Duo 10 g, 0–50% EtOAc/cyclohexane, linear gradient) followed by reverse phase HPLC (20–80% $CH_3CN/H_2O$, linear gradient with constant 0.1% v/v TFA additive). The pooled HPLC product fractions were neutralised with saturated $NaHCO_3$ and extracted with EtOAc (3x). The combined organic layers were dried over $Na_2SO_4$, filtered and evaporated to dryness to afford the title compound as a blue-green solid (39 mg, 23%). $^1$H NMR (400 MHz, $(CD_3)_2SO$, 5 mM DMF) δ 9.09 (t, *J* = 5.5 Hz, 1H), 7.83 (d, *J* = 6.9 Hz, 1H), 7.02 (d, *J* = 8.7 Hz, 4H), 6.70 (d, J = 8.7 Hz, 4H), 3.59 (t, *J* = 6.6 Hz, 2H), 3.54 – 3.49 (m, 4H), 3.48 – 3.44 (m, 2H), 3.40 (q, *J* = 5.6 Hz, 2H), 2.89 (d, *J* = 5.2 Hz, 12H), 1.67 (p, *J* = 6.7 Hz, 2H), 1.45 (p, *J* = 6.8 Hz, 2H), 1.39 – 1.21 (m, 6H). $^{19}$F NMR (376 MHz, $(CD_3)_2SO$) δ -110.15 (d, *J* = 6.2 Hz, 1F), -113.84 (d, *J* = 6.1 Hz, 1F). Analytical HPLC: $t_R$ = 4.6 min, ≥99% purity, 5–95% $CH_3CN/H_2O$, gradient with constant 0.1% formic acid additive, 8 min run, 0.5 mL/min flow, UV detection at 254 nm. HRMS (ESI) calcd for $C_{35}H_{43}ClF_2N_3O_5$ $[M+H]^+$ 658.2854, found 658.2845.

### NMR Spectra and HPLC Traces

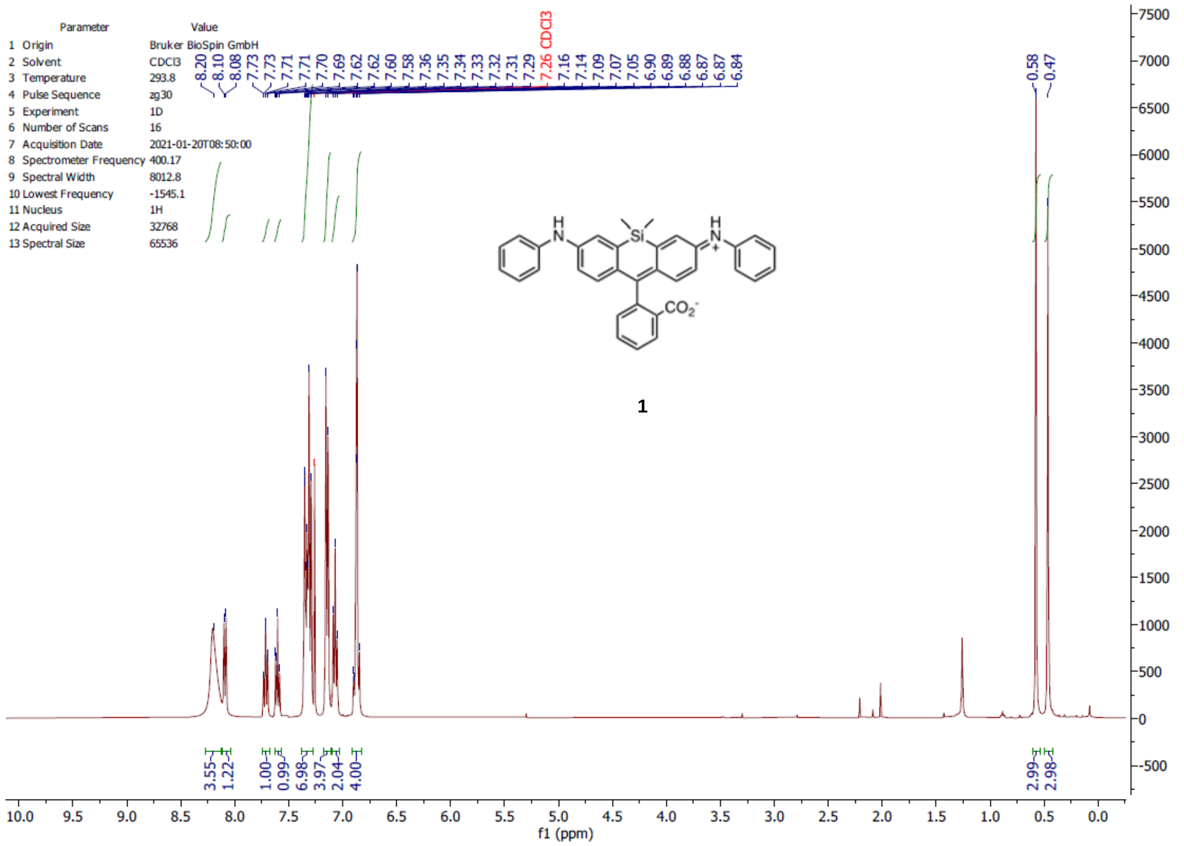

1

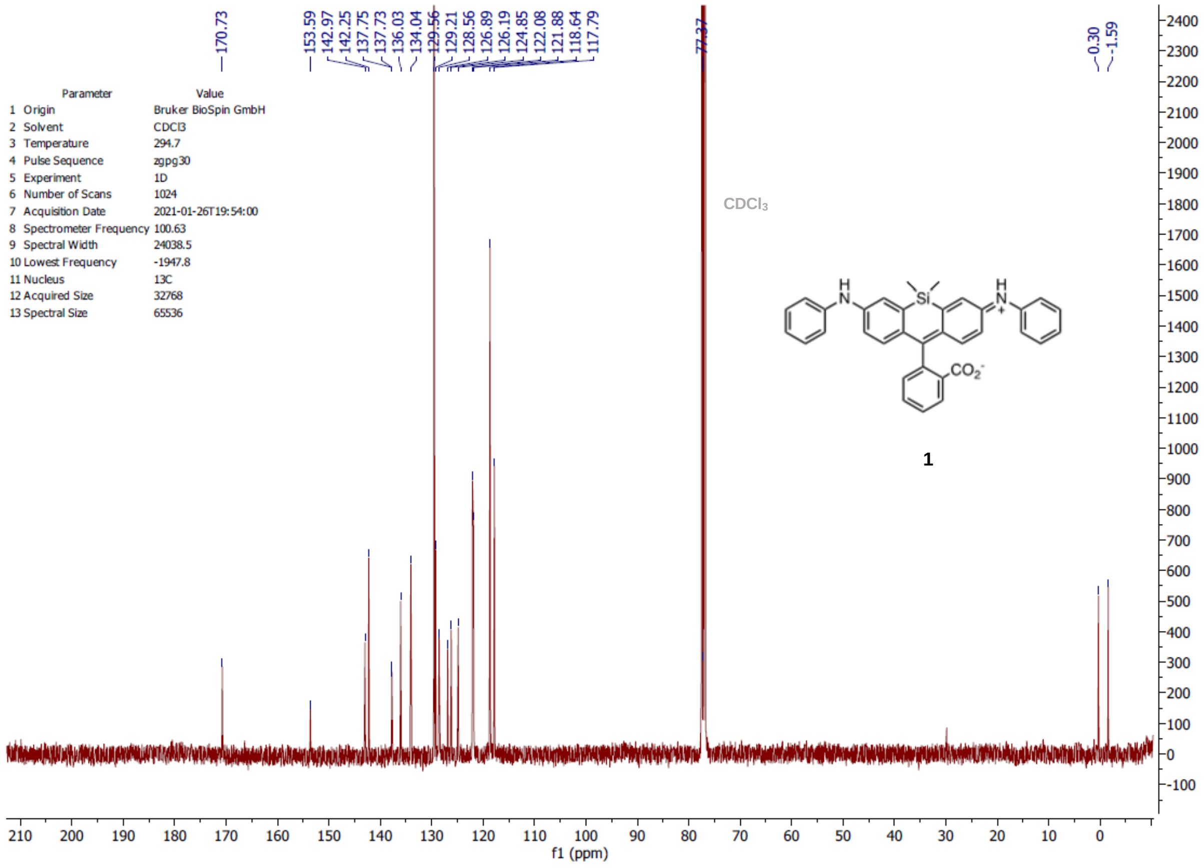

1

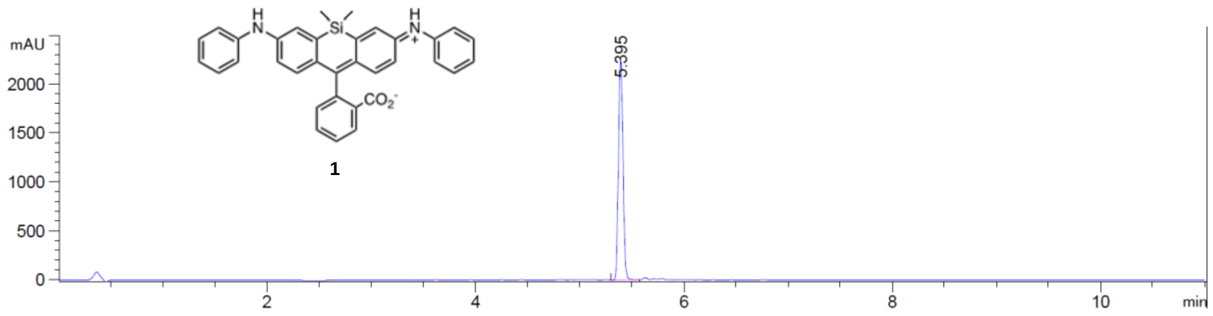
mAU
2000
1500
1000
500
0
5.395
2
4
6
8
10
min
1

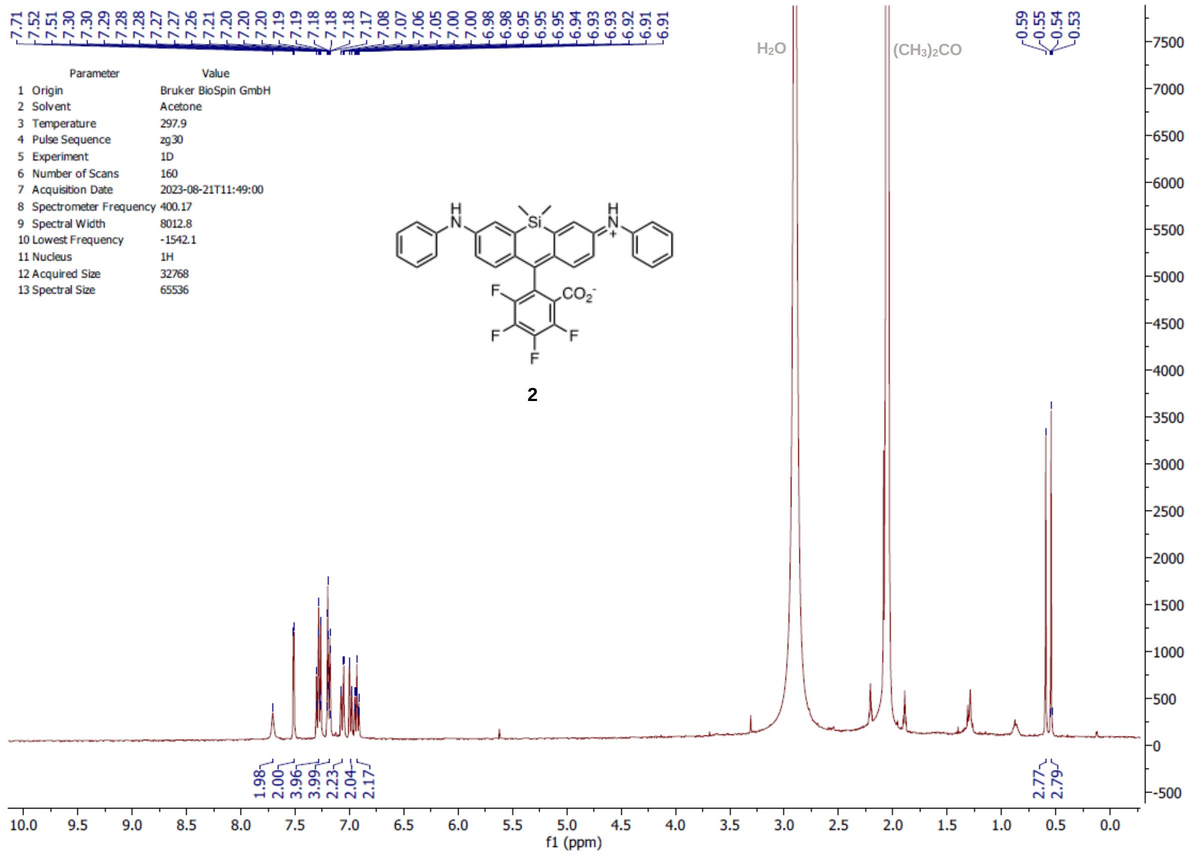
Parameter Value
1 Origin Bruker BioSpin GmbH
2 Solvent Acetone
3 Temperature 297.9
4 Pulse Sequence zg30
5 Experiment 1D
6 Number of Scans 160
7 Acquisition Date 2023-08-21T11:49:00
8 Spectrometer Frequency 400.17
9 Spectral Width 8012.8
10 Lowest Frequency -1542.1
11 Nucleus 1H
12 Acquired Size 32768
13 Spectral Size 65536
H2O
(CH3)2CO
2
f1 (ppm)

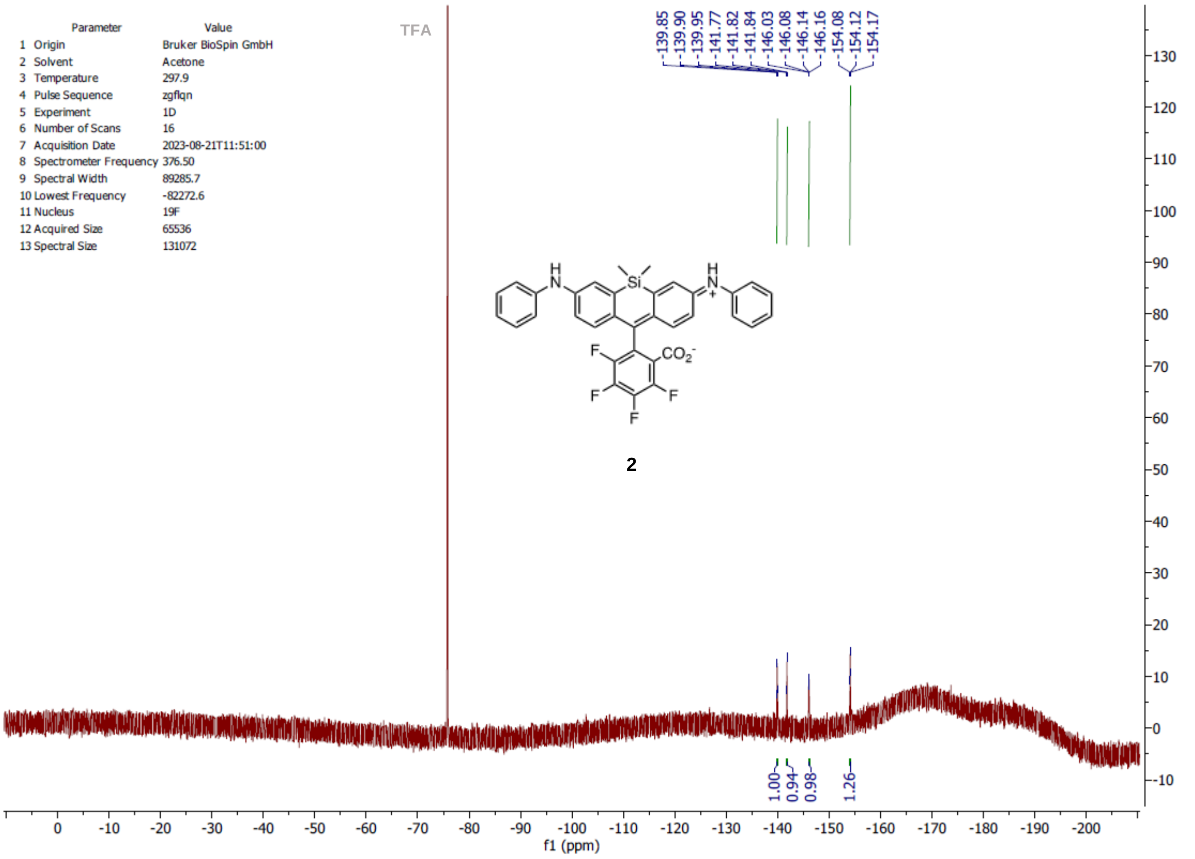
Parameter Value
1 Origin Bruker BioSpin GmbH
2 Solvent Acetone
3 Temperature 297.9
4 Pulse Sequence zgflqn
5 Experiment 1D
6 Number of Scans 16
7 Acquisition Date 2023-08-21T11:51:00
8 Spectrometer Frequency 376.50
9 Spectral Width 89285.7
10 Lowest Frequency -82272.6
11 Nucleus 19F
12 Acquired Size 65536
13 Spectral Size 131072
TFA
2
f1 (ppm)

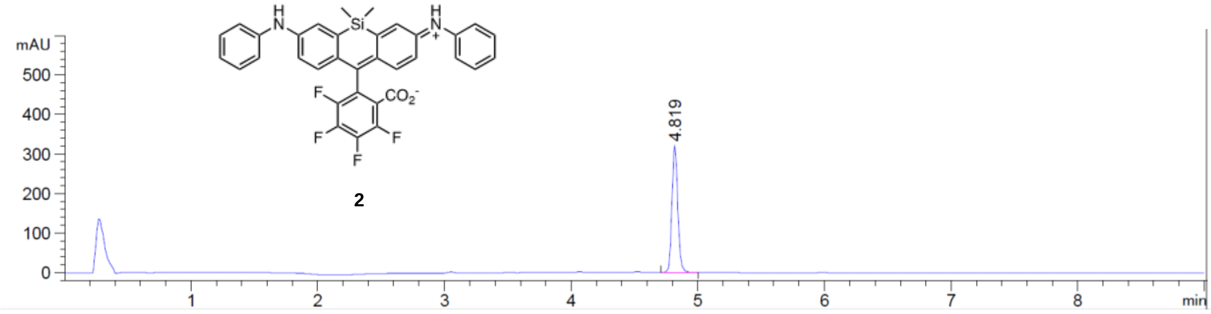

2

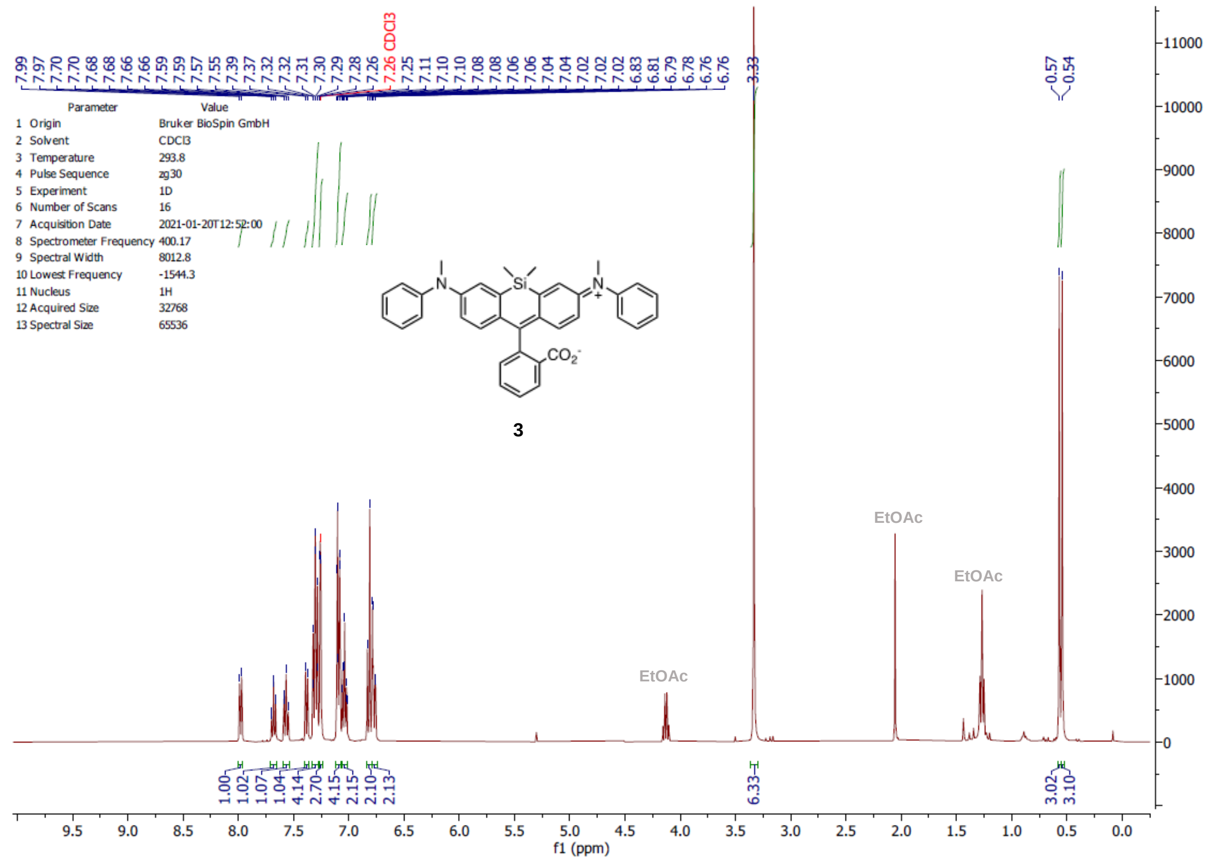

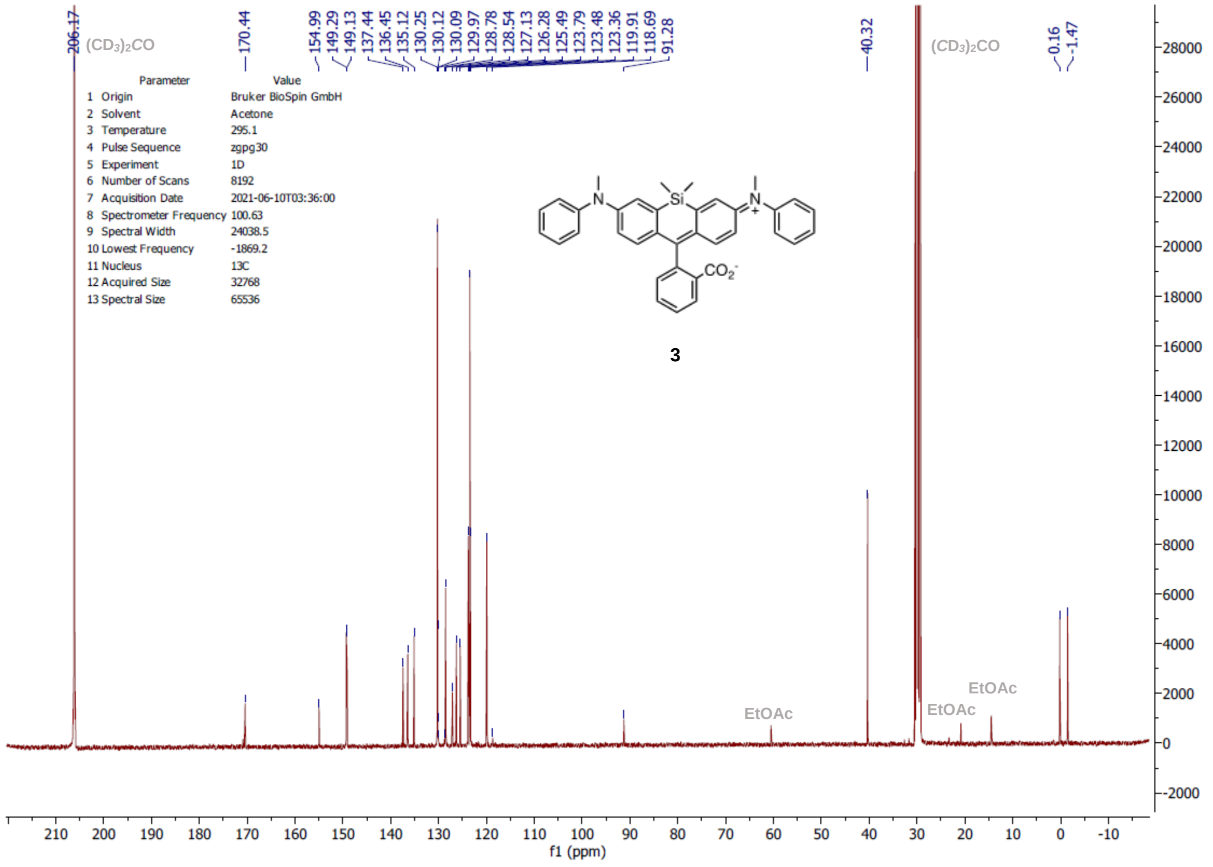

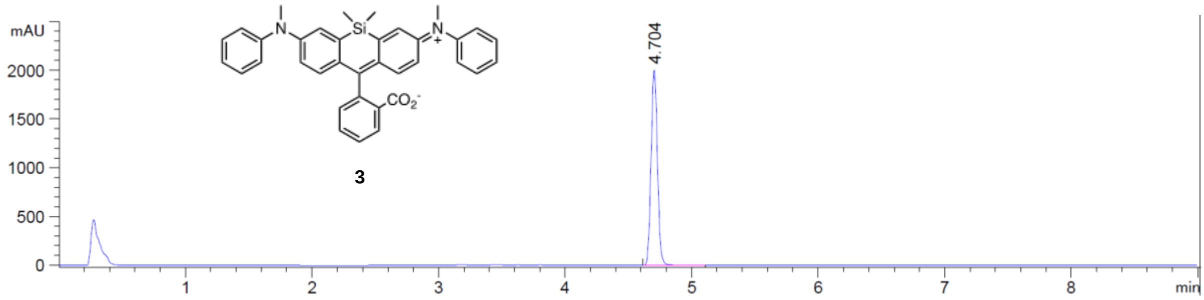
mAU
2000
1500
1000
500
0
1
2
3
4
5
6
7
8
min
4.704
Si
N
N
+
CO2-
3

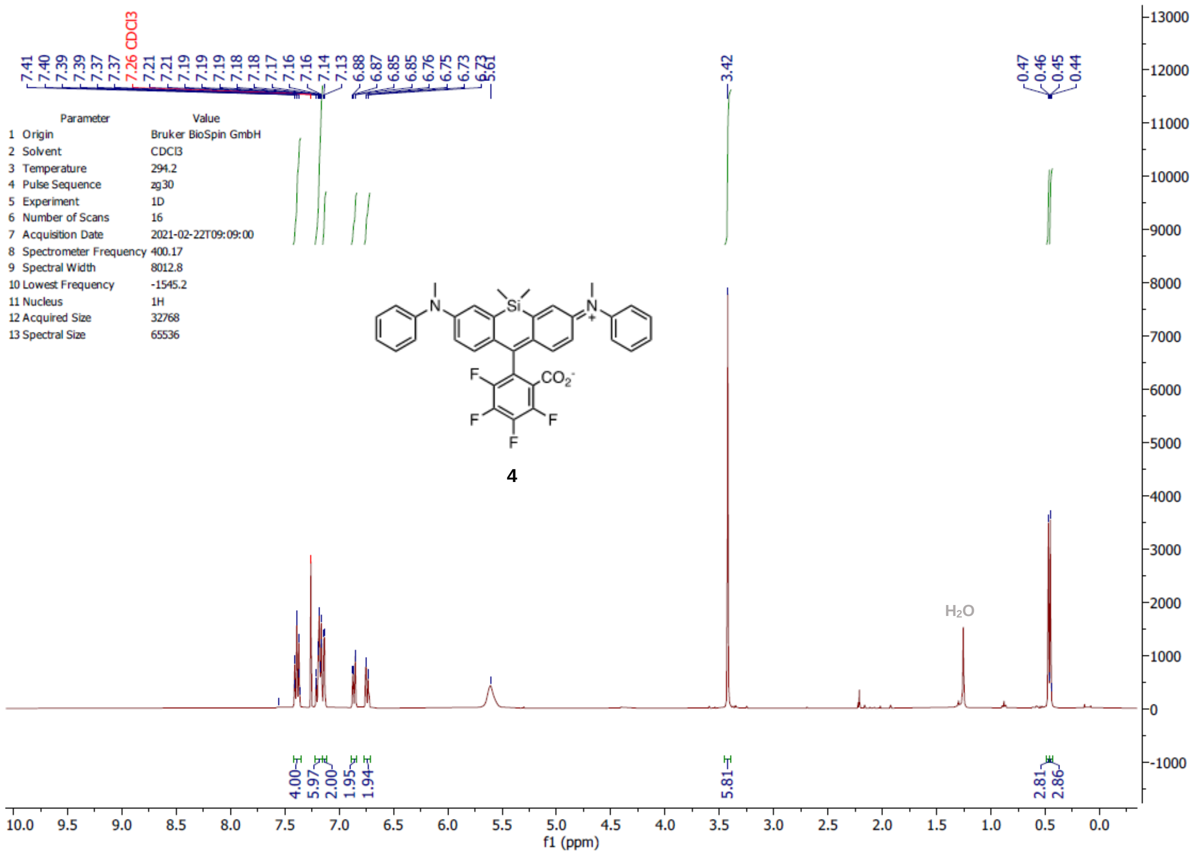
7.41 7.40 7.39 7.39 7.37 7.37 7.26 CDCl3 7.21 7.21 7.19 7.19 7.19 7.18 7.18 7.17 7.16 7.16 7.14 7.13 6.88 6.87 6.85 6.85 6.76 6.75 6.73 6.73 5.61
3.42
0.47 0.46 0.45 0.44
Parameter Value
1 Origin Bruker BioSpin GmbH
2 Solvent CDCl3
3 Temperature 294.2
4 Pulse Sequence zg30
5 Experiment 1D
6 Number of Scans 16
7 Acquisition Date 2021-02-22T09:09:00
8 Spectrometer Frequency 400.17
9 Spectral Width 8012.8
10 Lowest Frequency -1545.2
11 Nucleus 1H
12 Acquired Size 32768
13 Spectral Size 65536
4
H2O
4.00 5.97 2.00 1.95 1.94
5.81
2.81 2.86
f1 (ppm)

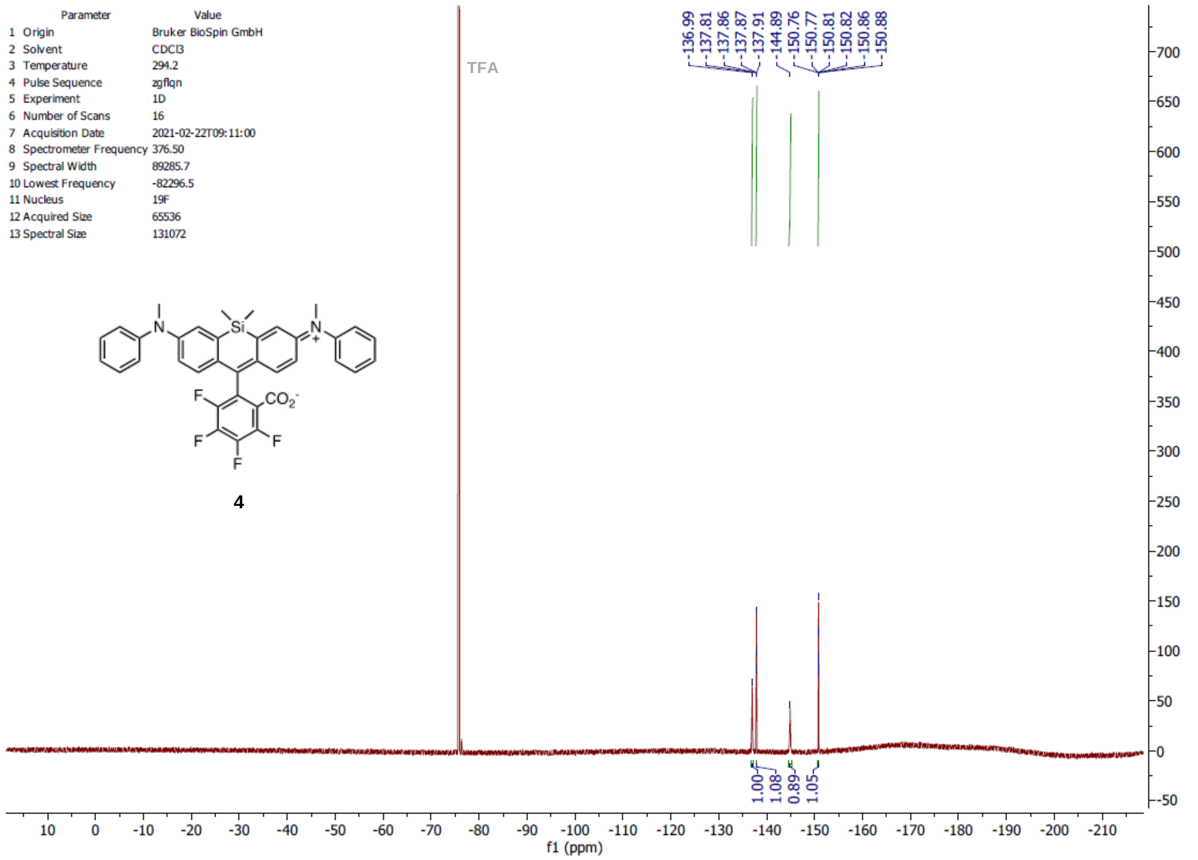
Parameter Value
1 Origin Bruker BioSpin GmbH
2 Solvent CDCl3
3 Temperature 294.2
4 Pulse Sequence zgflqn
5 Experiment 1D
6 Number of Scans 16
7 Acquisition Date 2021-02-22T09:11:00
8 Spectrometer Frequency 376.50
9 Spectral Width 89285.7
10 Lowest Frequency -82296.5
11 Nucleus 19F
12 Acquired Size 65536
13 Spectral Size 131072
TFA
-136.99 -137.81 -137.86 -137.87 -137.91 -144.89 -150.76 -150.77 -150.81 -150.82 -150.86 -150.88
4
1.00 1.08 0.89 1.05
f1 (ppm)

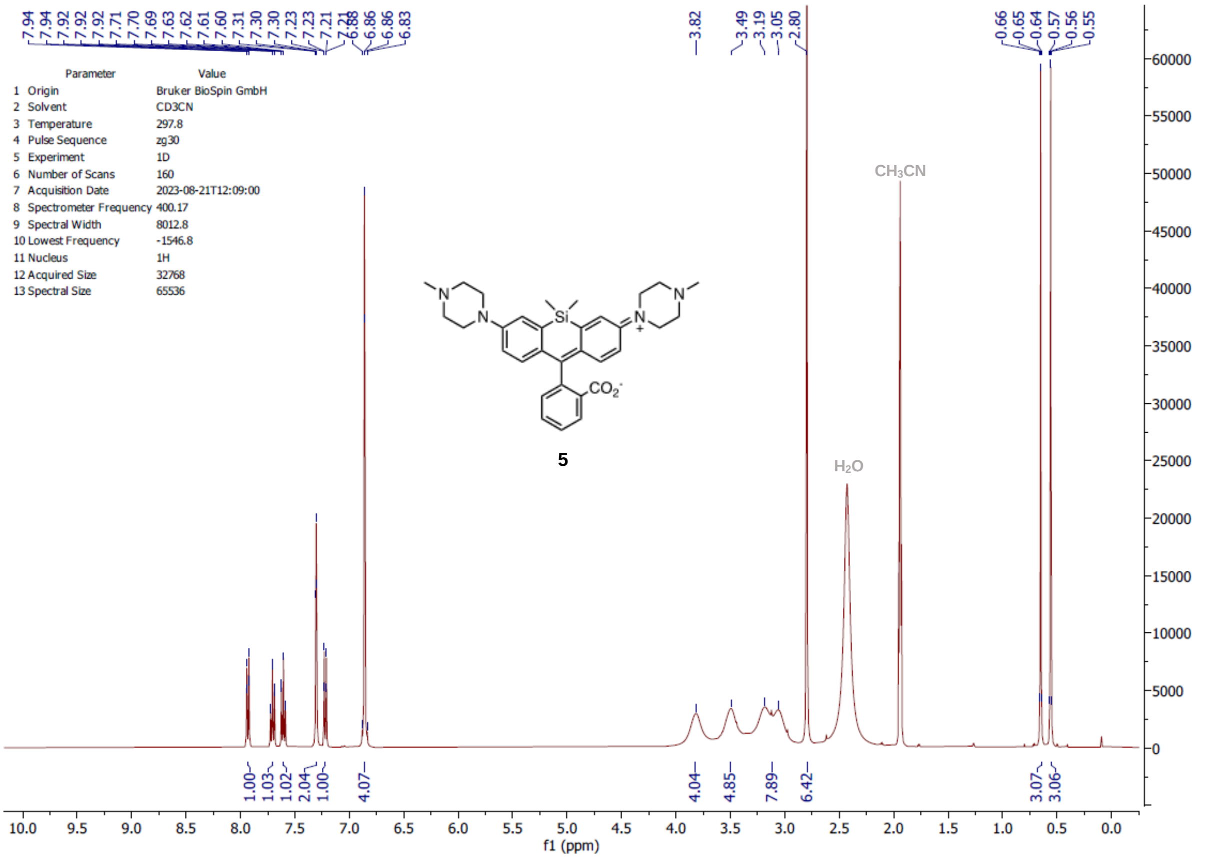
7.94 7.94 7.92 7.92 7.92 7.71 7.70 7.69 7.63 7.62 7.61 7.60 7.31 7.30 7.30 7.23 7.23 7.21 7.21 6.88 6.86 6.86 6.83
3.82
3.49 3.19 3.05 2.80
0.66 0.65 0.64 0.57 0.56 0.55
Parameter Value
1 Origin Bruker BioSpin GmbH
2 Solvent CD3CN
3 Temperature 297.8
4 Pulse Sequence zg30
5 Experiment 1D
6 Number of Scans 160
7 Acquisition Date 2023-08-21T12:09:00
8 Spectrometer Frequency 400.17
9 Spectral Width 8012.8
10 Lowest Frequency -1546.8
11 Nucleus 1H
12 Acquired Size 32768
13 Spectral Size 65536
Si
N
CO2-
5
CH3CN
H2O
1.00 1.03 1.02 2.04 1.00 4.07 4.04 4.85 7.89 6.42 3.07 3.06
f1 (ppm)

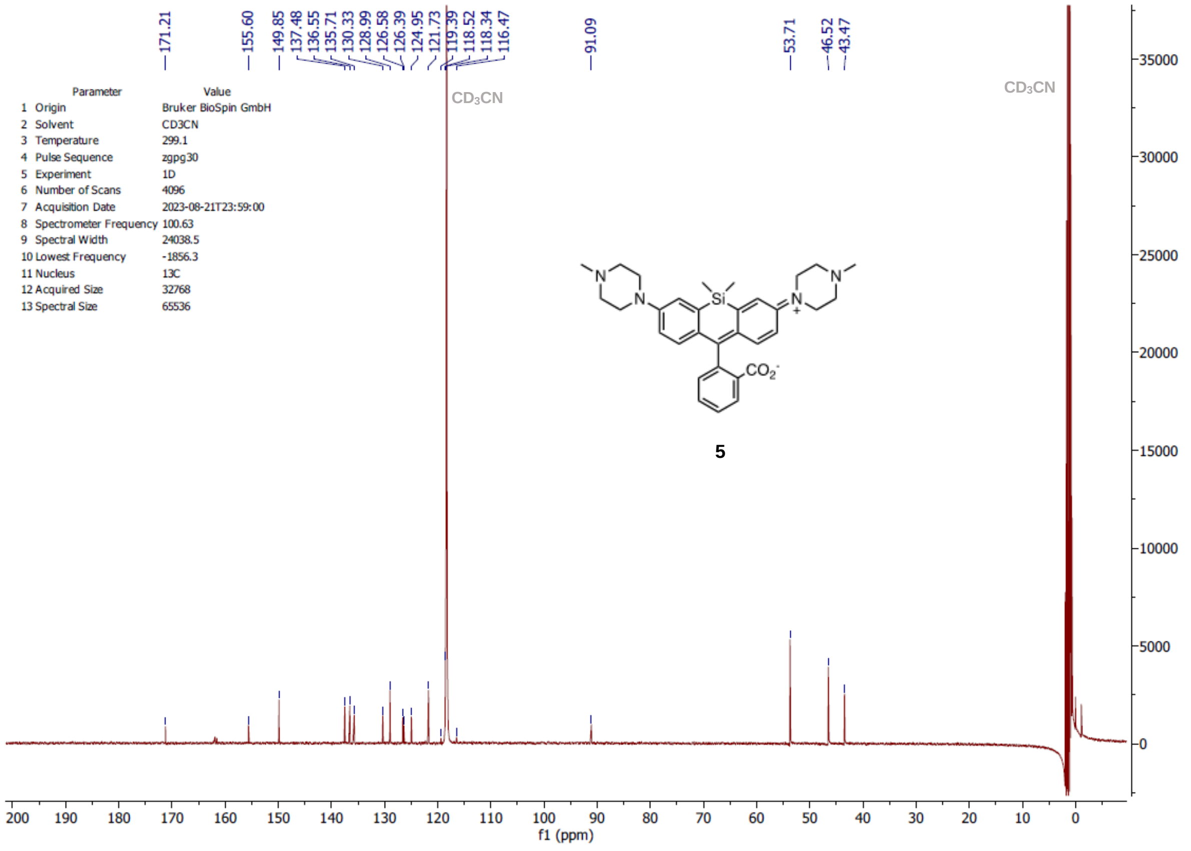
171.21
155.60
149.85 137.48 136.55 135.71 130.33 128.99 126.58 126.39 124.95 121.73 119.39 118.52 118.34 116.47
91.09
53.71
46.52 43.47
Parameter Value
1 Origin Bruker BioSpin GmbH
2 Solvent CD3CN
3 Temperature 299.1
4 Pulse Sequence zgpg30
5 Experiment 1D
6 Number of Scans 4096
7 Acquisition Date 2023-08-21T23:59:00
8 Spectrometer Frequency 100.63
9 Spectral Width 24038.5
10 Lowest Frequency -1856.3
11 Nucleus 13C
12 Acquired Size 32768
13 Spectral Size 65536
CD3CN
CD3CN
Si
N
CO2-
5
f1 (ppm)

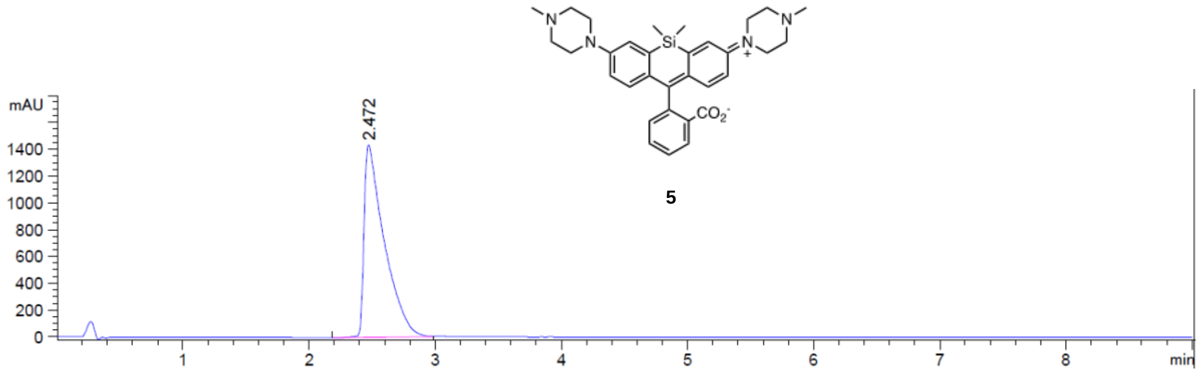

5

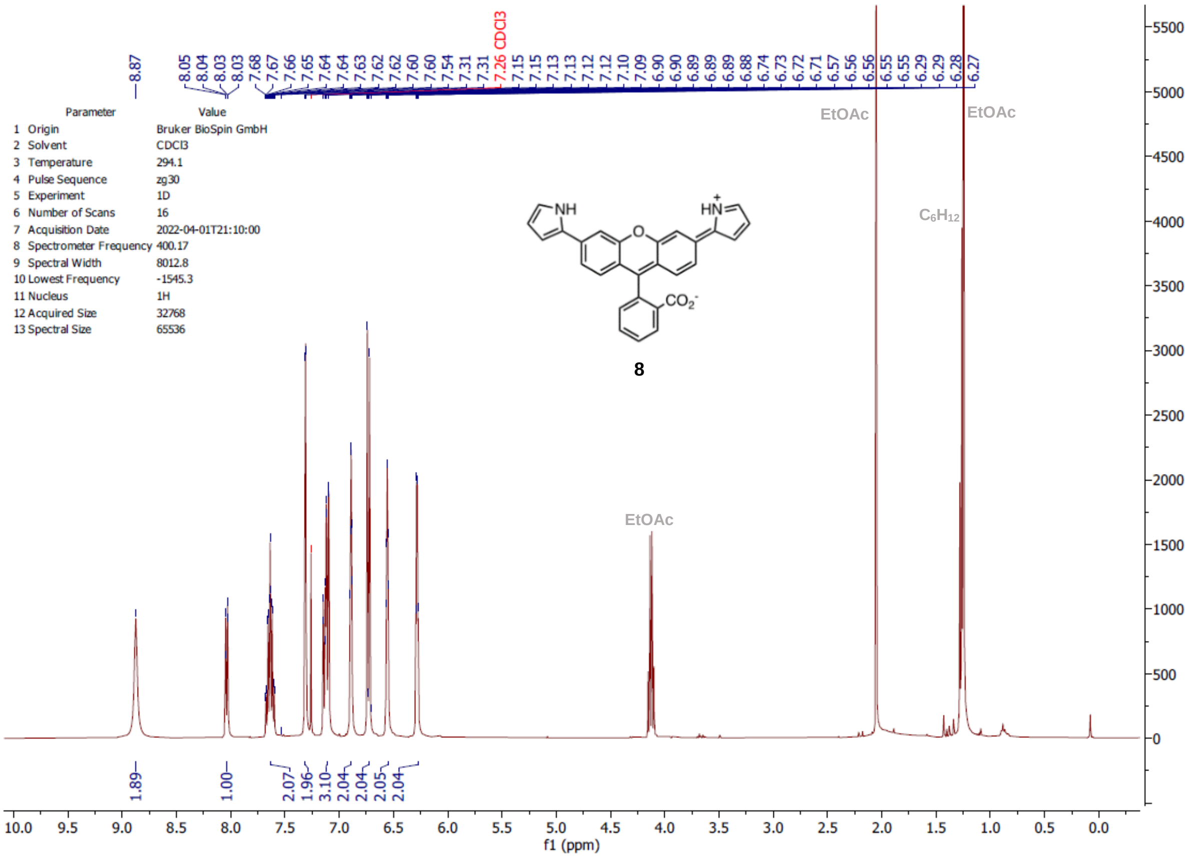
8.87
8.05 8.04 8.03 8.03 7.68 7.67 7.66 7.65 7.64 7.64 7.63 7.62 7.62 7.60 7.60 7.54 7.31 7.31 7.26 CDCl3 7.15 7.15 7.13 7.13 7.12 7.10 7.09 6.90 6.90 6.89 6.89 6.88 6.74 6.73 6.72 6.71 6.57 6.56 6.56 6.55 6.29 6.29 6.28 6.27
Parameter Value
1 Origin Bruker BioSpin GmbH
2 Solvent CDCl3
3 Temperature 294.1
4 Pulse Sequence zg30
5 Experiment 1D
6 Number of Scans 16
7 Acquisition Date 2022-04-01T21:10:00
8 Spectrometer Frequency 400.17
9 Spectral Width 8012.8
10 Lowest Frequency -1545.3
11 Nucleus 1H
12 Acquired Size 32768
13 Spectral Size 65536
EtOAc
EtOAc
EtOAc
$C_6H_{12}$
NH
HN
$CO_2^-$
8
1.89
1.00
2.07
1.96
3.10
2.04
2.04
2.05
2.04
f1 (ppm)

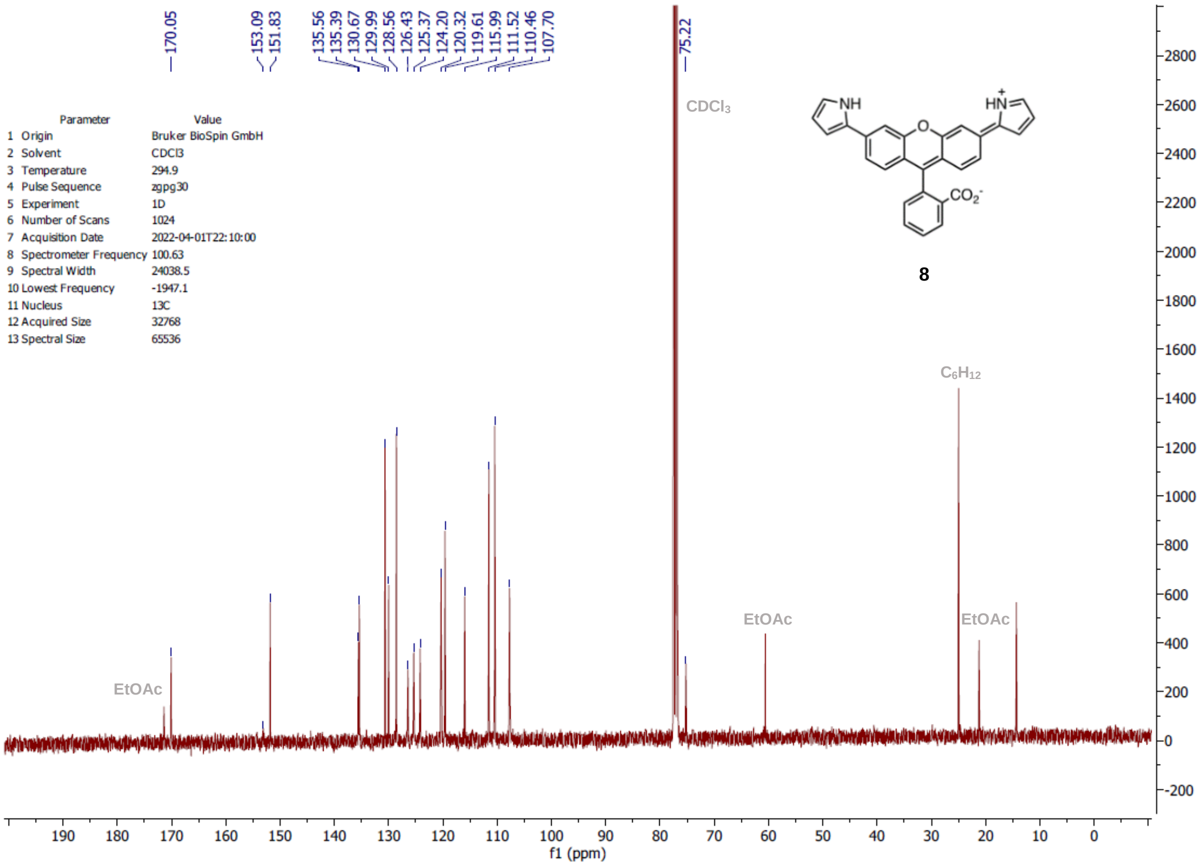
170.05
153.09 151.83
135.56 135.39 130.67 129.99 128.56 126.43 125.37 124.20 120.32 119.61 115.99 111.52 110.46 107.70
75.22
Parameter Value
1 Origin Bruker BioSpin GmbH
2 Solvent CDCl3
3 Temperature 294.9
4 Pulse Sequence zgpg30
5 Experiment 1D
6 Number of Scans 1024
7 Acquisition Date 2022-04-01T22:10:00
8 Spectrometer Frequency 100.63
9 Spectral Width 24038.5
10 Lowest Frequency -1947.1
11 Nucleus 13C
12 Acquired Size 32768
13 Spectral Size 65536
$CDCl_3$
NH
HN
$CO_2^-$
8
$C_6H_{12}$
EtOAc
EtOAc
EtOAc
f1 (ppm)

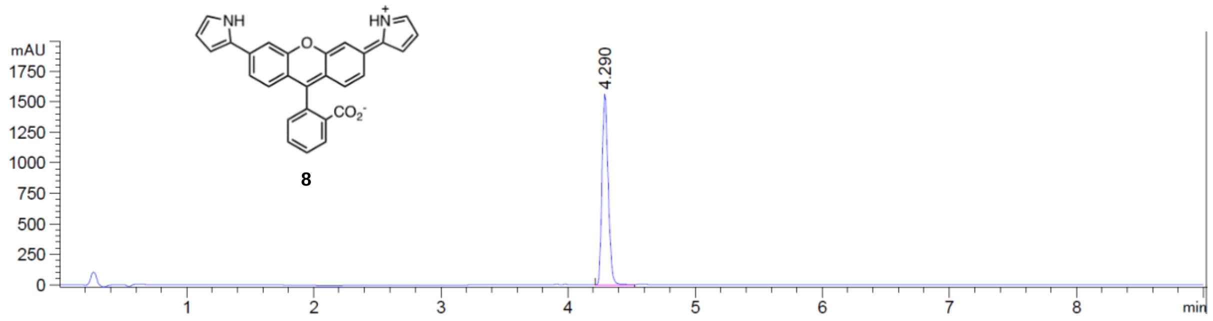
NH
HN+
CO2-
8
mAU
1750
1500
1250
1000
750
500
250
0
4.290
1
2
3
4
5
6
7
8
min

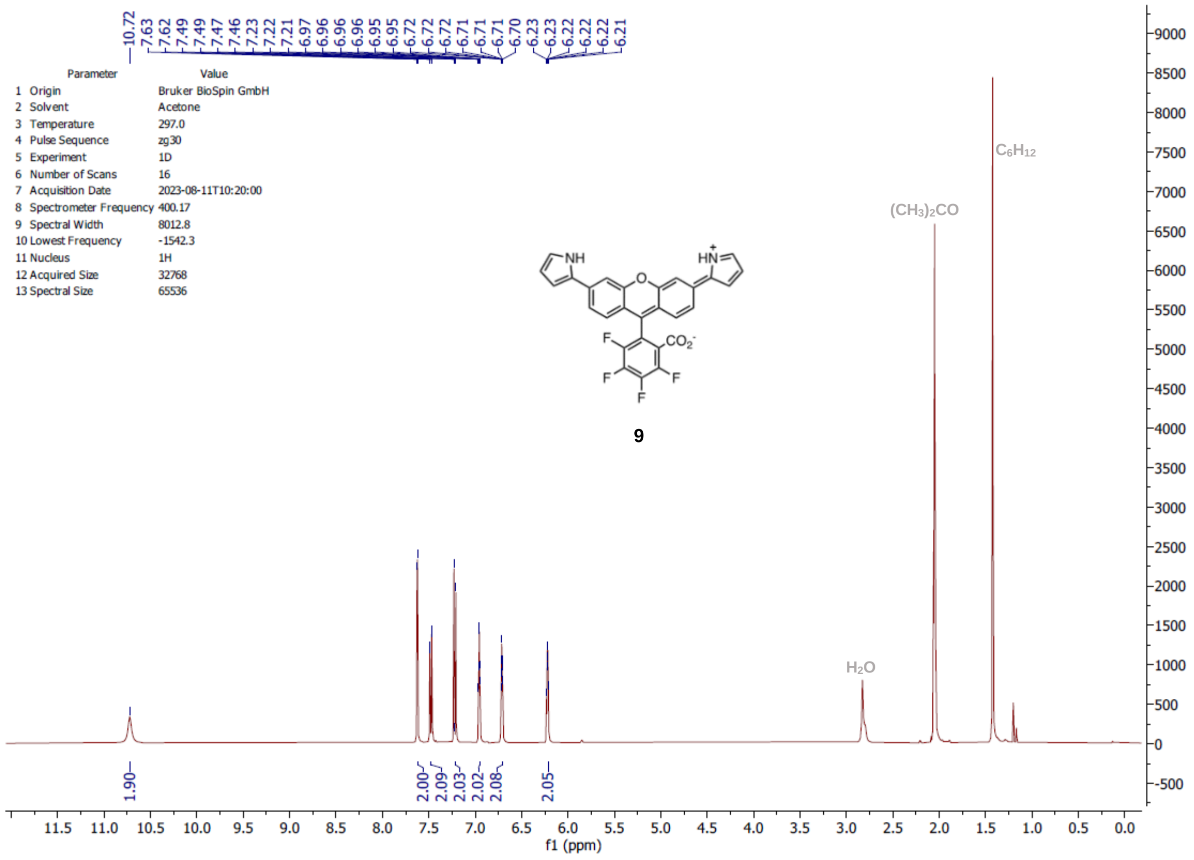

10.72
7.63
7.62
7.49
7.47
7.46
7.23
7.22
7.21
6.97
6.96
6.96
6.96
6.95
6.95
6.72
6.72
6.72
6.71
6.71
6.71
6.70
6.23
6.22
6.22
6.22
6.21
Parameter Value
1 Origin Bruker BioSpin GmbH
2 Solvent Acetone
3 Temperature 297.0
4 Pulse Sequence zg30
5 Experiment 1D
6 Number of Scans 16
7 Acquisition Date 2023-08-11T10:20:00
8 Spectrometer Frequency 400.17
9 Spectral Width 8012.8
10 Lowest Frequency -1542.3
11 Nucleus 1H
12 Acquired Size 32768
13 Spectral Size 65536
NH
HN+
O
F
CO2-
F
F
F
9
C6H12
(CH3)2CO
H2O
1.90
2.00
2.09
2.03
2.02
2.08
2.05
f1 (ppm)

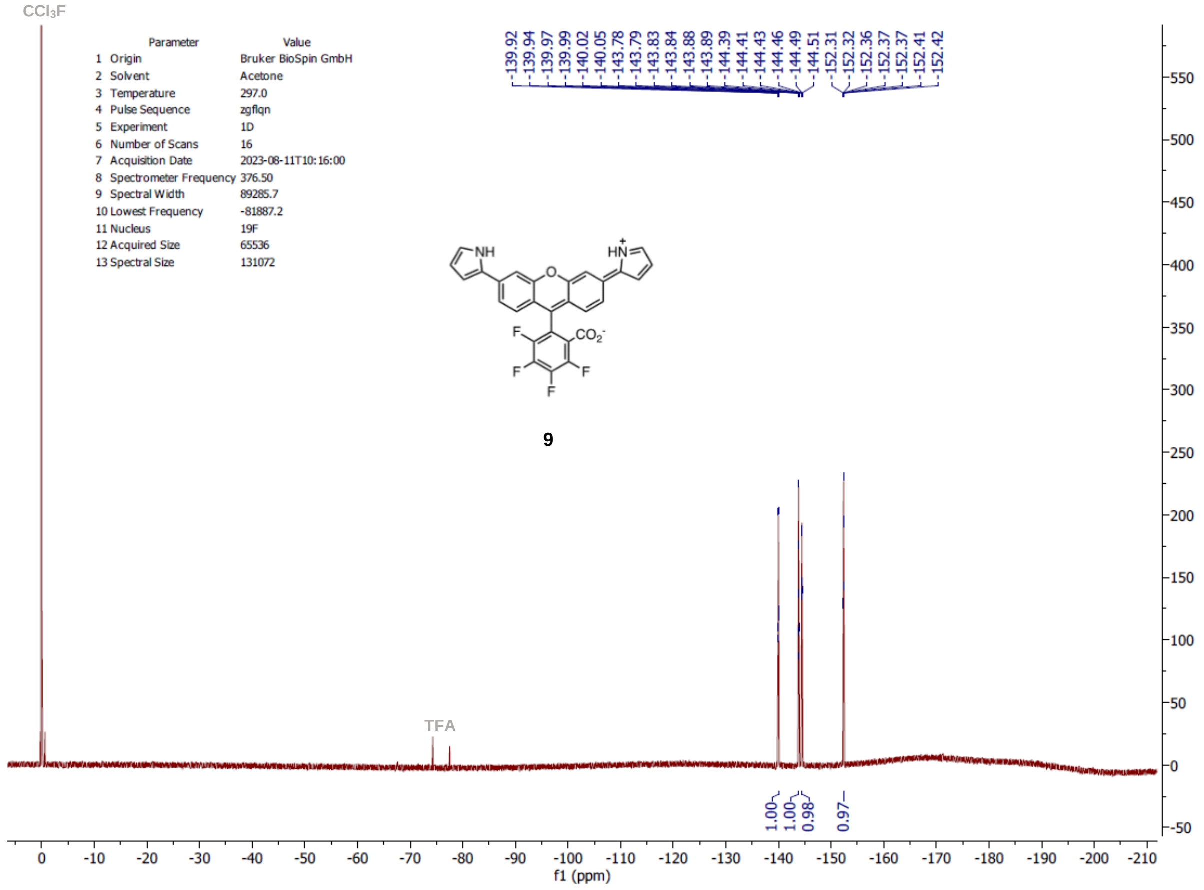

CCl3F
Parameter Value
1 Origin Bruker BioSpin GmbH
2 Solvent Acetone
3 Temperature 297.0
4 Pulse Sequence zgflqn
5 Experiment 1D
6 Number of Scans 16
7 Acquisition Date 2023-08-11T10:16:00
8 Spectrometer Frequency 376.50
9 Spectral Width 89285.7
10 Lowest Frequency -81887.2
11 Nucleus 19F
12 Acquired Size 65536
13 Spectral Size 131072
-139.92
-139.94
-139.97
-139.99
-140.02
-140.05
-143.78
-143.79
-143.83
-143.84
-143.88
-143.89
-144.39
-144.41
-144.43
-144.46
-144.49
-144.51
-152.31
-152.32
-152.36
-152.37
-152.37
-152.41
-152.42
NH
HN+
O
F
CO2-
F
F
F
9
TFA
1.00
1.00
0.98
0.97
f1 (ppm)

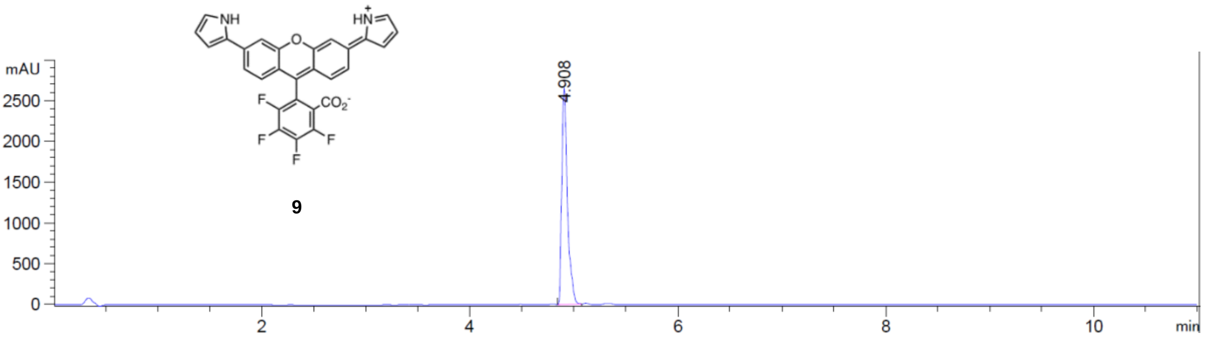

NH
HN+
O
F
CO2-
F
F
F
9
mAU
2500
2000
1500
1000
500
0
4.908
2
4
6
8
10
min

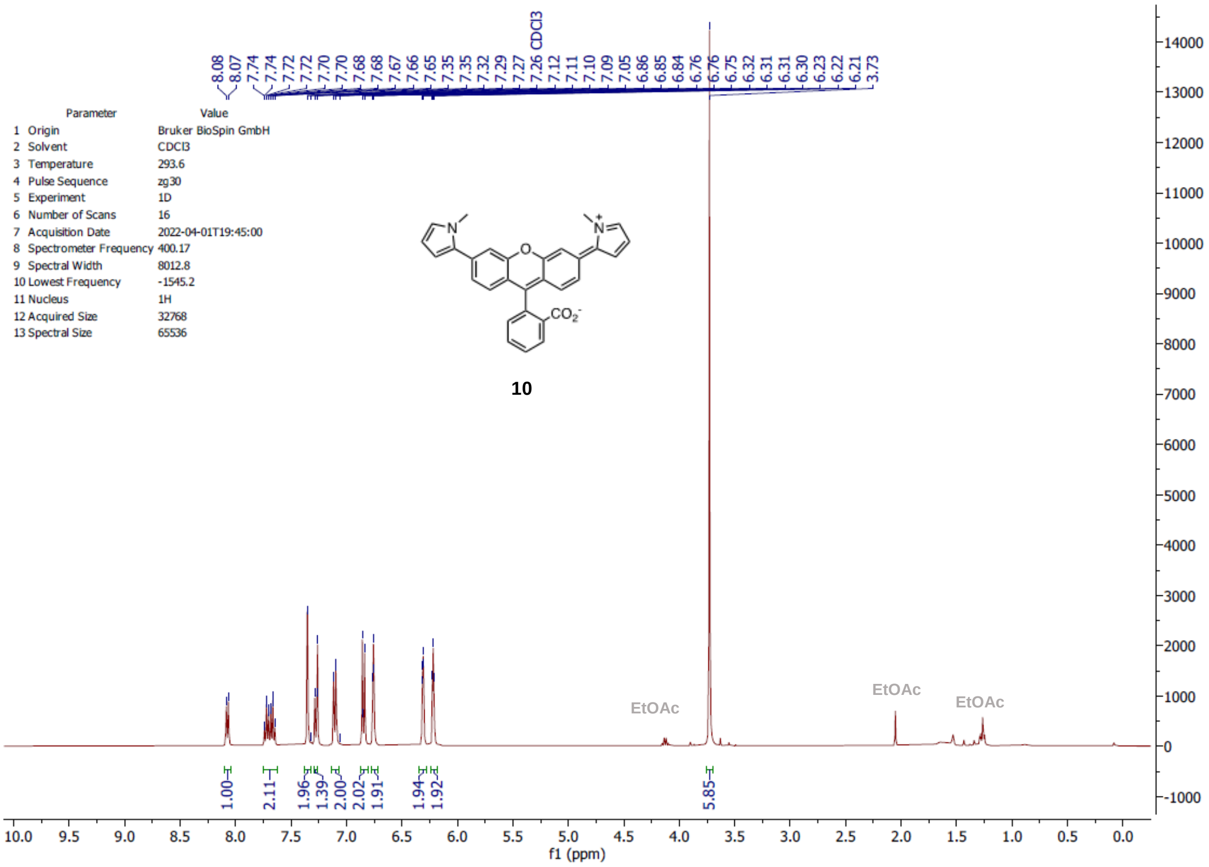

| Parameter | Value |
|---|---|
| 1 Origin | Bruker BioSpin GmbH |
| 2 Solvent | CDCl3 |
| 3 Temperature | 293.6 |
| 4 Pulse Sequence | zg30 |
| 5 Experiment | 1D |
| 6 Number of Scans | 16 |
| 7 Acquisition Date | 2022-04-01T19:45:00 |
| 8 Spectrometer Frequency | 400.17 |
| 9 Spectral Width | 8012.8 |
| 10 Lowest Frequency | -1545.2 |
| 11 Nucleus | 1H |
| 12 Acquired Size | 32768 |
| 13 Spectral Size | 65536 |

**10**

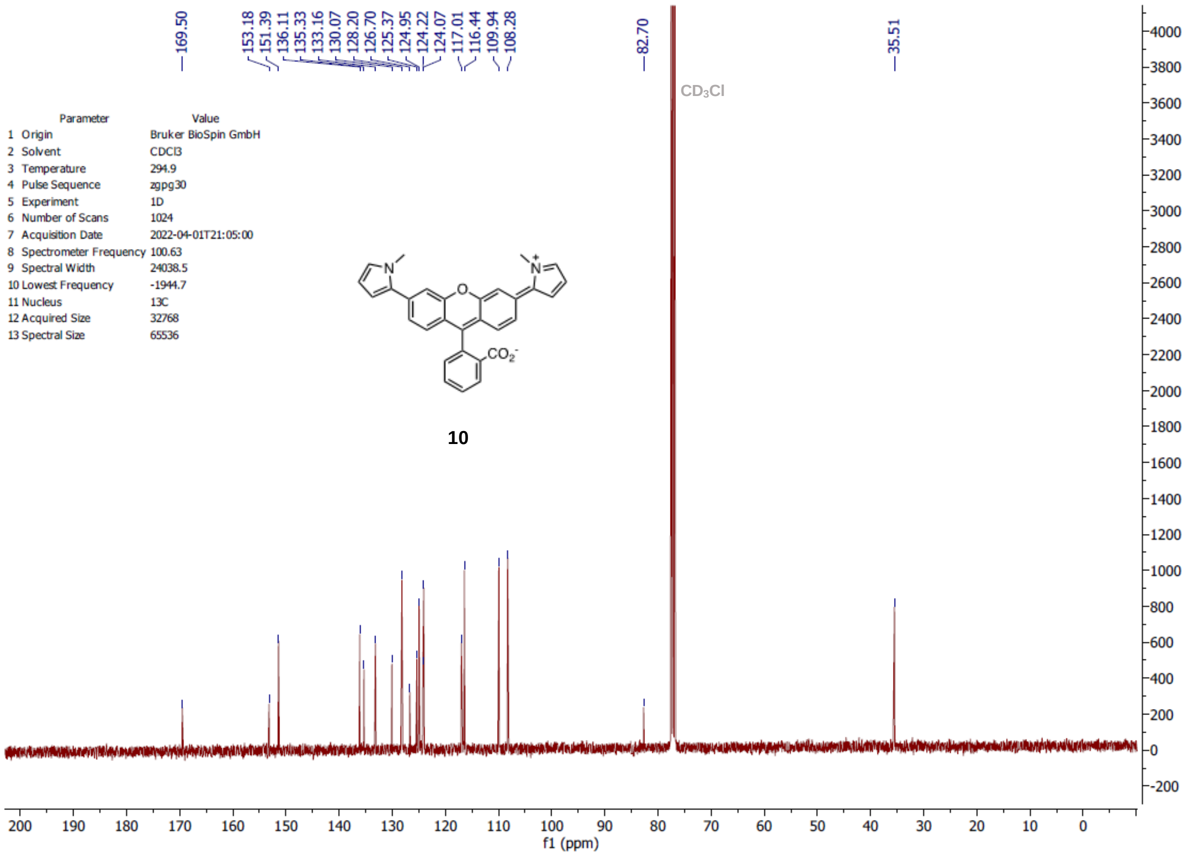

| Parameter | Value |
|---|---|
| 1 Origin | Bruker BioSpin GmbH |
| 2 Solvent | CDCl3 |
| 3 Temperature | 294.9 |
| 4 Pulse Sequence | zgpg30 |
| 5 Experiment | 1D |
| 6 Number of Scans | 1024 |
| 7 Acquisition Date | 2022-04-01T21:05:00 |
| 8 Spectrometer Frequency | 100.63 |
| 9 Spectral Width | 24038.5 |
| 10 Lowest Frequency | -1944.7 |
| 11 Nucleus | 13C |
| 12 Acquired Size | 32768 |
| 13 Spectral Size | 65536 |

**10**

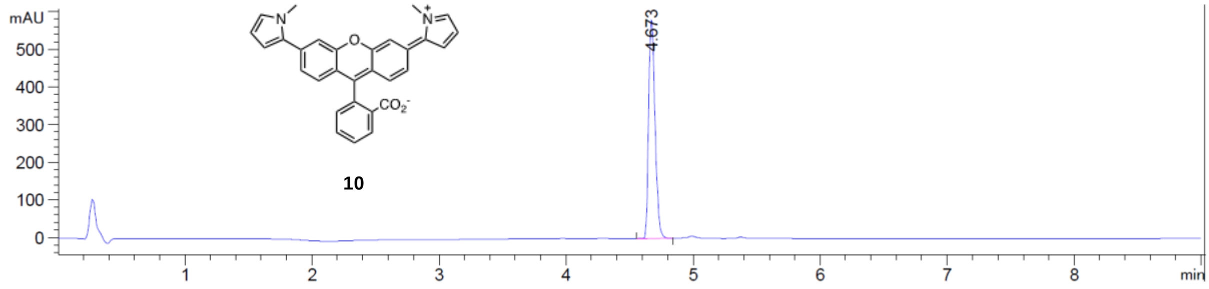
mAU
500
400
300
200
100
0
1
2
3
4
5
6
7
8
min
4.673
N
N+
O
$CO_2^-$
10

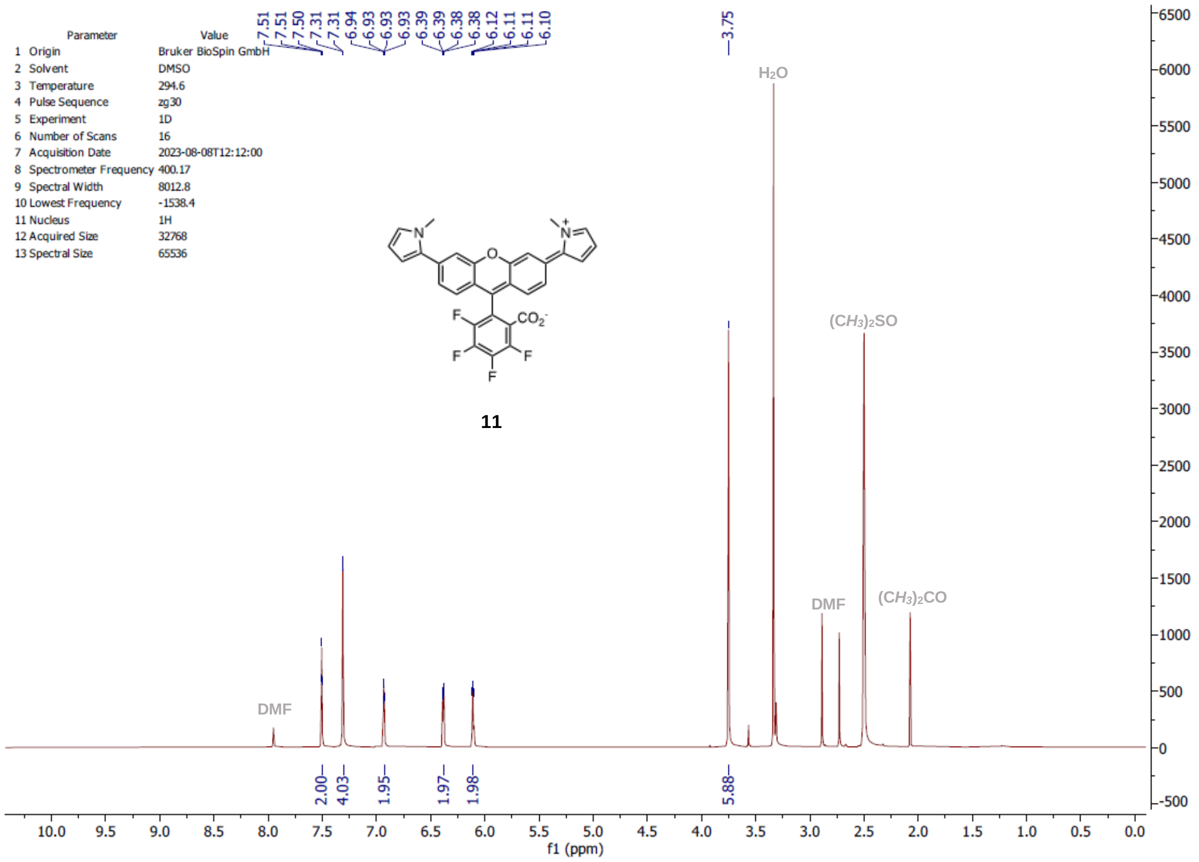
Parameter Value
1 Origin Bruker BioSpin GmbH
2 Solvent DMSO
3 Temperature 294.6
4 Pulse Sequence zg30
5 Experiment 1D
6 Number of Scans 16
7 Acquisition Date 2023-08-08T12:12:00
8 Spectrometer Frequency 400.17
9 Spectral Width 8012.8
10 Lowest Frequency -1538.4
11 Nucleus 1H
12 Acquired Size 32768
13 Spectral Size 65536
7.51 7.51 7.50 7.31 7.31 6.94 6.93 6.93 6.93 6.39 6.39 6.38 6.38 6.12 6.11 6.11 6.10
3.75
H2O
(CH3)2SO
DMF
(CH3)2CO
DMF
11
2.00 4.03 1.95 1.97 1.98 5.88
f1 (ppm)

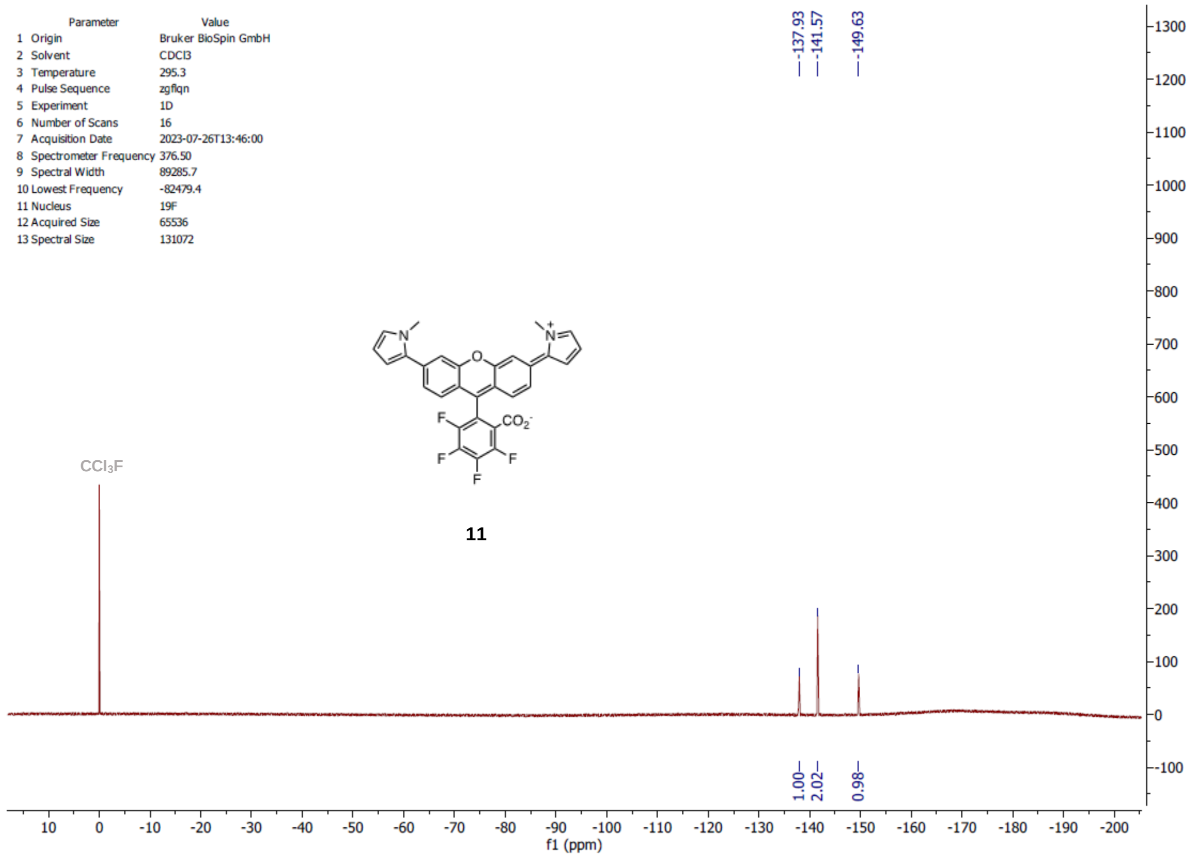
Parameter Value
1 Origin Bruker BioSpin GmbH
2 Solvent CDCl3
3 Temperature 295.3
4 Pulse Sequence zgflqn
5 Experiment 1D
6 Number of Scans 16
7 Acquisition Date 2023-07-26T13:46:00
8 Spectrometer Frequency 376.50
9 Spectral Width 89285.7
10 Lowest Frequency -82479.4
11 Nucleus 19F
12 Acquired Size 65536
13 Spectral Size 131072
-137.93 -141.57 -149.63
CCl3F
11
1.00 2.02 0.98
f1 (ppm)

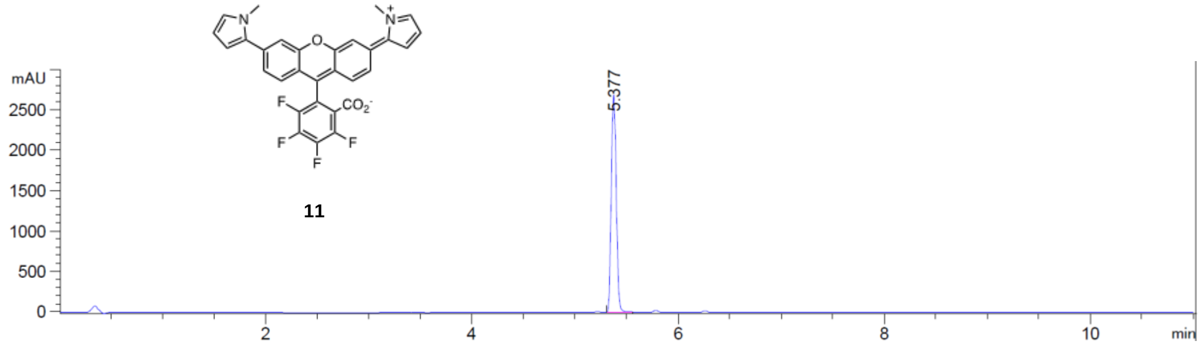
N
N+
O
F
CO2-
F
F
F
F
11
mAU
2500
2000
1500
1000
500
0
5.377
2
4
6
8
10
min

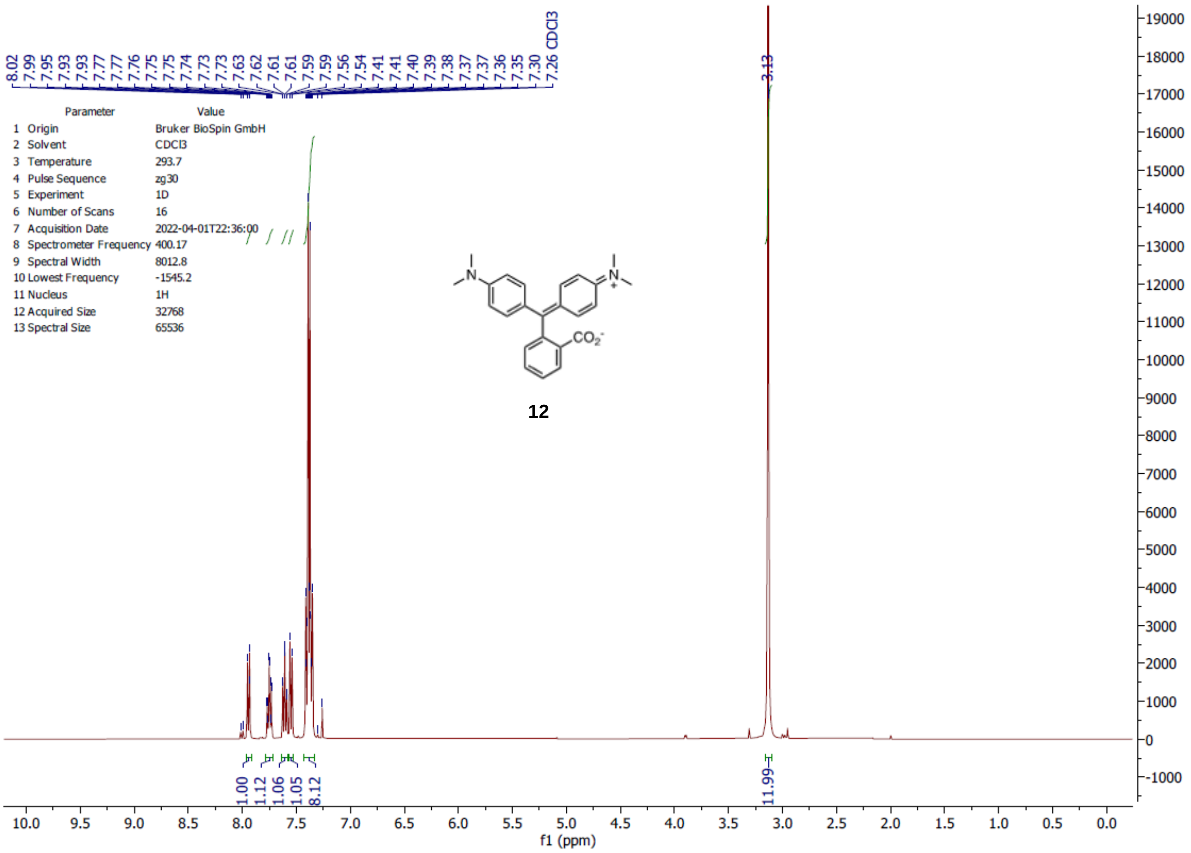

8.02 7.99 7.95 7.93 7.93 7.77 7.77 7.76 7.75 7.75 7.74 7.73 7.73 7.63 7.62 7.61 7.61 7.59 7.59 7.56 7.54 7.41 7.41 7.40 7.39 7.38 7.37 7.37 7.36 7.35 7.30 7.26 CDCl3
3.13
Parameter Value
1 Origin Bruker BioSpin GmbH
2 Solvent CDCl3
3 Temperature 293.7
4 Pulse Sequence zg30
5 Experiment 1D
6 Number of Scans 16
7 Acquisition Date 2022-04-01T22:36:00
8 Spectrometer Frequency 400.17
9 Spectral Width 8012.8
10 Lowest Frequency -1545.2
11 Nucleus 1H
12 Acquired Size 32768
13 Spectral Size 65536
12
1.00 1.12 1.06 1.05 8.12
11.99
f1 (ppm)

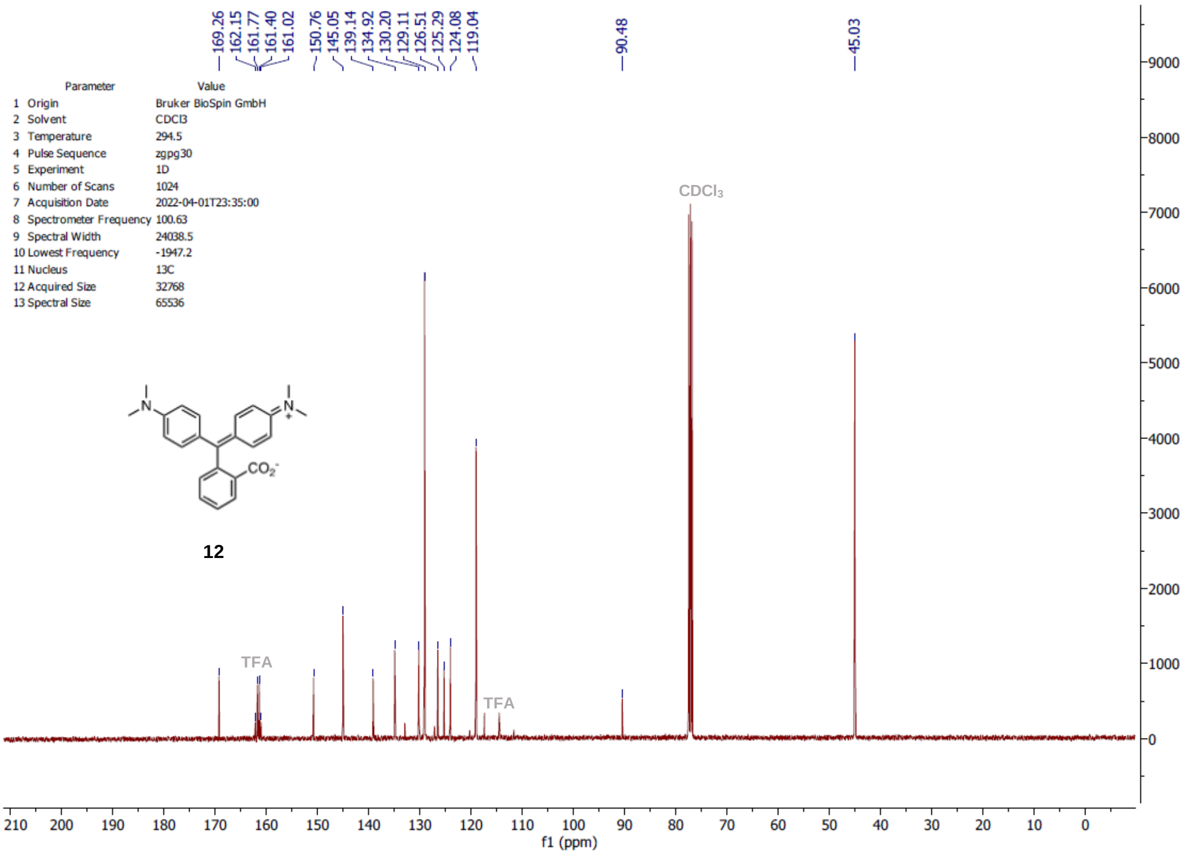

169.26 162.15 161.77 161.40 161.02 150.76 145.05 139.14 134.92 130.20 129.11 126.51 125.29 124.08 119.04
90.48
45.03
Parameter Value
1 Origin Bruker BioSpin GmbH
2 Solvent CDCl3
3 Temperature 294.5
4 Pulse Sequence zgpg30
5 Experiment 1D
6 Number of Scans 1024
7 Acquisition Date 2022-04-01T23:35:00
8 Spectrometer Frequency 100.63
9 Spectral Width 24038.5
10 Lowest Frequency -1947.2
11 Nucleus 13C
12 Acquired Size 32768
13 Spectral Size 65536
CDCl3
TFA
TFA
12
f1 (ppm)

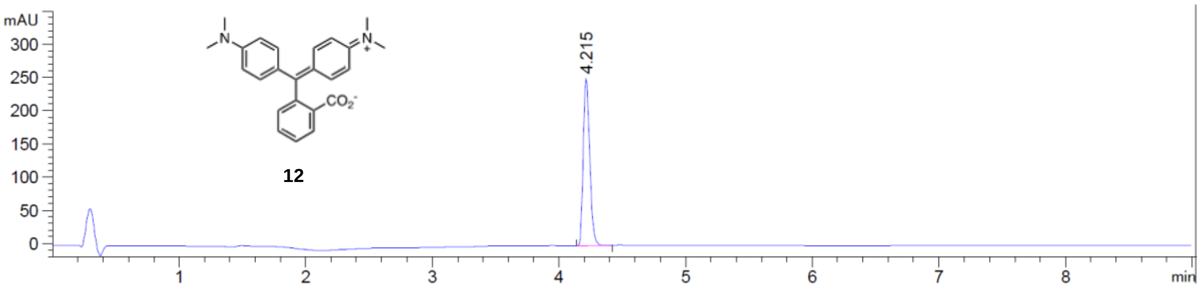
mAU
300
250
200
150
100
50
0
4.215
CO2-
12
1
2
3
4
5
6
7
8
min

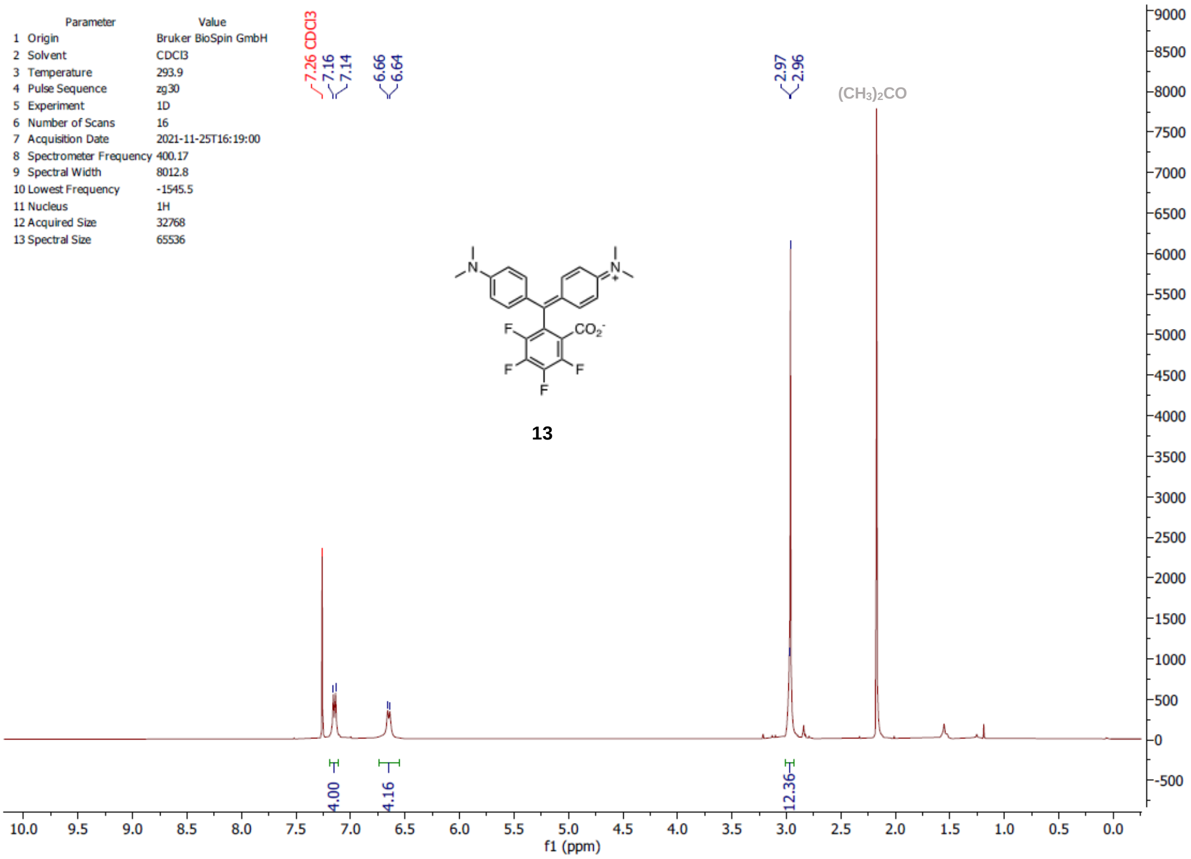
Parameter Value
1 Origin Bruker BioSpin GmbH
2 Solvent CDCl3
3 Temperature 293.9
4 Pulse Sequence zg30
5 Experiment 1D
6 Number of Scans 16
7 Acquisition Date 2021-11-25T16:19:00
8 Spectrometer Frequency 400.17
9 Spectral Width 8012.8
10 Lowest Frequency -1545.5
11 Nucleus 1H
12 Acquired Size 32768
13 Spectral Size 65536
7.26 CDCl3
7.16
7.14
6.66
6.64
2.97
2.96
(CH3)2CO
13
4.00
4.16
12.36
f1 (ppm)

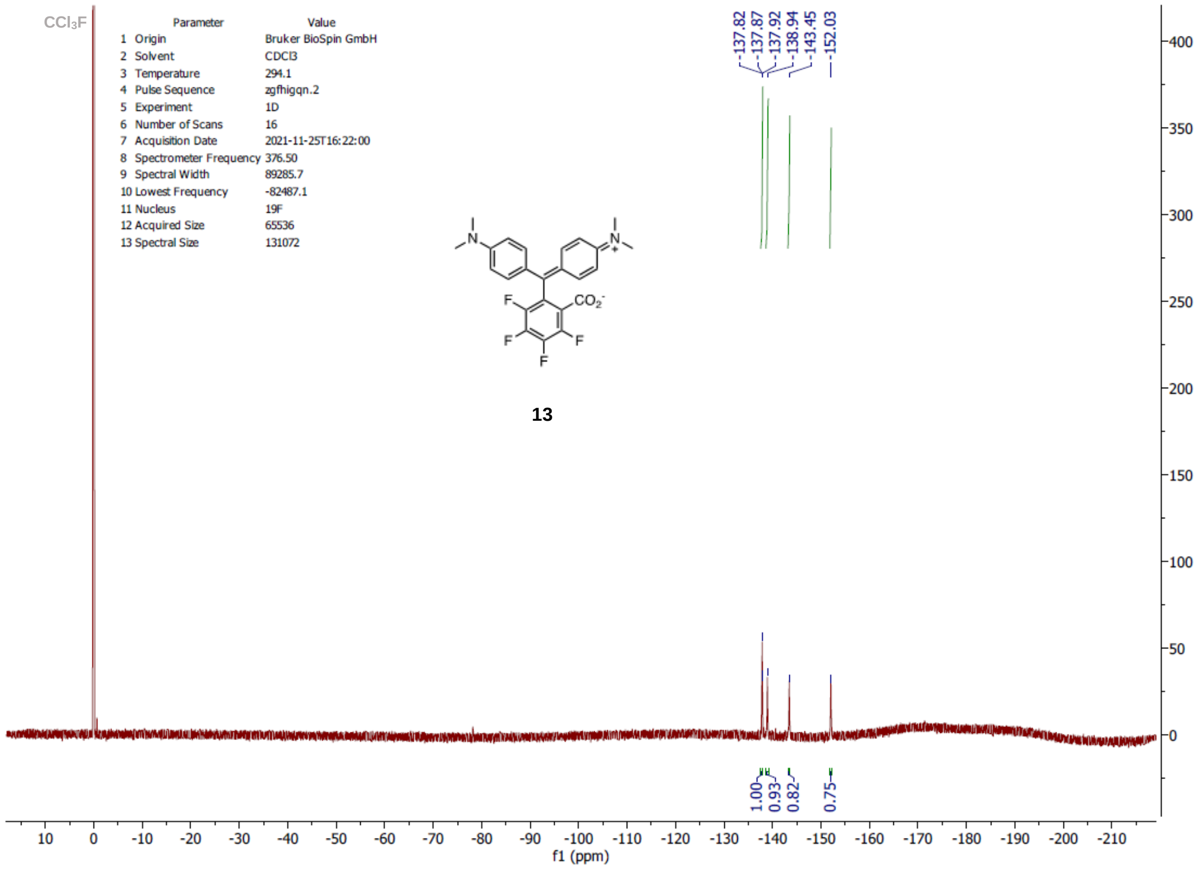
CCl3F
Parameter Value
1 Origin Bruker BioSpin GmbH
2 Solvent CDCl3
3 Temperature 294.1
4 Pulse Sequence zgfhigqn.2
5 Experiment 1D
6 Number of Scans 16
7 Acquisition Date 2021-11-25T16:22:00
8 Spectrometer Frequency 376.50
9 Spectral Width 89285.7
10 Lowest Frequency -82487.1
11 Nucleus 19F
12 Acquired Size 65536
13 Spectral Size 131072
-137.82
-137.87
-137.92
-138.94
-143.45
-152.03
13
1.00
0.93
0.82
0.75
f1 (ppm)

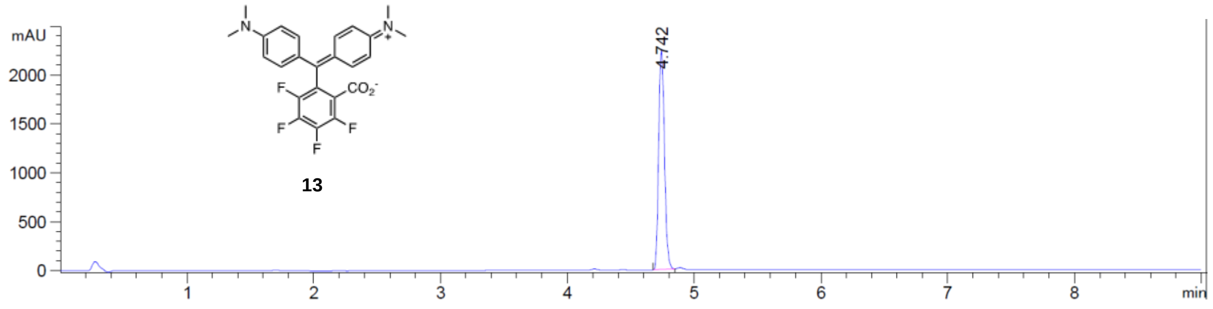
mAU
2000
1500
1000
500
0
4.742
13
1
2
3
4
5
6
7
8
min

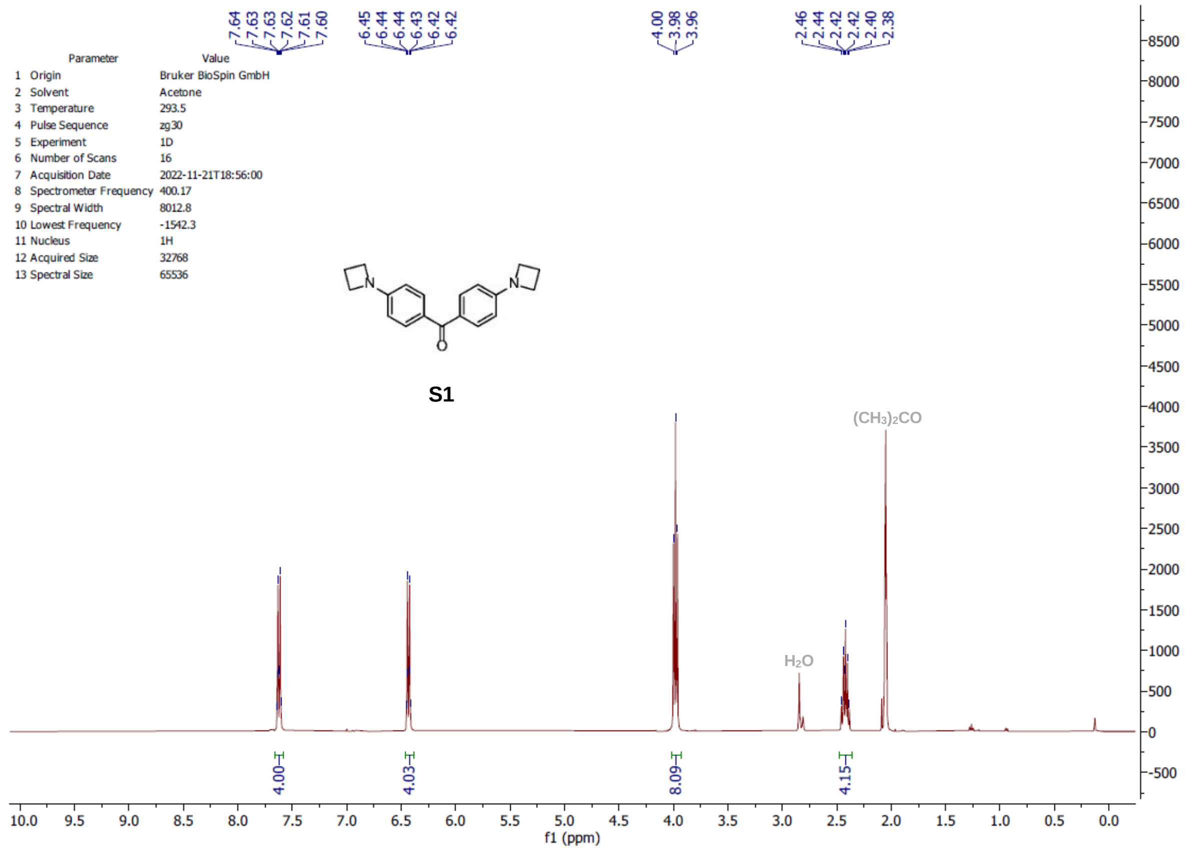
7.64
7.63
7.63
7.62
7.61
7.60
6.45
6.44
6.44
6.43
6.42
6.42
4.00
3.98
3.96
2.46
2.44
2.42
2.42
2.40
2.38
Parameter Value
1 Origin Bruker BioSpin GmbH
2 Solvent Acetone
3 Temperature 293.5
4 Pulse Sequence zg30
5 Experiment 1D
6 Number of Scans 16
7 Acquisition Date 2022-11-21T18:56:00
8 Spectrometer Frequency 400.17
9 Spectral Width 8012.8
10 Lowest Frequency -1542.3
11 Nucleus 1H
12 Acquired Size 32768
13 Spectral Size 65536
S1
(CH3)2CO
H2O
4.00
4.03
8.09
4.15
f1 (ppm)

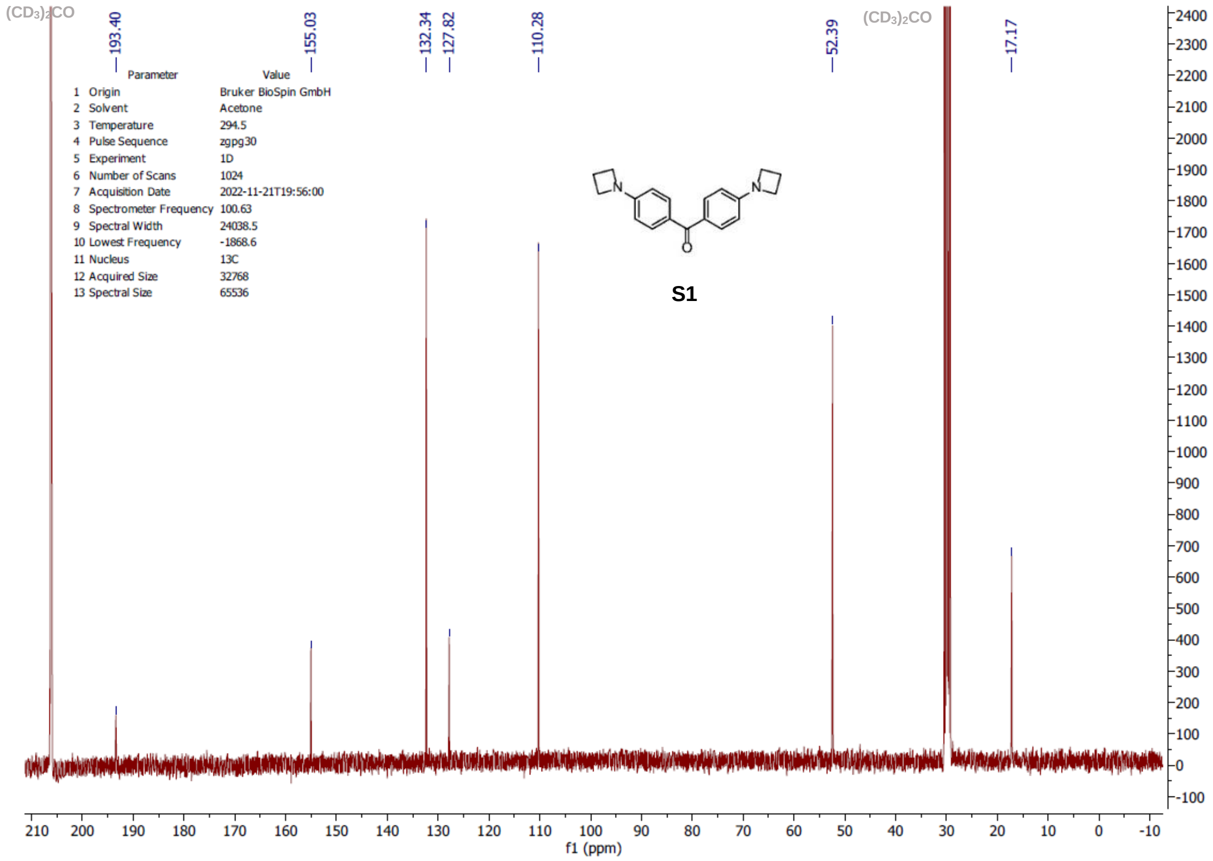
(CD3)2CO
193.40
155.03
132.34
127.82
110.28
52.39
(CD3)2CO
17.17
Parameter Value
1 Origin Bruker BioSpin GmbH
2 Solvent Acetone
3 Temperature 294.5
4 Pulse Sequence zgpg30
5 Experiment 1D
6 Number of Scans 1024
7 Acquisition Date 2022-11-21T19:56:00
8 Spectrometer Frequency 100.63
9 Spectral Width 24038.5
10 Lowest Frequency -1868.6
11 Nucleus 13C
12 Acquired Size 32768
13 Spectral Size 65536
S1
f1 (ppm)

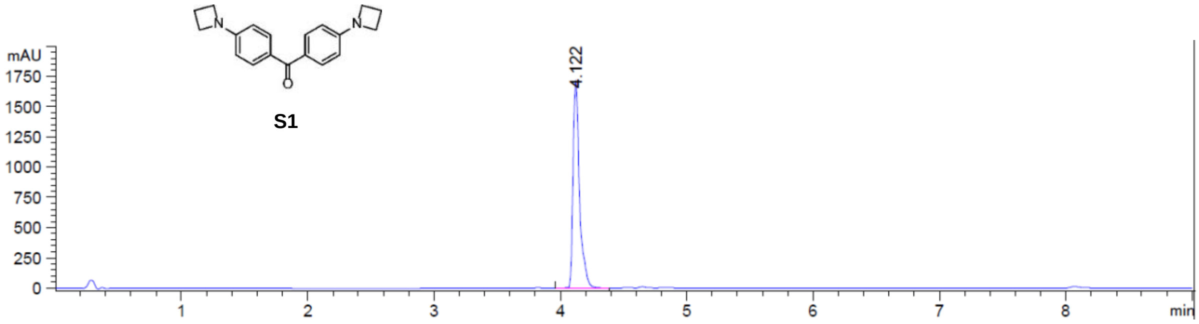
S1
N
O
mAU
1750
1500
1250
1000
750
500
250
0
4.122
1
2
3
4
5
6
7
8
min

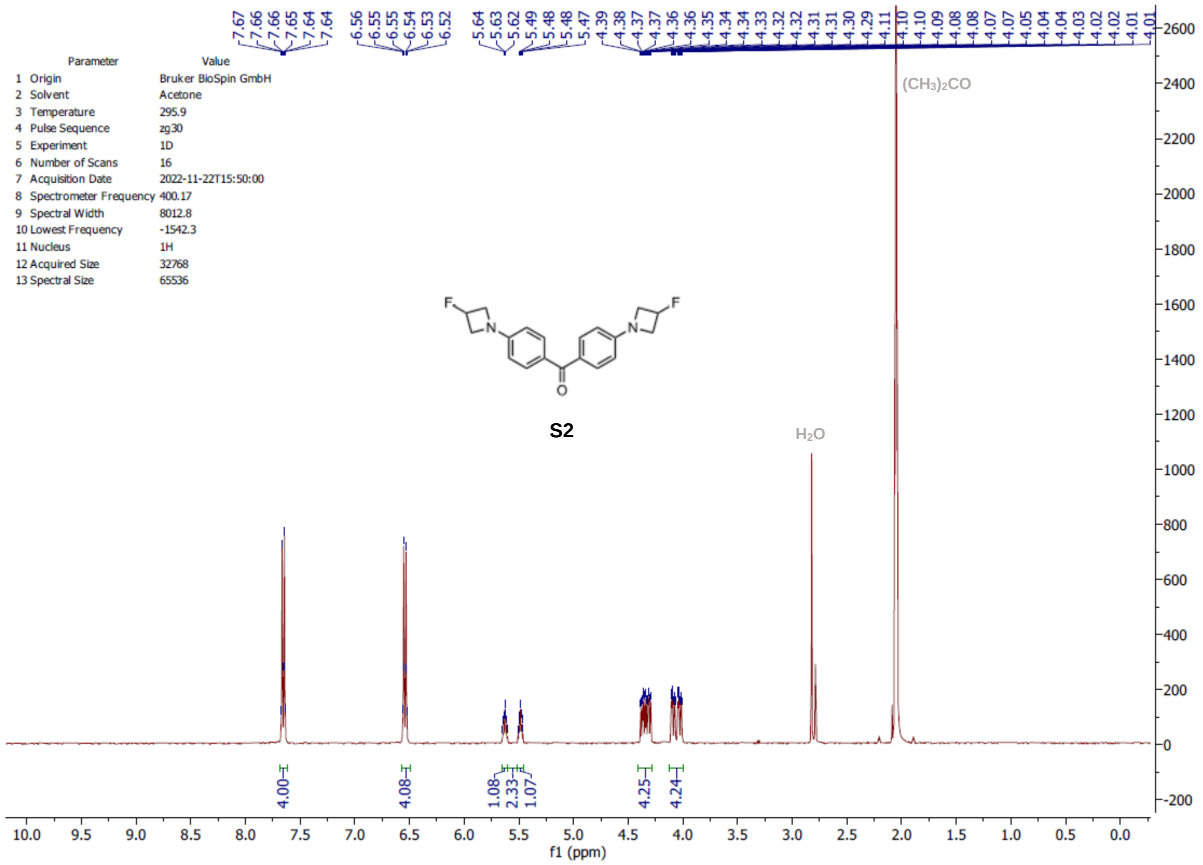
7.67 7.66 7.66 7.65 7.64 7.64
6.56 6.55 6.55 6.54 6.53 6.52
5.64 5.63 5.62 5.49 5.48 5.48 5.47
4.39 4.38 4.37 4.37 4.36 4.35 4.34 4.34 4.33 4.32 4.32 4.31 4.31 4.30 4.29 4.11 4.10 4.10 4.09 4.08 4.08 4.07 4.07 4.05 4.04 4.04 4.03 4.02 4.02 4.01 4.01
Parameter Value
1 Origin Bruker BioSpin GmbH
2 Solvent Acetone
3 Temperature 295.9
4 Pulse Sequence zg30
5 Experiment 1D
6 Number of Scans 16
7 Acquisition Date 2022-11-22T15:50:00
8 Spectrometer Frequency 400.17
9 Spectral Width 8012.8
10 Lowest Frequency -1542.3
11 Nucleus 1H
12 Acquired Size 32768
13 Spectral Size 65536
S2
H2O
(CH3)2CO
4.00 4.08 1.08 2.33 1.07 4.25 4.24
f1 (ppm)

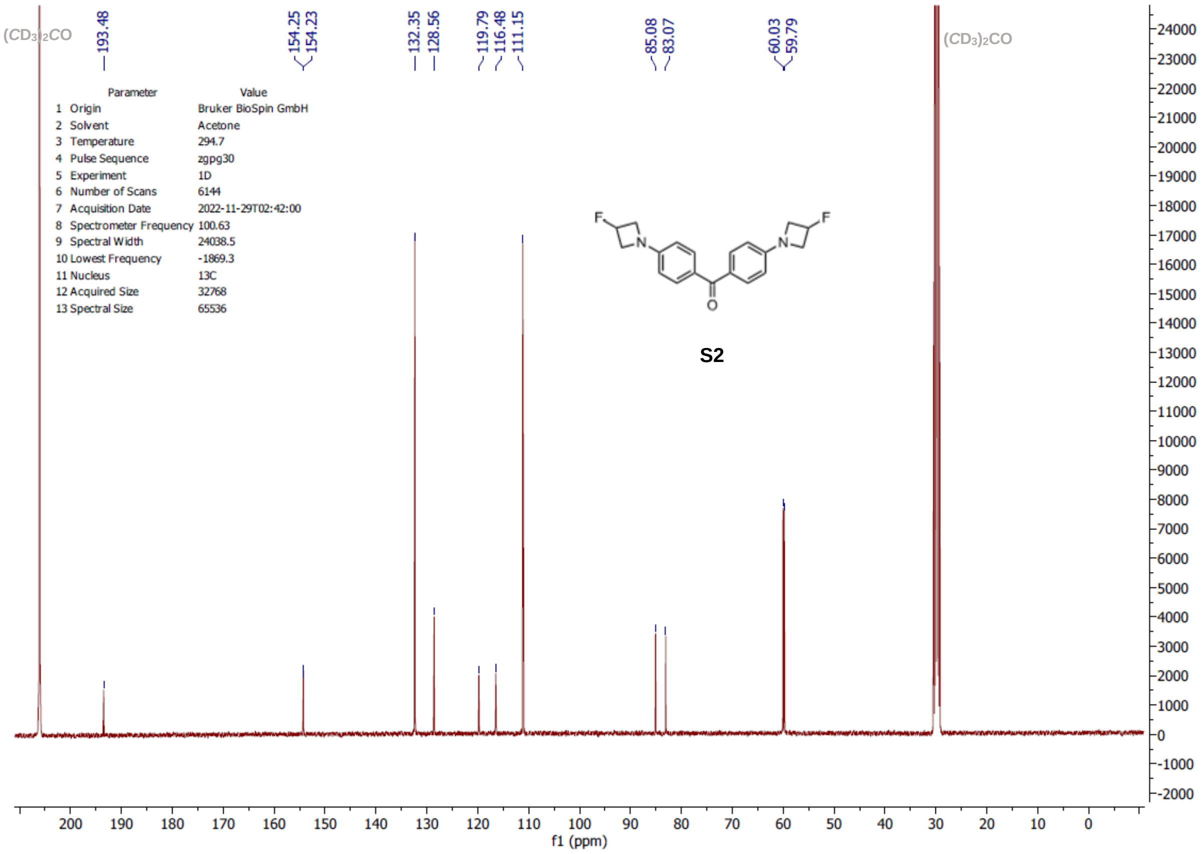
(CD3)2CO
193.48
154.25 154.23
132.35 128.56
119.79 116.48 111.15
85.08 83.07
60.03 59.79
(CD3)2CO
Parameter Value
1 Origin Bruker BioSpin GmbH
2 Solvent Acetone
3 Temperature 294.7
4 Pulse Sequence zgpg30
5 Experiment 1D
6 Number of Scans 6144
7 Acquisition Date 2022-11-29T02:42:00
8 Spectrometer Frequency 100.63
9 Spectral Width 24038.5
10 Lowest Frequency -1869.3
11 Nucleus 13C
12 Acquired Size 32768
13 Spectral Size 65536
S2
f1 (ppm)

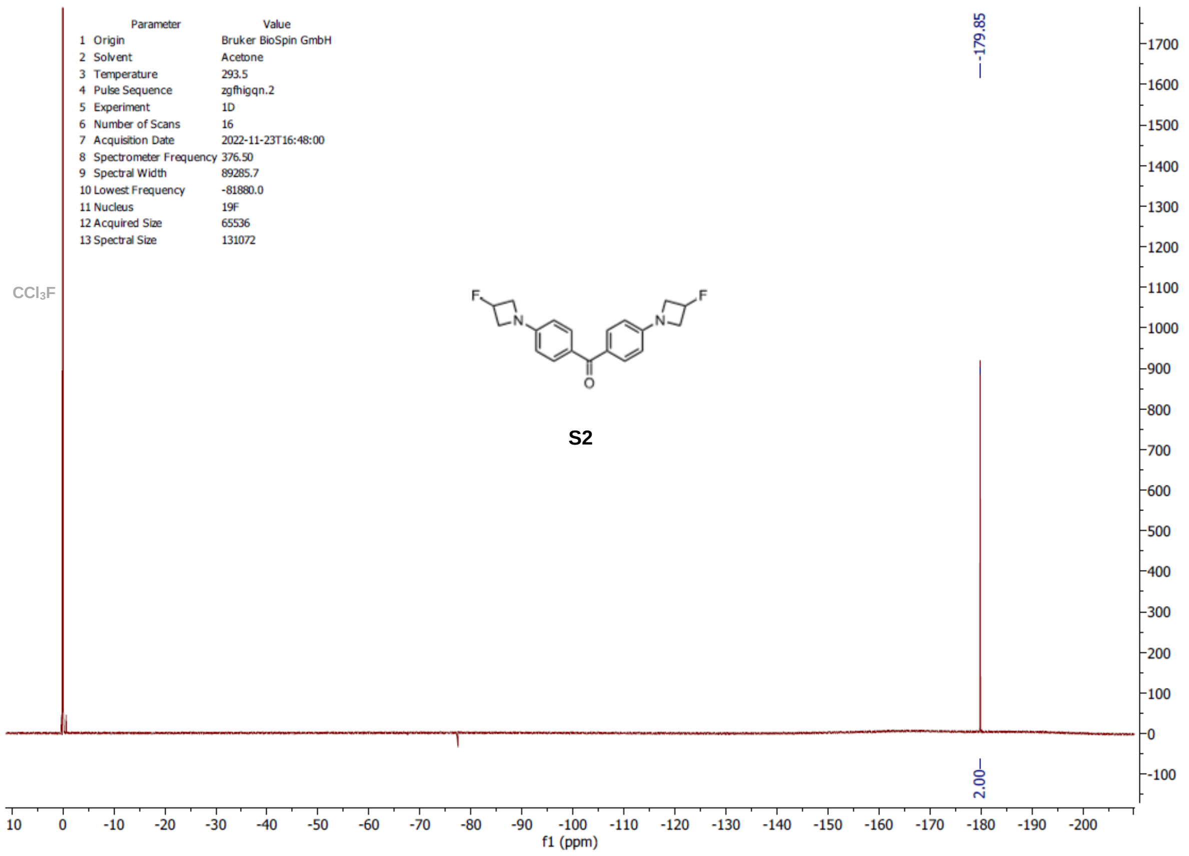

Parameter Value
1 Origin Bruker BioSpin GmbH
2 Solvent Acetone
3 Temperature 293.5
4 Pulse Sequence zgfhigqn.2
5 Experiment 1D
6 Number of Scans 16
7 Acquisition Date 2022-11-23T16:48:00
8 Spectrometer Frequency 376.50
9 Spectral Width 89285.7
10 Lowest Frequency -81880.0
11 Nucleus 19F
12 Acquired Size 65536
13 Spectral Size 131072
CCl3F
—179.85
S2
2.00
f1 (ppm)

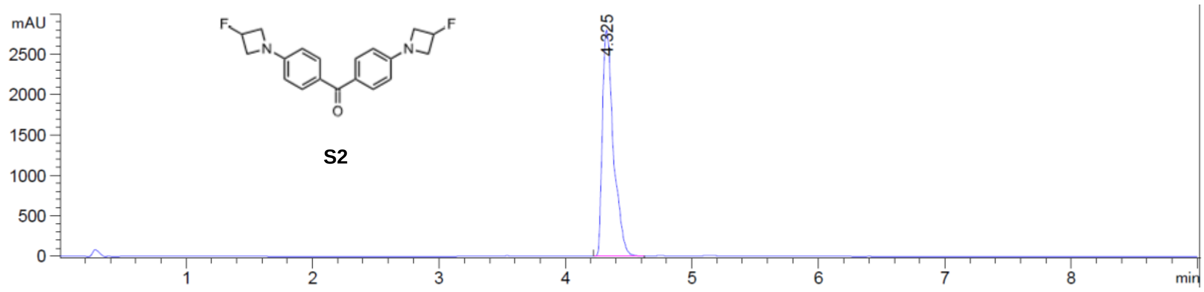

mAU
4.325
S2
min

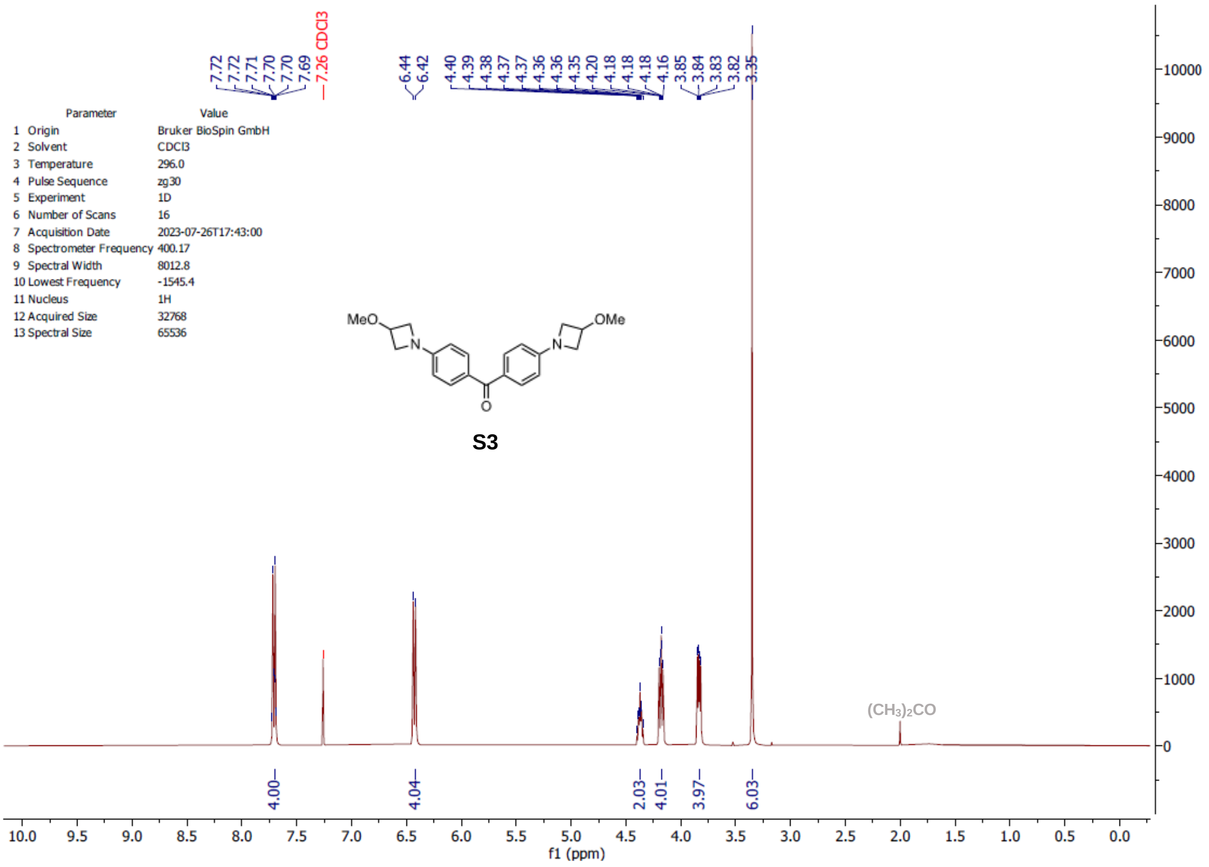
7.72
7.72
7.71
7.70
7.70
7.69
7.26 CDCl3
6.44
6.42
4.40
4.39
4.38
4.37
4.37
4.36
4.36
4.35
4.20
4.18
4.18
4.18
4.16
3.85
3.84
3.83
3.82
3.35
Parameter Value
1 Origin Bruker BioSpin GmbH
2 Solvent CDCl3
3 Temperature 296.0
4 Pulse Sequence zg30
5 Experiment 1D
6 Number of Scans 16
7 Acquisition Date 2023-07-26T17:43:00
8 Spectrometer Frequency 400.17
9 Spectral Width 8012.8
10 Lowest Frequency -1545.4
11 Nucleus 1H
12 Acquired Size 32768
13 Spectral Size 65536
MeO
N
N
OMe
O
S3
(CH3)2CO
4.00
4.04
2.03
4.01
3.97
6.03
f1 (ppm)

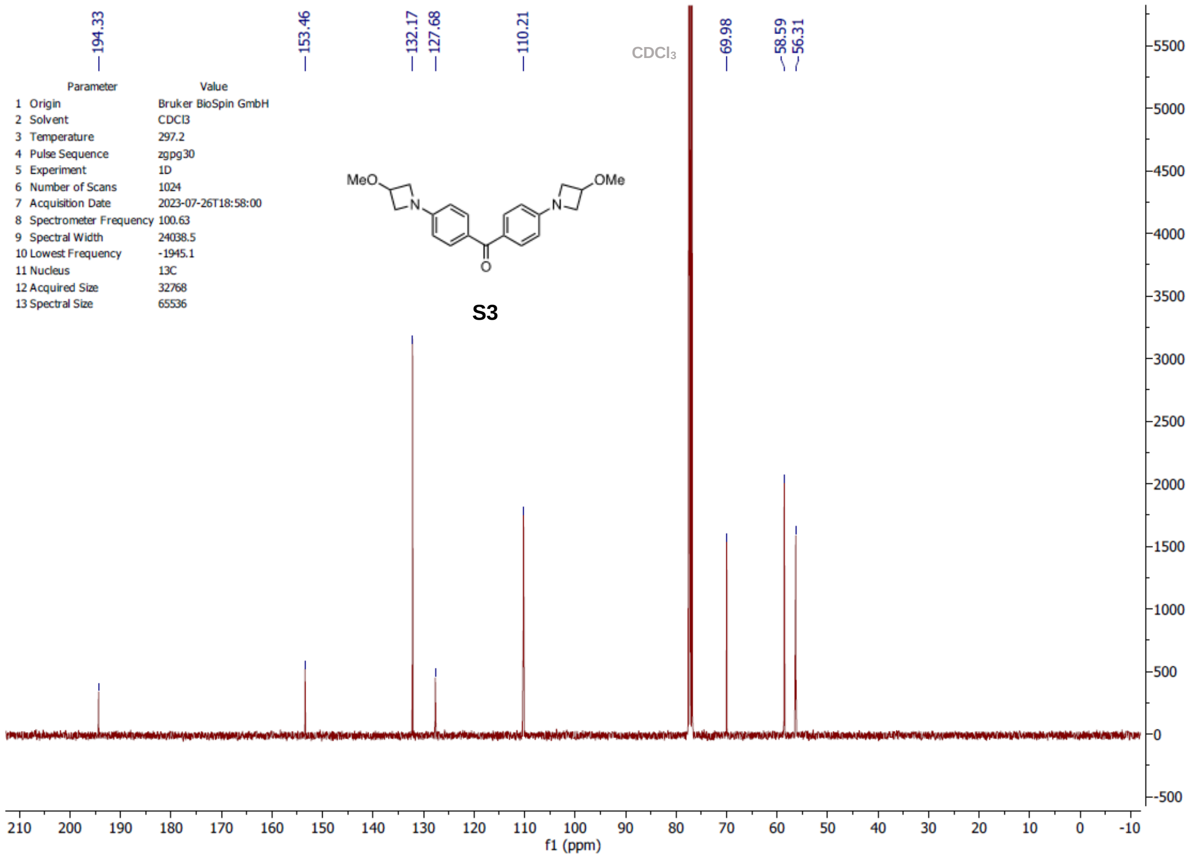
194.33
153.46
132.17
127.68
110.21
CDCl3
69.98
58.59
56.31
Parameter Value
1 Origin Bruker BioSpin GmbH
2 Solvent CDCl3
3 Temperature 297.2
4 Pulse Sequence zgpg30
5 Experiment 1D
6 Number of Scans 1024
7 Acquisition Date 2023-07-26T18:58:00
8 Spectrometer Frequency 100.63
9 Spectral Width 24038.5
10 Lowest Frequency -1945.1
11 Nucleus 13C
12 Acquired Size 32768
13 Spectral Size 65536
MeO
N
N
OMe
O
S3
f1 (ppm)

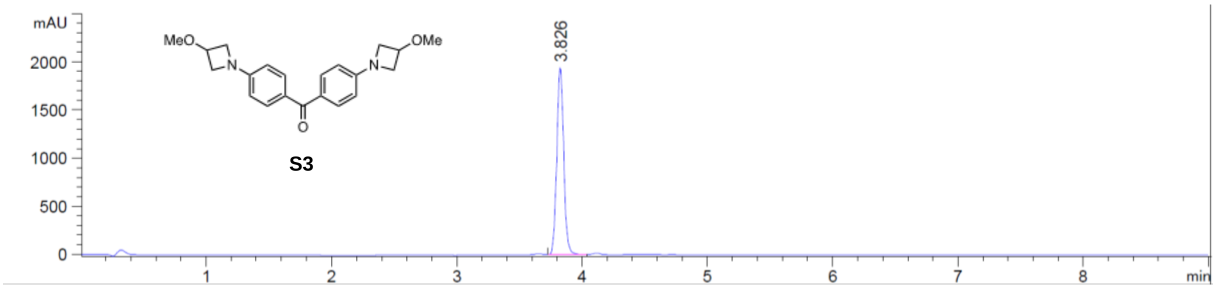
mAU
2000
1500
1000
500
0
MeO
N
O
N
OMe
S3
3.826
1
2
3
4
5
6
7
8
min

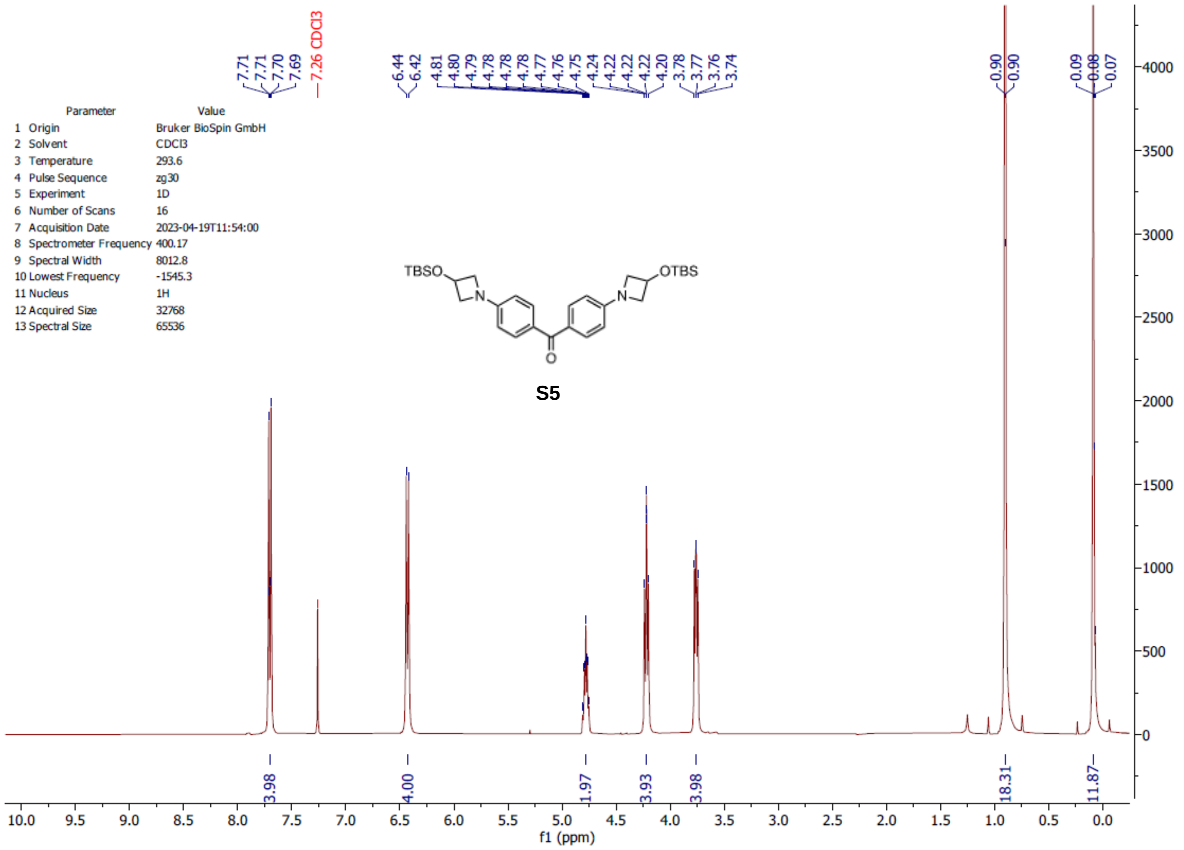

7.71
7.71
7.70
7.69
7.26 CDCl3
6.44
6.42
4.81
4.80
4.79
4.78
4.78
4.78
4.77
4.76
4.75
4.24
4.22
4.22
4.22
4.20
3.78
3.77
3.76
3.74
0.90
0.90
0.09
0.08
0.07
Parameter Value
1 Origin Bruker BioSpin GmbH
2 Solvent CDCl3
3 Temperature 293.6
4 Pulse Sequence zg30
5 Experiment 1D
6 Number of Scans 16
7 Acquisition Date 2023-04-19T11:54:00
8 Spectrometer Frequency 400.17
9 Spectral Width 8012.8
10 Lowest Frequency -1545.3
11 Nucleus 1H
12 Acquired Size 32768
13 Spectral Size 65536
TBSO
N
N
OTBS
O
S5
3.98
4.00
1.97
3.93
3.98
18.31
11.87
f1 (ppm)

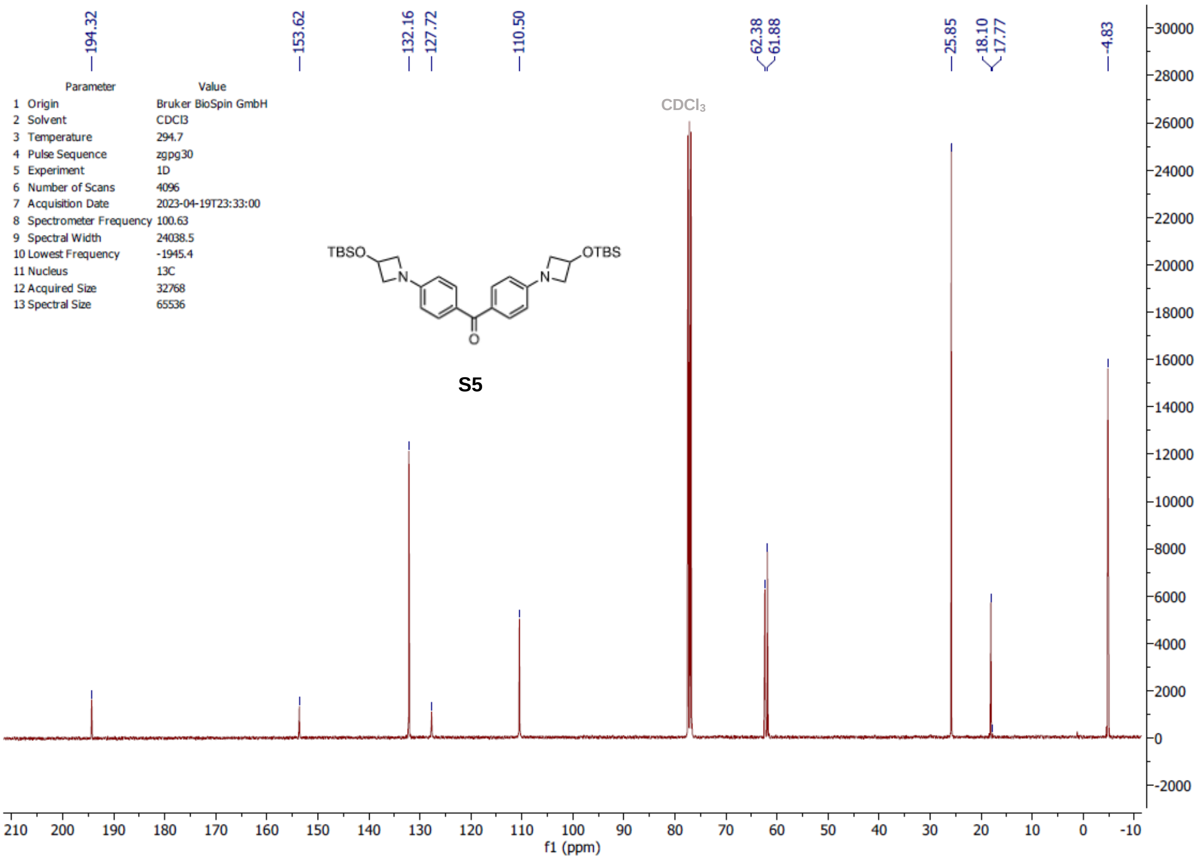

194.32
153.62
132.16
127.72
110.50
62.38
61.88
25.85
18.10
17.77
-4.83
Parameter Value
1 Origin Bruker BioSpin GmbH
2 Solvent CDCl3
3 Temperature 294.7
4 Pulse Sequence zgpg30
5 Experiment 1D
6 Number of Scans 4096
7 Acquisition Date 2023-04-19T23:33:00
8 Spectrometer Frequency 100.63
9 Spectral Width 24038.5
10 Lowest Frequency -1945.4
11 Nucleus 13C
12 Acquired Size 32768
13 Spectral Size 65536
CDCl₃
TBSO
N
N
OTBS
O
S5
f1 (ppm)

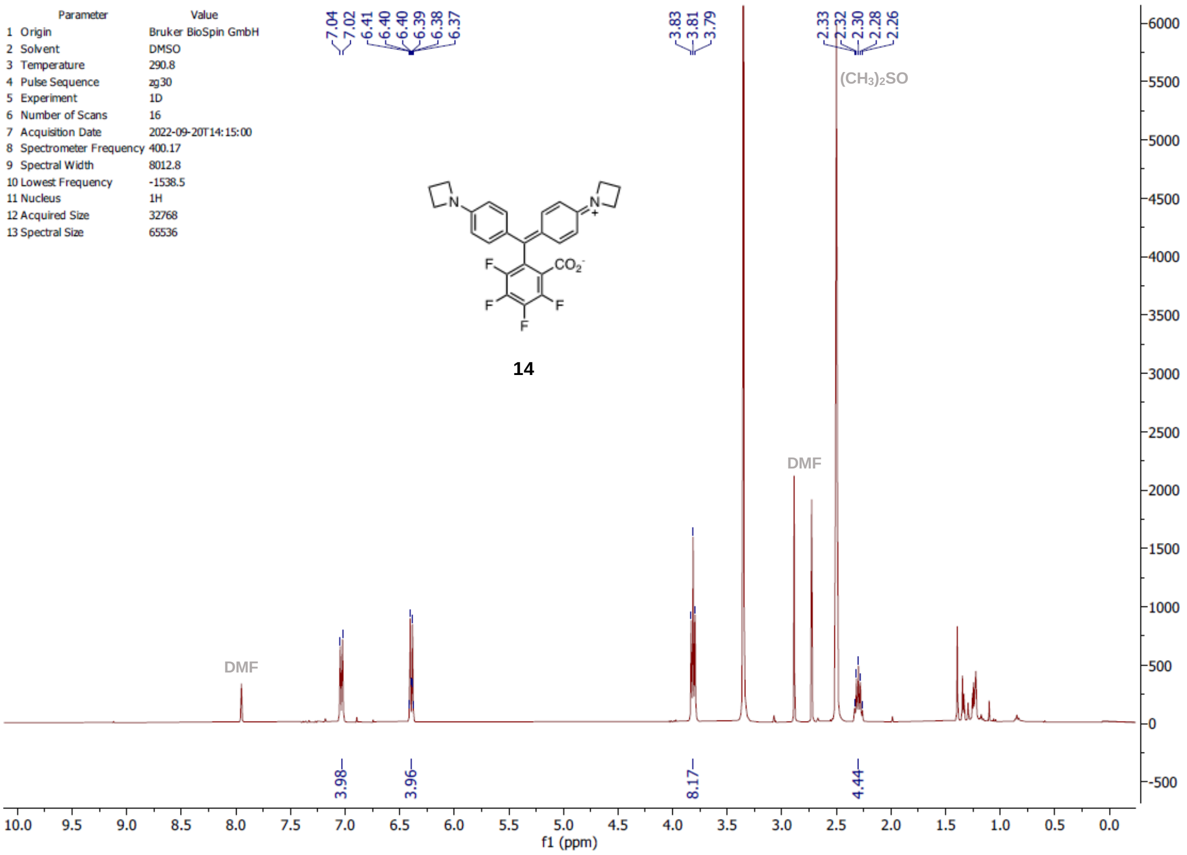
Parameter
Value
1 Origin
Bruker BioSpin GmbH
2 Solvent
DMSO
3 Temperature
290.8
4 Pulse Sequence
zg30
5 Experiment
1D
6 Number of Scans
16
7 Acquisition Date
2022-09-20T14:15:00
8 Spectrometer Frequency
400.17
9 Spectral Width
8012.8
10 Lowest Frequency
-1538.5
11 Nucleus
1H
12 Acquired Size
32768
13 Spectral Size
65536
14
DMF
(CH3)2SO
f1 (ppm)

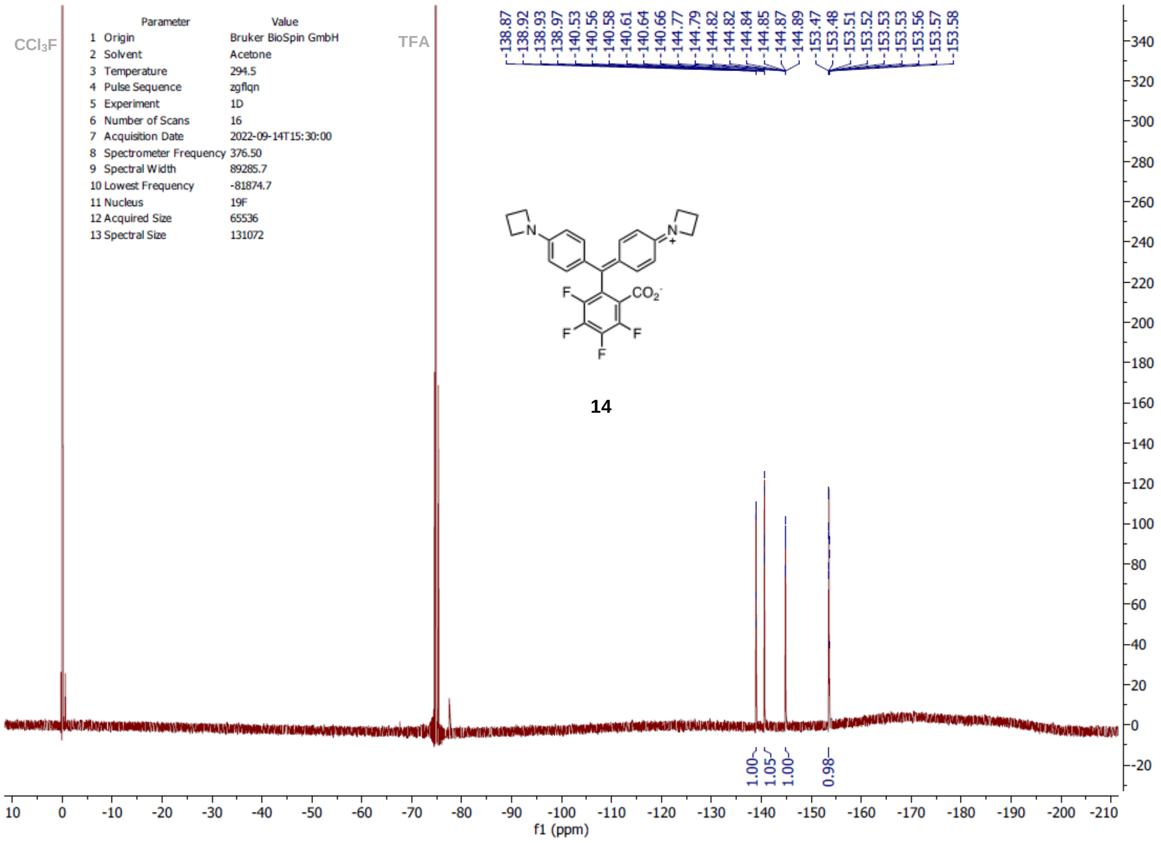
CCl3F
TFA
Parameter
Value
1 Origin
Bruker BioSpin GmbH
2 Solvent
Acetone
3 Temperature
294.5
4 Pulse Sequence
zgflqn
5 Experiment
1D
6 Number of Scans
16
7 Acquisition Date
2022-09-14T15:30:00
8 Spectrometer Frequency
376.50
9 Spectral Width
89285.7
10 Lowest Frequency
-81874.7
11 Nucleus
19F
12 Acquired Size
65536
13 Spectral Size
131072
14
f1 (ppm)

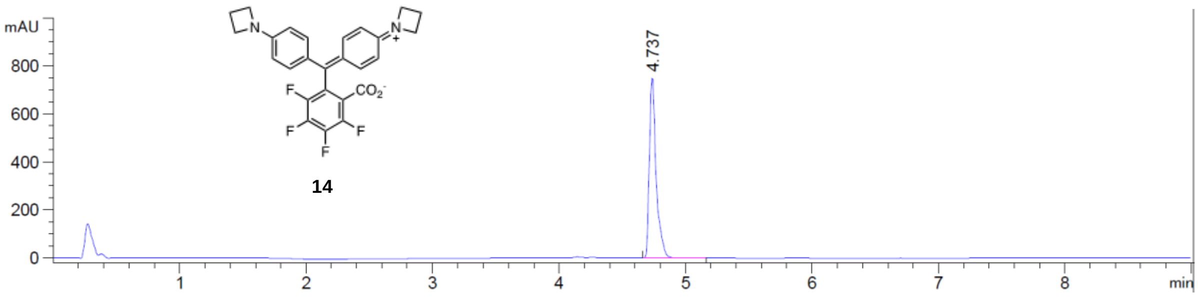
mAU
800
600
400
200
0
4.737
14
1
2
3
4
5
6
7
8
min

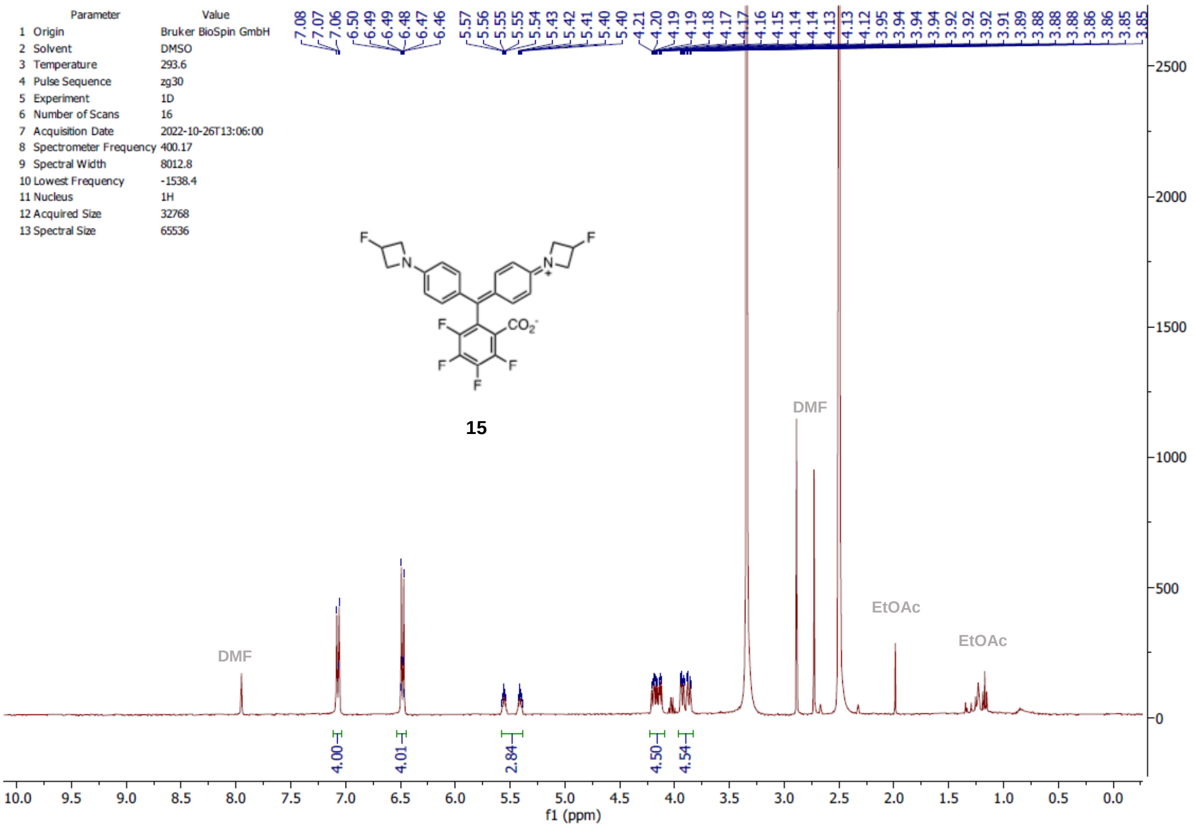

Parameter Value
1 Origin Bruker BioSpin GmbH
2 Solvent DMSO
3 Temperature 293.6
4 Pulse Sequence zg30
5 Experiment 1D
6 Number of Scans 16
7 Acquisition Date 2022-10-26T13:06:00
8 Spectrometer Frequency 400.17
9 Spectral Width 8012.8
10 Lowest Frequency -1538.4
11 Nucleus 1H
12 Acquired Size 32768
13 Spectral Size 65536
7.08 7.07 7.06 6.50 6.49 6.49 6.48 6.47 6.46 5.57 5.56 5.55 5.55 5.54 5.43 5.42 5.41 5.40 5.40 4.21 4.20 4.19 4.19 4.18 4.17 4.17 4.16 4.15 4.14 4.14 4.13 4.13 4.12 3.95 3.94 3.94 3.94 3.92 3.92 3.92 3.91 3.89 3.88 3.88 3.86 3.86 3.85 3.85
15
DMF
EtOAc
EtOAc
DMF
4.00
4.01
2.84
4.50
4.54
f1 (ppm)

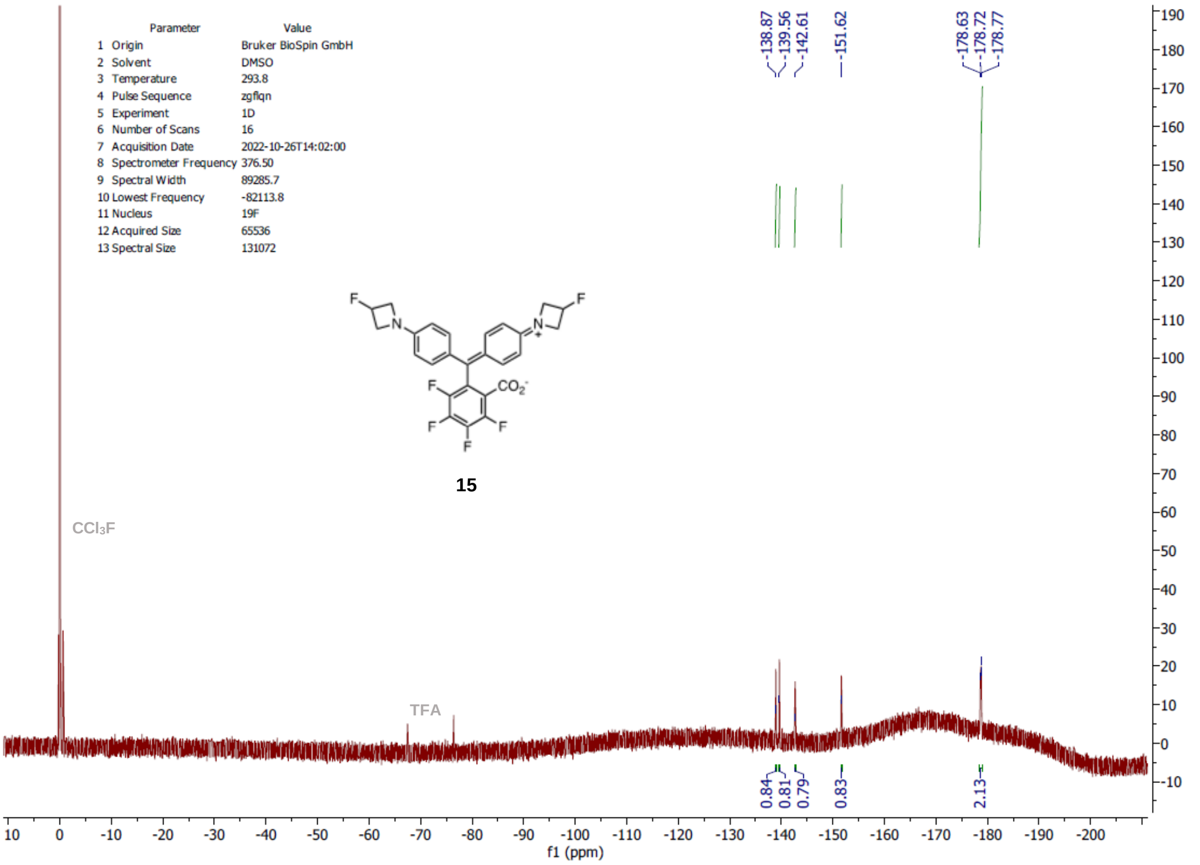

Parameter Value
1 Origin Bruker BioSpin GmbH
2 Solvent DMSO
3 Temperature 293.8
4 Pulse Sequence zgflqn
5 Experiment 1D
6 Number of Scans 16
7 Acquisition Date 2022-10-26T14:02:00
8 Spectrometer Frequency 376.50
9 Spectral Width 89285.7
10 Lowest Frequency -82113.8
11 Nucleus 19F
12 Acquired Size 65536
13 Spectral Size 131072
-138.87 -139.56 -142.61 -151.62 -178.63 -178.72 -178.77
15
CCl3F
TFA
0.84 0.81 0.79 0.83 2.13
f1 (ppm)

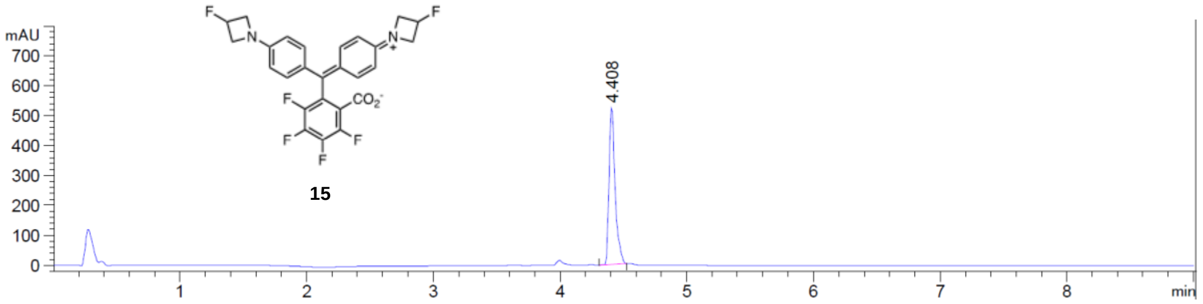

mAU
700
600
500
400
300
200
100
0
4.408
1
2
3
4
5
6
7
8
min
F
N
N
F
CO2-
F
F
F
F

**15**

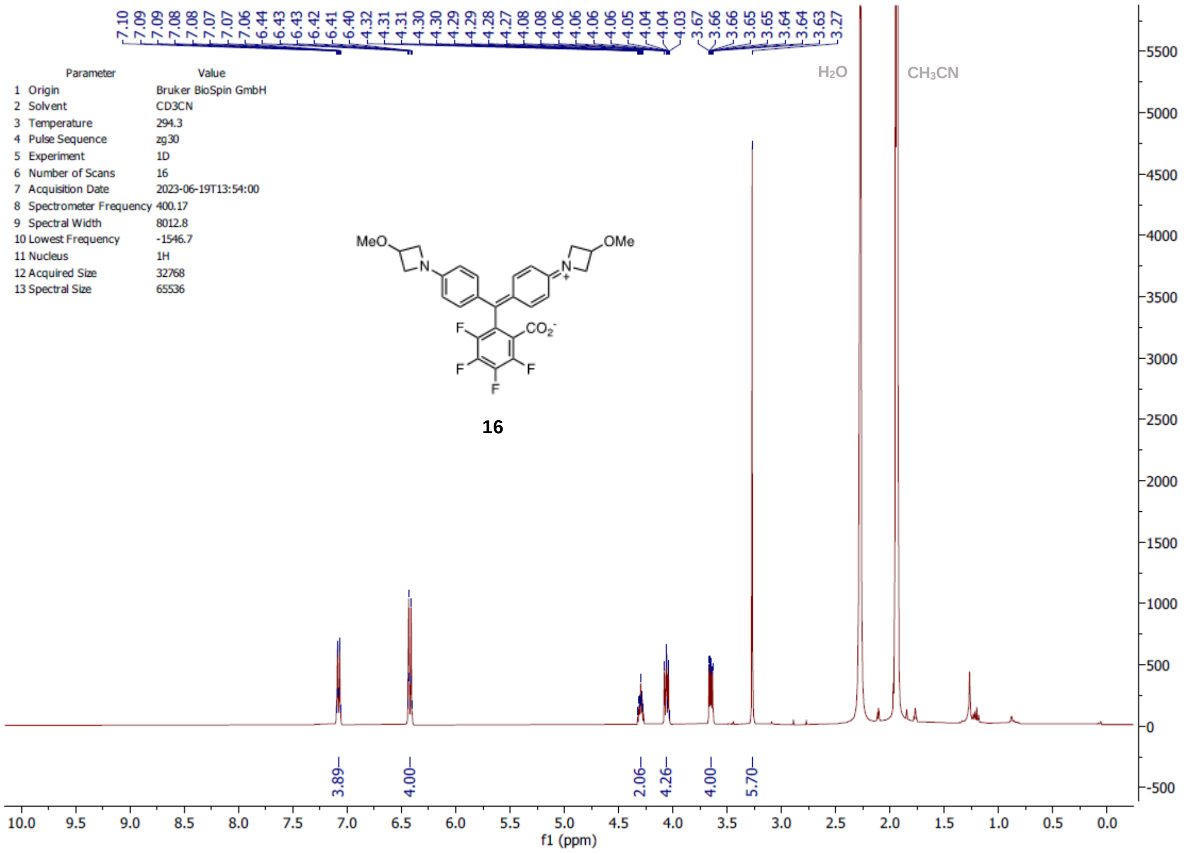
Parameter Value
1 Origin Bruker BioSpin GmbH
2 Solvent CD3CN
3 Temperature 294.3
4 Pulse Sequence zg30
5 Experiment 1D
6 Number of Scans 16
7 Acquisition Date 2023-06-19T13:54:00
8 Spectrometer Frequency 400.17
9 Spectral Width 8012.8
10 Lowest Frequency -1546.7
11 Nucleus 1H
12 Acquired Size 32768
13 Spectral Size 65536
H2O
CH3CN
MeO
OMe
16
3.89
4.00
2.06
4.26
4.00
5.70
f1 (ppm)

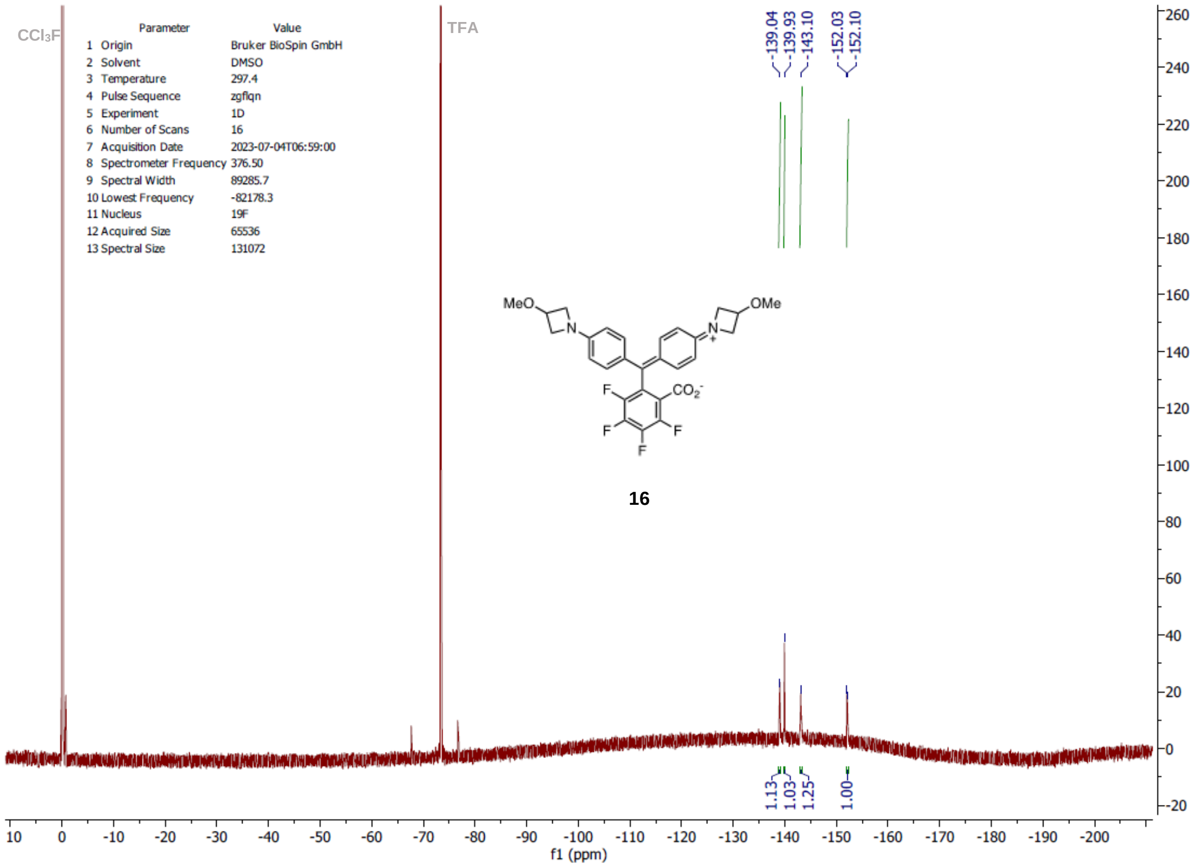
CCl3F
TFA
Parameter Value
1 Origin Bruker BioSpin GmbH
2 Solvent DMSO
3 Temperature 297.4
4 Pulse Sequence zgflqn
5 Experiment 1D
6 Number of Scans 16
7 Acquisition Date 2023-07-04T06:59:00
8 Spectrometer Frequency 376.50
9 Spectral Width 89285.7
10 Lowest Frequency -82178.3
11 Nucleus 19F
12 Acquired Size 65536
13 Spectral Size 131072
-139.04
-139.93
-143.10
-152.03
-152.10
MeO
OMe
16
1.13
1.03
1.25
1.00
f1 (ppm)

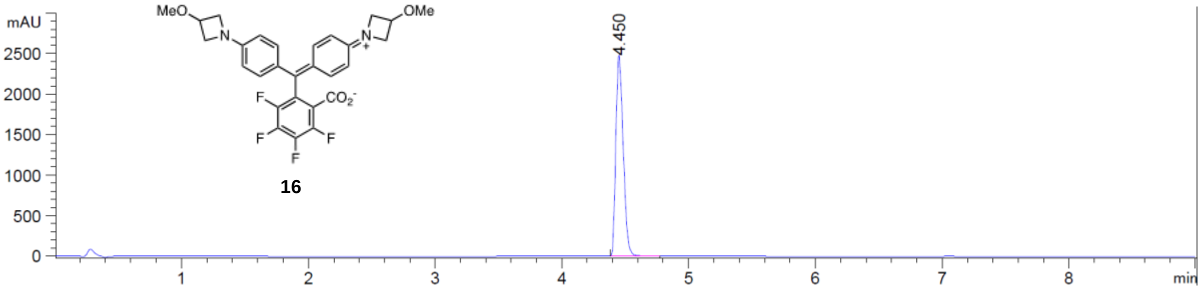
mAU
2500
2000
1500
1000
500
0
1
2
3
4
5
6
7
8
min
4.450
MeO
OMe
N
N
+
CO2-
F
F
F
F
16

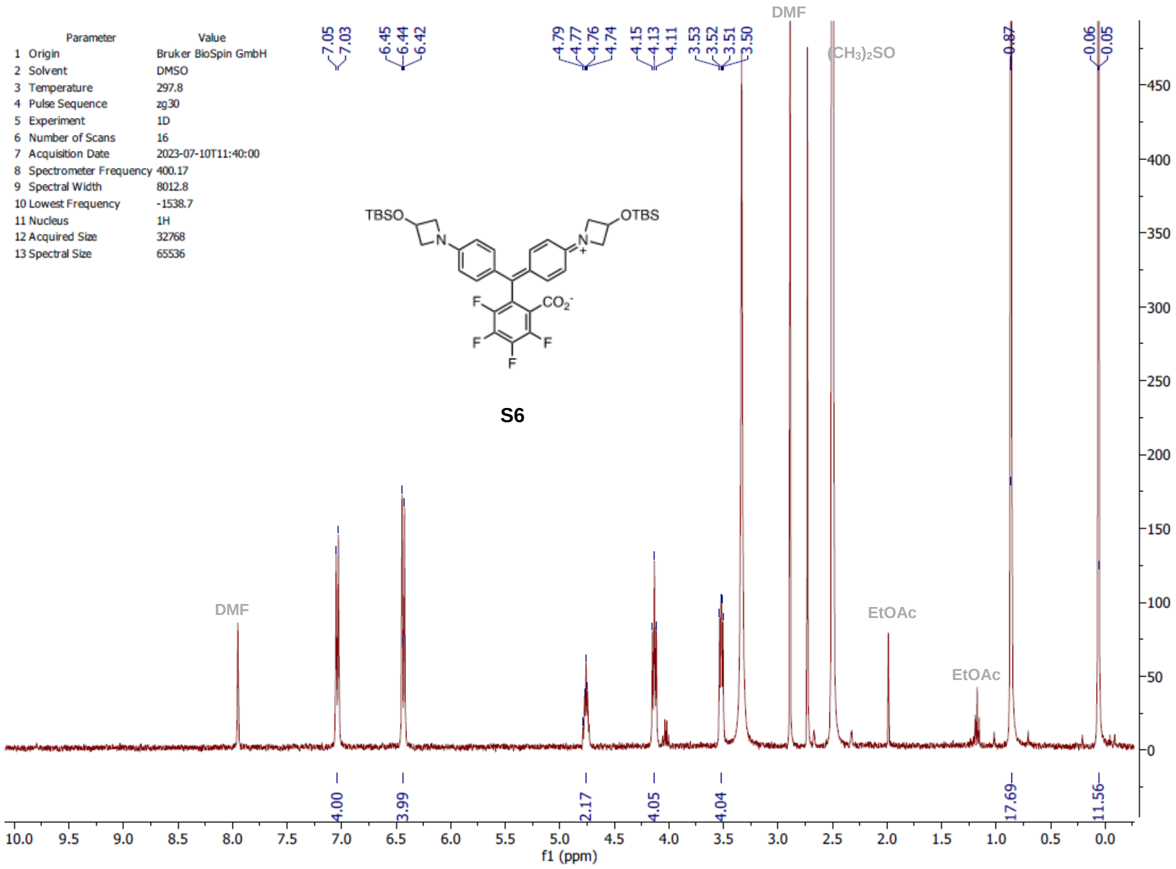
Parameter Value
1 Origin Bruker BioSpin GmbH
2 Solvent DMSO
3 Temperature 297.8
4 Pulse Sequence zg30
5 Experiment 1D
6 Number of Scans 16
7 Acquisition Date 2023-07-10T11:40:00
8 Spectrometer Frequency 400.17
9 Spectral Width 8012.8
10 Lowest Frequency -1538.7
11 Nucleus 1H
12 Acquired Size 32768
13 Spectral Size 65536
7.05
7.03
6.45
6.44
6.42
4.79
4.77
4.76
4.74
4.15
4.13
4.11
3.53
3.52
3.51
3.50
0.87
0.06
0.05
DMF
(CH3)2SO
EtOAc
TBSO
OTBS
N
CO2-
F
S6
4.00
3.99
2.17
4.05
4.04
17.69
11.56
f1 (ppm)

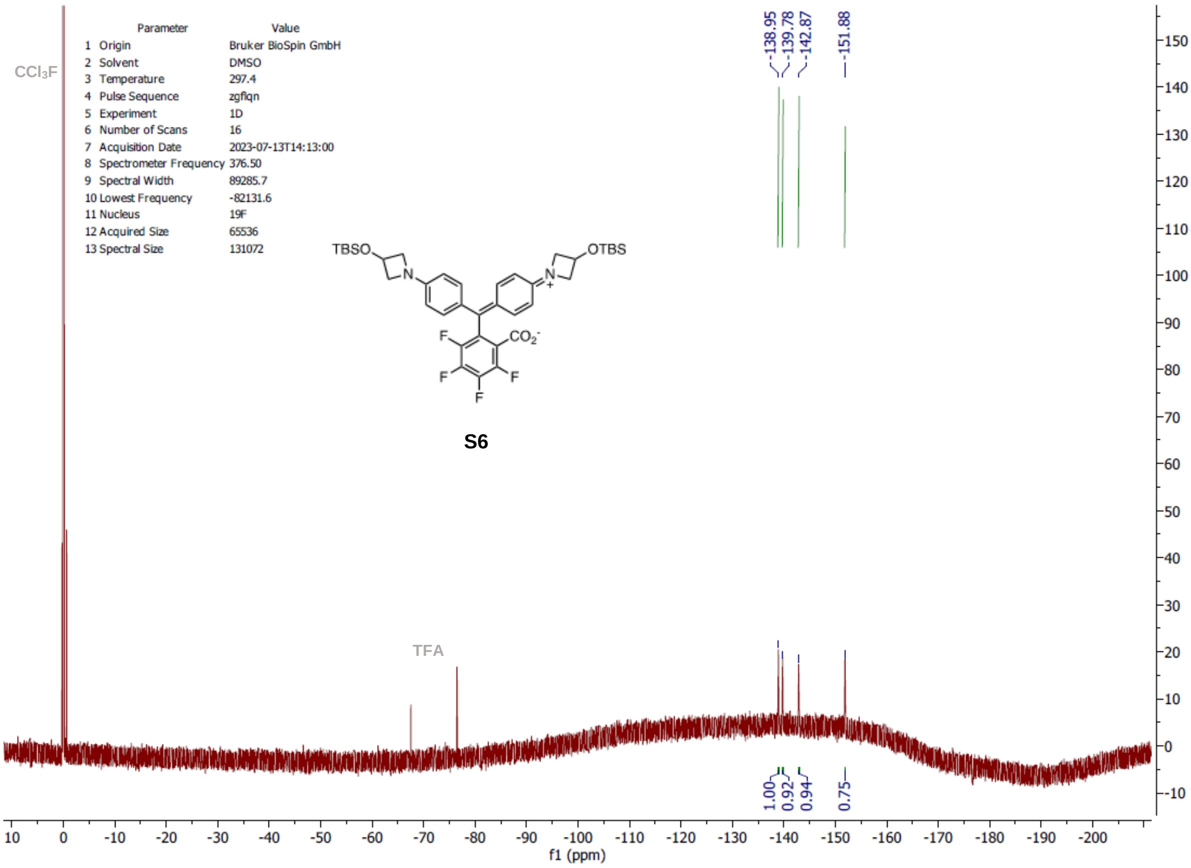
Parameter Value
1 Origin Bruker BioSpin GmbH
2 Solvent DMSO
3 Temperature 297.4
4 Pulse Sequence zgflqn
5 Experiment 1D
6 Number of Scans 16
7 Acquisition Date 2023-07-13T14:13:00
8 Spectrometer Frequency 376.50
9 Spectral Width 89285.7
10 Lowest Frequency -82131.6
11 Nucleus 19F
12 Acquired Size 65536
13 Spectral Size 131072
CCl3F
-138.95
-139.78
-142.87
-151.88
TFA
TBSO
OTBS
N
CO2-
F
S6
1.00
0.92
0.94
0.75
f1 (ppm)

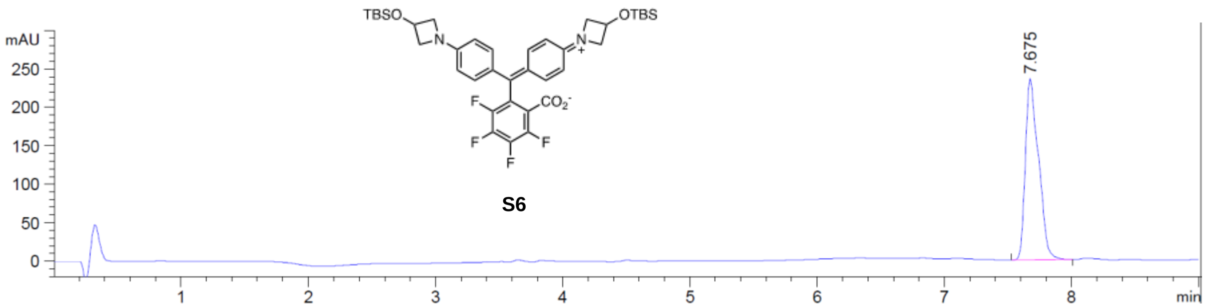
TBSO
OTBS
N
N
F
$CO_2^-$
F
F
F
F
S6
mAU
250
200
150
100
50
0
7.675
1
2
3
4
5
6
7
8
min

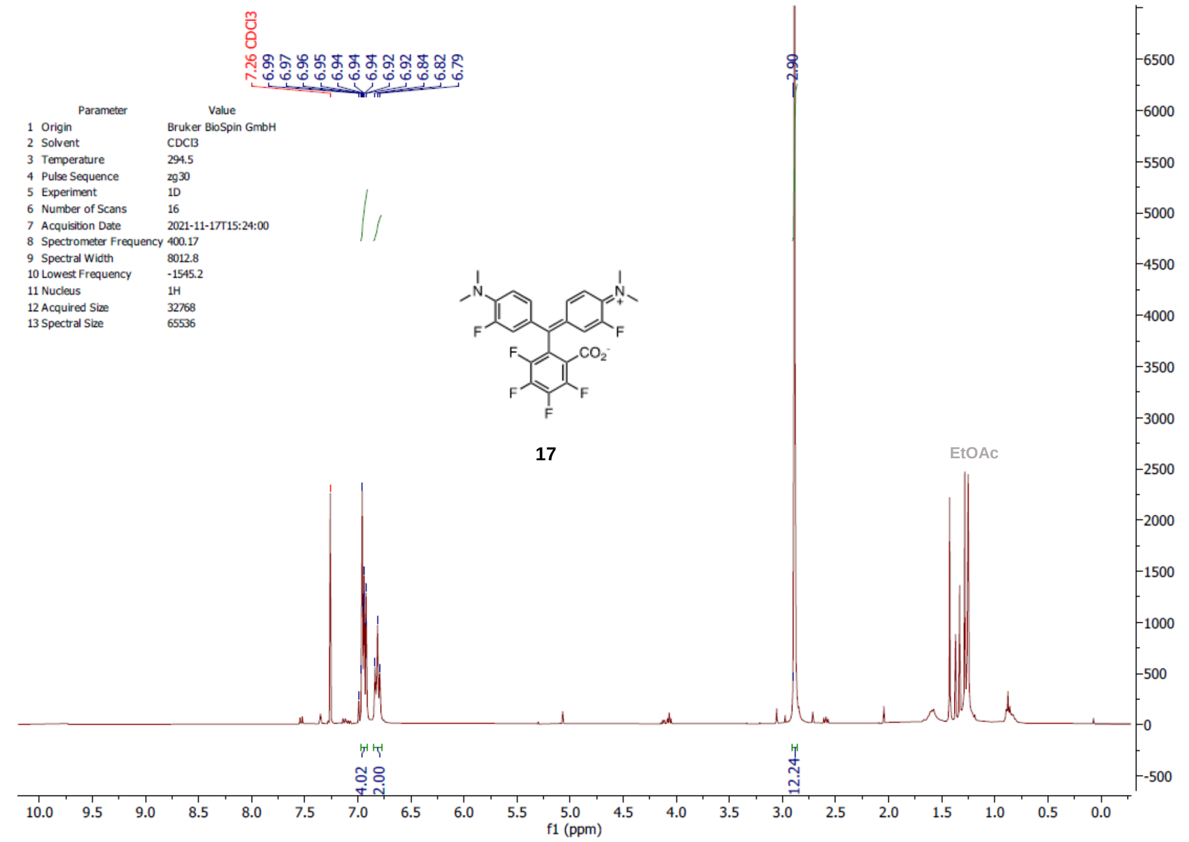

7.26 CDCl3
6.99
6.97
6.96
6.95
6.94
6.94
6.94
6.94
6.92
6.92
6.92
6.84
6.82
6.79
2.90
Parameter Value
1 Origin Bruker BioSpin GmbH
2 Solvent CDCl3
3 Temperature 294.5
4 Pulse Sequence zg30
5 Experiment 1D
6 Number of Scans 16
7 Acquisition Date 2021-11-17T15:24:00
8 Spectrometer Frequency 400.17
9 Spectral Width 8012.8
10 Lowest Frequency -1545.2
11 Nucleus 1H
12 Acquired Size 32768
13 Spectral Size 65536
17
EtOAc
4.02
2.00
12.24
f1 (ppm)

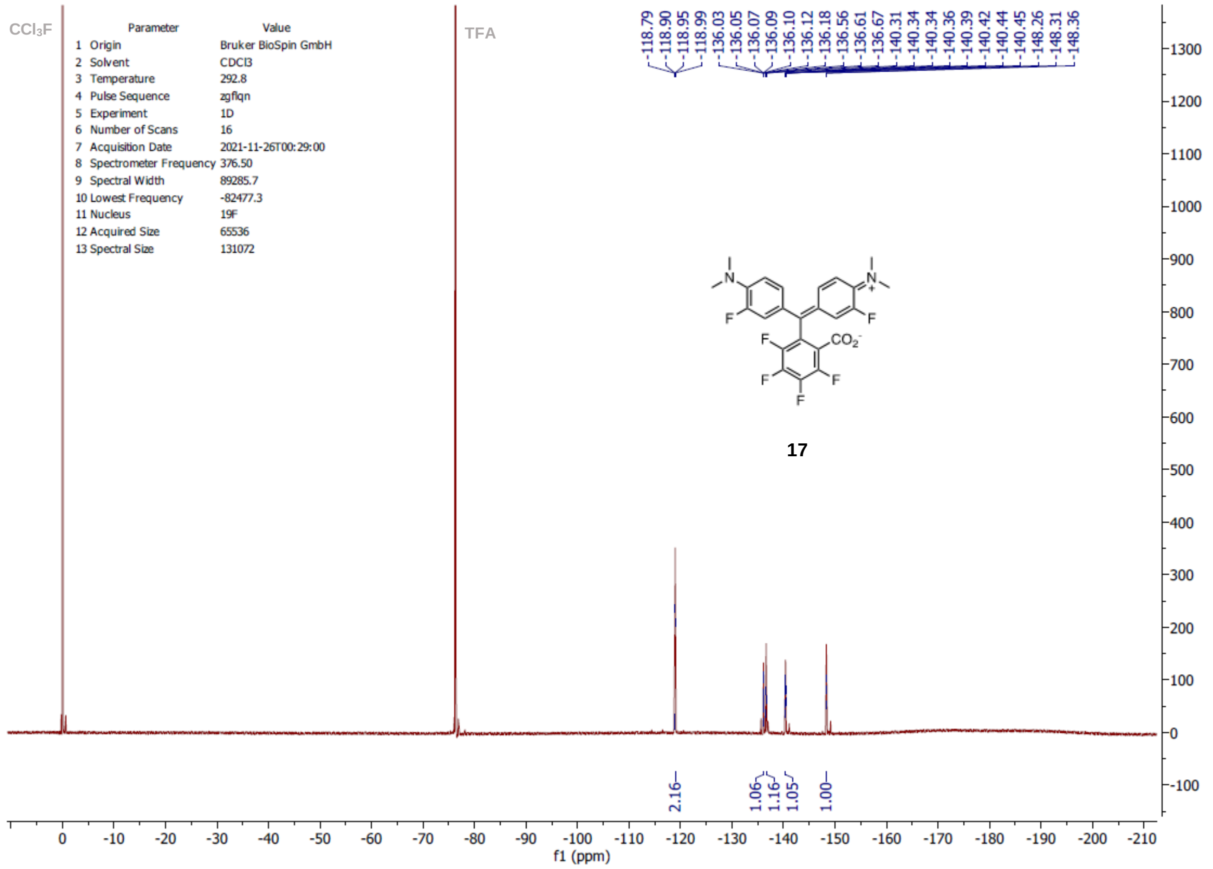

CCl3F
TFA
-118.79
-118.90
-118.95
-118.99
-136.03
-136.05
-136.07
-136.09
-136.10
-136.12
-136.18
-136.56
-136.61
-136.67
-140.31
-140.34
-140.34
-140.36
-140.39
-140.42
-140.44
-140.45
-148.26
-148.31
-148.36
Parameter Value
1 Origin Bruker BioSpin GmbH
2 Solvent CDCl3
3 Temperature 292.8
4 Pulse Sequence zgflqn
5 Experiment 1D
6 Number of Scans 16
7 Acquisition Date 2021-11-26T00:29:00
8 Spectrometer Frequency 376.50
9 Spectral Width 89285.7
10 Lowest Frequency -82477.3
11 Nucleus 19F
12 Acquired Size 65536
13 Spectral Size 131072
17
2.16
1.06
1.16
1.05
1.00
f1 (ppm)

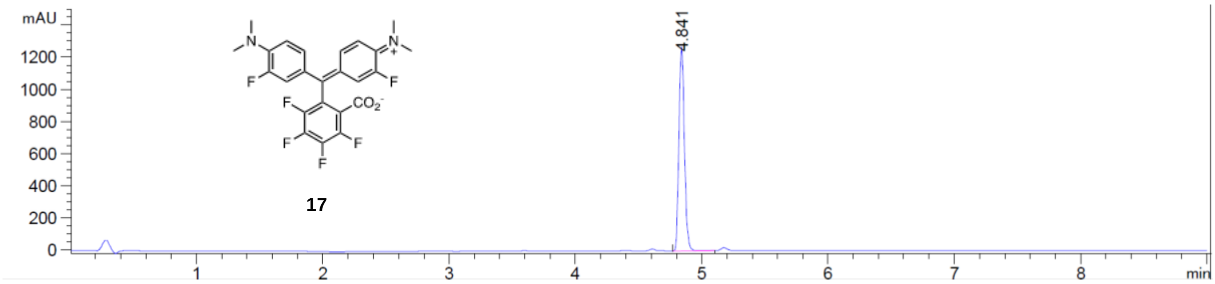

17

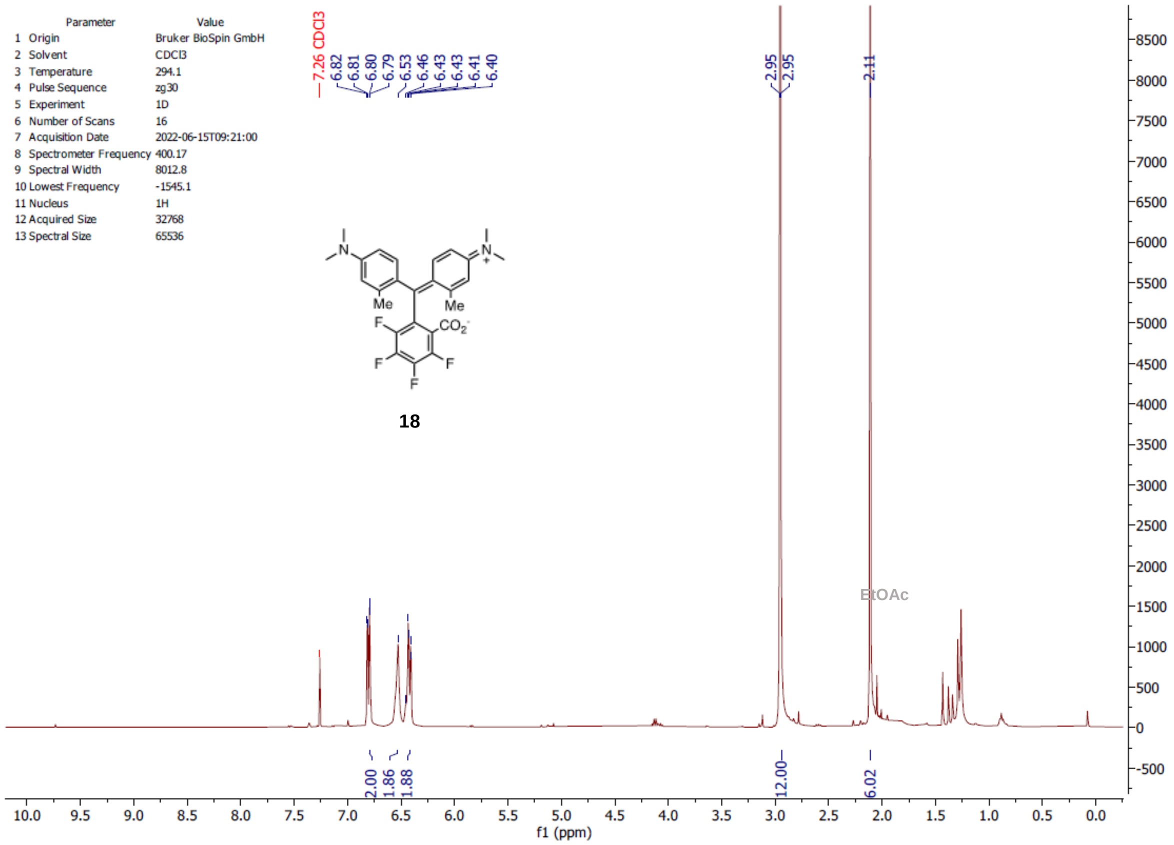
Parameter Value
1 Origin Bruker BioSpin GmbH
2 Solvent CDCl3
3 Temperature 294.1
4 Pulse Sequence zg30
5 Experiment 1D
6 Number of Scans 16
7 Acquisition Date 2022-06-15T09:21:00
8 Spectrometer Frequency 400.17
9 Spectral Width 8012.8
10 Lowest Frequency -1545.1
11 Nucleus 1H
12 Acquired Size 32768
13 Spectral Size 65536
7.26 CDCl3
6.82 6.81 6.80 6.79 6.53 6.46 6.43 6.43 6.41 6.40
2.95 2.95
2.11
Me
Me
CO2
F
F
F
F
18
EtOAc
2.00 1.86 1.88
12.00
6.02
f1 (ppm)

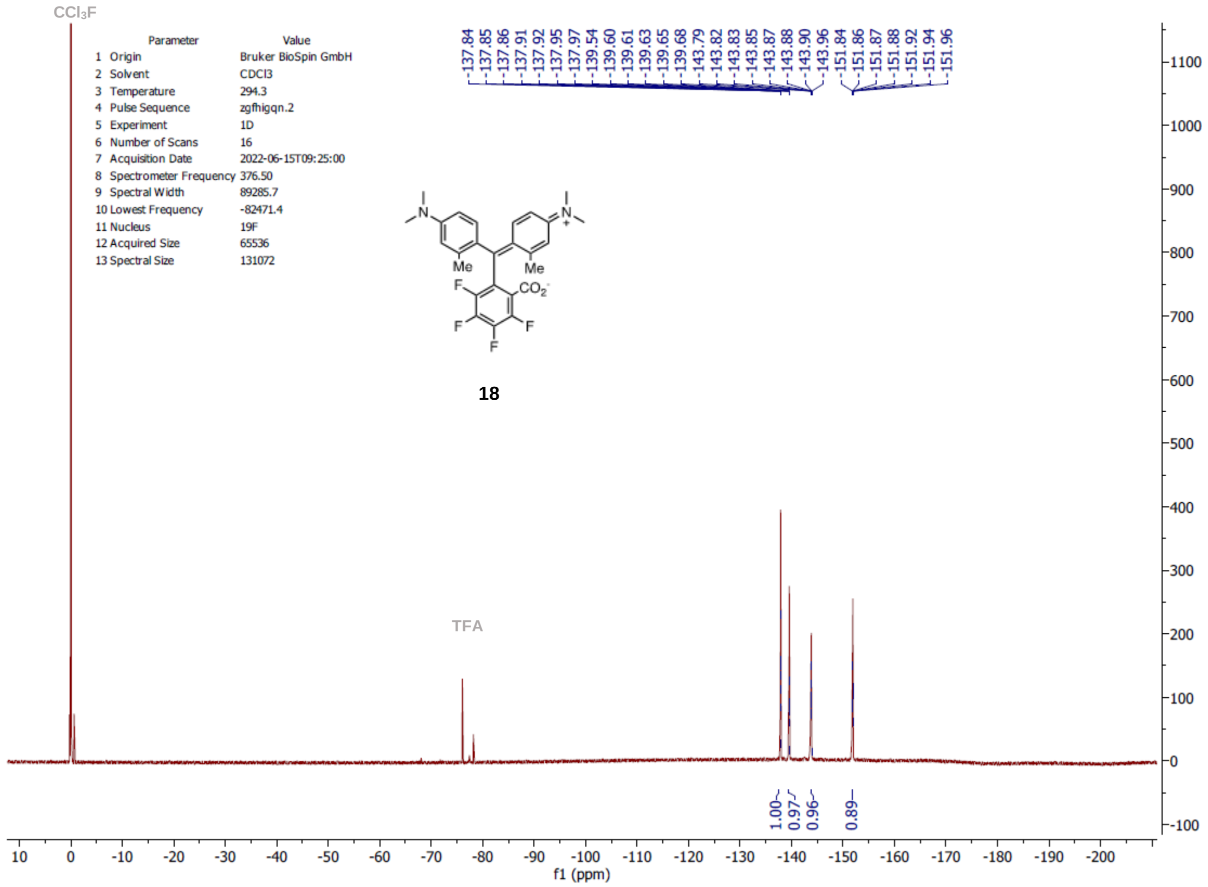
CCl3F
Parameter Value
1 Origin Bruker BioSpin GmbH
2 Solvent CDCl3
3 Temperature 294.3
4 Pulse Sequence zgfhigqn.2
5 Experiment 1D
6 Number of Scans 16
7 Acquisition Date 2022-06-15T09:25:00
8 Spectrometer Frequency 376.50
9 Spectral Width 89285.7
10 Lowest Frequency -82471.4
11 Nucleus 19F
12 Acquired Size 65536
13 Spectral Size 131072
-137.84 -137.85 -137.86 -137.91 -137.92 -137.95 -137.97 -139.54 -139.60 -139.61 -139.63 -139.65 -139.68 -143.79 -143.82 -143.83 -143.85 -143.87 -143.88 -143.90 -143.96 -151.84 -151.86 -151.87 -151.88 -151.92 -151.94 -151.96
Me
Me
CO2
F
F
F
F
18
TFA
1.00 0.97 0.96 0.89
f1 (ppm)

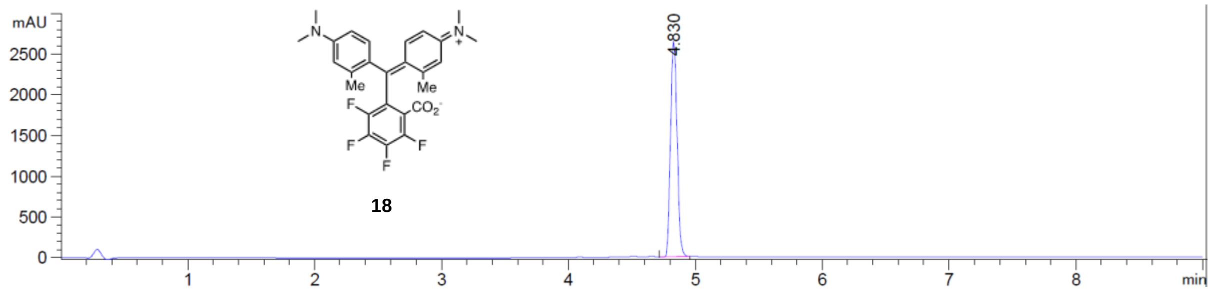
mAU
2500
2000
1500
1000
500
0
1
2
3
4
5
6
7
8
min
4.830
Me
Me
F
$CO_2^-$
F
F
F
18

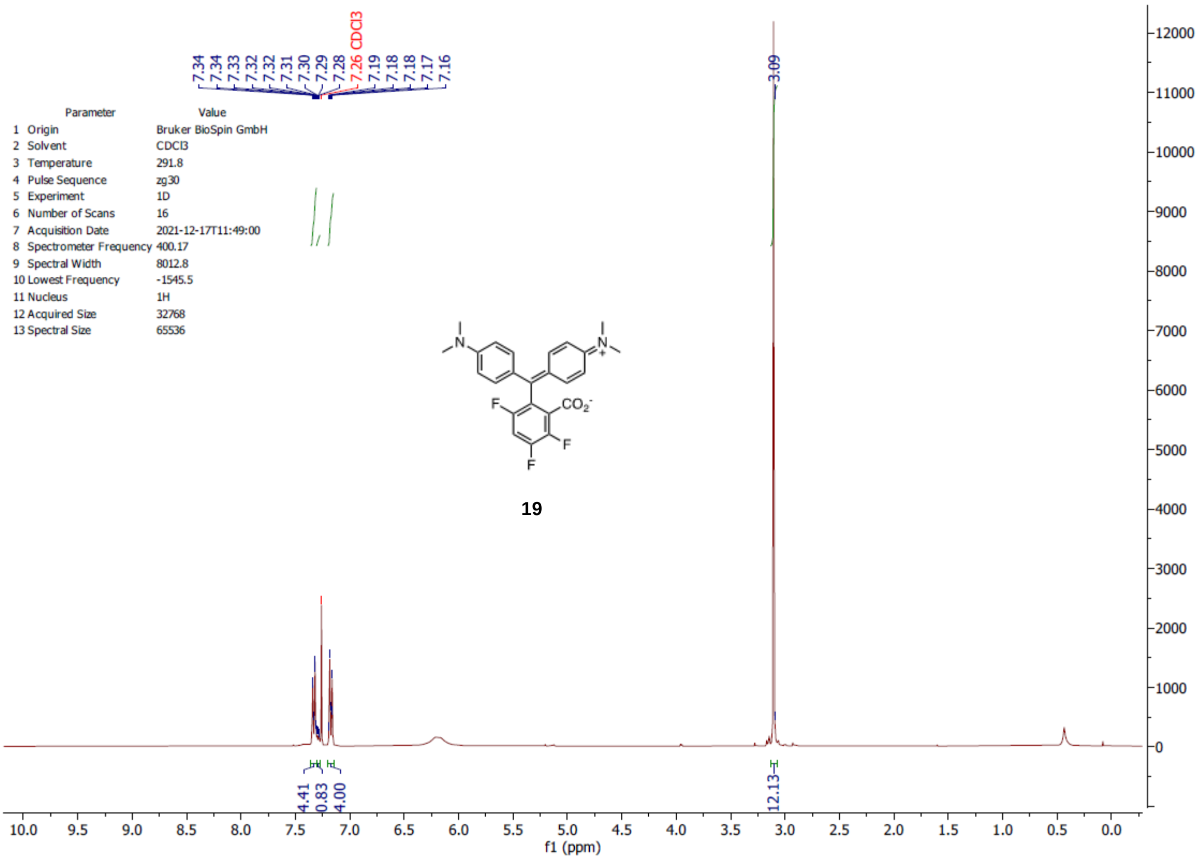

7.34
7.34
7.33
7.32
7.32
7.31
7.30
7.29
7.28
7.26 CDCl3
7.19
7.18
7.18
7.17
7.16
3.09
Parameter Value
1 Origin Bruker BioSpin GmbH
2 Solvent CDCl3
3 Temperature 291.8
4 Pulse Sequence zg30
5 Experiment 1D
6 Number of Scans 16
7 Acquisition Date 2021-12-17T11:49:00
8 Spectrometer Frequency 400.17
9 Spectral Width 8012.8
10 Lowest Frequency -1545.5
11 Nucleus 1H
12 Acquired Size 32768
13 Spectral Size 65536
19
4.41
0.83
4.00
12.13
f1 (ppm)

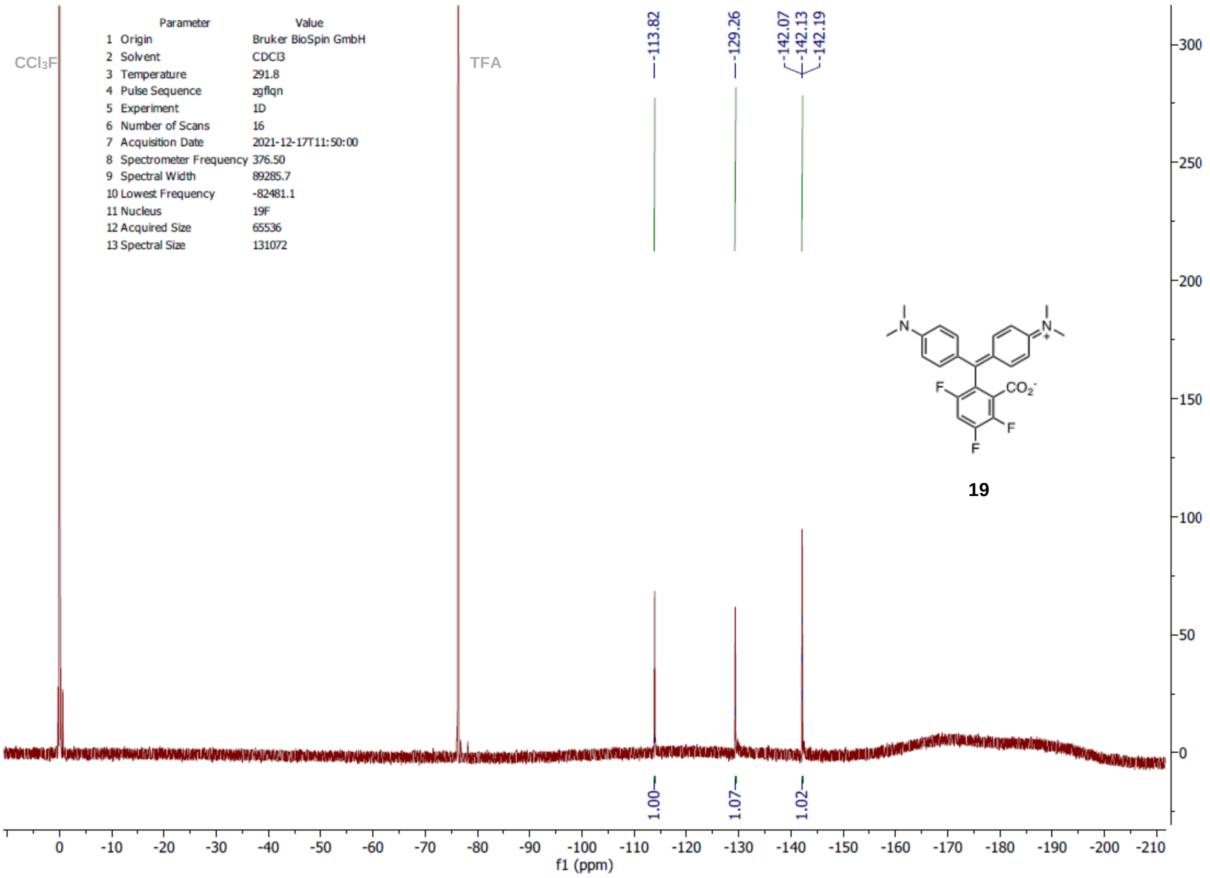

CCl3F
TFA
-113.82
-129.26
-142.07
-142.13
-142.19
Parameter Value
1 Origin Bruker BioSpin GmbH
2 Solvent CDCl3
3 Temperature 291.8
4 Pulse Sequence zgflqn
5 Experiment 1D
6 Number of Scans 16
7 Acquisition Date 2021-12-17T11:50:00
8 Spectrometer Frequency 376.50
9 Spectral Width 89285.7
10 Lowest Frequency -82481.1
11 Nucleus 19F
12 Acquired Size 65536
13 Spectral Size 131072
19
1.00
1.07
1.02
f1 (ppm)

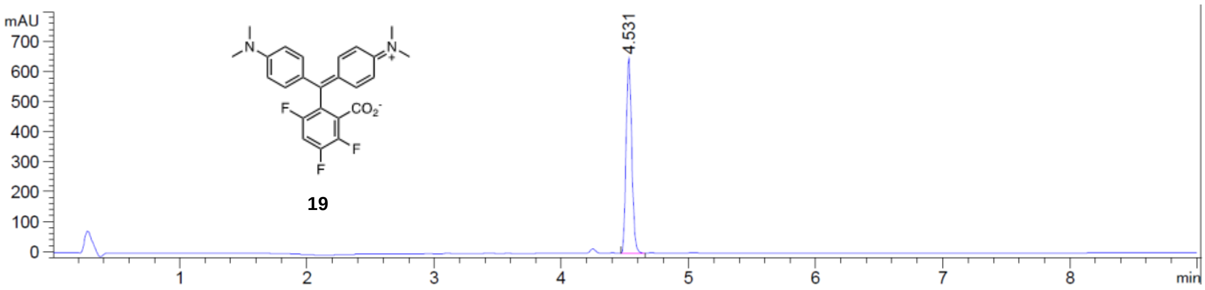
mAU
700
600
500
400
300
200
100
0
4.531
CO2-
F
F
F
19
1
2
3
4
5
6
7
8
min

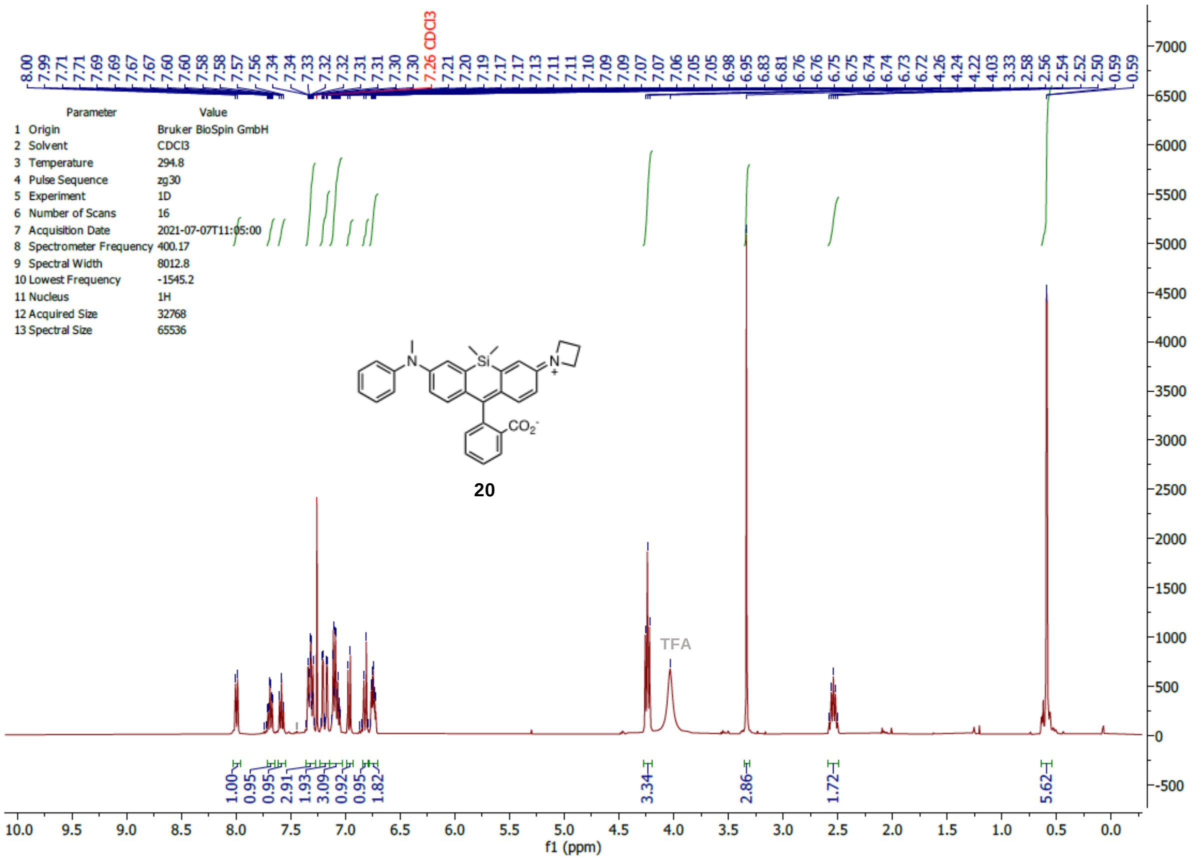
8.00 7.99 7.71 7.71 7.69 7.69 7.67 7.67 7.60 7.60 7.58 7.58 7.57 7.56 7.34 7.34 7.33 7.32 7.32 7.31 7.31 7.30 7.30 7.26 CDCl3 7.21 7.20 7.19 7.17 7.17 7.13 7.11 7.11 7.10 7.09 7.09 7.07 7.07 7.06 7.05 7.05 6.98 6.95 6.83 6.81 6.76 6.76 6.75 6.75 6.74 6.74 6.73 6.72 4.26 4.24 4.22 4.03 3.33 2.58 2.56 2.54 2.52 2.50 0.59 0.59
Parameter Value
1 Origin Bruker BioSpin GmbH
2 Solvent CDCl3
3 Temperature 294.8
4 Pulse Sequence zg30
5 Experiment 1D
6 Number of Scans 16
7 Acquisition Date 2021-07-07T11:05:00
8 Spectrometer Frequency 400.17
9 Spectral Width 8012.8
10 Lowest Frequency -1545.2
11 Nucleus 1H
12 Acquired Size 32768
13 Spectral Size 65536
20
TFA
1.00 0.95 0.95 2.91 1.93 3.09 0.92 0.95 1.82 3.34 2.86 1.72 5.62
f1 (ppm)

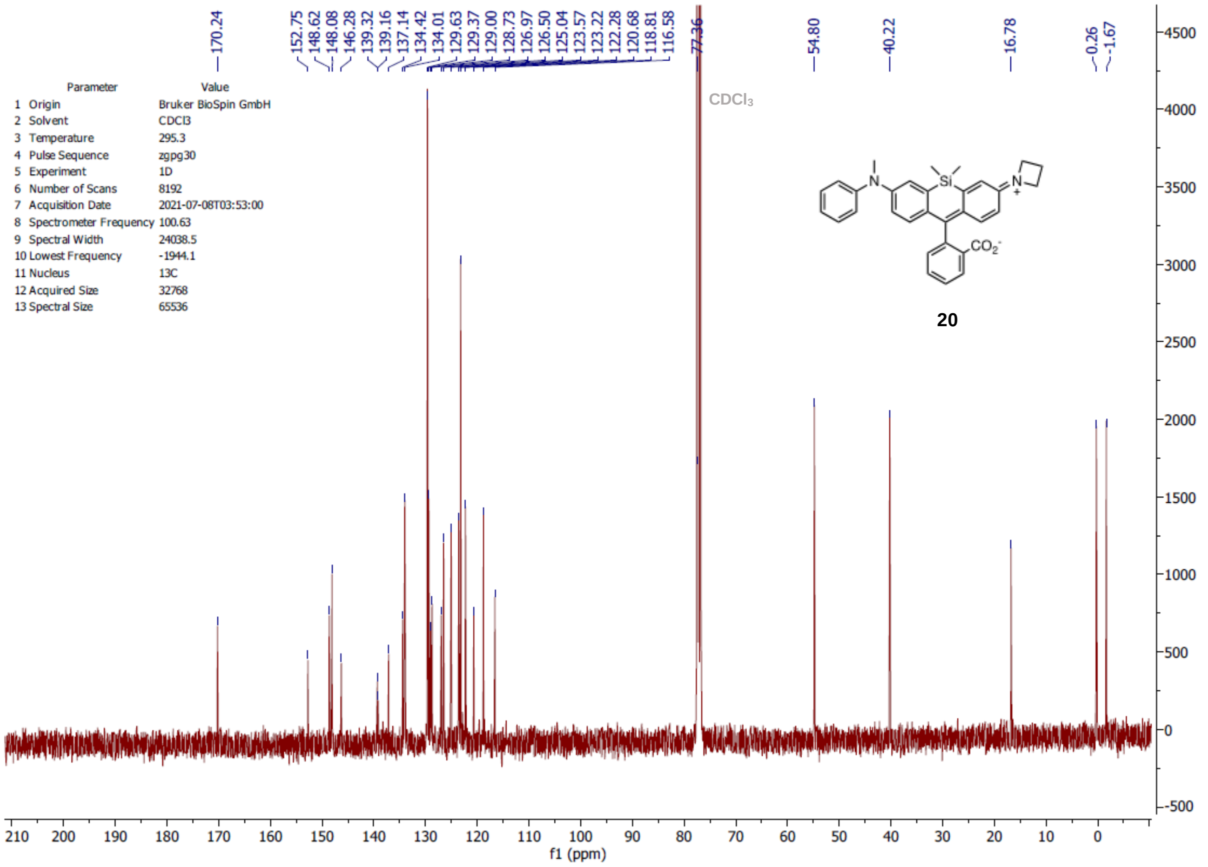
170.24 152.75 148.62 148.08 146.28 139.32 139.16 137.14 134.42 134.01 129.63 129.37 129.00 128.73 126.97 126.50 125.04 123.57 123.22 122.28 120.68 118.81 116.58 77.36 54.80 40.22 16.78 -0.26 -1.67
CDCl3
Parameter Value
1 Origin Bruker BioSpin GmbH
2 Solvent CDCl3
3 Temperature 295.3
4 Pulse Sequence zgpg30
5 Experiment 1D
6 Number of Scans 8192
7 Acquisition Date 2021-07-08T03:53:00
8 Spectrometer Frequency 100.63
9 Spectral Width 24038.5
10 Lowest Frequency -1944.1
11 Nucleus 13C
12 Acquired Size 32768
13 Spectral Size 65536
20
f1 (ppm)

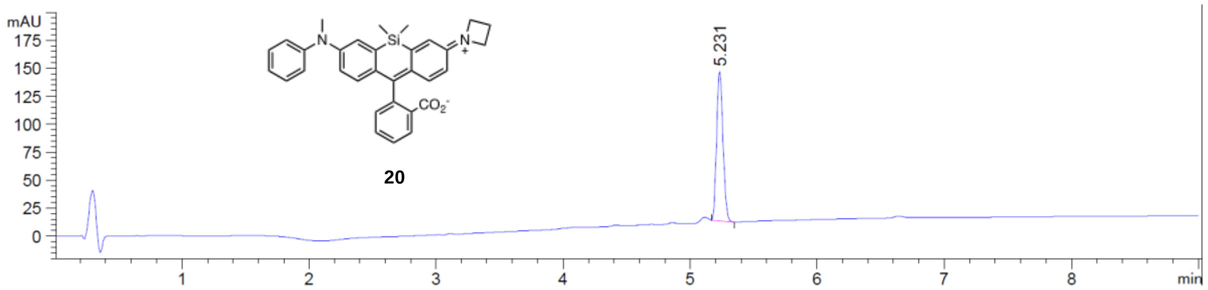
mAU
175
150
125
100
75
50
25
0
5.231
20
1
2
3
4
5
6
7
8
min

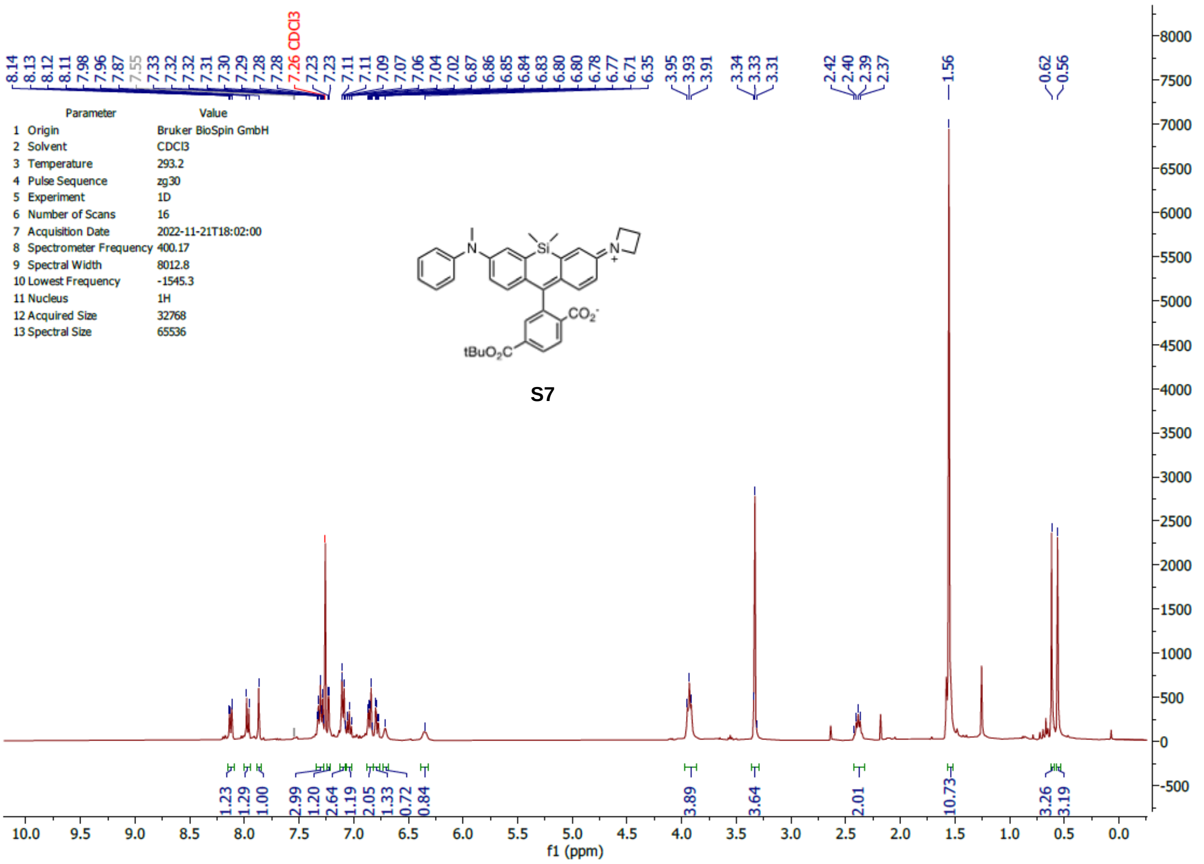
Parameter Value
1 Origin Bruker BioSpin GmbH
2 Solvent CDCl3
3 Temperature 293.2
4 Pulse Sequence zg30
5 Experiment 1D
6 Number of Scans 16
7 Acquisition Date 2022-11-21T18:02:00
8 Spectrometer Frequency 400.17
9 Spectral Width 8012.8
10 Lowest Frequency -1545.3
11 Nucleus 1H
12 Acquired Size 32768
13 Spectral Size 65536
7.26 CDCl3
tBuO2C
CO2-
S7
f1 (ppm)

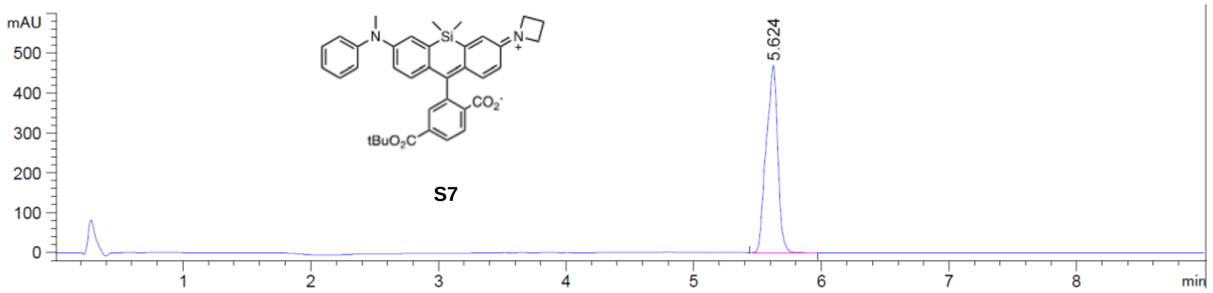
mAU
5.624
tBuO2C
CO2-
S7
min

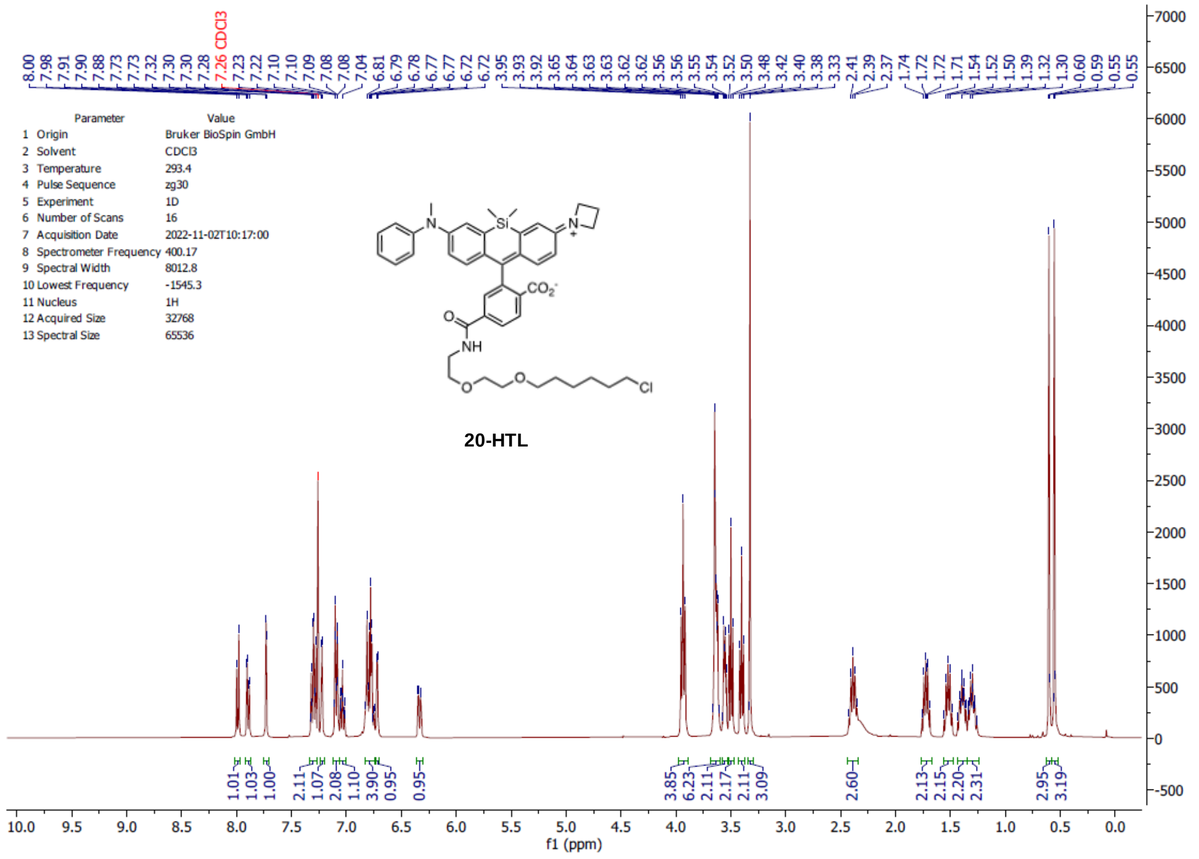
Parameter Value
1 Origin Bruker BioSpin GmbH
2 Solvent CDCl3
3 Temperature 293.4
4 Pulse Sequence zg30
5 Experiment 1D
6 Number of Scans 16
7 Acquisition Date 2022-11-02T10:17:00
8 Spectrometer Frequency 400.17
9 Spectral Width 8012.8
10 Lowest Frequency -1545.3
11 Nucleus 1H
12 Acquired Size 32768
13 Spectral Size 65536
20-HTL
f1 (ppm)

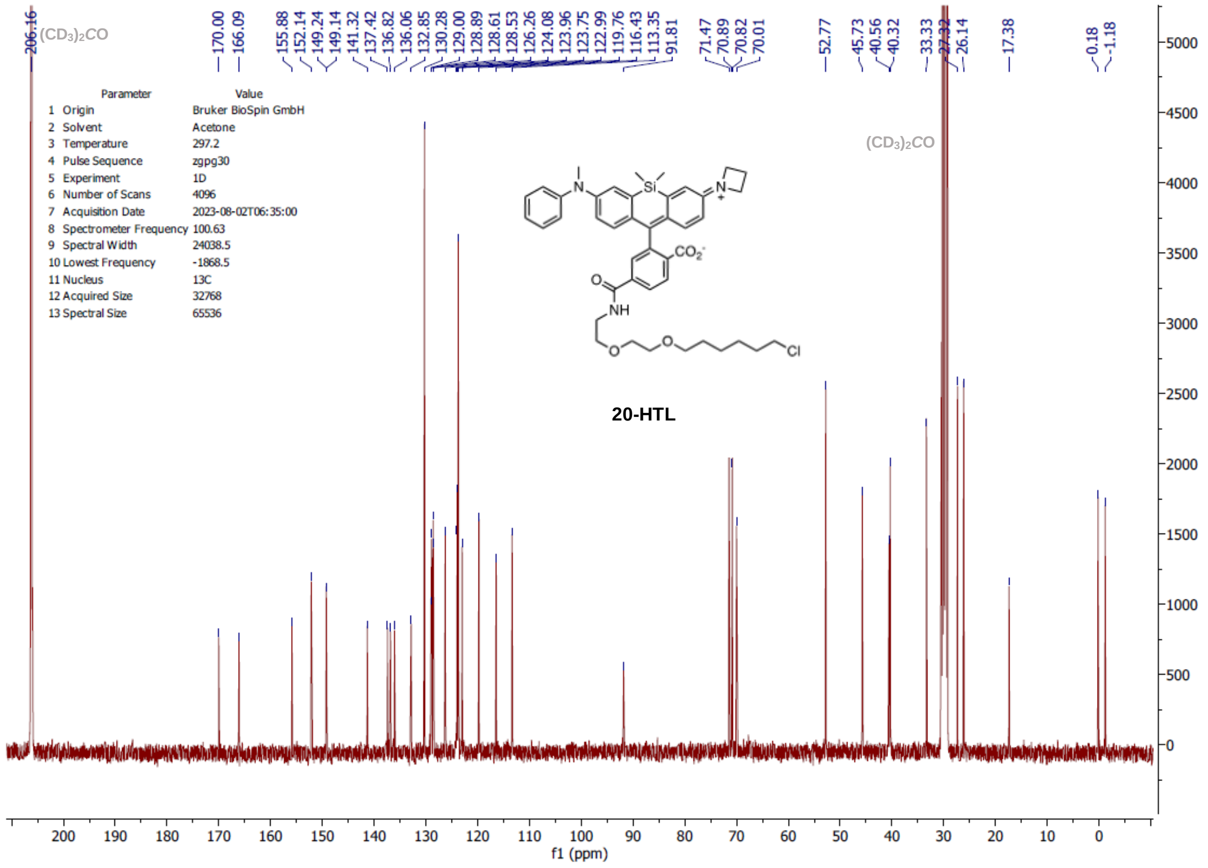
Parameter Value
1 Origin Bruker BioSpin GmbH
2 Solvent Acetone
3 Temperature 297.2
4 Pulse Sequence zgpg30
5 Experiment 1D
6 Number of Scans 4096
7 Acquisition Date 2023-08-02T06:35:00
8 Spectrometer Frequency 100.63
9 Spectral Width 24038.5
10 Lowest Frequency -1868.5
11 Nucleus 13C
12 Acquired Size 32768
13 Spectral Size 65536
(CD3)2CO
20-HTL
f1 (ppm)

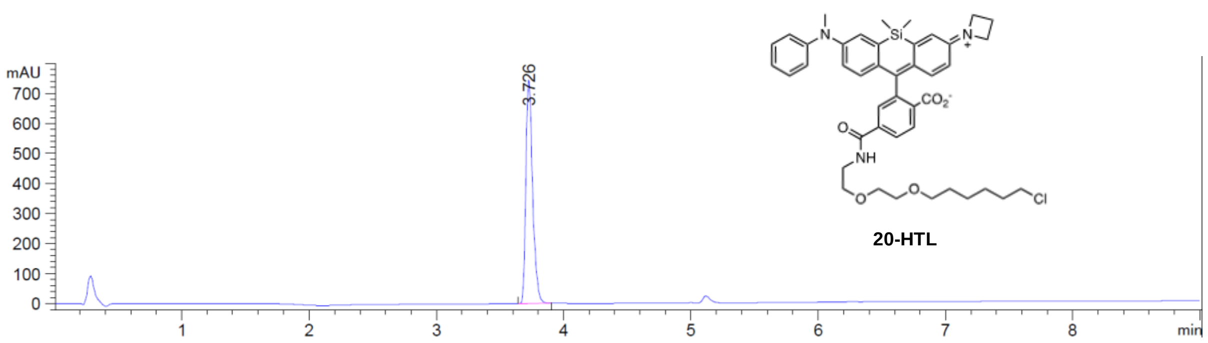

mAU
700
600
500
400
300
200
100
0
3.726
1
2
3
4
5
6
7
8
min
20-HTL

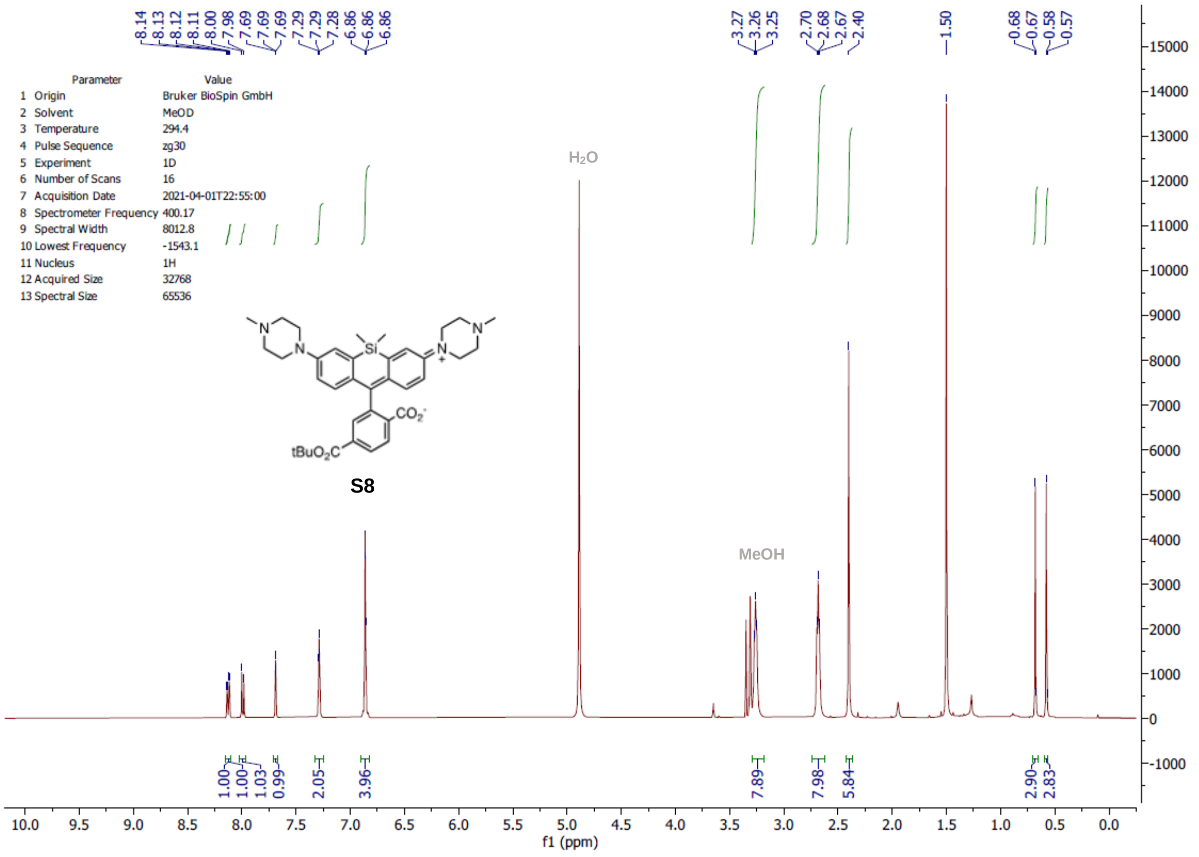

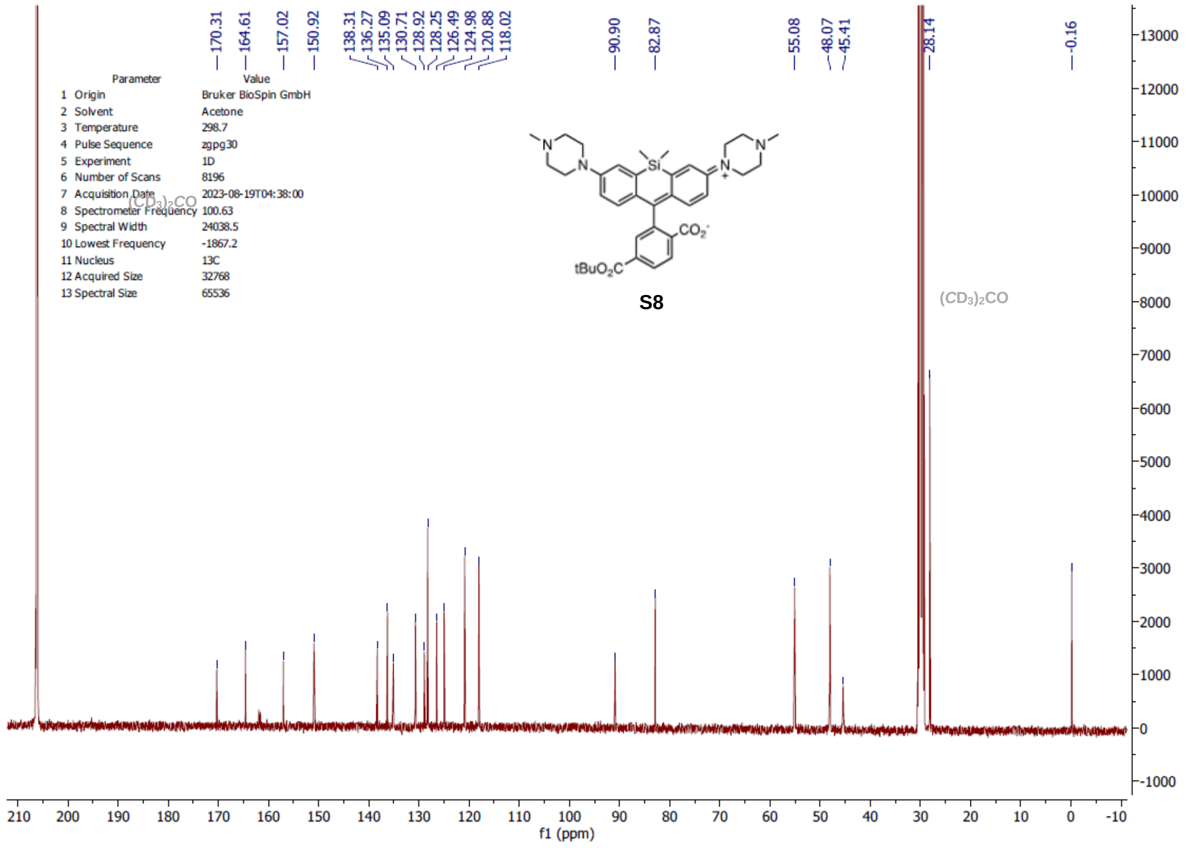

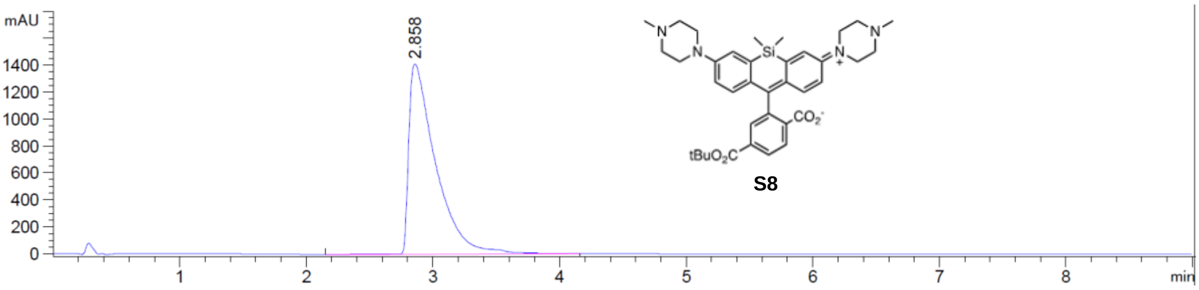
mAU
1400
1200
1000
800
600
400
200
0
2.858
1
2
3
4
5
6
7
8
min
tBuO2C
CO2-
Si
N
S8

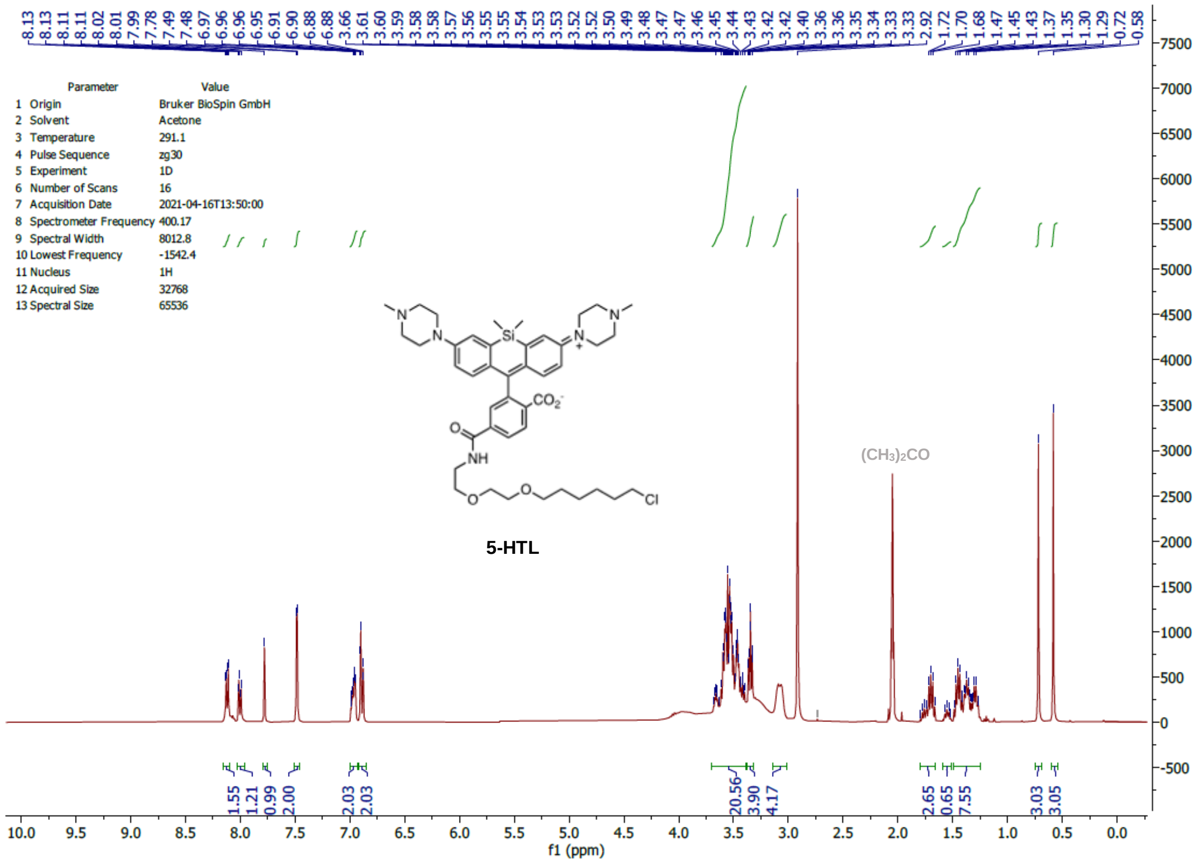
8.13 8.13 8.11 8.11 8.02 8.01 7.99 7.78 7.49 7.48 6.97 6.96 6.96 6.95 6.91 6.90 6.88 6.88 3.66 3.61 3.60 3.59 3.58 3.58 3.57 3.56 3.55 3.55 3.54 3.53 3.53 3.52 3.52 3.50 3.49 3.48 3.47 3.47 3.46 3.45 3.44 3.43 3.42 3.42 3.40 3.36 3.36 3.35 3.34 3.33 3.33 2.92 1.72 1.70 1.68 1.47 1.45 1.43 1.37 1.35 1.30 1.29 0.72 0.58
Parameter Value
1 Origin Bruker BioSpin GmbH
2 Solvent Acetone
3 Temperature 291.1
4 Pulse Sequence zg30
5 Experiment 1D
6 Number of Scans 16
7 Acquisition Date 2021-04-16T13:50:00
8 Spectrometer Frequency 400.17
9 Spectral Width 8012.8
10 Lowest Frequency -1542.4
11 Nucleus 1H
12 Acquired Size 32768
13 Spectral Size 65536
5-HTL
(CH3)2CO
1.55 1.21 0.99 2.00 2.03 2.03 20.56 3.90 4.17 2.65 0.65 7.55 3.03 3.05
f1 (ppm)

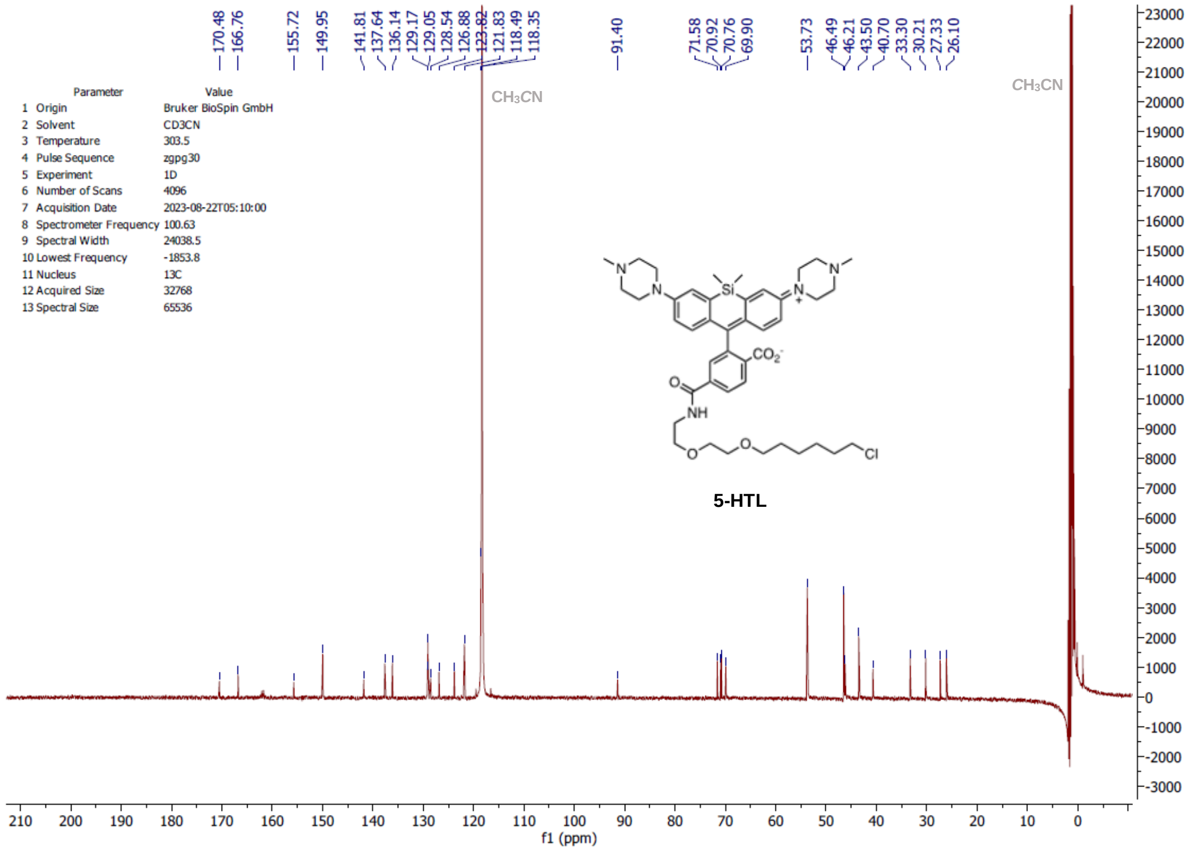
170.48 166.76 155.72 149.95 141.81 137.64 136.14 129.17 129.05 128.54 126.88 122.82 121.83 118.49 118.35 91.40 71.58 70.92 70.76 69.90 53.73 46.49 46.21 43.50 40.70 33.30 30.21 27.33 26.10
Parameter Value
1 Origin Bruker BioSpin GmbH
2 Solvent CD3CN
3 Temperature 303.5
4 Pulse Sequence zgpg30
5 Experiment 1D
6 Number of Scans 4096
7 Acquisition Date 2023-08-22T05:10:00
8 Spectrometer Frequency 100.63
9 Spectral Width 24038.5
10 Lowest Frequency -1853.8
11 Nucleus 13C
12 Acquired Size 32768
13 Spectral Size 65536
CH3CN
CH3CN
5-HTL
f1 (ppm)

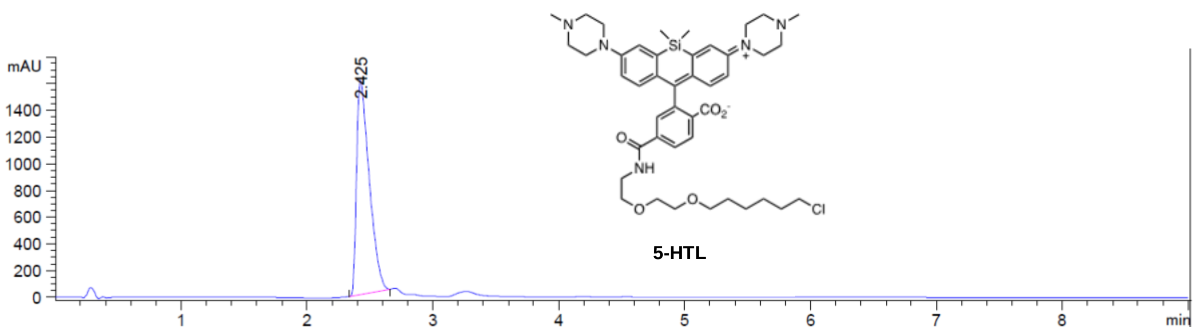

mAU
1400
1200
1000
800
600
400
200
0
2.425
1
2
3
4
5
6
7
8
min
NH
CO2
Cl
5-HTL

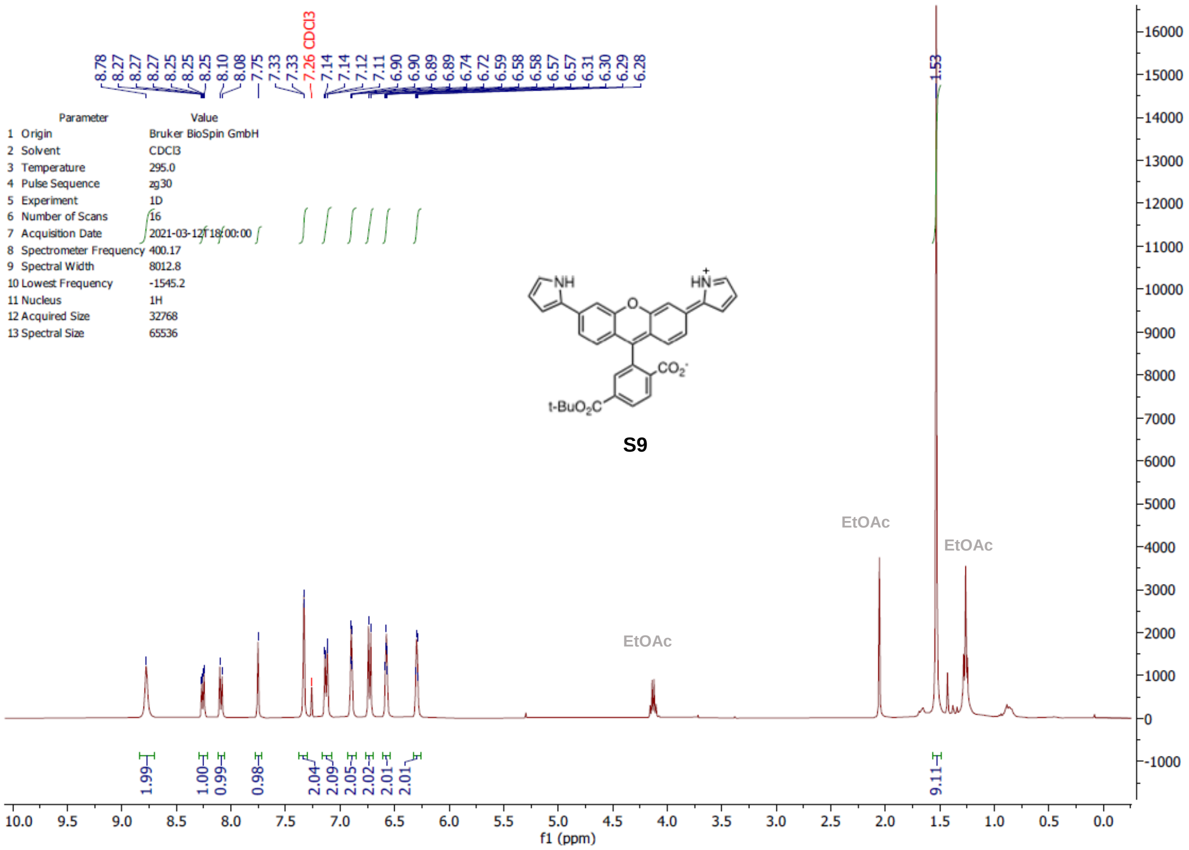
8.78
8.27
8.27
8.27
8.25
8.25
8.25
8.10
8.08
7.75
7.33
7.33
7.26 CDCl3
7.14
7.14
7.12
7.11
6.90
6.90
6.89
6.89
6.74
6.72
6.59
6.58
6.58
6.57
6.57
6.31
6.30
6.29
6.28
1.53
Parameter Value
1 Origin Bruker BioSpin GmbH
2 Solvent CDCl3
3 Temperature 295.0
4 Pulse Sequence zg30
5 Experiment 1D
6 Number of Scans 16
7 Acquisition Date 2021-03-12T18:00:00
8 Spectrometer Frequency 400.17
9 Spectral Width 8012.8
10 Lowest Frequency -1545.2
11 Nucleus 1H
12 Acquired Size 32768
13 Spectral Size 65536
NH
HN+
O
CO2-
t-BuO2C
S9
EtOAc
EtOAc
EtOAc
1.99
1.00
0.99
0.98
2.04
2.09
2.05
2.02
2.01
2.01
9.11
f1 (ppm)

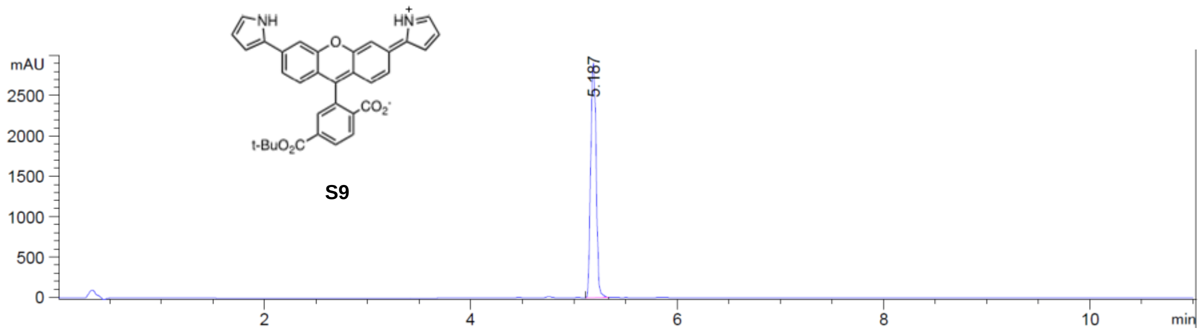
mAU
NH
HN+
O
CO2-
t-BuO2C
S9
5.187
min

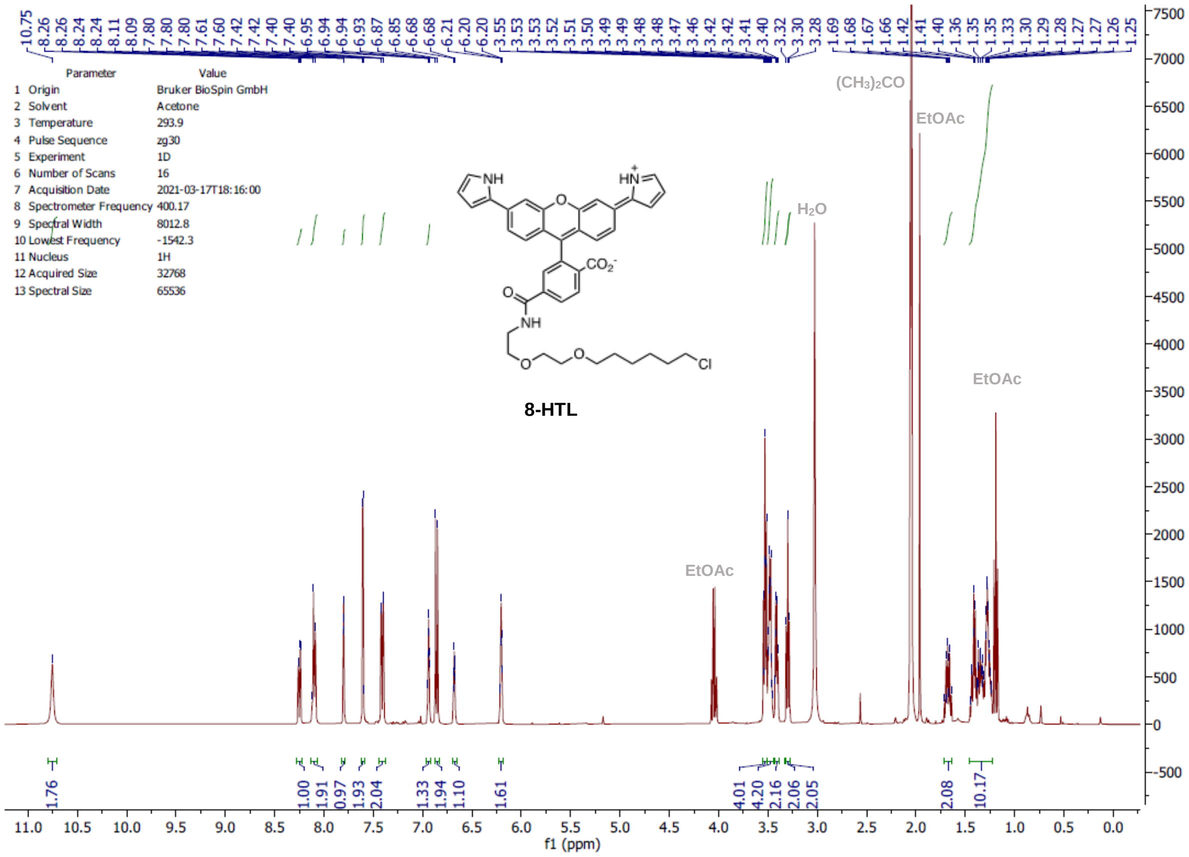
10.75 8.26 8.26 8.24 8.24 8.11 8.09 7.80 7.80 7.80 7.61 7.60 7.42 7.42 7.40 6.95 6.94 6.94 6.93 6.87 6.85 6.68 6.68 6.21 6.20 6.20 3.55 3.53 3.53 3.52 3.51 3.50 3.49 3.49 3.48 3.48 3.47 3.46 3.42 3.42 3.41 3.40 3.32 3.30 3.28 1.69 1.68 1.67 1.66 1.42 1.41 1.40 1.36 1.35 1.35 1.33 1.30 1.29 1.28 1.27 1.27 1.26 1.25
Parameter Value
1 Origin Bruker BioSpin GmbH
2 Solvent Acetone
3 Temperature 293.9
4 Pulse Sequence zg30
5 Experiment 1D
6 Number of Scans 16
7 Acquisition Date 2021-03-17T18:16:00
8 Spectrometer Frequency 400.17
9 Spectral Width 8012.8
10 Lowest Frequency -1542.3
11 Nucleus 1H
12 Acquired Size 32768
13 Spectral Size 65536
(CH3)2CO
EtOAc
H2O
EtOAc
EtOAc
NH
HN+
O
CO2-
NH
Cl
8-HTL
1.76 1.00 1.91 0.97 1.93 2.04 1.33 1.94 1.10 1.61 4.01 4.20 2.16 2.06 2.05 2.08 10.17
f1 (ppm)

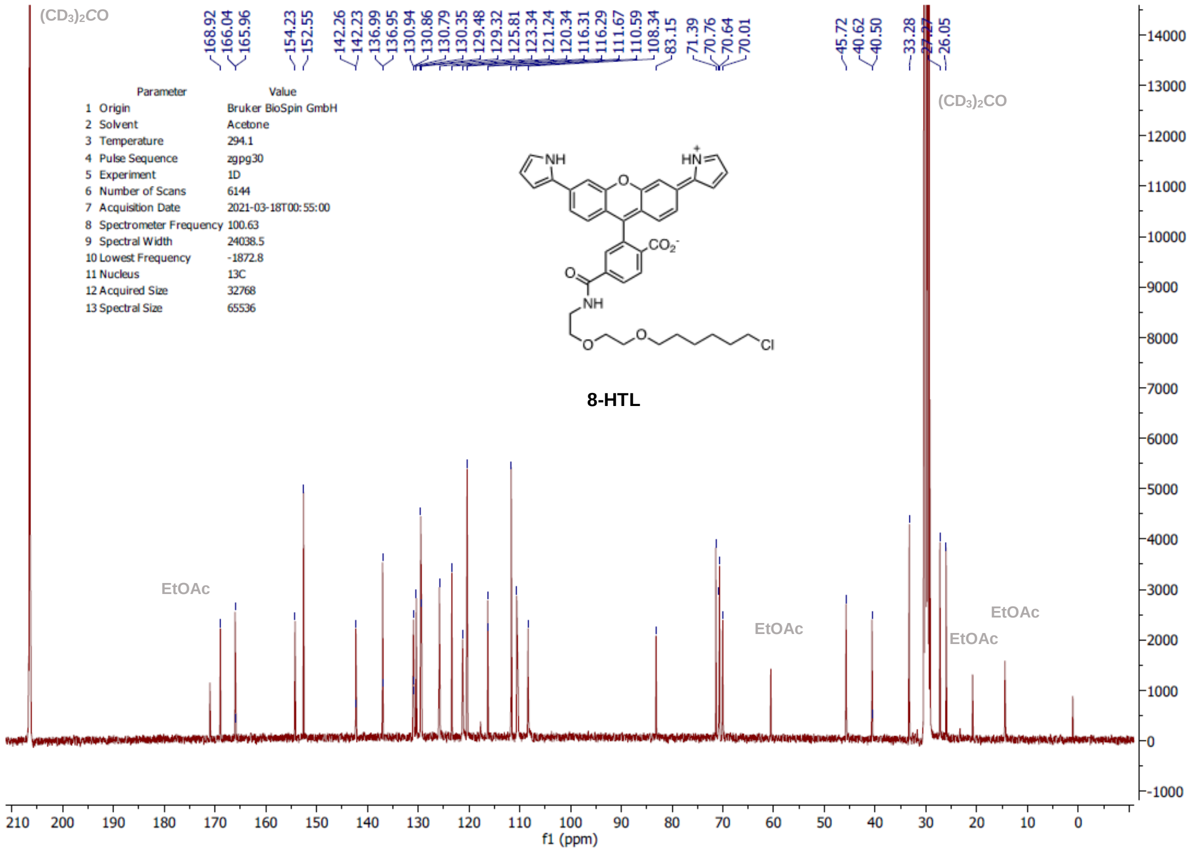
(CD3)2CO
168.92 166.04 165.96 154.23 152.55 142.26 142.23 136.99 136.95 130.94 130.86 130.79 130.35 129.48 129.32 125.81 123.34 121.24 120.34 116.31 116.29 111.67 110.59 108.34 83.15 71.39 70.76 70.64 70.01 45.72 40.62 40.50 33.28 26.05
Parameter Value
1 Origin Bruker BioSpin GmbH
2 Solvent Acetone
3 Temperature 294.1
4 Pulse Sequence zgpg30
5 Experiment 1D
6 Number of Scans 6144
7 Acquisition Date 2021-03-18T00:55:00
8 Spectrometer Frequency 100.63
9 Spectral Width 24038.5
10 Lowest Frequency -1872.8
11 Nucleus 13C
12 Acquired Size 32768
13 Spectral Size 65536
(CD3)2CO
NH
HN+
CO2-
NH
Cl
8-HTL
EtOAc
EtOAc
EtOAc
EtOAc
f1 (ppm)

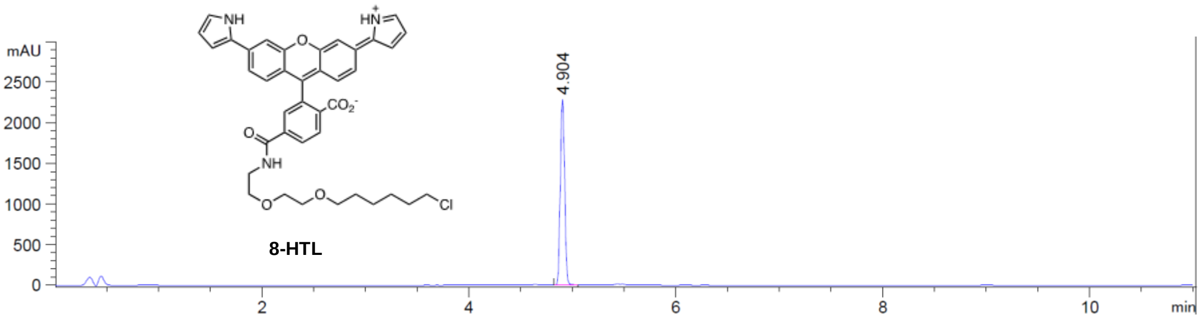
mAU
2500
2000
1500
1000
500
0
2
4
6
8
10
min
4.904
NH
HN
O
CO2-
O
NH
O
O
Cl
8-HTL

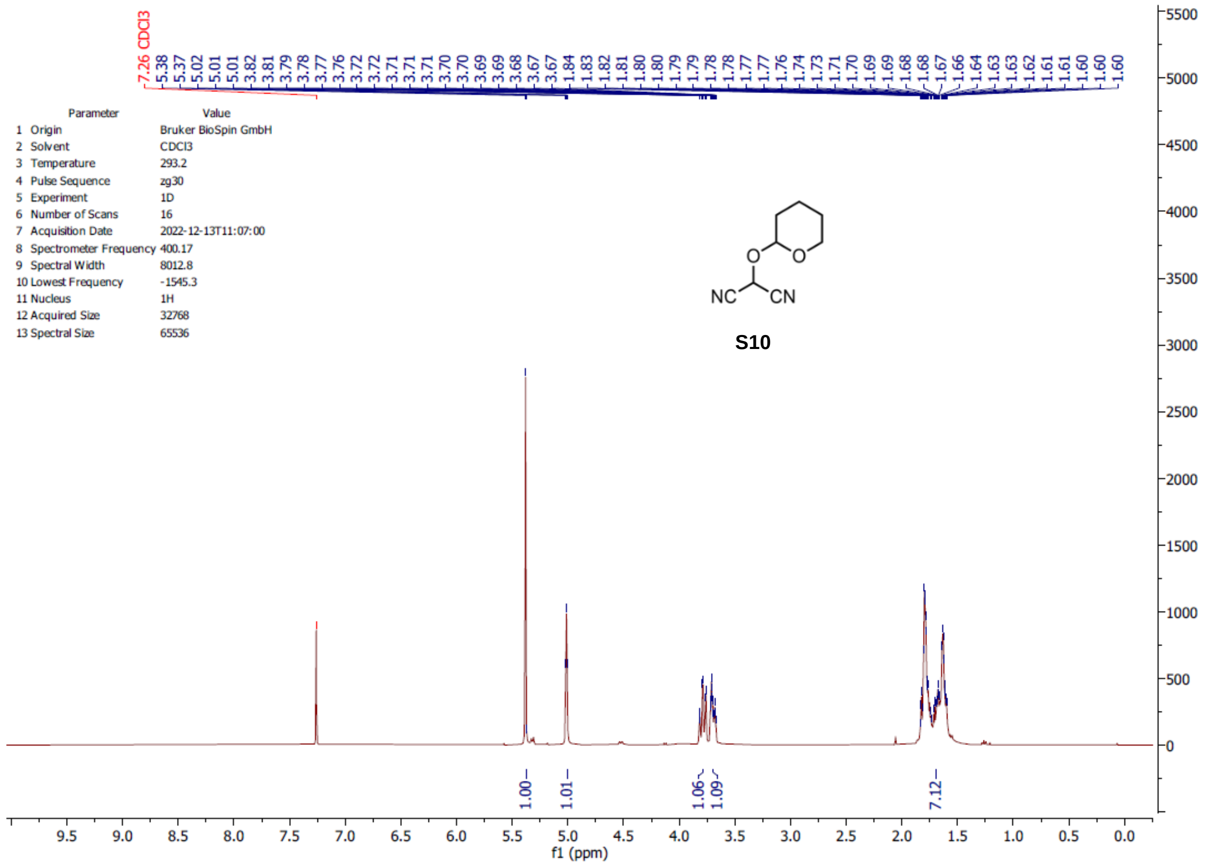
7.26 CDCl3
5.38
5.37
5.02
5.01
3.82
3.81
3.79
3.78
3.77
3.76
3.72
3.72
3.71
3.71
3.71
3.70
3.70
3.69
3.69
3.68
3.67
3.67
1.84
1.83
1.82
1.81
1.80
1.80
1.79
1.79
1.78
1.78
1.77
1.77
1.76
1.74
1.73
1.71
1.70
1.69
1.69
1.68
1.68
1.67
1.66
1.64
1.63
1.63
1.62
1.61
1.61
1.60
1.60
1.60
Parameter Value
1 Origin Bruker BioSpin GmbH
2 Solvent CDCl3
3 Temperature 293.2
4 Pulse Sequence zg30
5 Experiment 1D
6 Number of Scans 16
7 Acquisition Date 2022-12-13T11:07:00
8 Spectrometer Frequency 400.17
9 Spectral Width 8012.8
10 Lowest Frequency -1545.3
11 Nucleus 1H
12 Acquired Size 32768
13 Spectral Size 65536
O
O
NC
CN
S10
1.00
1.01
1.06
1.09
7.12
f1 (ppm)

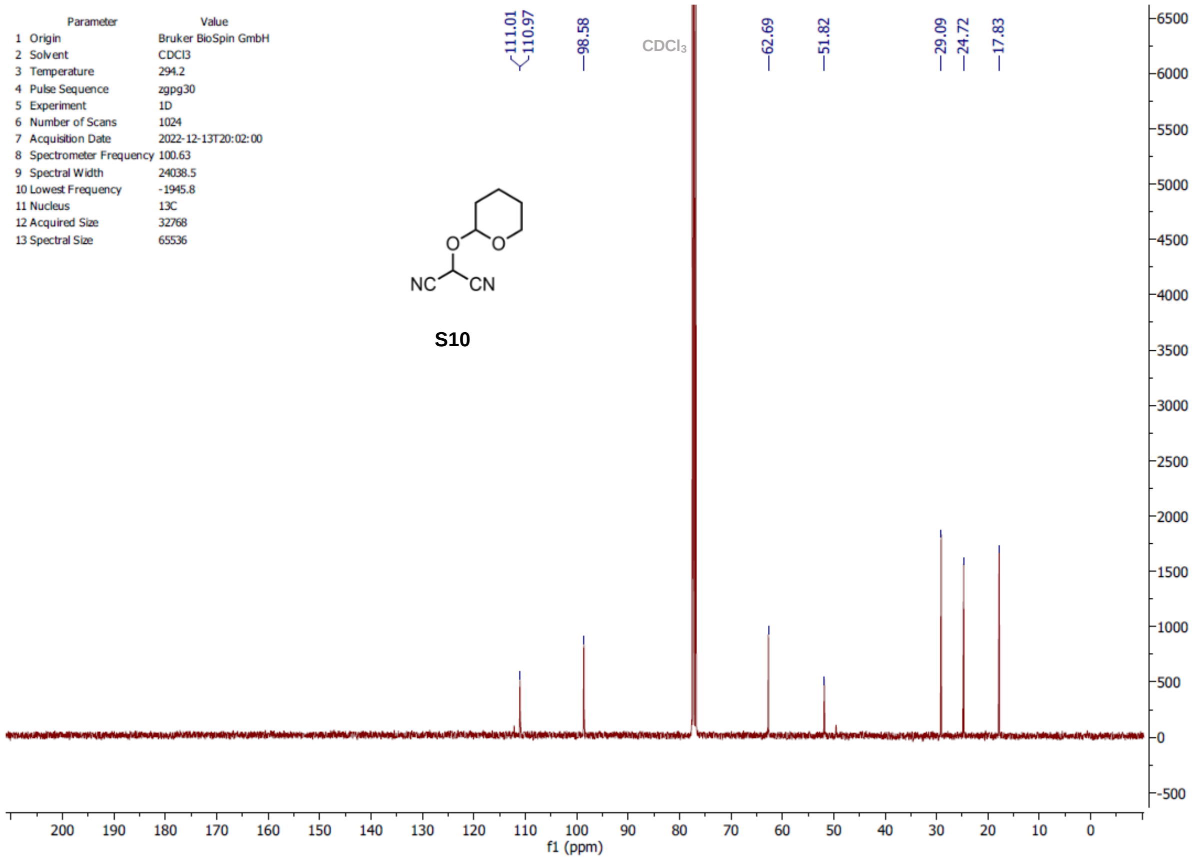
111.01
110.97
98.58
CDCl3
62.69
51.82
29.09
24.72
17.83
Parameter Value
1 Origin Bruker BioSpin GmbH
2 Solvent CDCl3
3 Temperature 294.2
4 Pulse Sequence zgpg30
5 Experiment 1D
6 Number of Scans 1024
7 Acquisition Date 2022-12-13T20:02:00
8 Spectrometer Frequency 100.63
9 Spectral Width 24038.5
10 Lowest Frequency -1945.8
11 Nucleus 13C
12 Acquired Size 32768
13 Spectral Size 65536
O
O
NC
CN
S10
f1 (ppm)

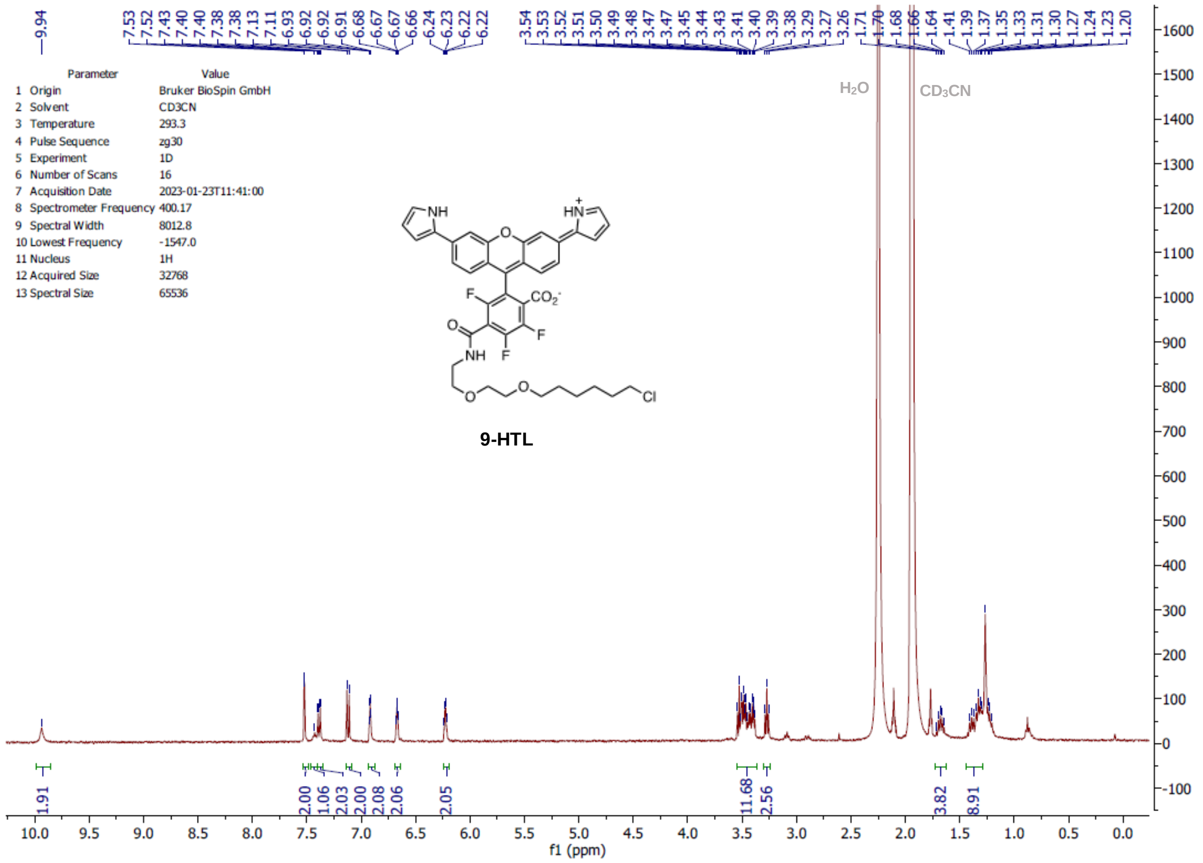
9.94
7.53 7.52 7.43 7.40 7.40 7.38 7.38 7.13 7.11 6.93 6.92 6.92 6.91 6.68 6.67 6.67 6.66 6.24 6.23 6.22 6.22
3.54 3.53 3.52 3.51 3.50 3.49 3.48 3.47 3.47 3.45 3.44 3.43 3.41 3.40 3.39 3.38 3.29 3.27 3.26 1.71 1.70 1.68 1.66 1.64 1.41 1.39 1.37 1.35 1.33 1.31 1.30 1.27 1.24 1.23 1.20
Parameter Value
1 Origin Bruker BioSpin GmbH
2 Solvent CD3CN
3 Temperature 293.3
4 Pulse Sequence zg30
5 Experiment 1D
6 Number of Scans 16
7 Acquisition Date 2023-01-23T11:41:00
8 Spectrometer Frequency 400.17
9 Spectral Width 8012.8
10 Lowest Frequency -1547.0
11 Nucleus 1H
12 Acquired Size 32768
13 Spectral Size 65536
H2O
CD3CN
9-HTL
1.91 2.00 1.06 2.03 2.00 2.08 2.06 2.05 11.68 2.56 3.82 8.91
f1 (ppm)

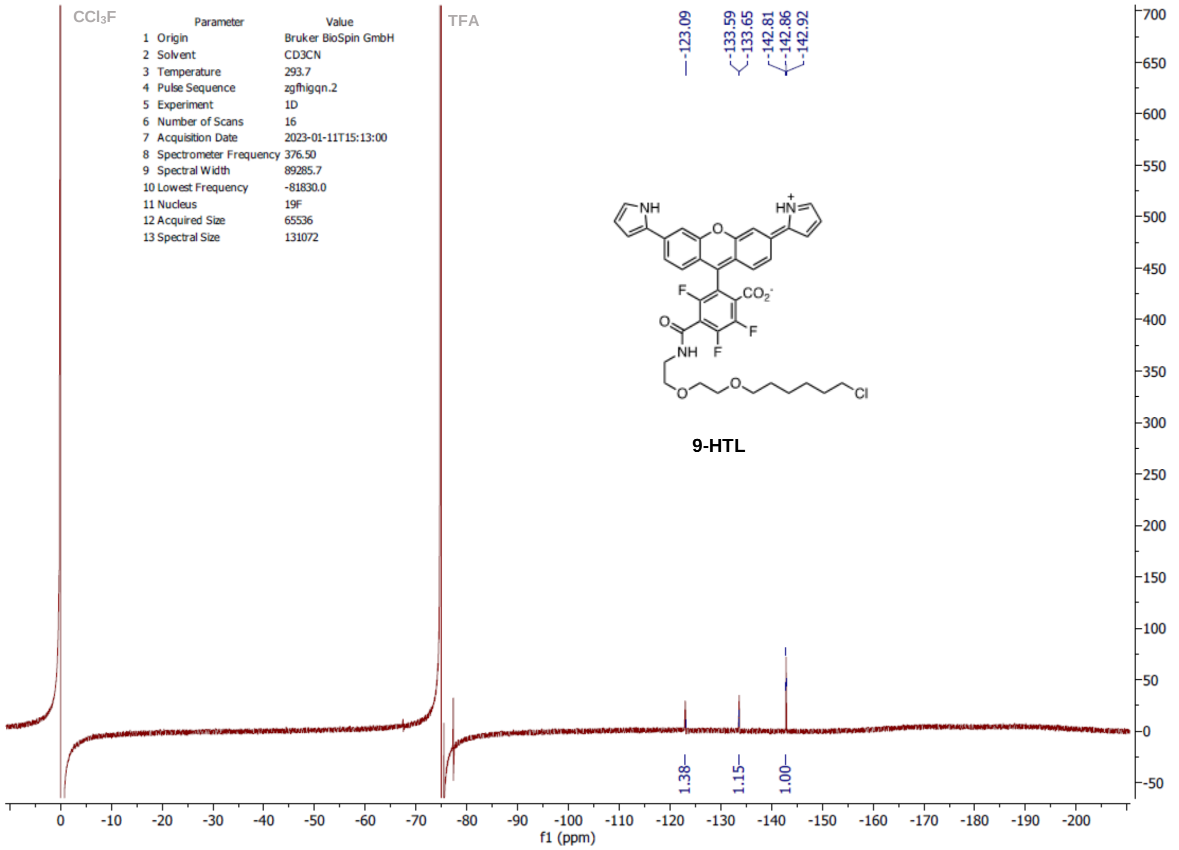
CCl3F
TFA
-123.09
-133.59 -133.65
-142.81 -142.86 -142.92
Parameter Value
1 Origin Bruker BioSpin GmbH
2 Solvent CD3CN
3 Temperature 293.7
4 Pulse Sequence zgfhigqn.2
5 Experiment 1D
6 Number of Scans 16
7 Acquisition Date 2023-01-11T15:13:00
8 Spectrometer Frequency 376.50
9 Spectral Width 89285.7
10 Lowest Frequency -81830.0
11 Nucleus 19F
12 Acquired Size 65536
13 Spectral Size 131072
9-HTL
1.38 1.15 1.00
f1 (ppm)

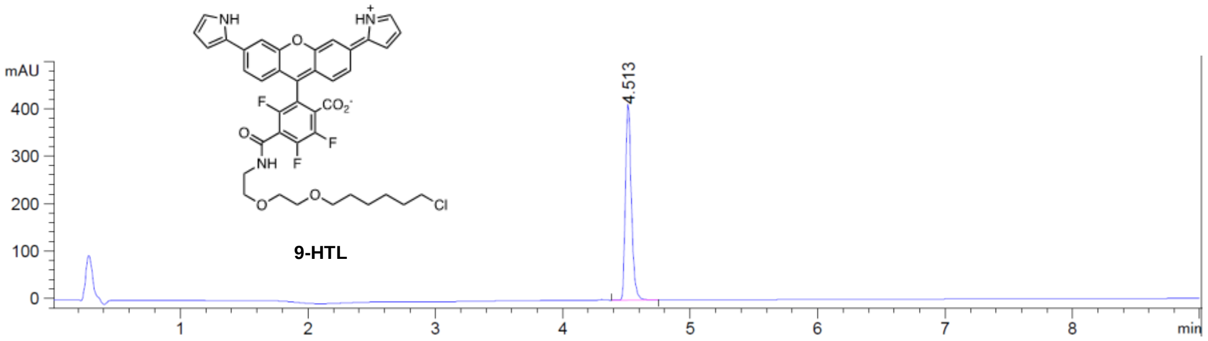

9-HTL

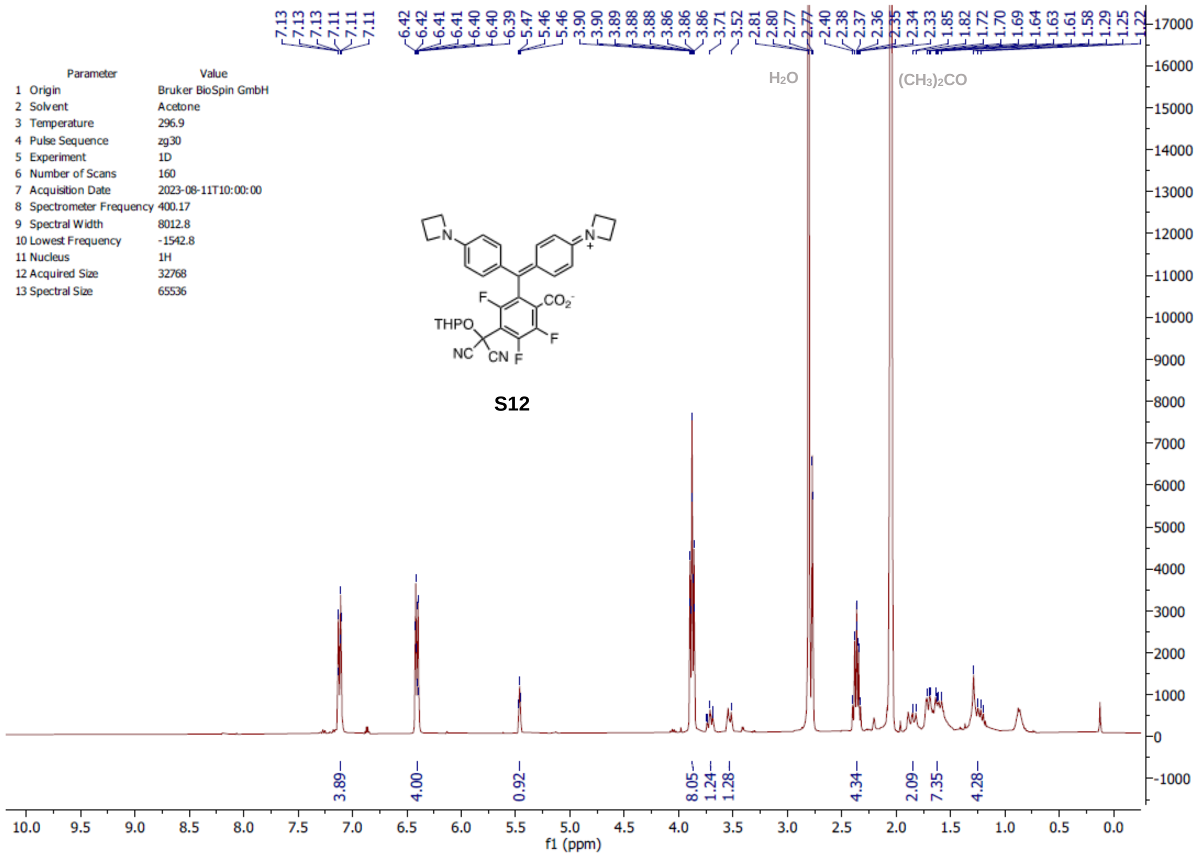

Parameter Value
1 Origin Bruker BioSpin GmbH
2 Solvent Acetone
3 Temperature 296.9
4 Pulse Sequence zg30
5 Experiment 1D
6 Number of Scans 160
7 Acquisition Date 2023-08-11T10:00:00
8 Spectrometer Frequency 400.17
9 Spectral Width 8012.8
10 Lowest Frequency -1542.8
11 Nucleus 1H
12 Acquired Size 32768
13 Spectral Size 65536
S12
H2O
(CH3)2CO
f1 (ppm)

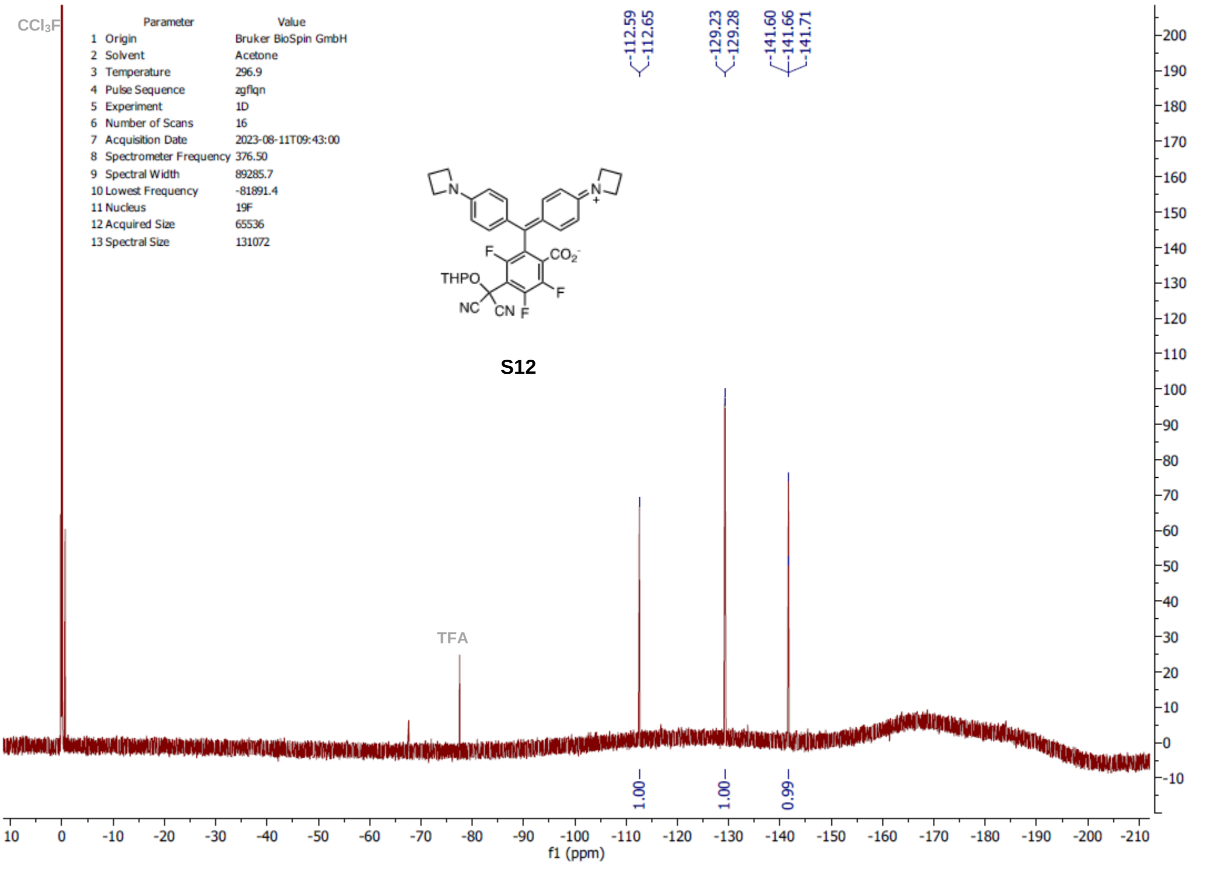

CCl3F
Parameter Value
1 Origin Bruker BioSpin GmbH
2 Solvent Acetone
3 Temperature 296.9
4 Pulse Sequence zgflqn
5 Experiment 1D
6 Number of Scans 16
7 Acquisition Date 2023-08-11T09:43:00
8 Spectrometer Frequency 376.50
9 Spectral Width 89285.7
10 Lowest Frequency -81891.4
11 Nucleus 19F
12 Acquired Size 65536
13 Spectral Size 131072
-112.59
-112.65
-129.23
-129.28
-141.60
-141.66
-141.71
S12
TFA
f1 (ppm)

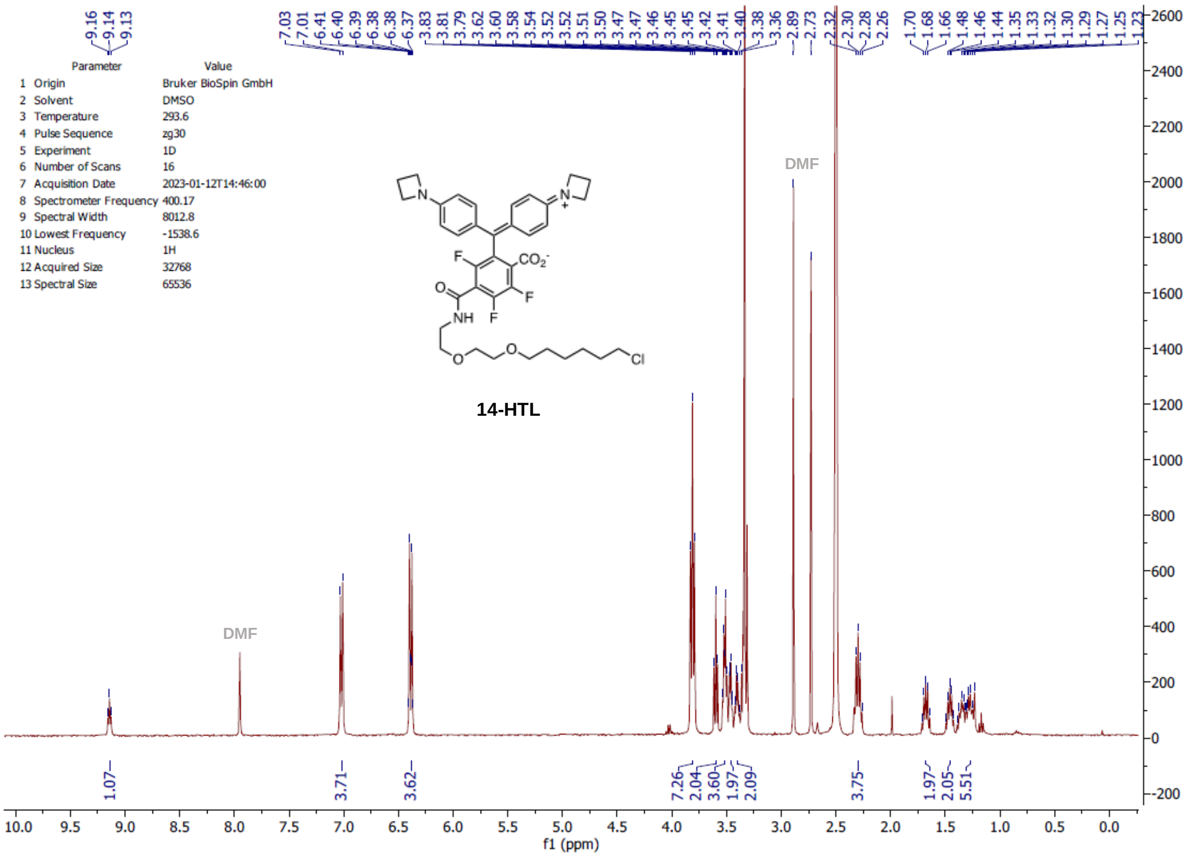
Parameter Value
1 Origin Bruker BioSpin GmbH
2 Solvent DMSO
3 Temperature 293.6
4 Pulse Sequence zg30
5 Experiment 1D
6 Number of Scans 16
7 Acquisition Date 2023-01-12T14:46:00
8 Spectrometer Frequency 400.17
9 Spectral Width 8012.8
10 Lowest Frequency -1538.6
11 Nucleus 1H
12 Acquired Size 32768
13 Spectral Size 65536
DMF
DMF
14-HTL
1.07
3.71
3.62
7.26
2.04
3.60
1.97
2.09
3.75
1.97
2.05
5.51
f1 (ppm)

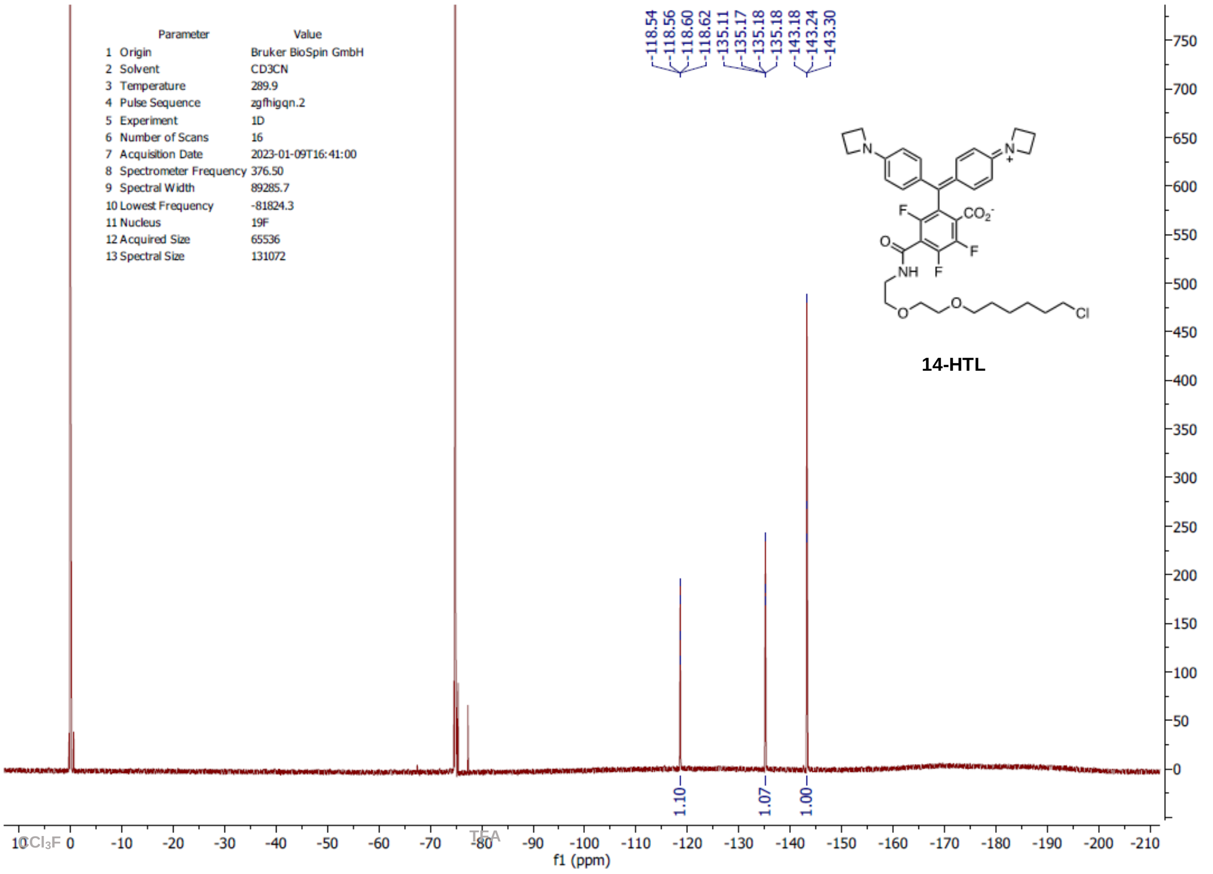
Parameter Value
1 Origin Bruker BioSpin GmbH
2 Solvent CD3CN
3 Temperature 289.9
4 Pulse Sequence zgfhigqn.2
5 Experiment 1D
6 Number of Scans 16
7 Acquisition Date 2023-01-09T16:41:00
8 Spectrometer Frequency 376.50
9 Spectral Width 89285.7
10 Lowest Frequency -81824.3
11 Nucleus 19F
12 Acquired Size 65536
13 Spectral Size 131072
14-HTL
1.10
1.07
1.00
CCl3F
TFA
f1 (ppm)

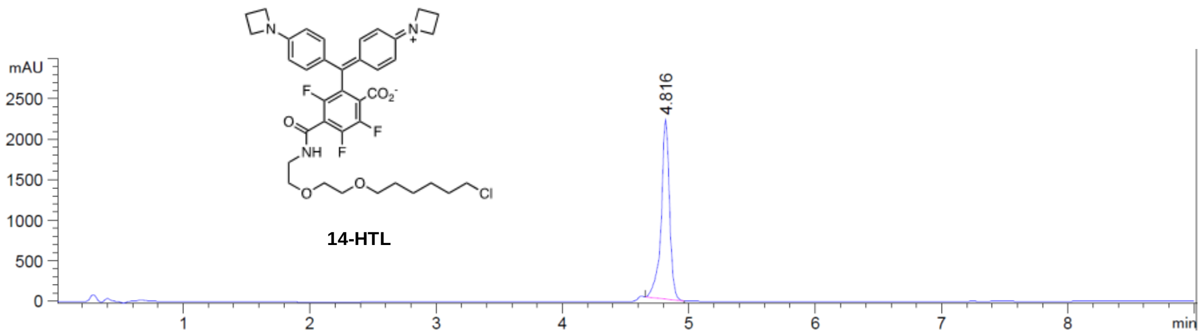

**14-HTL**

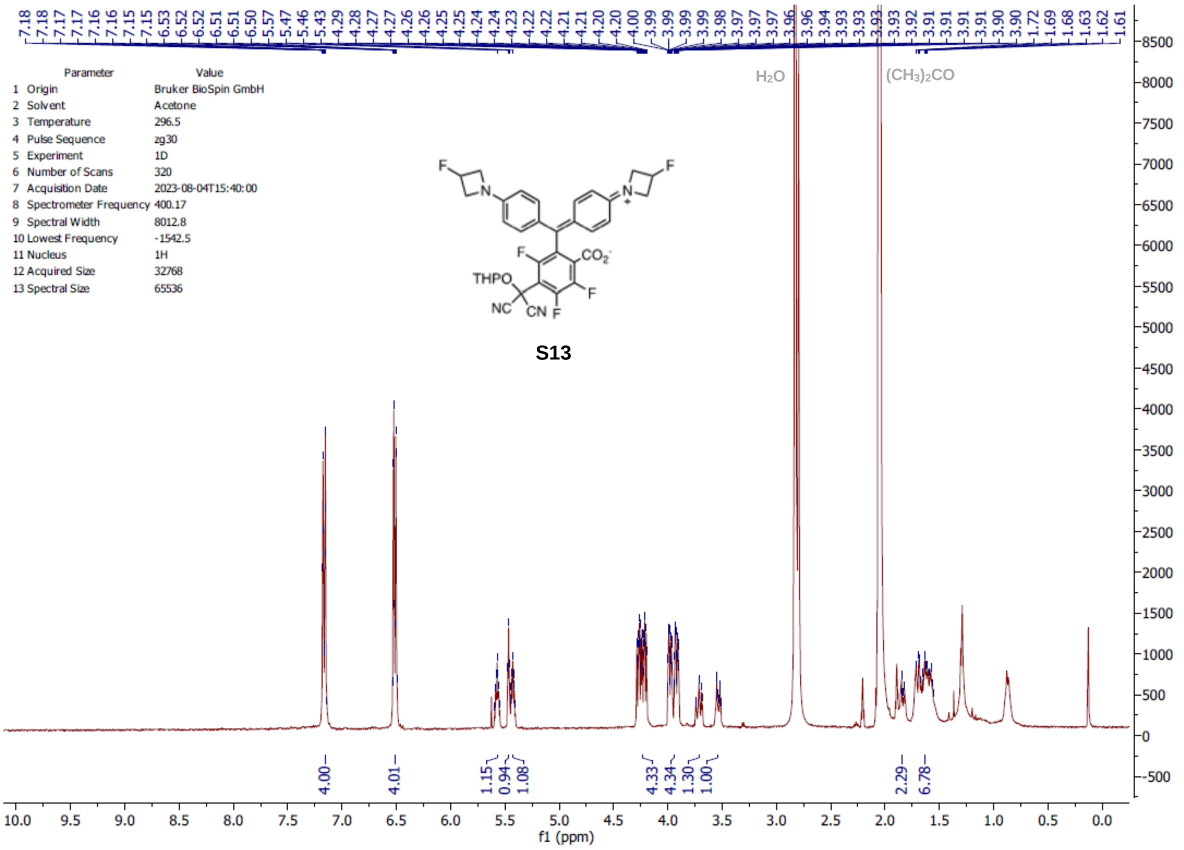

| Parameter | Value |
| --- | --- |
| 1 Origin | Bruker BioSpin GmbH |
| 2 Solvent | Acetone |
| 3 Temperature | 296.5 |
| 4 Pulse Sequence | zg30 |
| 5 Experiment | 1D |
| 6 Number of Scans | 320 |
| 7 Acquisition Date | 2023-08-04T15:40:00 |
| 8 Spectrometer Frequency | 400.17 |
| 9 Spectral Width | 8012.8 |
| 10 Lowest Frequency | -1542.5 |
| 11 Nucleus | 1H |
| 12 Acquired Size | 32768 |
| 13 Spectral Size | 65536 |

S13

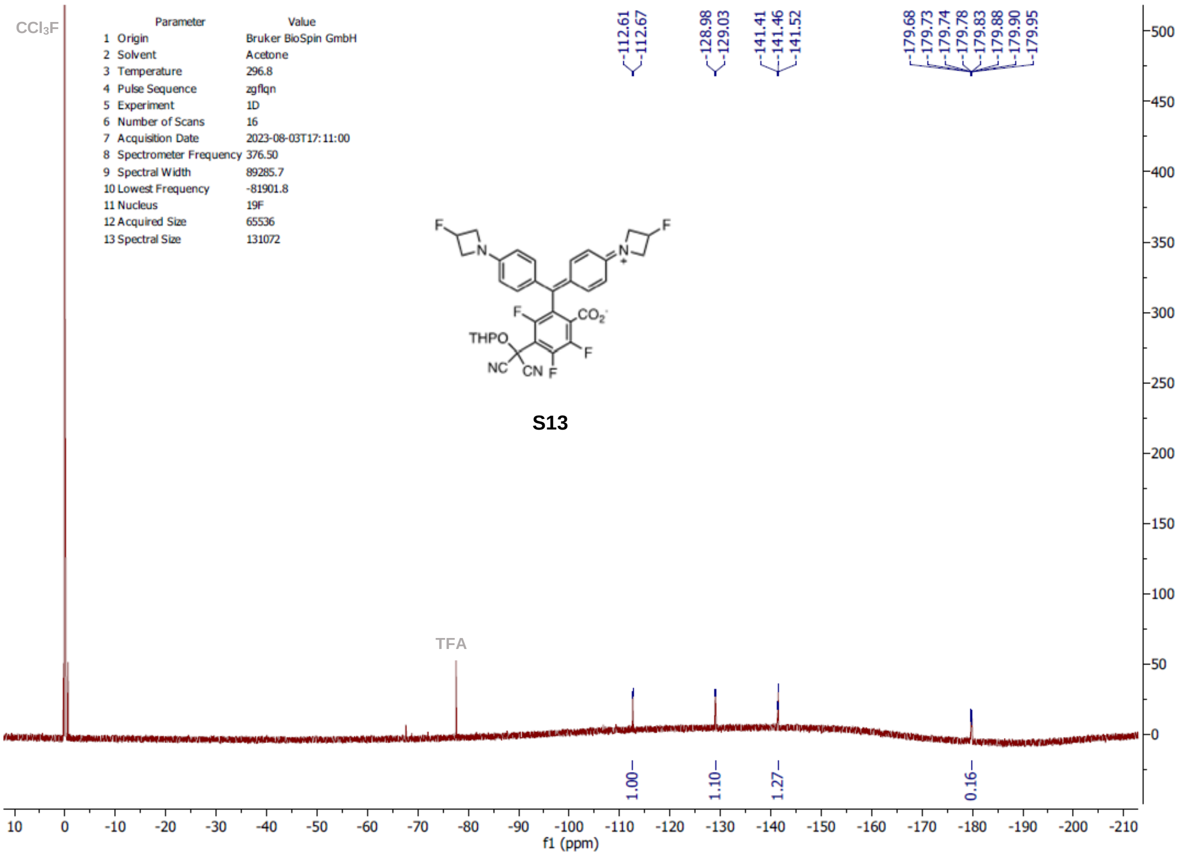

| Parameter | Value |
| --- | --- |
| 1 Origin | Bruker BioSpin GmbH |
| 2 Solvent | Acetone |
| 3 Temperature | 296.8 |
| 4 Pulse Sequence | zgflqn |
| 5 Experiment | 1D |
| 6 Number of Scans | 16 |
| 7 Acquisition Date | 2023-08-03T17:11:00 |
| 8 Spectrometer Frequency | 376.50 |
| 9 Spectral Width | 89285.7 |
| 10 Lowest Frequency | -81901.8 |
| 11 Nucleus | 19F |
| 12 Acquired Size | 65536 |
| 13 Spectral Size | 131072 |

S13

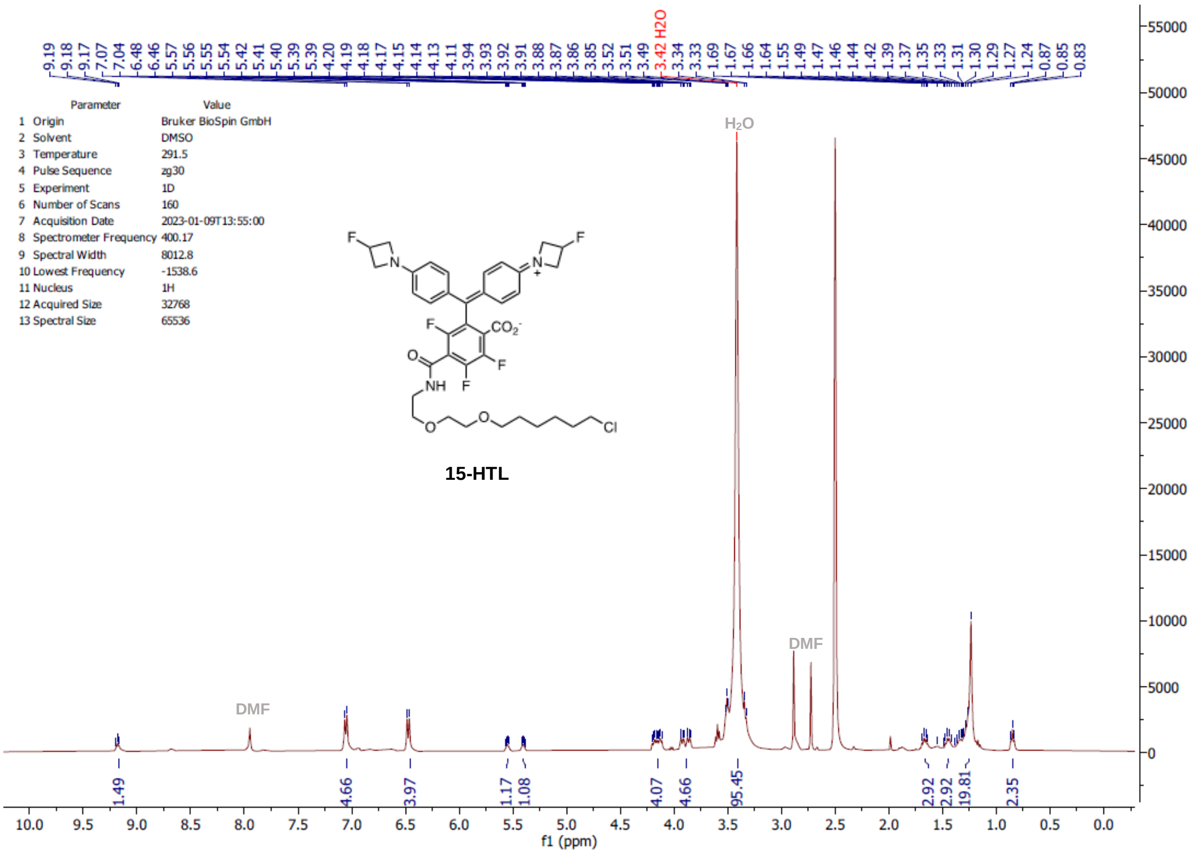
9.19 9.18 9.17 7.07 7.04 6.48 6.46 5.57 5.56 5.55 5.54 5.42 5.41 5.40 5.39 5.39 4.20 4.19 4.18 4.17 4.15 4.14 4.13 4.11 3.94 3.93 3.92 3.91 3.88 3.87 3.86 3.85 3.52 3.51 3.49 3.42 H2O 3.34 3.33 1.69 1.67 1.66 1.64 1.55 1.49 1.47 1.46 1.44 1.42 1.39 1.37 1.35 1.33 1.31 1.30 1.29 1.27 1.24 0.87 0.85 0.83
Parameter Value
1 Origin Bruker BioSpin GmbH
2 Solvent DMSO
3 Temperature 291.5
4 Pulse Sequence zg30
5 Experiment 1D
6 Number of Scans 160
7 Acquisition Date 2023-01-09T13:55:00
8 Spectrometer Frequency 400.17
9 Spectral Width 8012.8
10 Lowest Frequency -1538.6
11 Nucleus 1H
12 Acquired Size 32768
13 Spectral Size 65536
H2O
DMF
DMF
15-HTL
1.49 4.66 3.97 1.17 1.08 4.07 4.66 95.45 2.92 2.92 19.81 2.35
f1 (ppm)

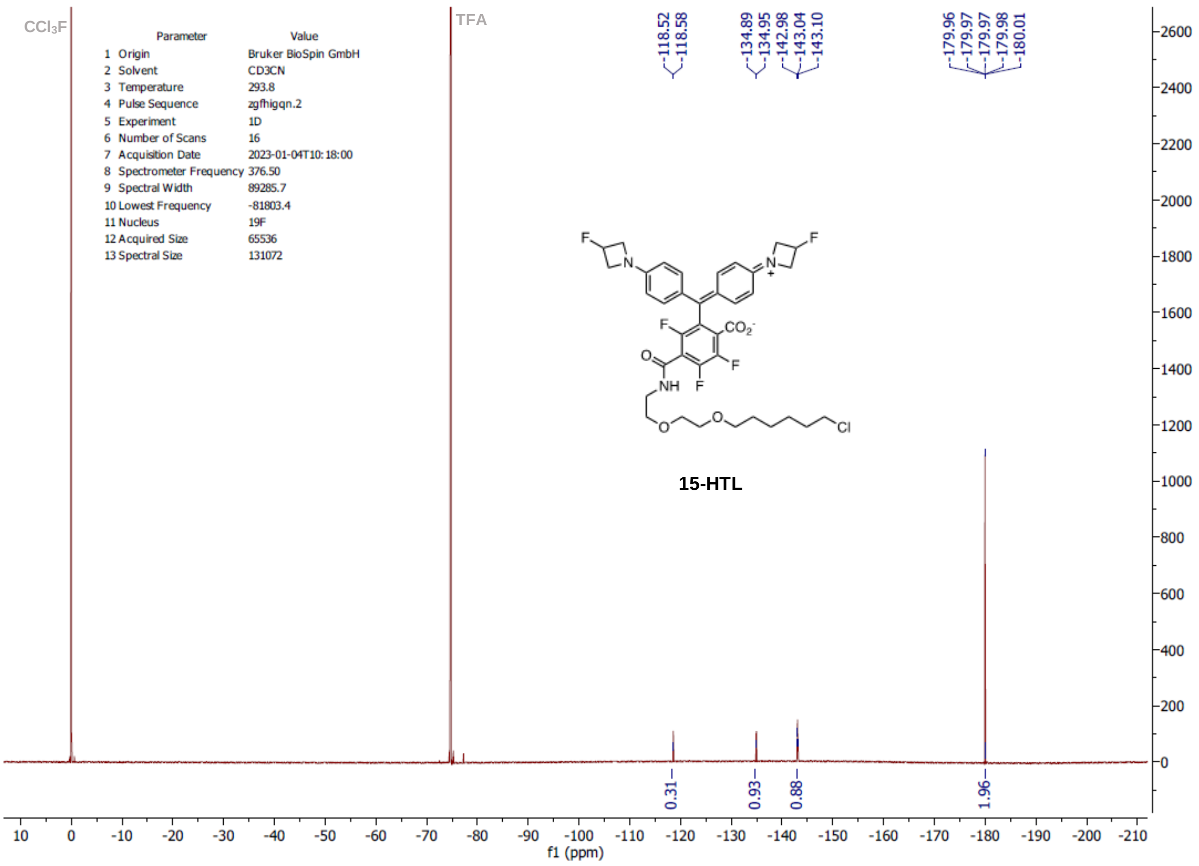
CCl3F
TFA
-118.52 -118.58 -134.89 -134.95 -142.98 -143.04 -143.10 -179.96 -179.97 -179.97 -179.98 -180.01
Parameter Value
1 Origin Bruker BioSpin GmbH
2 Solvent CD3CN
3 Temperature 293.8
4 Pulse Sequence zgfhigqn.2
5 Experiment 1D
6 Number of Scans 16
7 Acquisition Date 2023-01-04T10:18:00
8 Spectrometer Frequency 376.50
9 Spectral Width 89285.7
10 Lowest Frequency -81803.4
11 Nucleus 19F
12 Acquired Size 65536
13 Spectral Size 131072
15-HTL
0.31 0.93 0.88 1.96
f1 (ppm)

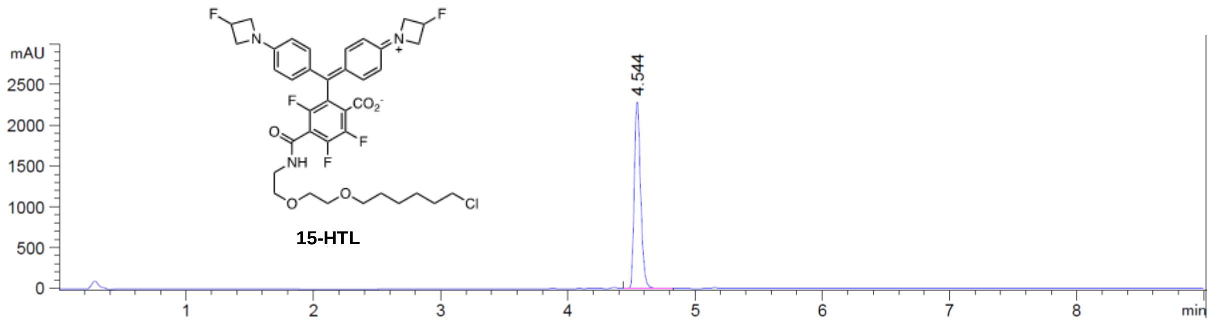

15-HTL

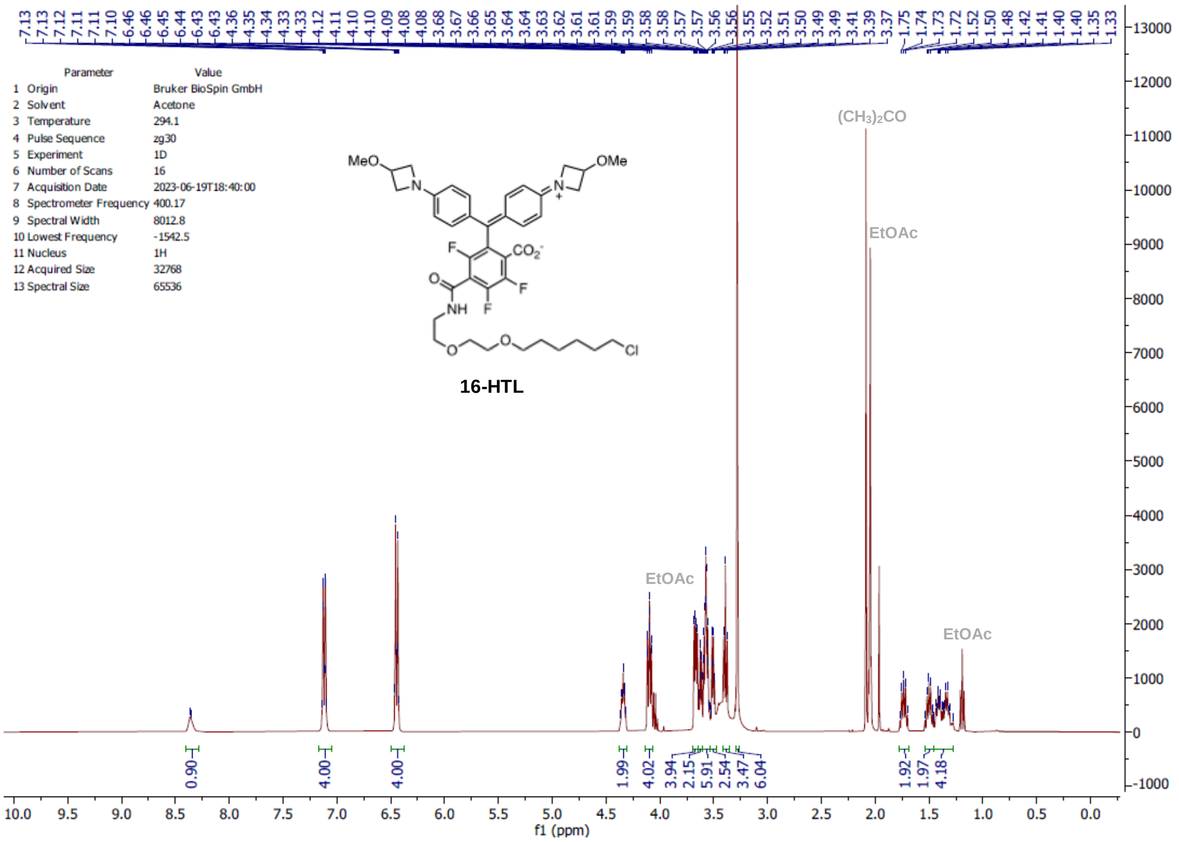
Parameter Value
1 Origin Bruker BioSpin GmbH
2 Solvent Acetone
3 Temperature 294.1
4 Pulse Sequence zg30
5 Experiment 1D
6 Number of Scans 16
7 Acquisition Date 2023-06-19T18:40:00
8 Spectrometer Frequency 400.17
9 Spectral Width 8012.8
10 Lowest Frequency -1542.5
11 Nucleus 1H
12 Acquired Size 32768
13 Spectral Size 65536
16-HTL
(CH3)2CO
EtOAc
f1 (ppm)

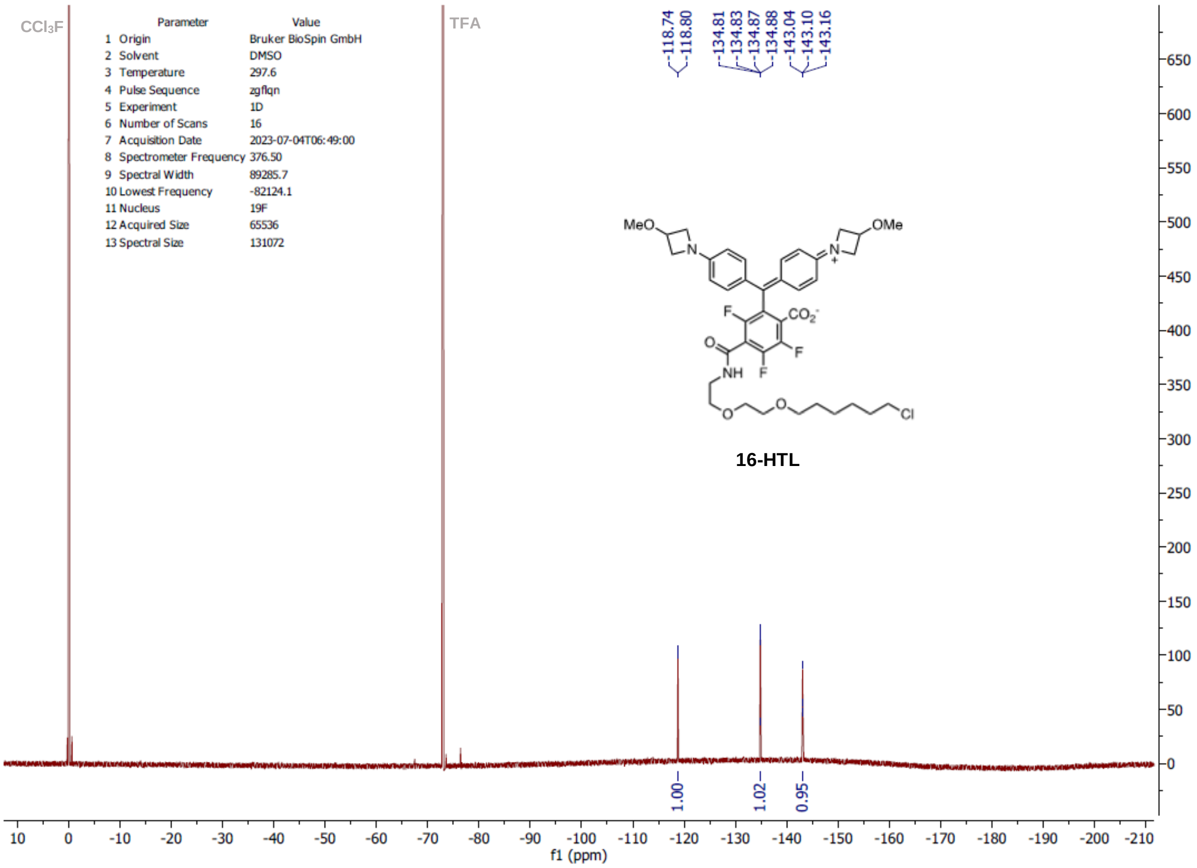
CCl3F
TFA
Parameter Value
1 Origin Bruker BioSpin GmbH
2 Solvent DMSO
3 Temperature 297.6
4 Pulse Sequence zgflqn
5 Experiment 1D
6 Number of Scans 16
7 Acquisition Date 2023-07-04T06:49:00
8 Spectrometer Frequency 376.50
9 Spectral Width 89285.7
10 Lowest Frequency -82124.1
11 Nucleus 19F
12 Acquired Size 65536
13 Spectral Size 131072
-118.74
-118.80
-134.81
-134.83
-134.87
-134.88
-143.04
-143.10
-143.16
16-HTL
1.00
1.02
0.95
f1 (ppm)

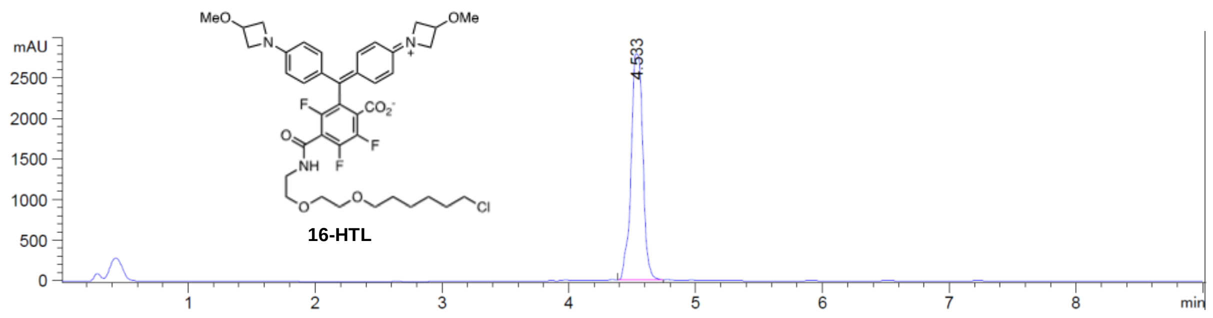
16-HTL
mAU
2500
2000
1500
1000
500
0
4.533
1
2
3
4
5
6
7
8
min

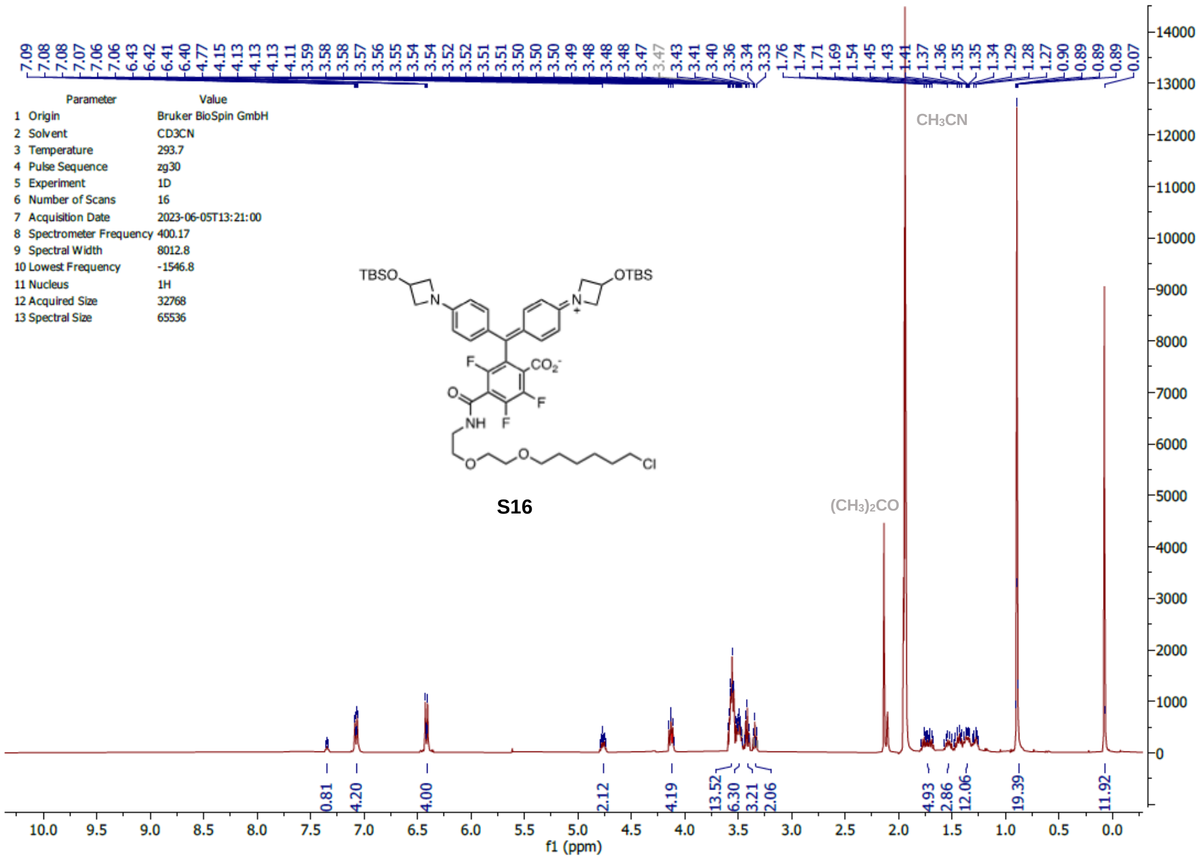
7.09 7.08 7.08 7.07 7.06 7.06 6.43 6.42 6.41 6.40 4.77 4.15 4.13 4.13 4.13 4.11 3.59 3.58 3.58 3.57 3.56 3.55 3.54 3.54 3.52 3.52 3.51 3.51 3.50 3.50 3.50 3.49 3.48 3.48 3.48 3.47 3.47 3.43 3.41 3.40 3.36 3.34 3.33 1.76 1.74 1.71 1.69 1.54 1.45 1.43 1.41 1.37 1.36 1.35 1.35 1.34 1.29 1.28 1.27 0.90 0.89 0.89 0.89 0.07
Parameter Value
1 Origin Bruker BioSpin GmbH
2 Solvent CD3CN
3 Temperature 293.7
4 Pulse Sequence zg30
5 Experiment 1D
6 Number of Scans 16
7 Acquisition Date 2023-06-05T13:21:00
8 Spectrometer Frequency 400.17
9 Spectral Width 8012.8
10 Lowest Frequency -1546.8
11 Nucleus 1H
12 Acquired Size 32768
13 Spectral Size 65536
CH3CN
(CH3)2CO
TBSO
OTBS
N
F
CO2-
O
NH
Cl
S16
0.81 4.20 4.00 2.12 4.19 13.52 6.30 3.21 2.06 4.93 2.86 12.06 19.39 11.92
f1 (ppm)

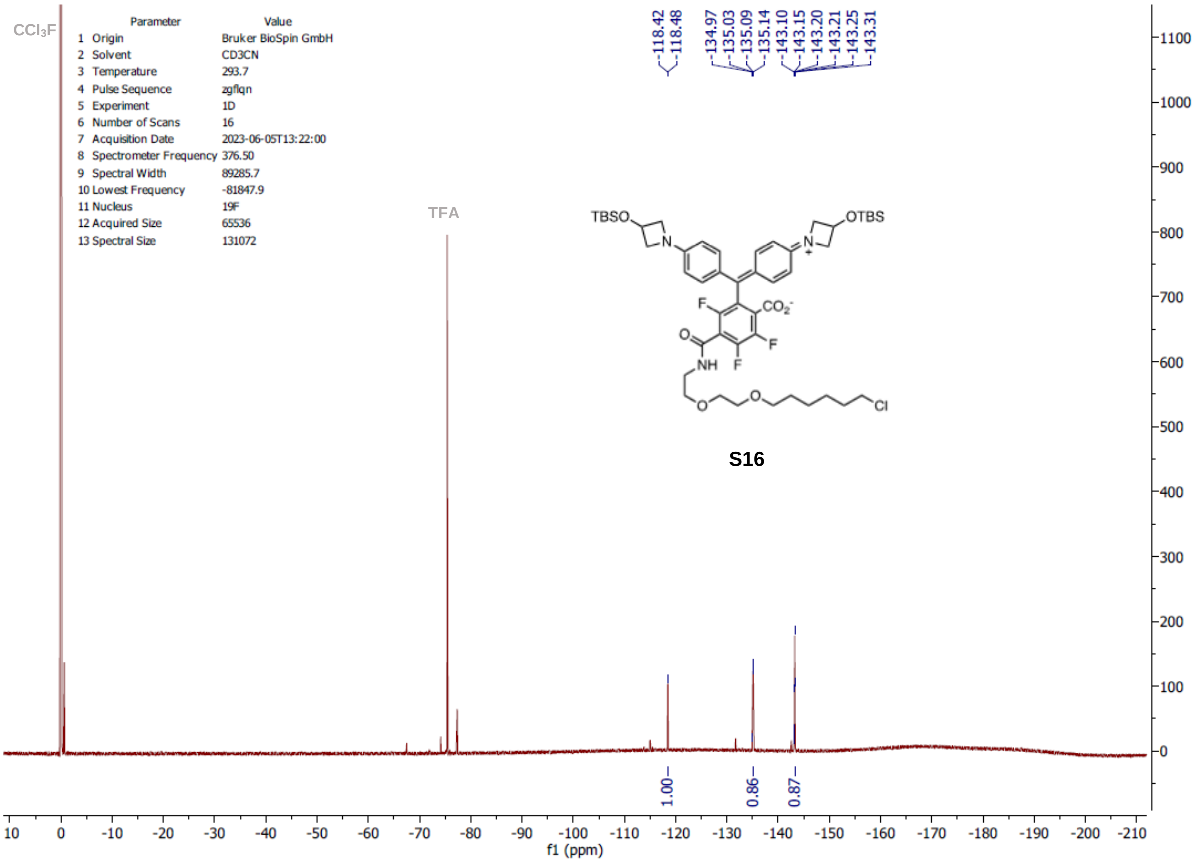
-118.42 -118.48 -134.97 -135.03 -135.09 -135.14 -143.10 -143.15 -143.20 -143.21 -143.25 -143.31
CCl3F
Parameter Value
1 Origin Bruker BioSpin GmbH
2 Solvent CD3CN
3 Temperature 293.7
4 Pulse Sequence zgflqn
5 Experiment 1D
6 Number of Scans 16
7 Acquisition Date 2023-06-05T13:22:00
8 Spectrometer Frequency 376.50
9 Spectral Width 89285.7
10 Lowest Frequency -81847.9
11 Nucleus 19F
12 Acquired Size 65536
13 Spectral Size 131072
TFA
TBSO
OTBS
N
F
CO2-
O
NH
Cl
S16
1.00 0.86 0.87
f1 (ppm)

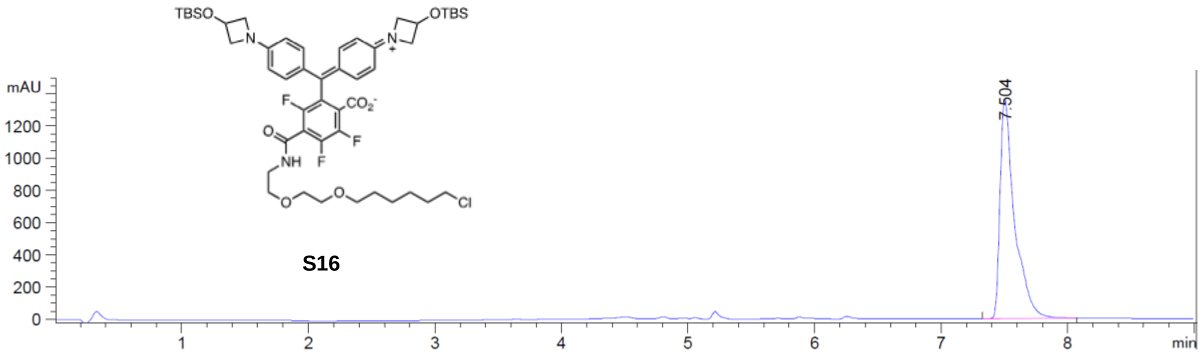
TBSO
OTBS
N
N
+
F
$CO_2^-$
O
NH
F
F
O
O
Cl
S16
mAU
1200
1000
800
600
400
200
0
1
2
3
4
5
6
7
8
min
7.504

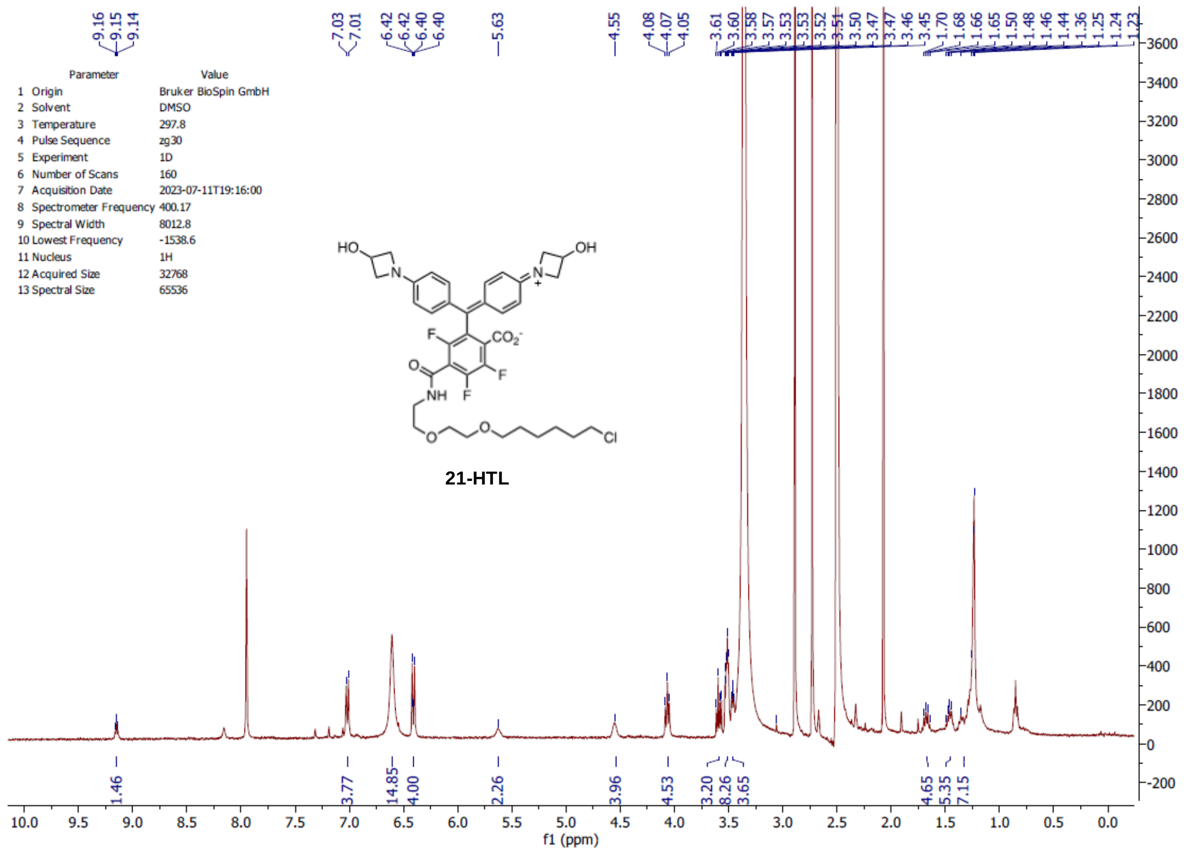
Parameter Value
1 Origin Bruker BioSpin GmbH
2 Solvent DMSO
3 Temperature 297.8
4 Pulse Sequence zg30
5 Experiment 1D
6 Number of Scans 160
7 Acquisition Date 2023-07-11T19:16:00
8 Spectrometer Frequency 400.17
9 Spectral Width 8012.8
10 Lowest Frequency -1538.6
11 Nucleus 1H
12 Acquired Size 32768
13 Spectral Size 65536
21-HTL
f1 (ppm)

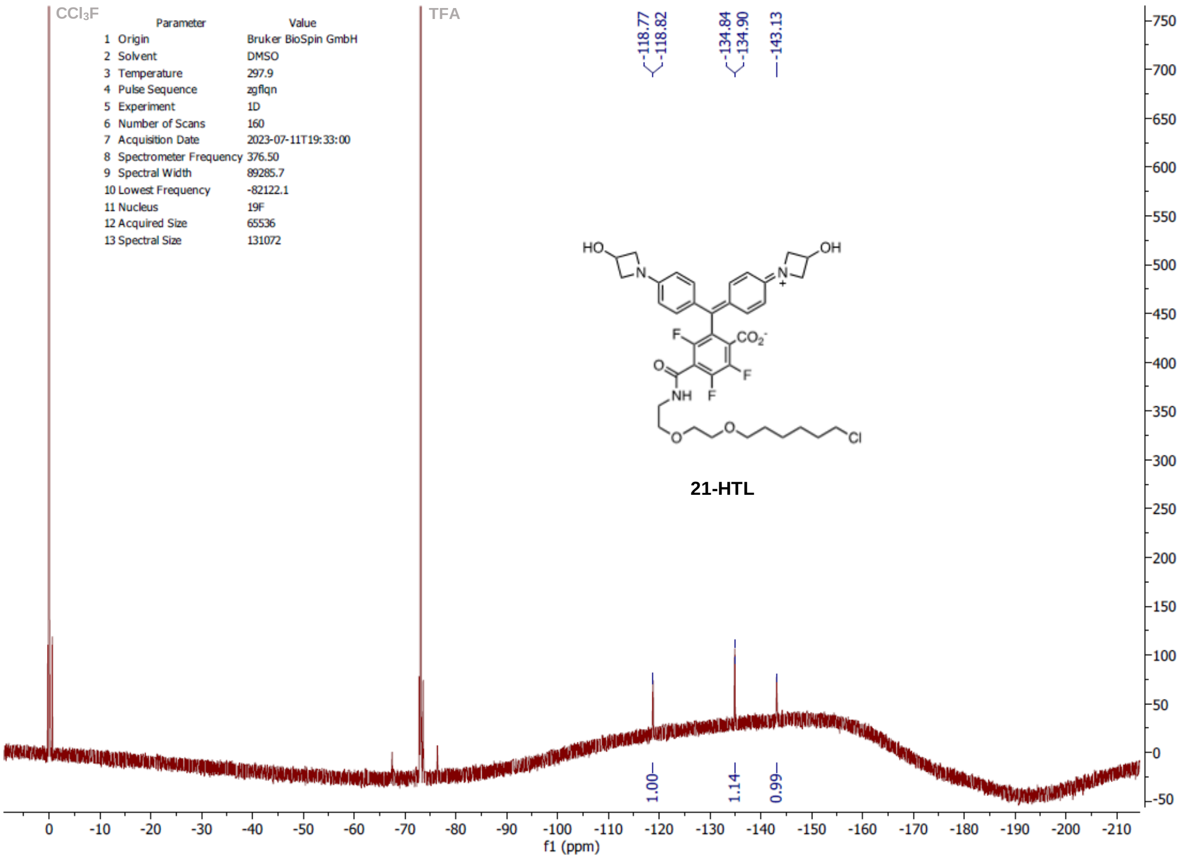
CCl3F
TFA
Parameter Value
1 Origin Bruker BioSpin GmbH
2 Solvent DMSO
3 Temperature 297.9
4 Pulse Sequence zgflqn
5 Experiment 1D
6 Number of Scans 160
7 Acquisition Date 2023-07-11T19:33:00
8 Spectrometer Frequency 376.50
9 Spectral Width 89285.7
10 Lowest Frequency -82122.1
11 Nucleus 19F
12 Acquired Size 65536
13 Spectral Size 131072
-118.77
-118.82
-134.84
-134.90
-143.13
1.00
1.14
0.99
21-HTL
f1 (ppm)

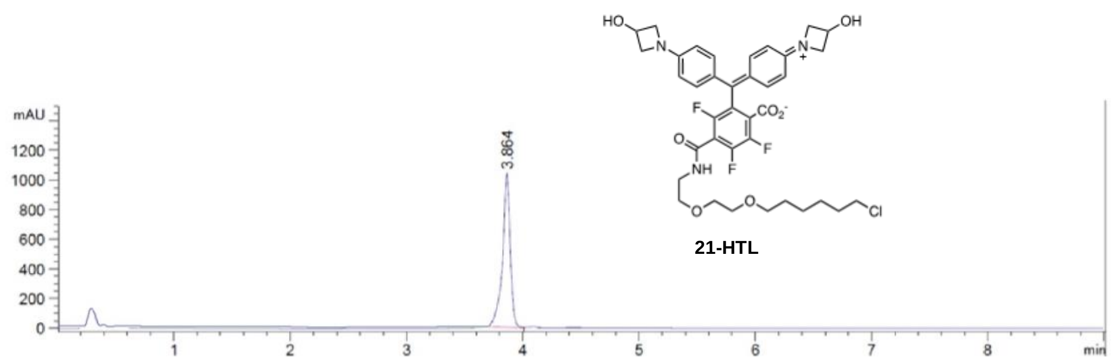

mAU
1200
1000
800
600
400
200
0
3.864
1
2
3
4
5
6
7
8
min
HO
N
OH
F
$CO_2^-$
O
F
NH
F
O
O
Cl
21-HTL

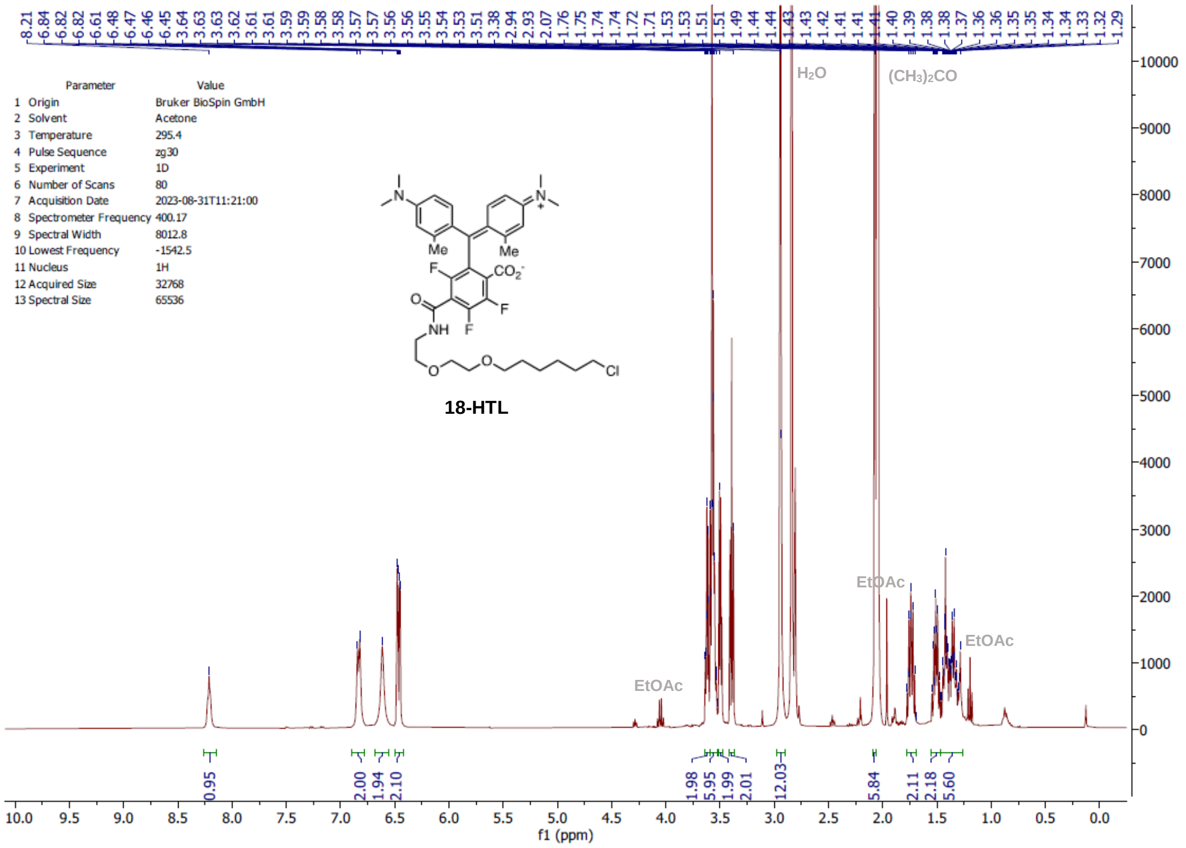
8.21 6.84 6.82 6.82 6.61 6.48 6.47 6.46 6.45 3.64 3.63 3.63 3.62 3.61 3.61 3.59 3.59 3.58 3.58 3.57 3.57 3.56 3.56 3.55 3.54 3.53 3.51 3.38 2.94 2.93 2.07 1.76 1.75 1.74 1.74 1.72 1.71 1.53 1.53 1.51 1.51 1.49 1.44 1.44 1.43 1.43 1.42 1.41 1.41 1.41 1.40 1.39 1.38 1.38 1.37 1.36 1.36 1.35 1.35 1.34 1.34 1.33 1.32 1.29
Parameter Value
1 Origin Bruker BioSpin GmbH
2 Solvent Acetone
3 Temperature 295.4
4 Pulse Sequence zg30
5 Experiment 1D
6 Number of Scans 80
7 Acquisition Date 2023-08-31T11:21:00
8 Spectrometer Frequency 400.17
9 Spectral Width 8012.8
10 Lowest Frequency -1542.5
11 Nucleus 1H
12 Acquired Size 32768
13 Spectral Size 65536
18-HTL
H2O
(CH3)2CO
EtOAc
EtOAc
EtOAc
0.95 2.00 1.94 2.10 1.98 5.95 1.99 2.01 12.03 5.84 2.11 2.18 5.60
f1 (ppm)

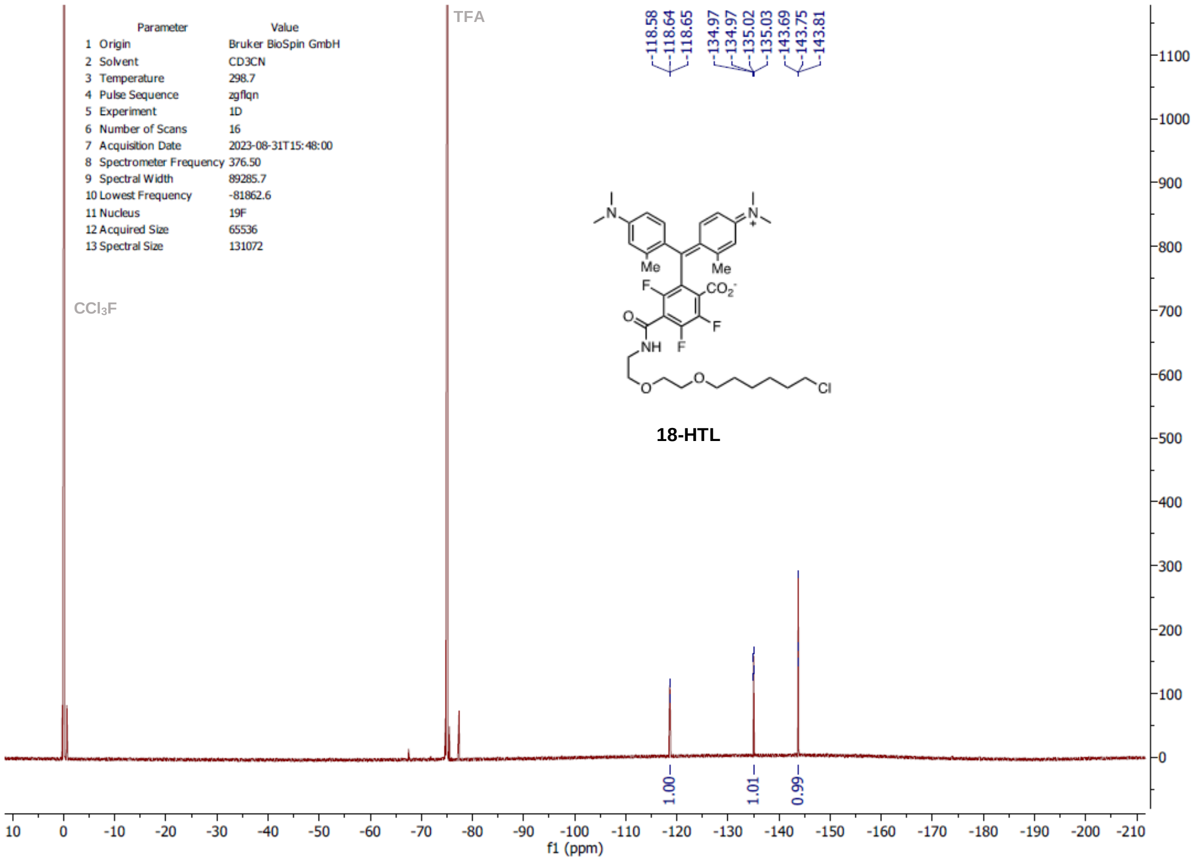
-118.58 -118.64 -118.65 -134.97 -134.97 -135.02 -135.03 -143.69 -143.75 -143.81
Parameter Value
1 Origin Bruker BioSpin GmbH
2 Solvent CD3CN
3 Temperature 298.7
4 Pulse Sequence zgflqn
5 Experiment 1D
6 Number of Scans 16
7 Acquisition Date 2023-08-31T15:48:00
8 Spectrometer Frequency 376.50
9 Spectral Width 89285.7
10 Lowest Frequency -81862.6
11 Nucleus 19F
12 Acquired Size 65536
13 Spectral Size 131072
TFA
CCl3F
18-HTL
1.00 1.01 0.99
f1 (ppm)

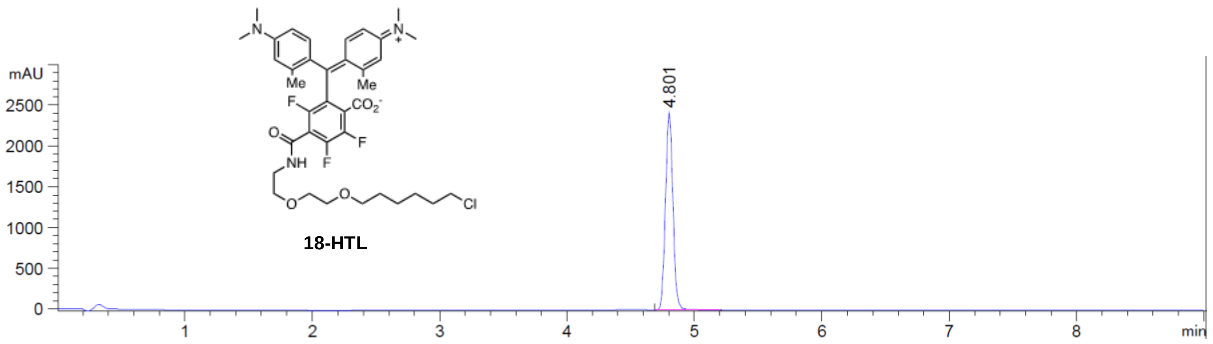
18-HTL
mAU
2500
2000
1500
1000
500
0
4.801
1
2
3
4
5
6
7
8
min

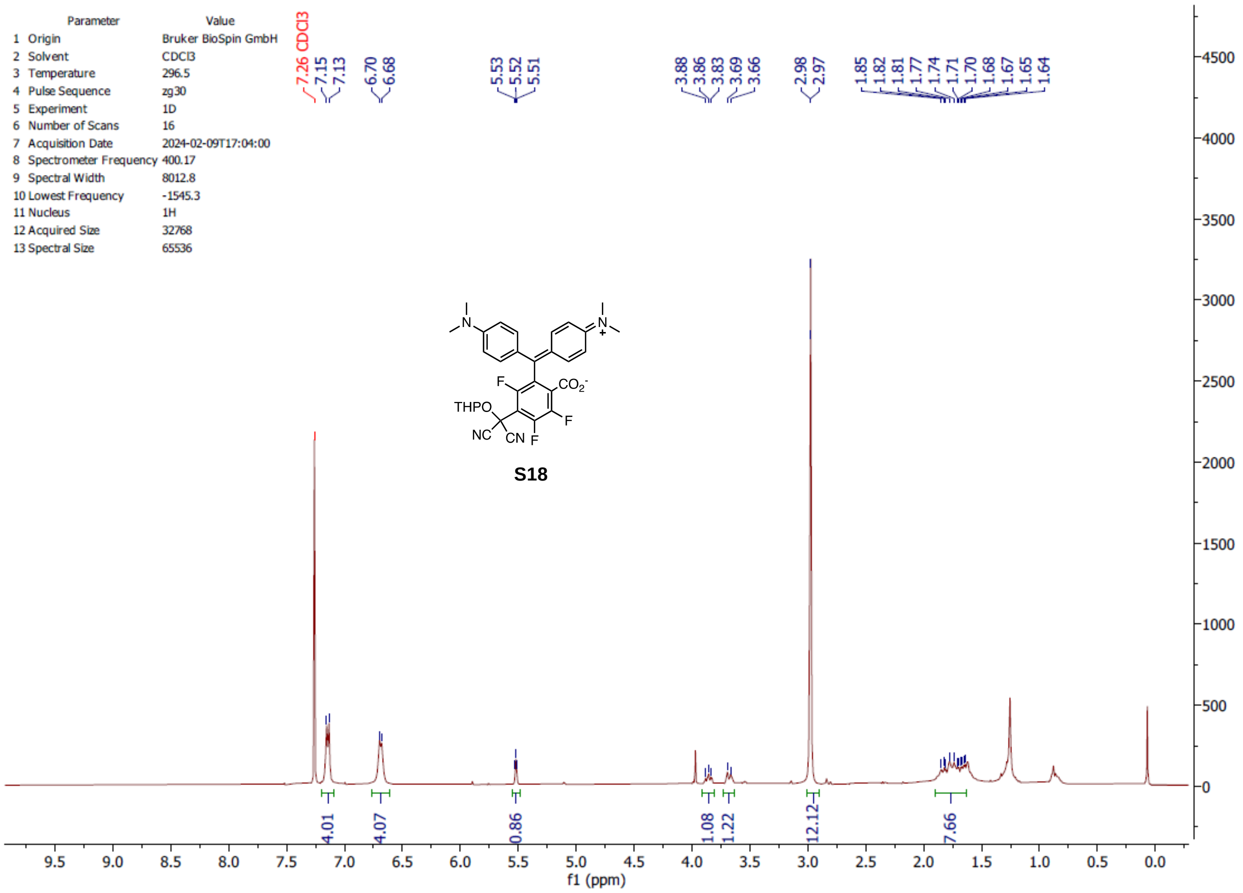
Parameter Value
1 Origin Bruker BioSpin GmbH
2 Solvent CDCl3
3 Temperature 296.5
4 Pulse Sequence zg30
5 Experiment 1D
6 Number of Scans 16
7 Acquisition Date 2024-02-09T17:04:00
8 Spectrometer Frequency 400.17
9 Spectral Width 8012.8
10 Lowest Frequency -1545.3
11 Nucleus 1H
12 Acquired Size 32768
13 Spectral Size 65536
7.26 CDCl3
7.15
7.13
6.70
6.68
5.53
5.52
5.51
3.88
3.86
3.83
3.69
3.66
2.98
2.97
1.85
1.82
1.81
1.77
1.74
1.71
1.70
1.68
1.67
1.65
1.64
THPO
NC
CN
F
CO2-
N
N+
S18
4.01
4.07
0.86
1.08
1.22
12.12
7.66
f1 (ppm)

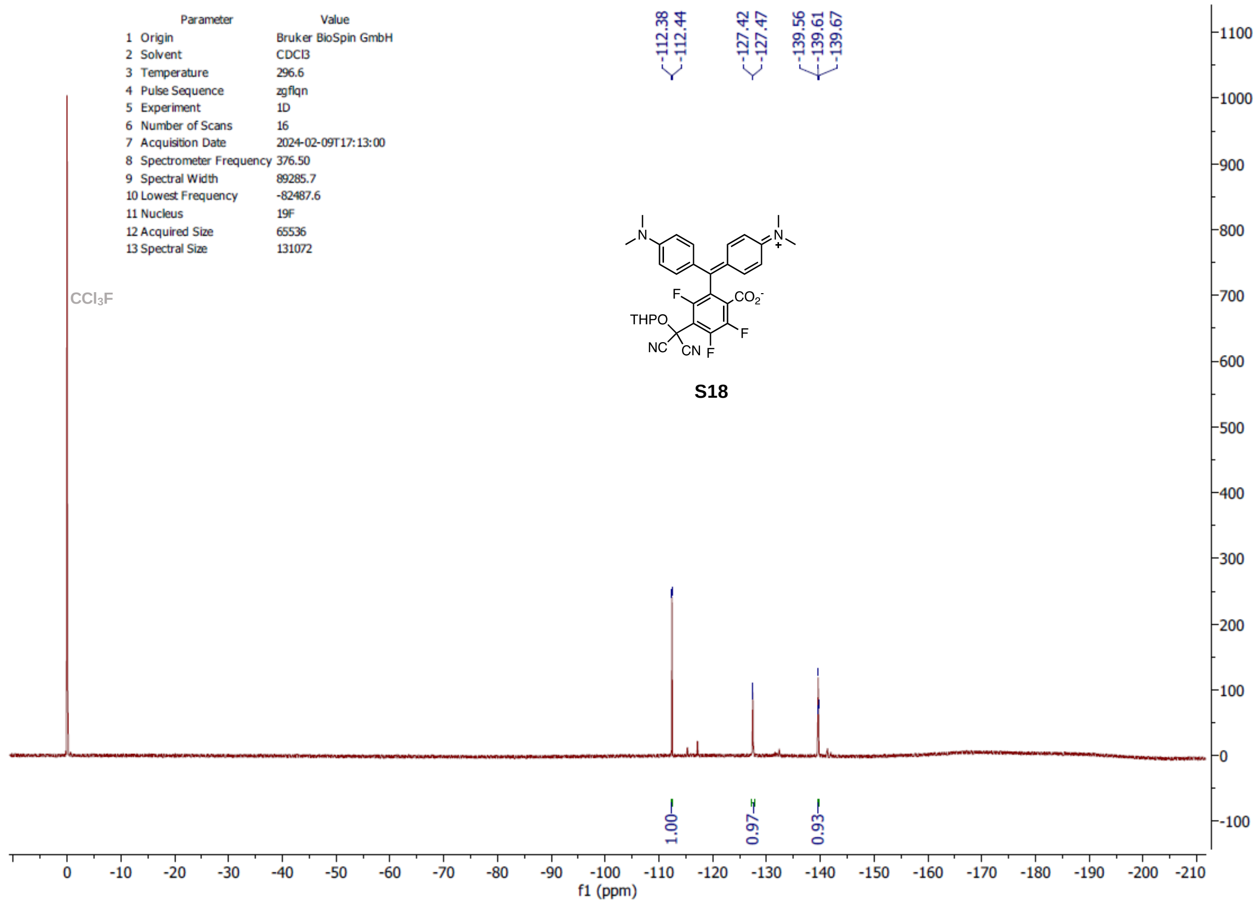
Parameter Value
1 Origin Bruker BioSpin GmbH
2 Solvent CDCl3
3 Temperature 296.6
4 Pulse Sequence zgflqn
5 Experiment 1D
6 Number of Scans 16
7 Acquisition Date 2024-02-09T17:13:00
8 Spectrometer Frequency 376.50
9 Spectral Width 89285.7
10 Lowest Frequency -82487.6
11 Nucleus 19F
12 Acquired Size 65536
13 Spectral Size 131072
-112.38
-112.44
-127.42
-127.47
-139.56
-139.61
-139.67
CCl3F
THPO
NC
CN
F
CO2-
N
N+
S18
1.00
0.97
0.93
f1 (ppm)

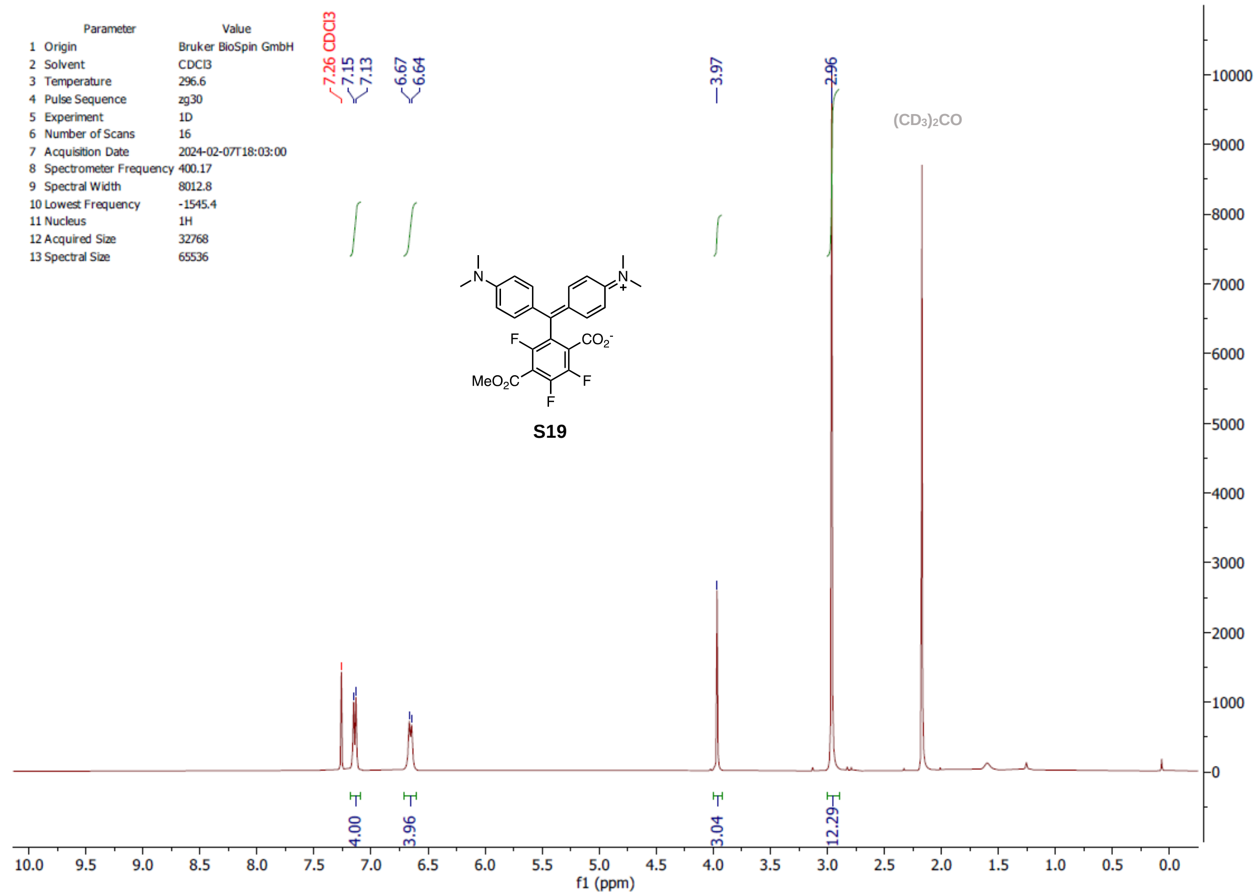
Parameter Value
1 Origin Bruker BioSpin GmbH
2 Solvent CDCl3
3 Temperature 296.6
4 Pulse Sequence zg30
5 Experiment 1D
6 Number of Scans 16
7 Acquisition Date 2024-02-07T18:03:00
8 Spectrometer Frequency 400.17
9 Spectral Width 8012.8
10 Lowest Frequency -1545.4
11 Nucleus 1H
12 Acquired Size 32768
13 Spectral Size 65536
7.26 CDCl3
7.15
7.13
6.67
6.64
3.97
2.96
(CD3)2CO
MeO2C
CO2-
F
S19
4.00
3.96
3.04
12.29
f1 (ppm)

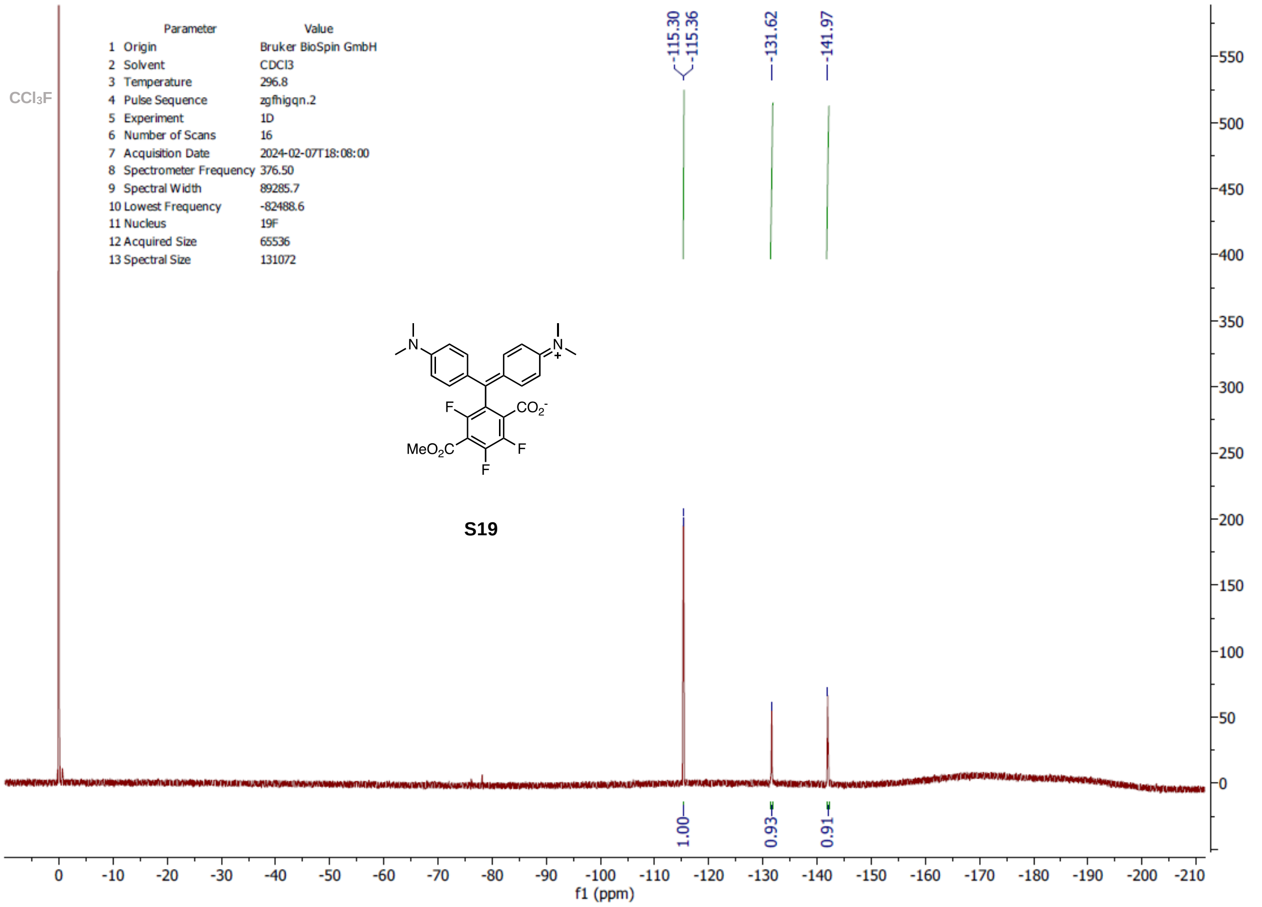
Parameter Value
1 Origin Bruker BioSpin GmbH
2 Solvent CDCl3
3 Temperature 296.8
4 Pulse Sequence zgfhigqn.2
5 Experiment 1D
6 Number of Scans 16
7 Acquisition Date 2024-02-07T18:08:00
8 Spectrometer Frequency 376.50
9 Spectral Width 89285.7
10 Lowest Frequency -82488.6
11 Nucleus 19F
12 Acquired Size 65536
13 Spectral Size 131072
CCl3F
-115.30
-115.36
-131.62
-141.97
MeO2C
CO2-
F
S19
1.00
0.93
0.91
f1 (ppm)

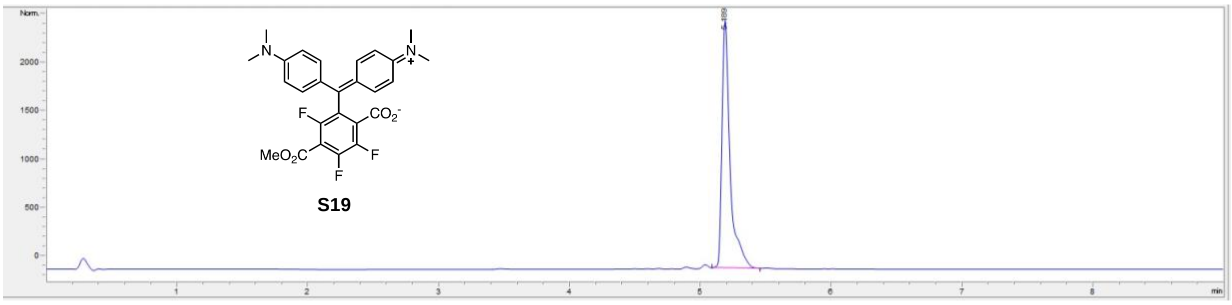
S19

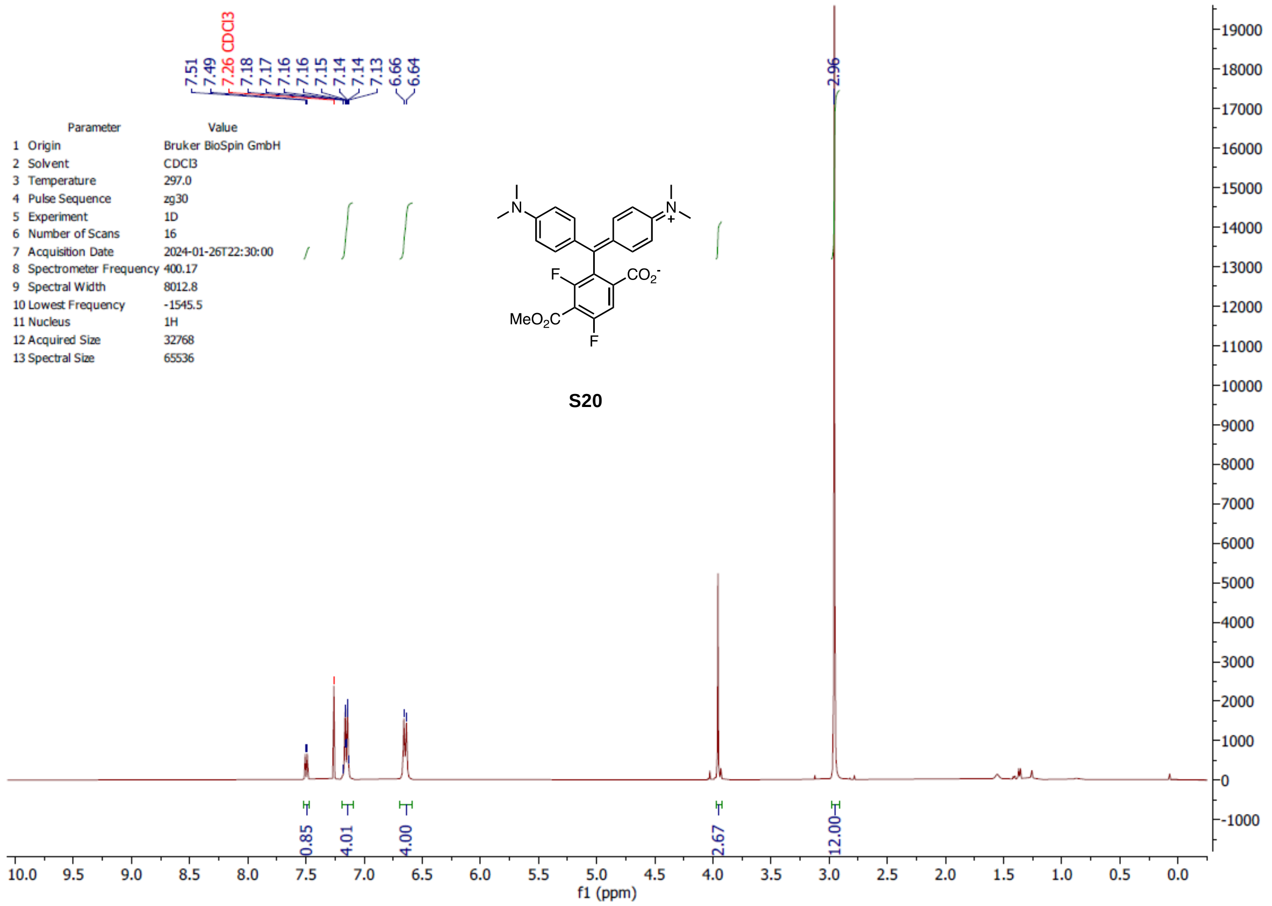
7.51 7.49 7.26 CDCl3 7.18 7.17 7.16 7.16 7.15 7.14 7.14 7.13 6.66 6.64
2.96
Parameter Value
1 Origin Bruker BioSpin GmbH
2 Solvent CDCl3
3 Temperature 297.0
4 Pulse Sequence zg30
5 Experiment 1D
6 Number of Scans 16
7 Acquisition Date 2024-01-26T22:30:00
8 Spectrometer Frequency 400.17
9 Spectral Width 8012.8
10 Lowest Frequency -1545.5
11 Nucleus 1H
12 Acquired Size 32768
13 Spectral Size 65536
F
$CO_2^-$
$MeO_2C$
F
S20
0.85 4.01 4.00 2.67 12.00
f1 (ppm)

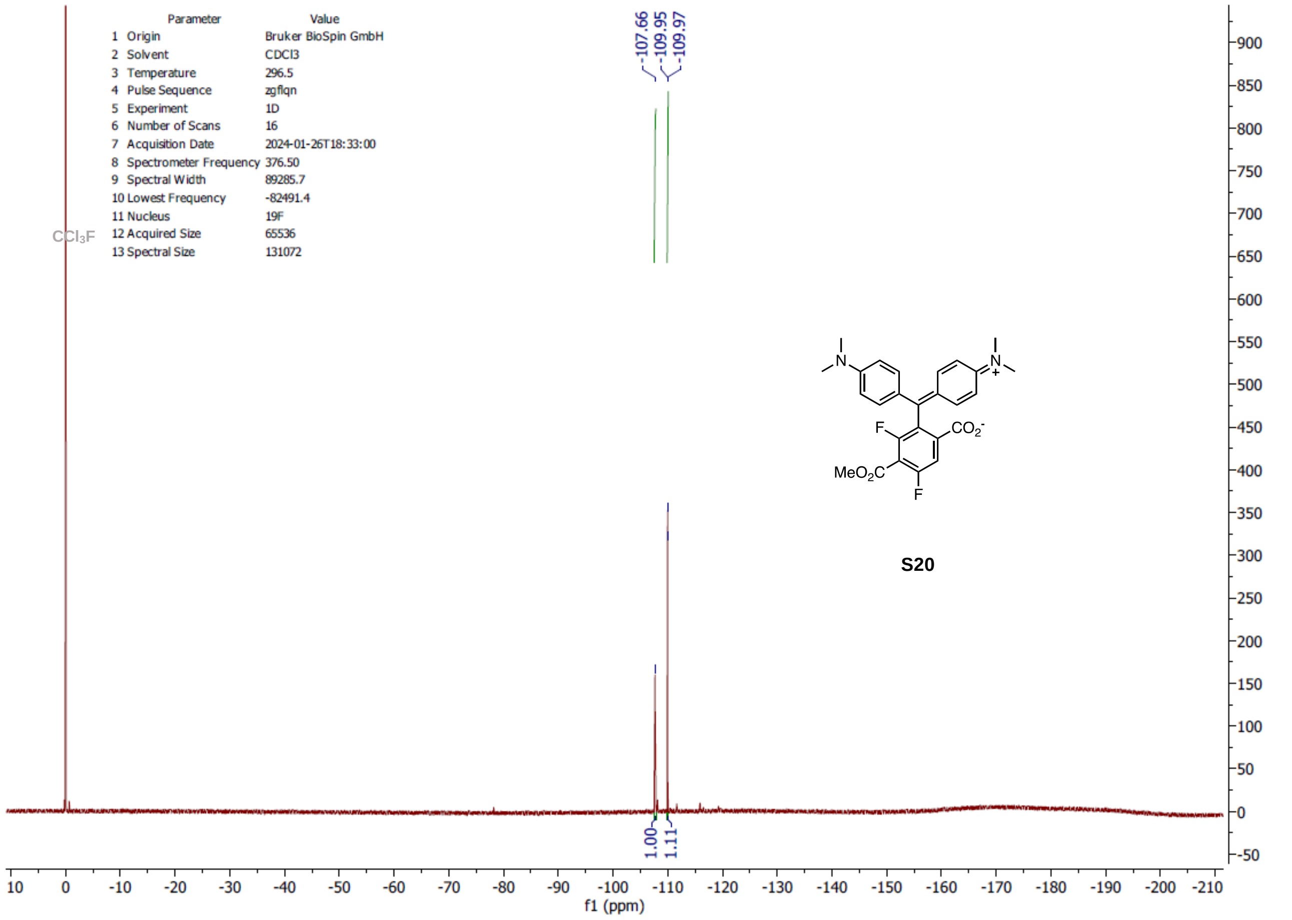
Parameter Value
1 Origin Bruker BioSpin GmbH
2 Solvent CDCl3
3 Temperature 296.5
4 Pulse Sequence zgflqn
5 Experiment 1D
6 Number of Scans 16
7 Acquisition Date 2024-01-26T18:33:00
8 Spectrometer Frequency 376.50
9 Spectral Width 89285.7
10 Lowest Frequency -82491.4
11 Nucleus 19F
12 Acquired Size 65536
13 Spectral Size 131072
$CCl_3F$
-107.66 -109.95 -109.97
F
$CO_2^-$
$MeO_2C$
F
S20
1.00 1.11
f1 (ppm)

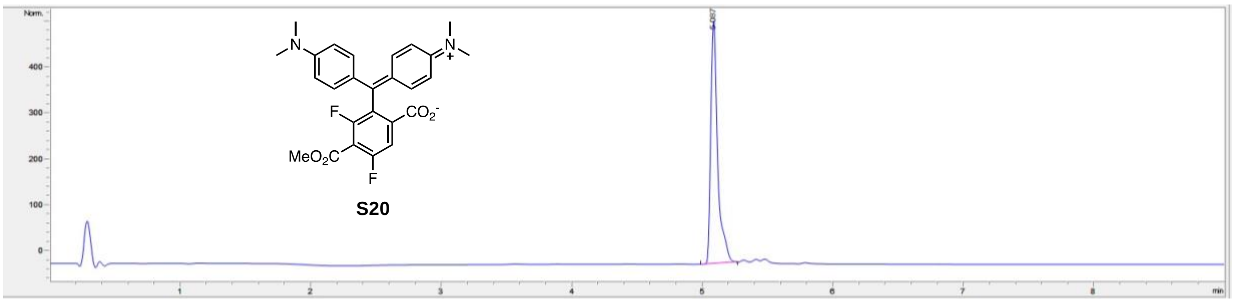

Norm.
400
300
200
100
0
1
2
3
4
5
6
7
8
min
F
$CO_2^-$
$MeO_2C$
F
S20

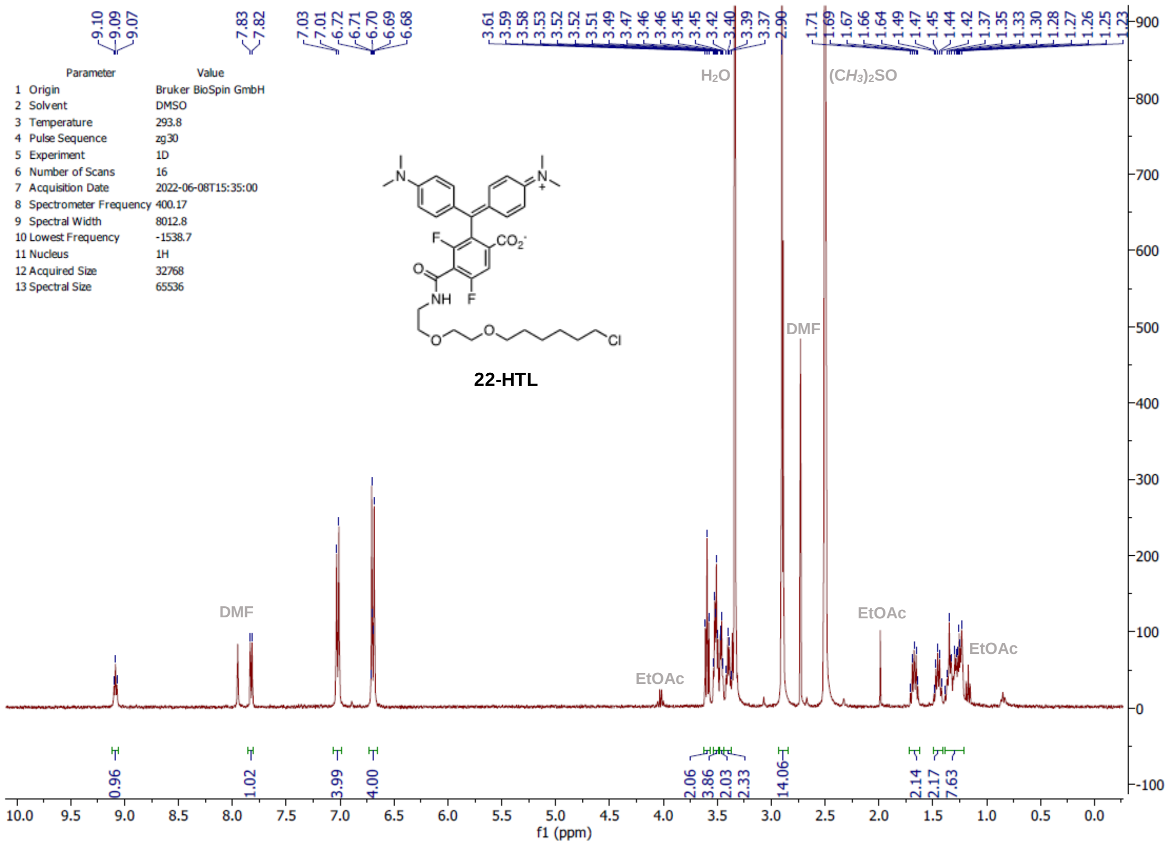
Parameter Value
1 Origin Bruker BioSpin GmbH
2 Solvent DMSO
3 Temperature 293.8
4 Pulse Sequence zg30
5 Experiment 1D
6 Number of Scans 16
7 Acquisition Date 2022-06-08T15:35:00
8 Spectrometer Frequency 400.17
9 Spectral Width 8012.8
10 Lowest Frequency -1538.7
11 Nucleus 1H
12 Acquired Size 32768
13 Spectral Size 65536
22-HTL
H2O
(CH3)2SO
DMF
EtOAc
f1 (ppm)

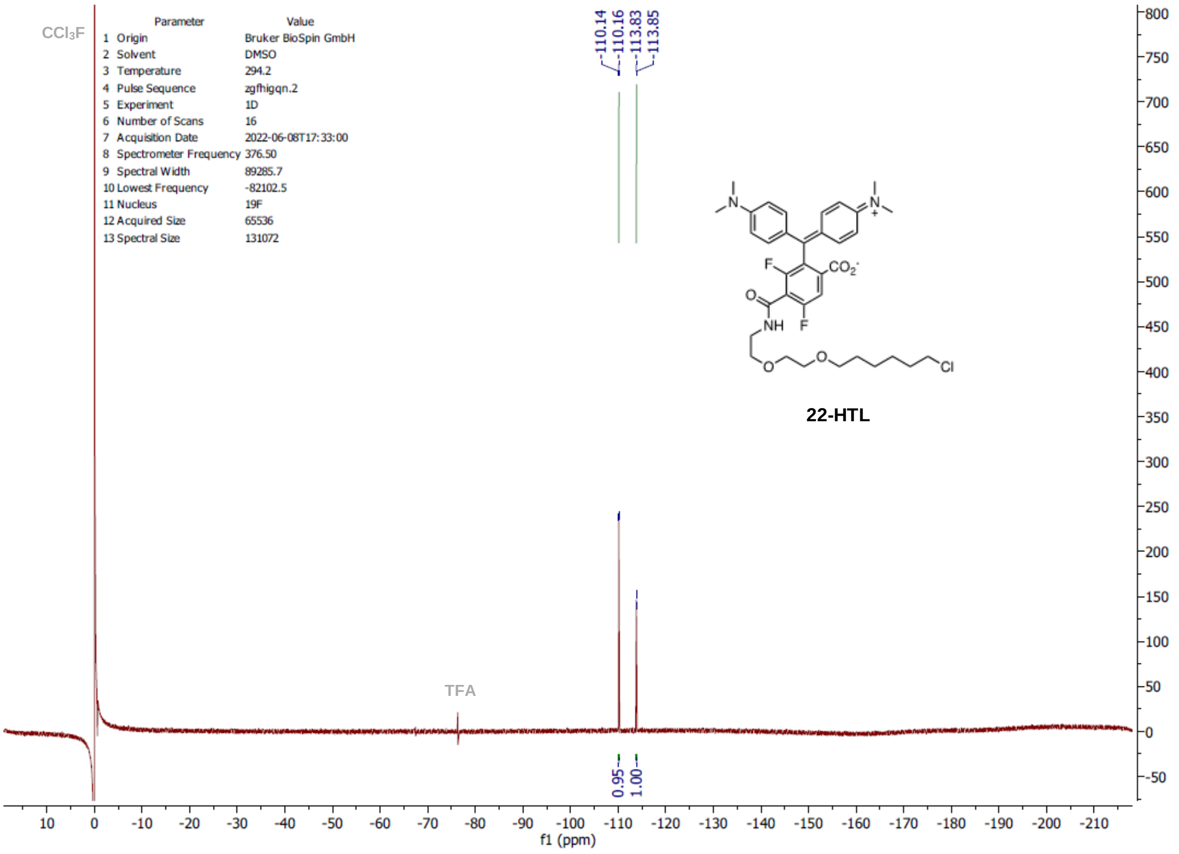
CCl3F
Parameter Value
1 Origin Bruker BioSpin GmbH
2 Solvent DMSO
3 Temperature 294.2
4 Pulse Sequence zgfhigqn.2
5 Experiment 1D
6 Number of Scans 16
7 Acquisition Date 2022-06-08T17:33:00
8 Spectrometer Frequency 376.50
9 Spectral Width 89285.7
10 Lowest Frequency -82102.5
11 Nucleus 19F
12 Acquired Size 65536
13 Spectral Size 131072
-110.14
-110.16
-113.83
-113.85
22-HTL
TFA
0.95
1.00
f1 (ppm)

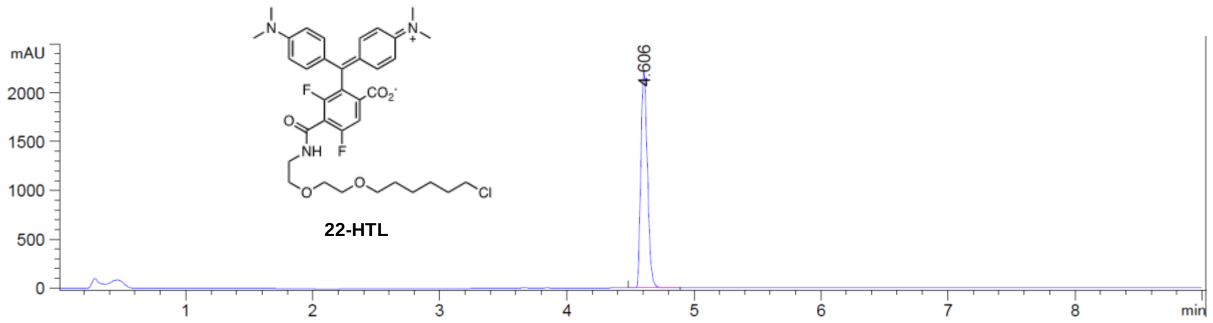

**22-HTL**
